# European Farmhouse Brewing Yeasts Form a Distinct Genetic Group

**DOI:** 10.1101/2024.04.04.588101

**Authors:** Richard Preiss, Eugene Fletcher, Lars Marius Garshol, Barret Foster, Emine Ozsahin, Mark Lubberts, George van der Merwe, Kristoffer Krogerus

## Abstract

The brewing industry is constantly evolving, driven by the quest for novel flavors and fermentation characteristics that cater to evolving consumer preferences. This study explores the genetic and phenotypic diversity of European farmhouse yeasts, traditionally used in rural brewing practices and maintained outside of pure culture industrial yeast selection. We isolated landrace brewing yeast strains from diverse geographical locations across Europe, including Norway, Lithuania, Latvia, Russia, and also included African farmhouse brewing strains from Ghana. Our genomic analysis using long-read and short-read whole genome sequencing uncovered a genetically distinct group that diverges from industrial brewing yeasts. This group, which is closely related to ale brewing strains, is preliminarily named the “European Farmhouse” group and shows greater predicted admixture from Asian fermentation strains. Through genomic and phenotypic analyses, including flavor metabolite analysis via HS-GC-MS, sugar metabolite analysis via HPLC, and wort fermentation analysis, we found a broad spectrum of fermentation capabilities, from rapid and efficient fermentation to unique aroma and flavor compound profiles, potentially offering novel traits for brewing applications. This study highlights the importance of preservation of brewing cultural heritage knowledge and resources including yeast cultures.

**Key Points:**

- A large set of geographically diverse farmhouse brewing strains were characterized
- Norwegian and Baltic farmhouse brewing strains form a distinct genetic group
- Farmhouse strains show considerable diversity in fermentation and flavour formation

## Introduction

Understanding the genetic diversity and characteristics of beer yeast is essential for the brewing industry to develop new flavours, aromas, and fermentation traits that meet modern consumer preferences and changing industry conditions. Currently, the beer industry is rapidly evolving, with an increasing number of craft breweries worldwide. These breweries rely on the diversity of yeasts to create unique beers that set them apart in a competitive market. Therefore, it is vital to study and preserve all available brewing yeast strains. Today, the majority of the commercially used brewing strains tend to cluster into one of two independently domesticated ‘Beer’ groups (Gallone et al. 2016; Gonçalves et al. 2016; Peter et al. 2018; Krogerus et al. 2019). However, non-industrialised brewing strains are still likely to be maintained in traditional brewing systems across the globe (Cubillos et al. 2019).

Farmhouse brewing is the practice where farmers brewed beer for use in their own households from their own grain. Historically, farmhouse brewing was nearly universal among farmers across Europe (Garshol 2020a), and also widespread in other parts of the world. The farmers maintained their own yeast cultures in isolation from those used in modern industrial brewing, and their fermentation practices differed substantially from those in modern brewing. Specifically, the farmers fermented at much higher temperatures (Garshol 2020b) and much faster (Garshol 2022) than modern brewers. Drying the yeast was also very common (Garshol 2020a). These yeasts are known as “farmhouse yeasts”, and because they consist of mixed populations of many strains that have never been isolated into monocultures of single strains, they have also been referred to as “landrace yeasts”.

Despite the widespread use of farmhouse yeasts in farmhouse brewing, little is known about their genetic makeup and their potential to offer new flavours and fermentation traits relevant to modern brewing practices. Farmhouse yeasts are potentially unique due to their isolation and adaptation to specific environments over long periods, sometimes resulting in genetic variations that are not present in industrial brewing yeasts. As such, farmhouse yeasts can offer significant potential for the brewing industry. Beyond commercial applications, the study and preservation of traditional yeasts contributes to the knowledge of cultural heritage in brewing.

In contrast to industrial brewing yeasts which are maintained typically by cryopreservation, traditional farmhouse brewers today use simpler methods such as reusing refrigerated yeast or drying the yeast between batches of beer. These cultures often contain multiple strains of yeast, whereas most commercially produced yeast strains are a monoculture. Traditional farmhouse brewing practices also differ from modern/industrial brewing in several other ways. A wider diversity of beer production methods are used in traditional farmhouse brewing, including techniques such as not boiling the wort (Norway, Denmark, Sweden, Finland, Estonia, Latvia, Lithuania), boiling the mash (Finland), extensive use of juniper and juniper infusions (Norway, Finland, Sweden, Estonia, Latvia, Lithuania), baking the mash (Lithuania, Russia), using grains such as sorghum and millet (Ghana) (Djameh et al. 2019; Garshol 2020a).

Previous research has explored kveik yeast, a Norwegian type of farmhouse yeast, which has gained popularity due to its beneficial features in the brewery. Opportunities identified in kveik yeast include its ability to ferment at high temperatures, reducing energy consumption in the brewing process, and its unique flavour profiles which can provide new beer styles to meet the growing consumer demand (Preiss et al. 2018; Krogerus et al. 2018b). To date, kveik has been demonstrated as useful in various fermentations including beer (Preiss et al. 2018; Kawa-Rygielska et al. 2021; Foster et al. 2022; Kawa-Rygielska et al. 2022; Dippel et al. 2022; Paszkot et al. 2023), Scotch Whiskey (Waymark and Hill 2021), and acid whey beverages (Luo et al. 2021). Recently it was also shown that kveik yeast has enhanced accumulation of trehalose compared to industrial brewing yeasts, which may explain its enhanced stress resistance (Foster et al. 2022). These traits show a broad range of potential for this Norwegian farmhouse yeast.

We hypothesized that farmhouse yeast strains isolated from a more geographically diverse set of sources may also show beneficial fermentation traits and unique genetics when studied. We isolated and selected a group of 35 landrace brewing yeast strains from the following geographic regions (Figure 1A):

- **Norway - West:** The fjord region of Norway, west of the central Norwegian mountain chain. All strains described as kveik in earlier studies are derived from this region.
- **Norway - East/north:** Collected from the village Ål in the Hallingdal region, and have informally been referred to as “gong”, after the local dialect word for yeast.
- **Norway - East/south:** From Tinn county in the province of Telemark, and have informally been known as “berm”. In this region, the brewers now use malt extract rather than mashing malt.
- **Lithuania:** From the northeastern region known as Aukštaitija, which for decades has been known as a hotspot of farmhouse brewing, with many brewers having started commercial breweries.
- **Latvia:** From the Latgale region in eastern Latvia, which culturally and linguistically differs substantially from the rest of Latvia. The strains were collected about 120km from the Lithuanian border.
- **Russia:** All strains were collected in the Chuvash Republic, a region on the south bank of the Volga, 600km east of Moscow, populated largely by the Chuvash people, who speak a Turkic language. All samples were collected from Chuvash speakers.
- **Ghana:** Collected from a pito brewer from Yendi in the northern region of Ghana, approximately 22 km from the northwestern border with Togo. Traditional brewers in this region mainly produce pito (sorghum/millet beer) with backslopped yeast from a previous brew.

**Figure 1.**
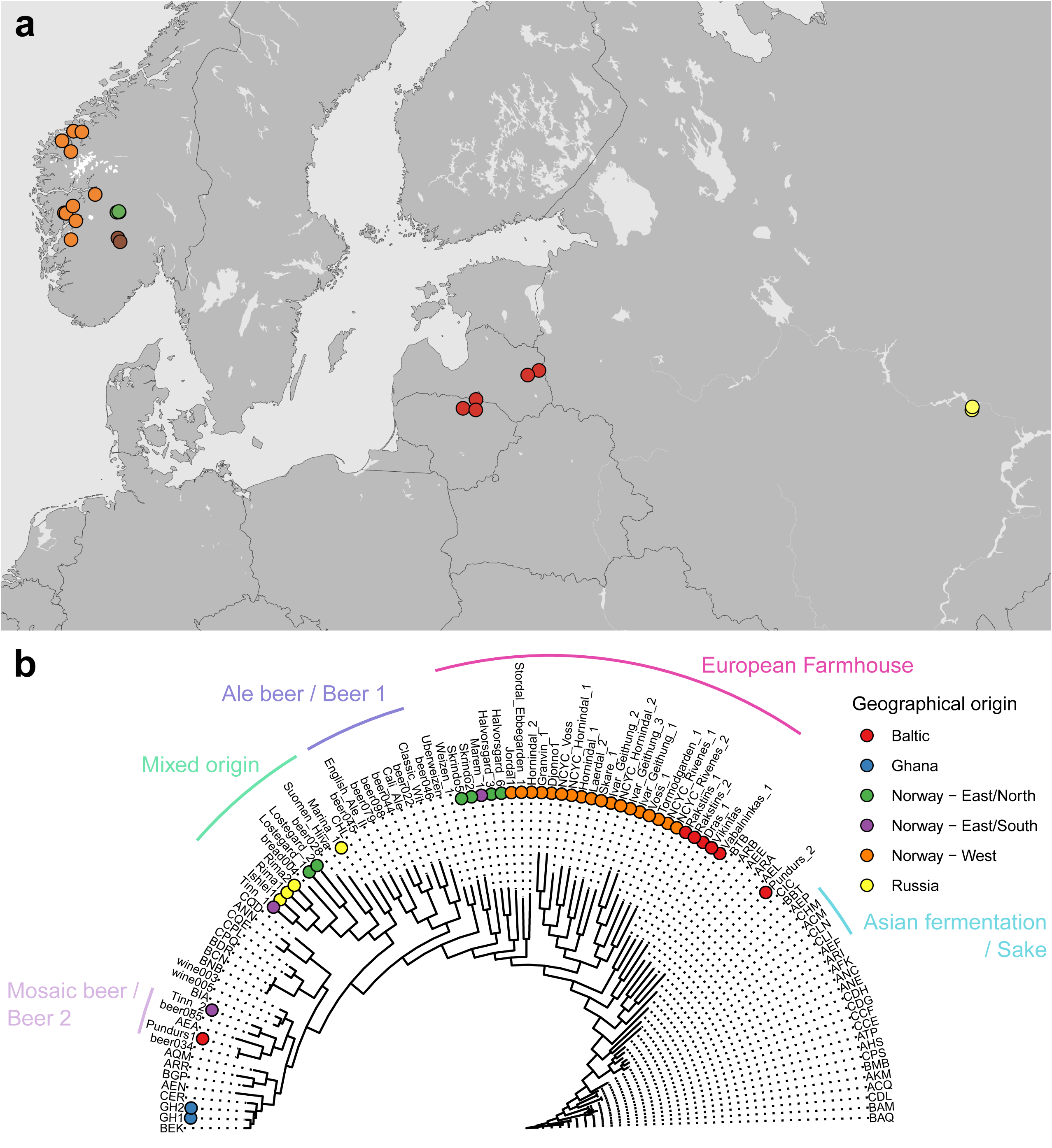
Phylogeny and geographical origin of the 35 *S. cerevisiae* landrace brewing strains sequenced in this study. **(A)** A map showing the regions from which the European landrace yeast strains were isolated from. **(B)** Maximum likelihood phylogenetic tree based on SNPs at 103555 sites in 105 *S. cerevisiae* strains. The set of strains include 35 landrace brewing strains studied here, 5 previously sequenced ‘kveik’ isolates, 5 control brewing strains, and 65 randomly selected strains from Peter et al. (2018) and Gallone et al. (2016) that represent the main *S. cerevisiae* populations. Information about the strains and their origin is available in Supplementary Table S1. The tree is rooted with the ‘China III’ clade as outgroup. Points in both (**A**) and (**B)** are coloured according to geographical origin of the strain.

These farmhouse yeasts, which have avoided the pure culture isolations and commercialization that industrial beer yeast have undergone, may be genetically distinct from industrial yeasts due to their adaptation to specific environments or niches over time, resulting in genetic and phenotypic variations unique to each strain or group. Early research on kveik isolates, for example, indicate that they are closely related, but genetically distinct, from ale brewing yeast (Preiss et al. 2018; Dondrup et al. 2023). Furthermore, the characterization of non-industrialised brewing yeast could further uncover the complex origins of brewing yeast (Fay et al. 2019; Abou Saada et al. 2022).

We further hypothesize that landrace yeasts may offer new combinations of flavour and fermentation traits relevant to modern brewing practices such as craft brewing. To test these hypotheses, we conducted both whole genome sequencing (short read and long read) and phenotypic analyses, such as flavour metabolite analysis via HS-GC-MS, sugar metabolite analysis via HPLC, and wort fermentation analysis.

## Materials & Methods

### Yeast strain isolation

European landrace yeast samples in either dry or liquid format were received at the research site in Canada from a source in Norway. Upon receipt, yeast samples were hydrated or inoculated into sterile beer wort (10 °P) at a rate of approximately 0.5 g / 10 mL. The samples were incubated overnight at 25 °C then aseptically streak plated onto WLN Agar and incubated for 5 days. Unique colony morphologies and colour patterns on WLN agar were selected and sub-streaked onto fresh WLN agar. The yeasts from Ghana were obtained in a powdered format from dried sediments of a pito brew received from a traditional brewer in Accra, Ghana. The samples were inoculated into 10 mL YPD and incubated with shaking at 30 °C for 24 hr after which single yeast colonies from the cultures were isolated on YPD agar plates. Pure cultures were cryopreserved at −80 °C prior to analysis. The colonies were verified as *Saccharomyces cerevisiae* by performing sequence analysis of the ITS region of the yeast isolates. Table 1 lists the strains selected for analysis. The additional strains included in the genomic analyses are listed in Supplementary Table S1).

**Table 1.**
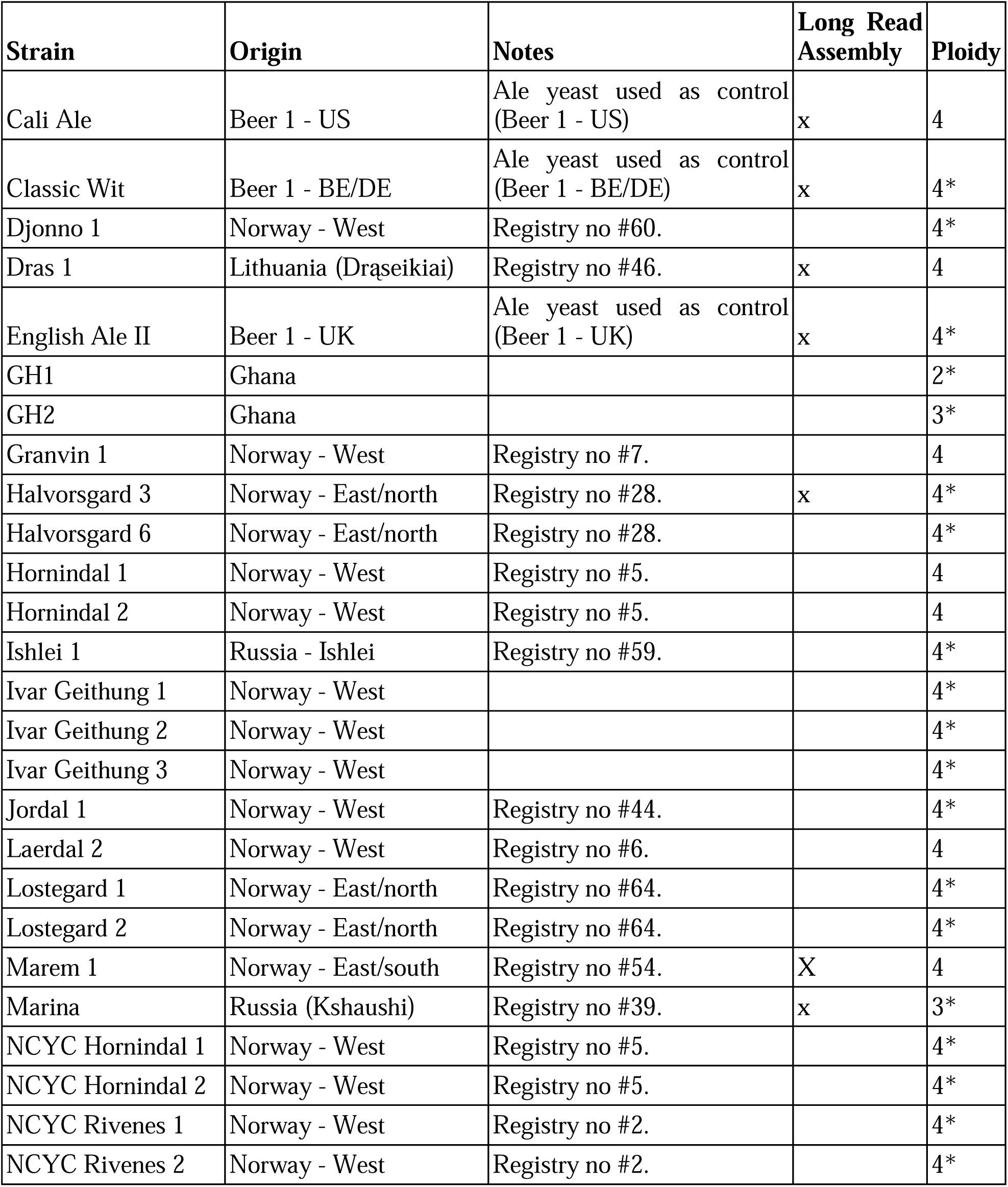
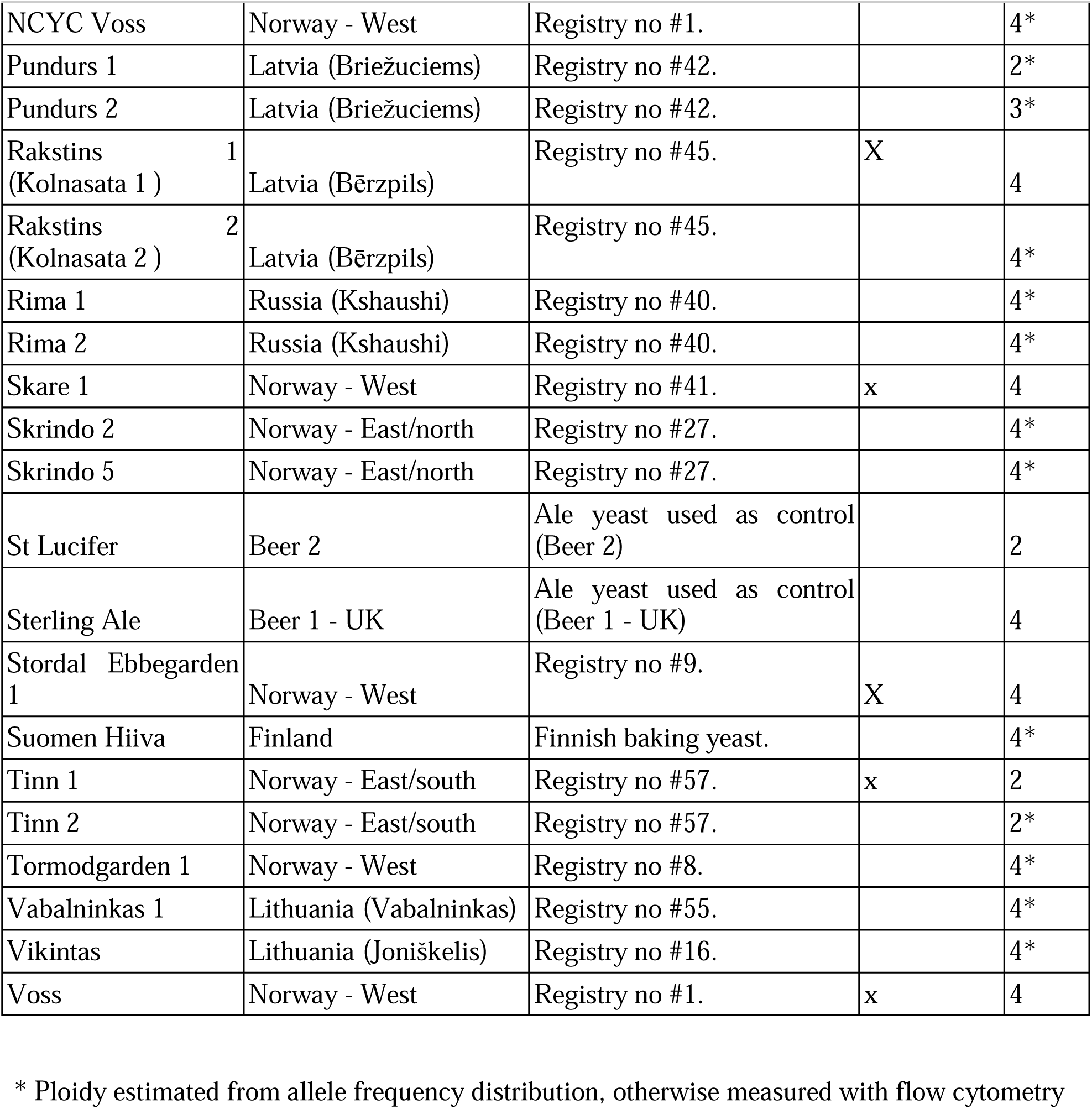
Strains selected for this study and their respective origins. All strains were sequenced with short-read sequencing.

### Yeast Genome Sequencing and Analysis

Short read Illumina sequencing was performed either as described in Foster et al. (2022) or otherwise DNA extractions were performed using the ZymoBIOMICS™ DNA Miniprep Kit (Zymo Research). Short-read sequencing of strains was performed at SeqCenter LLC (Pittsburgh, PA). Illumina sequencing libraries were prepared using the tagmentation-based and PCR-based Illumina DNA Prep kit and custom IDT 10bp unique dual indices (UDI) with a target insert size of 320 bp. No additional DNA fragmentation or size selection steps were performed. Illumina sequencing was performed on an Illumina NovaSeq 6000 sequencer in one or more multiplexed shared-flow-cell runs, producing 2×151bp paired-end reads. Demultiplexing, quality control and adapter trimming was performed with bcl-convert1 (v4.1.5).

Long-read sequencing experiments were carried out at the University of Guelph, University of Waterloo and VTT. In the two former locations, HMW yeast genomic DNA was extracted as prescribed by Oxford Nanopore according to Denis et al. (2018). Sequencing was performed with the Oxford Nanopore MinION instrument on a R9.4 flow cell using the SQK-LSK109 ligation sequencing kit. At VTT, DNA was extracted using a Monarch® HMW DNA Extraction Kit for Tissue (New England Biolabs) and the supplied protocol for yeast. Sequencing was carried out on an Oxford Nanopore MinION MK1C instrument on a R10.3 flow cell using the SQK-LSK110 ligation sequencing kit. Reads were basecalled using Guppy version 5.0.11 using the ‘super high accuracy’ model.

Reads were trimmed to a minimum length and quality of 1000 bp and Q8 using NanoFilt (De Coster et al. 2018). Genomes were de novo assembled from the long reads using Flye version 2.9 (Kolmogorov et al. 2019). Assemblies were first polished using Medaka version 1.4.3, and then with the short reads using NextPolish version 1.3.1 (Hu et al. 2020). Short reads were trimmed and filtered with fastp using default settings (version 0.20.1; Chen et al. 2018). Trimmed reads were aligned to a *S. cerevisiae* S288C reference genome (Engel et al. 2014) using BWA-MEM (Li and Durbin 2009), and alignments were sorted and duplicates were marked with sambamba (version 0.7.1; Tarasov et al. 2015). Variants were jointly called in all strains using FreeBayes (version 1.32; Garrison and Marth 2012). Variant calling used the following settings: --min-base-quality 30 --min-mapping-quality 30 --min-alternate-fraction 0.25 – min-repeat-entropy 0.5 --use-best-n-alleles 70 -p 2. The resulting VCF file was filtered to remove variants with a quality score less than 1000 and with a sequencing depth below 10 per sample using BCFtools (Li 2011). Variants were annotated with SnpEff (Cingolani et al. 2012). Sequencing coverage was estimated with mosdepth (version 0.2.6; Pedersen and Quinlan 2018). Chromosome copy numbers were estimated based on distribution of alternate allele frequencies, ploidy as measured by flow cytometry, and sequencing coverage.

For phylogenetic analysis, the variants were filtered to retain only single nucleotide polymorphisms and remove sites with a minor allele frequency less than 5%. The filtered SNP matrix was converted to PHYLIP format (https://github.com/edgardomortiz/vcf2phylip). A random allele was selected for heterozygous sites. A maximum likelihood phylogenetic tree was generated using IQ-TREE (version 2.0.3; Nguyen et al. 2015) run with the ‘GTR+G4’ model and 1000 bootstrap replicates (Minh et al. 2013). In addition, neighbour-joining trees based on MinHash distances (Ondov et al. 2016) between genome assemblies were generated using mashtree (Katz et al. 2019). Mashtree was used with the long-read assemblies produced here, the short-read assemblies from Peter et al. (2018) and Gallone et al. (2016), and the long-read assemblies from O’Donnell et al. (2023).

Population structure was estimated based on the identified SNPs. The SNP matrix produced above was filtered using PLINK (version 1.9; Purcell et al. 2007) by removing sites in linkage disequilibrium (using a 50 SNP window size, 5 SNP step size, and pairwise threshold of 0.5) and with a minor allele frequency <5%. The thinned SNP matrix was then input into ADMIXTURE v.1.3.0 (Alexander et al. 2009), and was run for 1 to 20 ancestral populations (*K*). The number of ancestral populations (*K*) that best represented this dataset was chosen based on the lowest cross-validation error. The analysis was also re-run on a dataset produced by merging the SNP matrix available from the 1,011 yeast genomes dataset (Peter et al. 2018) with SNPs identified in the landrace strains sequenced here using the the ‘-@’ option in FreeBayes to force calls at all locations in the 1,011 gVCF file. VCF files were merged with BCFtools (Li 2011), and then filtered as described above.

Admixture between populations was further estimated from allele frequency data using both TreeMix (Pickrell and Pritchard 2012) and AdmixtureBayes (Nielsen et al. 2023). The thinned SNP matrix used as input to ADMIXTURE above was used for analysis. First, strains belonging to ‘Mosaic group 3’ were removed, and the VCF was converted to TreeMix format using the script from https://github.com/speciationgenomics/scripts/blob/master/vcf2treemix.sh. TreeMix (version 1.13) was run based on the pipeline available at https://github.com/carolindahms/TreeMix. To find the optimum number of migration events, the pipeline was run using the following options: migration events (*m*) ranging from 1 to 16, ten replicates per migration event, 500 bootstrap replicates, and China_III as the outgroup. OptM (Fitak 2021) was then used to find the optimum number of migration events for the data set (*m* = 6). The TreeMix pipeline was then rerun with *m* = 6 as above, but with 30 replicates, and a consensus tree was produced with BITE (Milanesi et al. 2017). *D, f_4_*-ratio and *f*-branch statistics were calculated with DSuite from all the polymorphic sites detected among the 105 strains (Malinsky et al. 2021). For AdmixtureBayes, which estimates a best-fitting admixture graph without priori information, the same input file as for TreeMix was used. AdmixtureBayes was run with 20 MCMC chains with 75,000,000 iterations (-n 1500000). Convergence was tested by discarding the first 30% of samples as burn-in.

Phasing of short reads was carried out using nPhase version 1.2.0 (Abou Saada et al. 2021) and the workflow outlined in (Abou Saada et al. 2022) for the strains that had also been long-read sequenced. The predicted haplotigs for each strain are visualised in Supplmentary Figure S1. The phased SNPs for each predicted haplotig, as output by nPhase, were then applied to the sequence of the *S. cerevisiae* S288C reference genome to produce sequences for each of the predicted haplotigs. These sequences were further split into 10 kbp windows. In addition to the phased haplotigs, 53 genomes, representative of the 19 main clades identified in (Peter et al. 2018), were obtained from NCBI. These genomes were aligned to the *S. cerevisiae* S288C reference genome using minimap2 version 2.17-r941, and variants were called using paftools.js (included within minimap2). The pairwise distances between all the phased haplotigs and the representative genomes within each of the 10kbp windows was calculated using snp-dists (https://github.com/tseemann/snp-dists). Within the resulting distance matrices for all 10 kbp windows, we identified among the haplotigs for each of the phased strains the minimum distance compared to each of the 53 representative genomes.

### Ploidy with flow cytometry

Ploidy of selected strains was measured using SYTOX Green staining and flow cytometry as described previously (Krogerus et al. 2017).

### Yeast Propagation and Beer Fermentation

To perform lab scale fermentations, yeast starters were grown in 20 mL wort at 25 °C for 24 hr. A cell count was performed on the yeast starters using a haemocytometer after which the starters were used to pitch fresh wort with an original gravity of 10.5 °P (specific gravity = 1.043 ± 0.0014) at a rate of 1 million cells/mL/°P. The wort used for the fermentation was made from pale barley malt (Rahr) and hopped with 2.4 g/L Cascade hops. Three biological replicates of the fermentations were carried out in 500 mL glass bottles fitted with airlocks and incubated without shaking at 25 °C for 7 days. Specific gravity readings of the fermenting wort were measured daily using a DMA35 handheld density meter (Anton Paar GmbH, Graz, Austria). At the end of the fermentation, 20 mL of the fermented wort was taken, filter-sterilized and stored for HPLC and GC-MS analysis.

### Sugar and ethanol analysis by HPLC

The fermented beer samples were prepared for HPLC analysis by sterile filtration using a 0.22 micron syringe filter. Samples were prepared in an HPLC vial containing 400 µL of filtered beer and 50 µL of 6% (v/v) isopropanol as the internal standard. Measurements were performed using HPLC with a refractive index detector (RI). An Aminex HPX-87H column was used, where 5mM sulfuric acid was the mobile phase. The conditions were: flow rate 0.6 mL/min, 620 psi, 60°C.

### Aroma compound analysis by HS-GC-MS

5mL of sterile-filtered beer samples and internal standard mix solution [1% (v/v) 3-octanol and 0.005% (v/v) ethyl methyl sulfide] of 20 µL were pipetted into a 20 mL HS vial sealed by a metal screwcap with a PTFE/silicone septa for quantitation of higher alcohols and esters by Agilent GC 7890B coupled with a PAL autosampler (RSI 85, CTC Analytics, Zwingen, Switzerland) and an MSD 5977B (Agilent, Santa Clara, CA, USA).

Beer samples were incubated at 40°C for 20 min to reach equilibrium, followed by injection of 1 mL headspace with syringe temperature set at 70°C, inlet temperature at 200°C, and a split ratio of 10. Separation of volatile compounds was carried out with a DB-WAX UI capillary column (60 m x 0.25 mm, 0.25 um film thickness, Agilent, Santa Clara, CA, USA) and helium as the carrier gas at a constant flow of 1.2 mL/min. The oven program was stated as below: initial temperature held at 35°C for 1 min; increased to 120°C by 15°C /min; increased to 180°C by 5°C /min; further increased to 250°C by 20°C/min, and held at 250°C for 5 min. The temperature of the transfer line to MS was set at 250°C. The detection was performed at TIC mode (m/z 25-250) with electron-impact ionization at 70 eV, ion source temperature at 230°C, and quadrupole temperature at 150°C.

Ethyl hexanoate, ethyl octanoate, ethyl decanoate, phenylethyl acetate (MilliporeSigma, St. Louis, MO, USA), and pre-made standard mixture containing acetaldehyde, N-propanol, ethyl acetate, isobutanol, isopropyl acetate, ethyl propanoate, active amyl alcohol, isoamyl alcohol, isobutyl acetate, ethyl butanoate, N-butyl acetate, and isoamyl acetate (SPEX CertiPrep Inc, Metuchen, NJ, USA) were used to establish standard calibration curves. 3-octanol was used as the internal standard for quantitation.

## Results

### Isolation and genomic analysis of landrace yeasts

A cohort of 35 landrace brewing yeasts was selected on the basis of geographic and cultural diversity from a yeast strain collection containing approximately 250 yeast isolates obtained from original farmhouse brewing cultures (Table 1; Figure 1A). Further details for the farmhouse brewing yeasts can be obtained from the Farmhouse Yeast Registry (Garshol 2020c)^1^. Previously studied and sequenced industrial ale strains as well as kveik strains were also included in the analysis (Preiss et al. 2018; Foster et al. 2022).

To clarify the relatedness and ancestry of the landrace strains, whole-genome sequencing of the included strains was performed. Short-read sequence data was previously available for six of the kveik strains included here (Preiss et al. 2018), while the remaining 30 landrace strains were short-read sequenced (average coverage of 125, ranging from 29.5 to 406). The majority of the landrace strains were tetraploid (28 out of 35; Table 1), with many strains (20 out of 35 aneuploid) showing chromosome copy number variations based on read coverage and allele frequency distributions (Supplementary Figures S2 and S3). Nine of the landrace strains were further long-read sequenced using nanopore sequencing. These nine strains included six of Norwegian origin (from all three sub-regions), and three of Baltic and Russian origin (Table 1). In addition to these landrace strains, long-read sequence data was also obtained for the six brewing and baking strains included as controls (Foster et al. 2022). First, long-read assemblies were produced for these strains. Genome assembly size ranged from 11.8 to 12.3 Mbp, contig number from 16 to 45, and N50 values from 751 to 927 kbp (statistics available in Supplementary Table S2). The metrics suggested high contiguity and completeness of the assemblies.

To better understand how these nine landrace strains were genetically related to other wild and domesticated *S. cerevisiae*, an alignment-free neighbour-joining tree was first produced based on minhash distances between the landrace strains and the short-read assemblies from the 1,011 yeast genomes data set (Peter et al. 2018) and Gallone et al. (2016). The resulting tree (Figure 2) grouped seven of the landrace strains as a sister group to the ‘Beer 1 / Ale beer’ group. This group included five Norwegian landrace strains, and, surprisingly, two of the Baltic strains (Dras 1 and Rakstins 1). The remaining two landrace strains (Tinn 1 and Marina 1) were placed in the ‘Mixed / Beer-Baking’ group. The six control strains grouped as expected in the ‘Beer 1’ and ‘Mixed’ groups.

**Figure 2.**
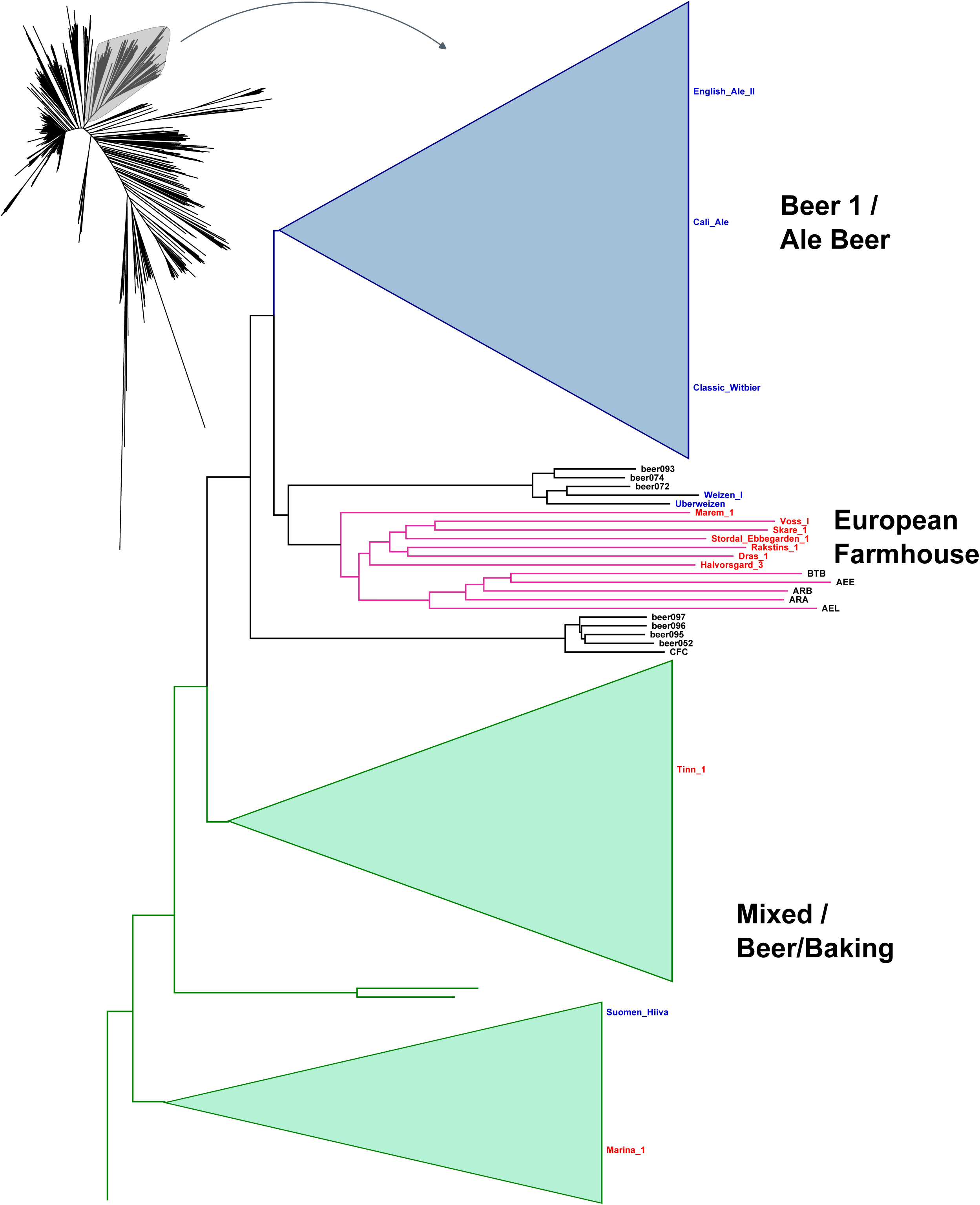
A neighbour-joining tree generated based on MinHash distances in the assemblies of 1,011 yeast genomes (Peter et al. 2018) and long-read assemblies of 9 landrace and 6 control strains generated here. Strain names of the landrace and control strains are coloured red and blue, respectively. The branch containing the ‘Mixed origin’, ‘European farmhouse’ and ‘Beer 1’/’Ale beer’ populations is highlighted in grey.

These results agree with previous work on kveik (Preiss et al. 2018), suggesting that ‘kveik’ strains are closely related to, yet genetically distinct from, domesticated ale yeasts. However, farmhouse strains from outside Norway also clustered with the ‘kveik’ strains, so the group does not seem to be geographically restricted to Norway. Hence, we decided to preliminarily rename this group ‘European Farmhouse’. Interestingly, five strains from the 1,011 yeast genomes set, identified as belonging to Mosaic group 3, also grouped in a sister branch to these ‘European Farmhouse’ strains. This could hint towards an admixed ancestry of these strains, as also hypothesized earlier by Preiss et al. (2018). To ensure that no artifacts were introduced from comparing long-read assemblies with short-read assemblies, a second neighbour-joining tree was produced based on minhash distances between the landrace strains and the recently published long-read assemblies in the ScRAP data set (O’Donnell et al. 2023). The resulting tree again placed the nine landrace and six control strains in the same groups, and the ‘European Farmhouse’ group once again formed a sister group to ‘Beer 1’ (Supplementary Figure S4).

Next, we expanded the phylogenetic analysis to the full set of 35 landrace strains. Single nucleotide polymorphisms (SNPs) were first identified from the short-read data. SNPs were also simultaneously called for five control brewing strains, five ‘kveik’ isolates sequenced by NCYC (Bioproject PRJEB42916), and 65 randomly selected *S. cerevisiae* strains representing 20 of the main *S. cerevisiae* populations identified by (Peter et al. 2018). A maximum-likelihood phylogenetic tree was then inferred from the resulting SNP matrix consisting of 103555 variable sites. As with the minhash trees, the majority of the landrace strains (23 of 35, as well as the five ‘kveik’ isolates sequenced by NCYC) grouped together, branching off before the ‘Beer 1’ and ‘Mixed’ strains (Figure 1B). Seven of the remaining strains grouped within the ‘Mixed’ group, two in ‘Beer 2’, and the two Ghanaian strains among the ‘African Beer’ strains. Of the total of 28 strains in the ‘European Farmhouse’ group, 23 were of Norwegian origin, while the remaining five were of Baltic origin. A sixth strain of Baltic origin grouped together with strains of Mosaic group 3. Like in the minhash tree produced from the long-read assemblies, the Baltic strains formed their own branches in the ‘European Farmhouse’ group. Interestingly, among the Norwegian strains, two sub-groups also form depending on geographical origin. One consists of the strains deriving from Norway - West, the so called ‘kveik’ strains, including those from e.g. Voss, Stordal Ebbegarden, and Skare, while the other consists of strains isolated in Eastern Norway, in the Norway - East/north and East/south regions, including those from e.g. Halvorsgard and Skrindo. Of the five landrace strains in the ‘Mixed’ group, four were of Russian origin, while Tinn 1 derives from Norway East/south. In contrast to the assembly-based tree, where the ‘European Farmhouse’ strains grouped as a sister branch between the ‘Beer 1’ and ‘Mixed’ groups, here in the SNP-based tree, the ‘European Farmhouse’ strains branched off closer to the root of the tree after the ‘Mosaic 3’ and ‘Asian fermentation’ strains. This is likely a result of the assembly-based tree not capturing the high levels of heterozygosity found in ‘European Farmhouse’ strains (ranging from 0.34 to 0.55%; Supplementary Table S1), which could indicate an admixed ancestry.

An extended SNP matrix was further produced containing all strains from 1,011 yeast genomes (Peter et al. 2018), the 35 landrace and five brewing control strains sequenced here, and the five sequenced NCYC ‘kveik’ strains. First, SNPs were recalled in the 45 strains based on all the sites in the 1,011 yeast genomes gVCF, after which the SNPs in the landrace and brewing strains were merged with those in the 1,011 yeast genomes data set. The resulting SNP matrix consisted of 310688 variable sites after filtering to retain only sites with a minor allele frequency > 1%. As in the assembly-based tree, the maximum-likelihood phylogenetic tree produced from this SNP matrix again grouped the ‘European Farmhouse’ strains as a sister branch to the ‘Beer 1’ group (Supplementary Figure S5). Within the ‘European Farmhouse’ group, the strains of Baltic origin again grouped separately to those of Norwegian origin. Together, the results so far highlight that ‘European Farmhouse’ strains are closely related but genetically distinct to ‘Beer 1’/’Ale beer’ brewing strains, and they may have an admixed ancestry. Fascinatingly, this group is not geographically restricted to Norway as indicated by previous preliminary studies restricted to the ‘kveik’ group. Rather, this group consists of farmhouse brewing strains that have been maintained by brewers in Norway and around the Baltic Sea, providing a valuable set of brewing strains genetically distinct from the ‘Beer 1’ strains.

### Admixture analysis reveals Asian influence on European Farmhouse genomes

To further investigate the possibility of admixed ancestry among the ‘European Farmhouse’ strains, population structure and recent admixture events were estimated based on the polymorphic sites among the 105 strains (Figure 1B) using a range of methods. First, the SNP matrix was filtered to remove sites in linkage disequilibrium and with minor allele frequencies <5%. Using ADMIXTURE, the number of populations that best represented this dataset was eight (Figure 3A). Here, the ‘European Farmhouse’ strains formed their own population, with a subset of the strains showing admixture with the ‘Beer 1’ and ‘Asian fermentation’ populations. The five ‘Mosaic group 3’ strains from the 1,011 yeast genomes set, which in both the minhash- and SNP-based phylogenies grouped close to the ‘European Farmhouse’ strains, showed distinct patterns compared to the latter. Because of the high representation of landrace strains in the set of 105 strains, the analysis was further repeated with the larger data set produced above by merging the 1,011 yeast genomes and the landrace and brewing strains sequenced here. After the same filtering as above, ADMIXTURE was rerun, and 16 populations best represented this expanded data set (Supplementary Figure S6). Now, the ‘European Farmhouse’ and ‘Beer 1’ strains grouped in the same population, with all ‘European Farmhouse’ strains also showing evidence of admixture from strains in the ‘Asian fermentation’ population (Figure 3B). Among the Norwegian landrace strains, the Norway East/(north and south) strains had a lower proportion of ‘Asian fermentation’ alleles compared to the Western ‘kveik’ strains. The Baltic landrace strains, on the other hand, showed a higher proportion of ‘Asian fermentation’ alleles compared to the Norwegian farmhouse strains, which geographically makes sense.

**Figure 3.**
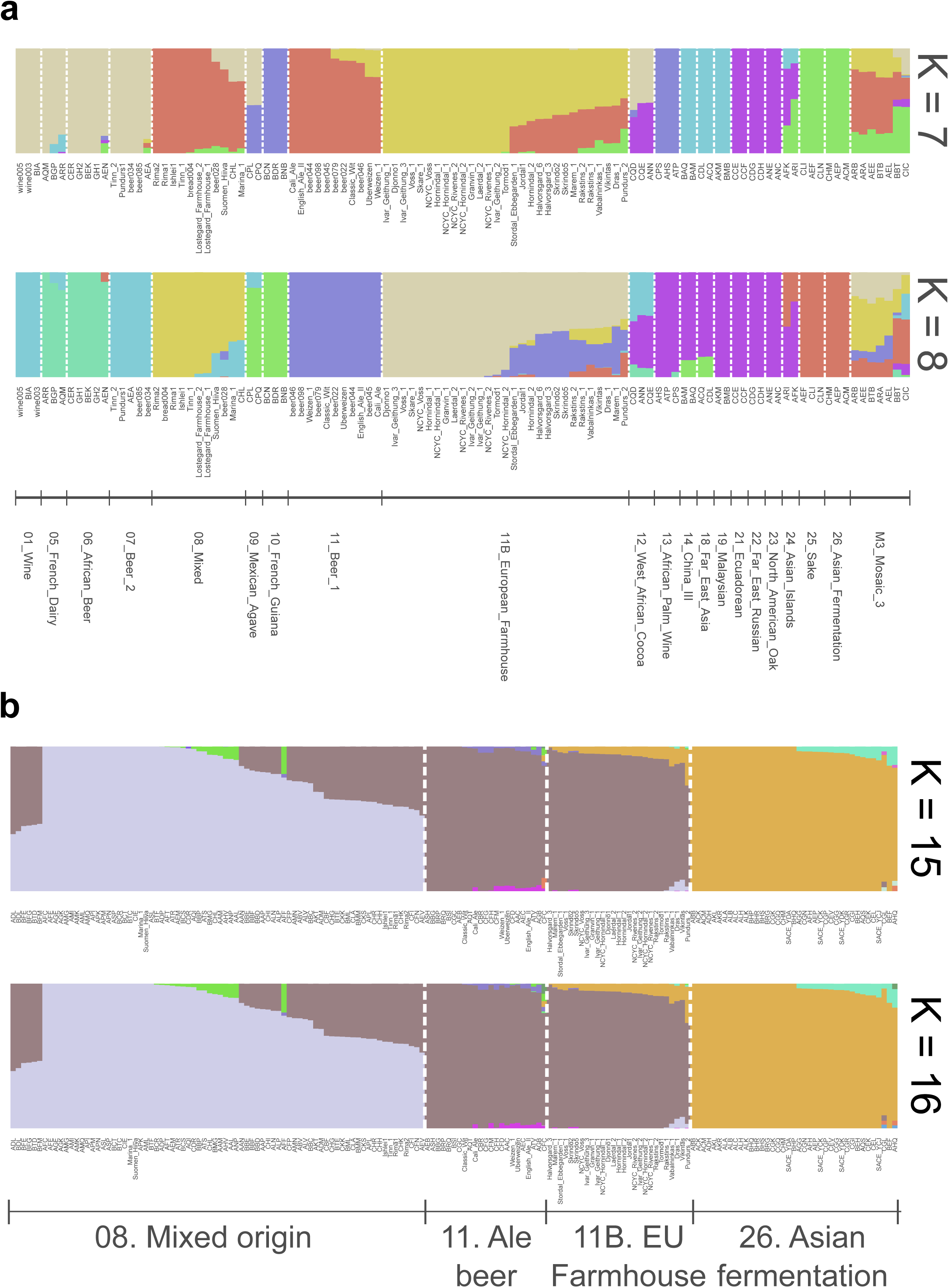
Population structure of the 35 *S. cerevisiae* landrace brewing strains and reference strain data sets. **(A)** Population structure of the set of 105 *S. cerevisiae* strains estimated with ADMIXTURE based on SNPs at 47954 sites. Each strain along the x-axis is represented by a vertical bar partitioned into colors based on estimated membership fractions to the resolved populations for K = 7 and 8 assumed ancestral populations. **(B)** Population structure of a merged set of 35 *S. cerevisiae* landrace brewing strains and the 1,011 yeast genomes strains estimated with ADMIXTURE based on SNPs at 31766 sites. Strains belonging to the ‘Mixed origin’, ‘Ale beer’, ‘Asian fermentation’, and ‘European Farmhouse’ populations are represented along the x-axis by a vertical bar partitioned into colors based on estimated membership fractions to the resolved populations for K = 15 and 16 assumed ancestral populations. Plots covering all strains for K = 8 to 20 assumed ancestral populations are available as Supplementary Figure S6.

To test for migration events, we also ran TreeMix and AdmixtureBayes on the thinned SNP matrix. With TreeMix, the optimal number of migration events that best explained the variation in the set was predicted to be six using the ‘Evanno’ method implemented in OptM (Fitak 2021). Consistent with previous work (Fay et al. 2019; Abou Saada et al. 2022), migration events from the Asian fermentation populations to both the root of the ‘Beer 1’-‘European Farmhouse’-‘Mixed origin’ branch (35% migration weight) and the Beer 2 population (14% migration weight) were predicted (Figure 4A). In addition, a third admixture event from the Asian fermentation populations to the ‘European Farmhouse’ population was observed (20% migration weight), providing additional evidence for the ‘kveik’ and other ‘European Farmhouse’ strains emerging from admixture between a ‘Beer 1’ ancestor and a strain of Asian origin. The ‘European Farmhouse’ strains therefore exhibit a higher proportion of Asian ancestry compared to ‘Beer 1’ strains. These migration events are also supported by *D, f_4_*-ratio and *f*-branch statistics calculated from the full set of polymorphic sites across the 105 strains (Supplementary Figure S7). The multiple migration events from the Asian fermentation populations into the brewing populations seems to suggest a fitness advantage for the ‘Wine – Asian fermentation’ intraspecific hybrid in a malt fermentation environment.

**Figure 4.**
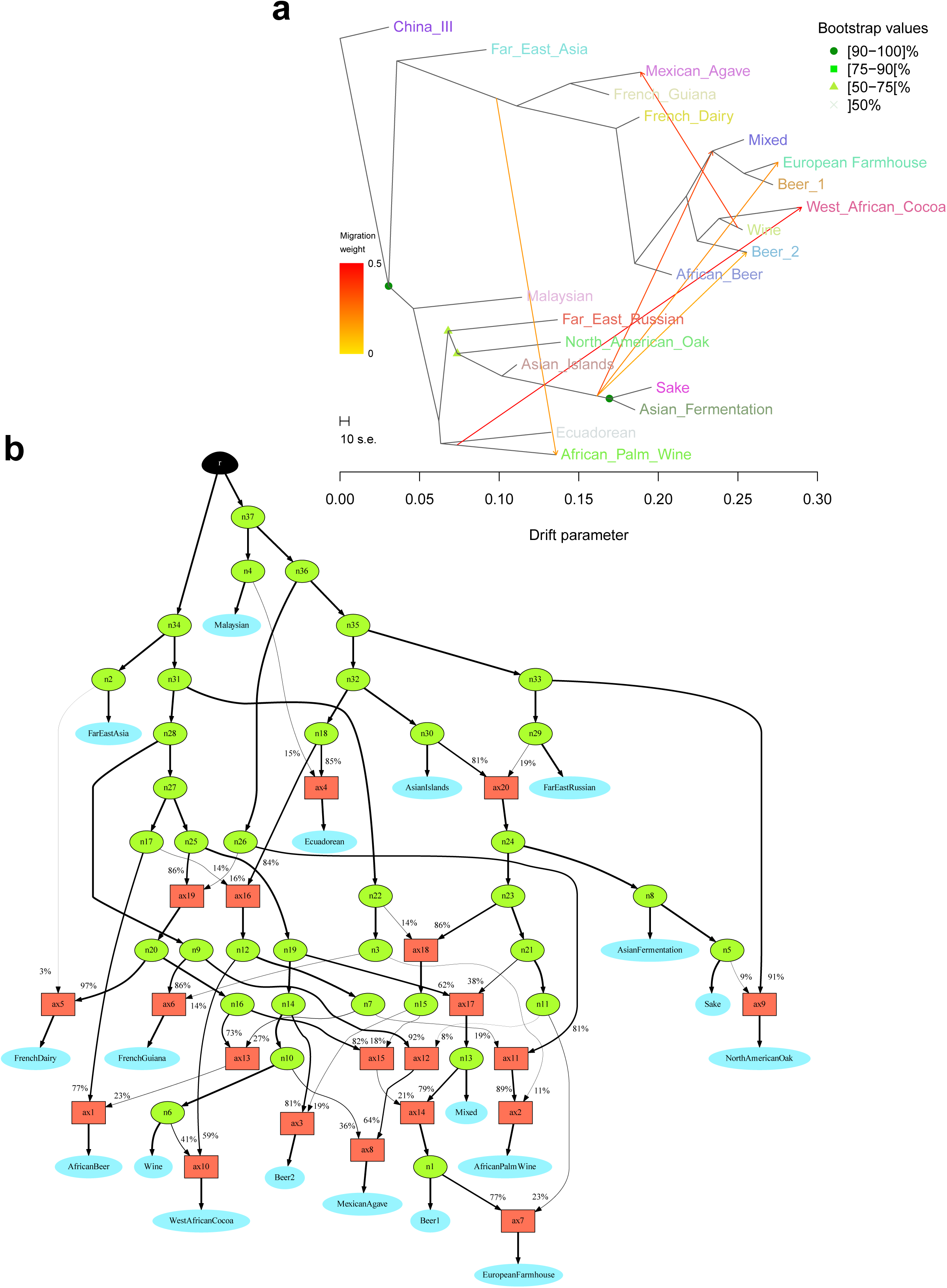
Phylogenetic networks inferred by (**A**) Treemix and (**B**) AdmixtureBayes in the set of 105 *S. cerevisiae* strains. (**A**) Using the ‘Evanno’ method implemented in OptM, six migration events between populations inferred by Treemix were selected and are shown with arrows indicating the direction toward the recipient population and coloured according to migration weight (ancestry percentage received from the donor). The scale bar shows ten times the average standard error of the entries in the sample covariance matrix. (**B**) The network graph topology with highest posterior probability as derived by AdmixtureBayes. The data set converged to and was best described by 20 admixture events. Red squares indicate admixture events, while blue ovals indicate populations. Percentage numbers on the branches and branch thickness represent admixture proportions.

AdmixtureBayes was also used to estimate an admixture graph for the set of 105 strains studied. The tool uses a reversible jump Monte Carlo Markov Chain algorithm to sample high-probability admixture graphs, and has been shown to infer graph topologies more accurately than TreeMix (Nielsen et al. 2023). Here, after running 75,000,000 iterations, the population structure was best described by 20 admixture events. The topology with highest posterior probability suggested the ‘European Farmhouse’ population formed from admixture between a ‘Beer 1’ ancestor (77% contribution) and a node derived from an ancestor of the ‘Asian Fermentation’ and ‘Sake’ branch (23% contribution) (Figure 4B). Likewise, admixture from these Asian fermentation populations was also predicted in the formation of the ‘Mixed origin’-‘Beer 1’ branch, as well as the ‘Beer 2’ population. Hence, results here further confirm our previous observations. AdmixtureBayes was also rerun on the extended SNP data set (strains of 1,011 yeast genomes and the 35 landrace strains studied here). Strains from 1,011 yeast genomes that had not been assigned to a clade, or belonged to one of the three mosaic regions, were excluded from the analysis. The resulting top topology also included 20 admixture events and predicted similar admixture events in the forming of the ‘European Farmhouse’ and other brewing and baking populations (Supplementary Figure S8).

Finally, SNP phasing was used to demonstrate the strong contribution from Asian fermentation strains in the ‘European Farmhouse’ strains. For 17 of the strains where long-read sequences were available (this included nine landrace strains), phased haplotypes were produced using nPhase (Abou Saada et al. 2021). Similar to the approach described by Abou Saada et al. (2022), the obtained haplotypes for each strain were then divided into 10 kbp windows. Within each of the 17 strains, and each 10 kbp window, the sequences of the strain’s haplotypes were compared to those of 53 genomes selected to represent the major populations described in Peter et al. (2018), by calculating the pairwise genetic distance. For each window, the haplotype with the smallest distance was retained. The genome-wide average minimum distance for each of the 17 phased strains compared to the 53 representative genomes was then plotted (Figure 5). Results reveal that compared to the included control ‘Beer 1’ strains, the ‘European Farmhouse’ strains contain haplotypes with a lower genetic distance to the ‘Asian Fermentation’ and ‘Sake’ strains. Two strains, AEL and BBT, from ‘Mosaic Region 3’ in the 1,011 yeast genomes (Peter et al. 2018) and ScRAP data sets (O’Donnell et al. 2023) were also included in the analysis, since they clustered close to the ‘European Farmhouse’ strains in phylogenetic analysis. Here, however, they showed distinct profiles to that of the ‘European Farmhouse’ strains, particularly showing a higher minimum genetic distance to the ‘Beer 1’ strains. Taken together, results here strongly support the hypothesis of kveik and other ‘European Farmhouse’ strains forming from the admixture of a ‘Beer 1’ ancestor and a strain with origins in Asian fermentations.

**Figure 5.**
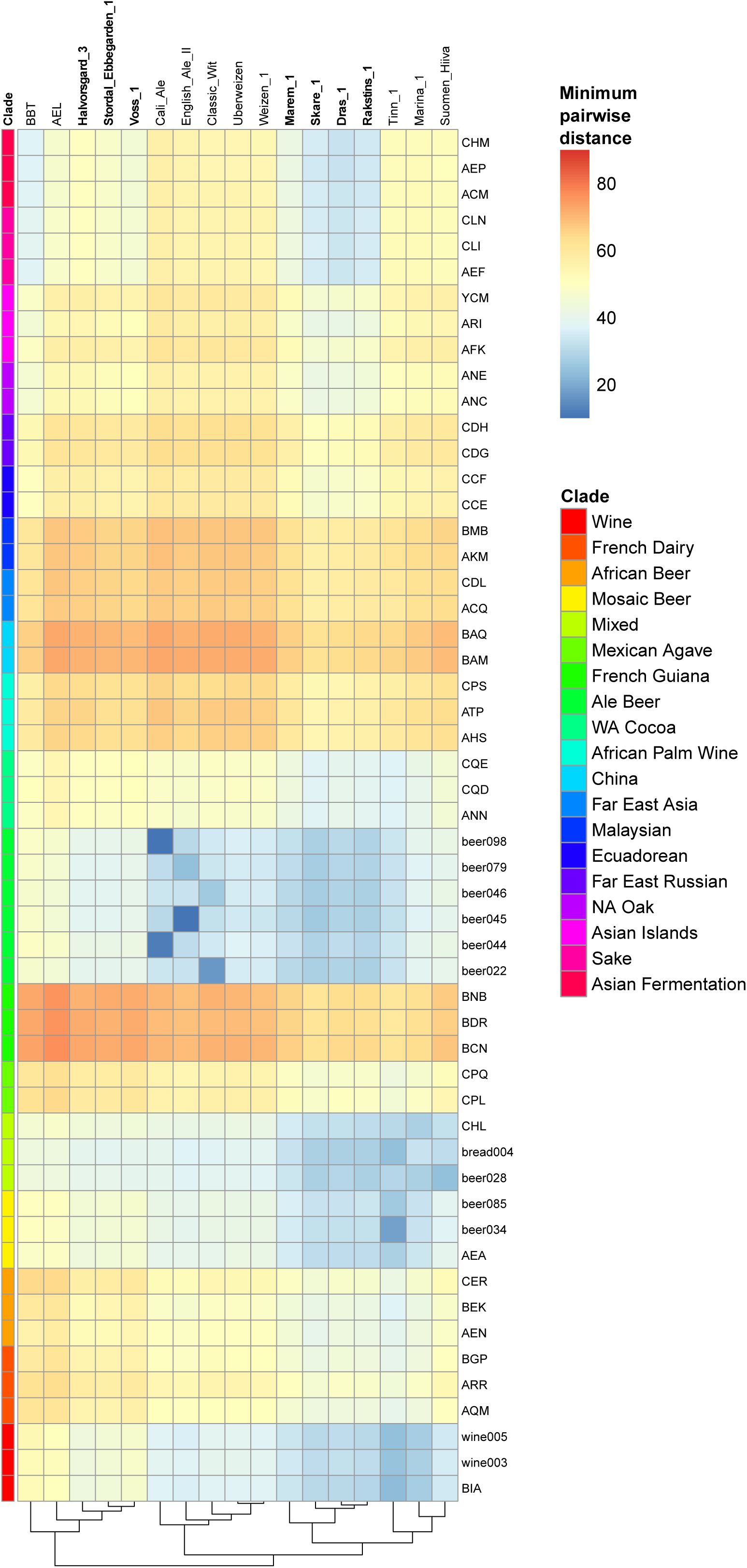
Minimum genetic distance among the 17 phased haplotypes (columns) compared to 53 *S. cerevisiae* reference strains representing the major populations found in Peter et al. (2018) (rows). Analysis was carried out on haplotypes that were divided into 10 kbp windows. For each of the 17 strains with phased haplotypes, and each 10 kbp window, the sequences of the strain’s haplotypes were compared to those of the 53 reference genomes by calculating the pairwise genetic distance. For each window, the haplotype with the smallest distance was retained. Heatmap squares are coloured based on the genome-wide average minimum pairwise distance observed between each phased and the reference strain. Hierarchical clustering was performed on the columns.

### Beer fermentation and metabolite analysis

While genomic analyses revealed that the studied landrace strains were genetically distinct from traditional brewing strains, we also wanted to test whether they were phenotypically different in beer fermentations. To test this, small-scale fermentations in a standard pale ale wort (100% Maris otter pale ale barley malt hopped with Cascade hops) were carried out with a selection of 21 landrace and five brewing strains. Fermentations were conducted at 25°C which was a compromise between the temperature optima typical of ale yeast fermentation (∼16-25°C) and farmhouse yeast fermentation (∼28-40°C). Overall, the majority of the tested landrace strains could efficiently ferment the wort, reaching final specific gravities as low as the brewing controls (Figure 6A; Supplementary Figure S9). The highest fermentation rates were observed among the landrace strains, with the ‘European Farmhouse’ strains Rakstins 1, Hornindal 1, and Skare 1 among the top performers. Interestingly, a number of ‘European Farmhouse’ strains in the inland Norway subgroup were also among the bottom performers, including Marem 1, Skrindo 2 and Skrindo 5. Overall, no statistical difference (*p* > 0.05; one-way ANOVA) was observed for fermentation rate between the different groups of strains (Figure 6B). Six landrace strains, including the three previously mentioned slow ‘European Farmhouse’ strains, the two Ghanaian strains, and Pundurs 1, produced beer with higher final gravity, suggesting inability to use either of the main wort sugars maltose or maltotriose. Indeed, these beers contained either residual maltose or maltotriose (Figure 6A).

**Figure 6.**
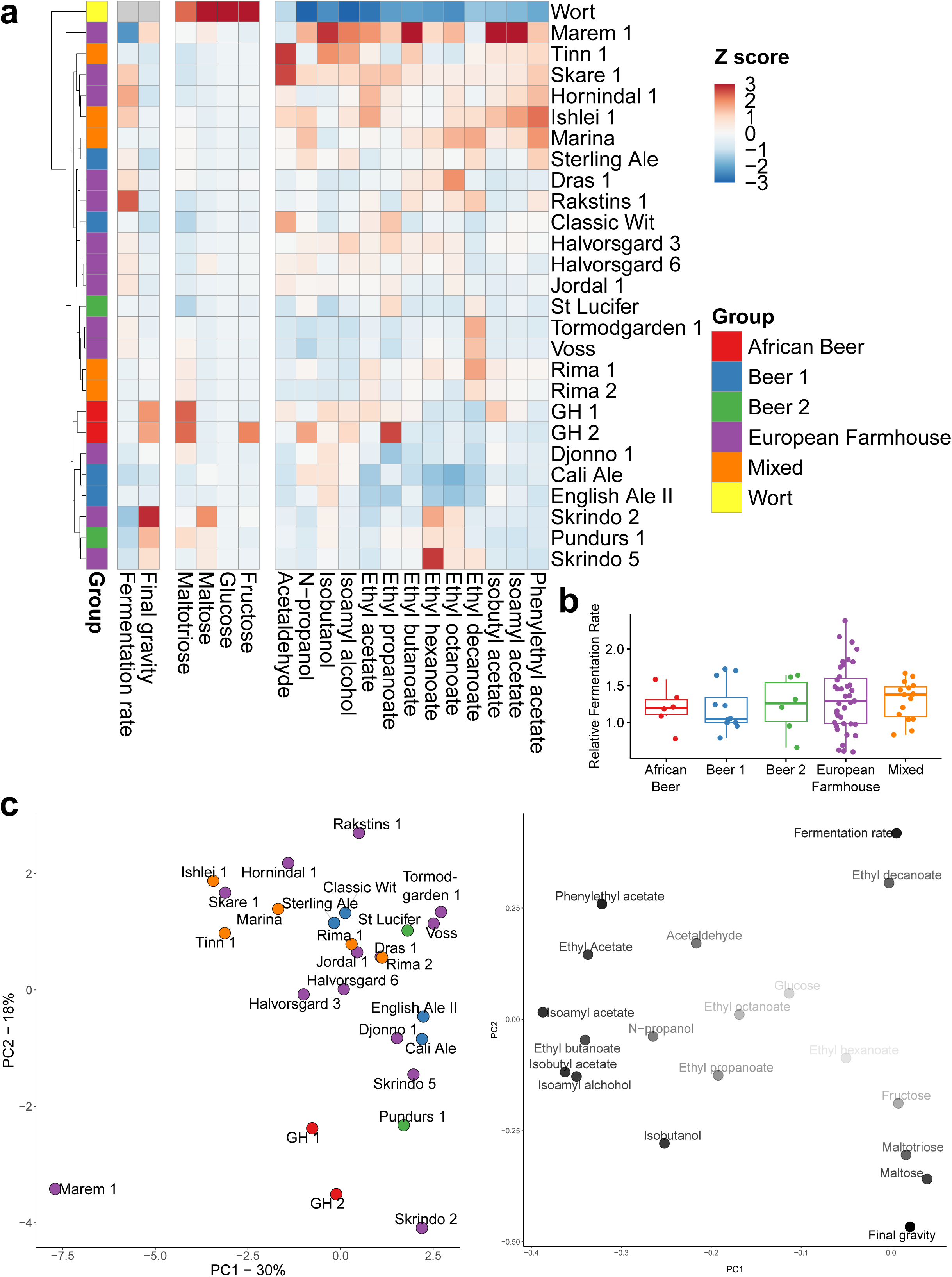
Beer fermentation and metabolite analysis reveals fermentation and flavour production opportunities among landrace yeasts. **(A)** A heatmap showing phenotypic diversity in fermentation performance, sugar consumption and flavour metabolite formation among the 26 landrace and control brewing strains included in the phenotypic analysis. The heatmap is coloured based on Z-scores, and rows are sorted by hierarchical clustering. **(B)** The average relative fermentation rate among the strains grouped by population. No statistical difference (*p* > 0.05; one-way ANOVA) was observed between the populations, but the ‘European Farmhouse’ group showed greatest variation. **(C)** Principal component analysis of the set of 19 phenotypic traits. Scores are shown to the left (strains coloured based on population), while loadings are shown to the right (opacity of the phenotypic traits increases with distance from origin).

Similarly to the fermentation rate, considerable variation in volatile aroma production was also observed within the landrace strains, with strains both among the top and bottom producers of higher alcohols and esters (Figure 6A). Compared to the control brewing strains, the studied landrace strains showed larger diversity for fermentation performance and aroma compound formation (Figure 6C). In regards to acetate ester production, the top producers included ‘European Farmhouse’ and ‘Mixed’ strains, such as Marem 1, Ishlei 1, Tinn 1, Skare 1, Hornindal 1, and Marina (Supplementary Figure S10). Marem 1, in particular, produced beers with high levels of flavour-active esters and was a clear outlier among the strains (Figure 6C). The lowest formation of esters, in contrast, was observed among the ‘Beer 1’ strains Cali Ale and English Ale II. Overall, total ester formation was significantly higher (*p* < 0.05; one-way ANOVA) among the strains from the ‘Mixed’ and ‘European Farmhouse’ groups, and lowest for those from the ‘Beer 1’ and ‘African Beer’ groups (Supplementary Figure S11).

### Linking genotype with phenotype

The small-scale wort fermentations revealed that a number of the landrace strains did not reach as high fermentation yields as the brewing strain controls, and that this was due to residual maltose and maltotriose (Figure 6). Indeed, the beers produced with the brewing controls had significantly lower concentrations of maltotriose than those produced with the landrace strains (*p* = 0.038; unpaired two-tailed t-test). The decreased use of maltotriose can be attributed mainly to the presence of up to three different loss-of-function mutations in the main maltotriose permease, *MAL11*, in many of the landrace strains (Figure 7A). In addition, strains like Djonno_1, Voss_1, Marina and the two Ghanaian strains, showed lower copy numbers or complete lack of *MAL11.* Nevertheless, most landrace strains fermented beer wort, and particularly maltose, efficiently, which could be expected based on their history of use. Among the whole set of 105 strains included in the genomics analysis, strains in the ‘Beer 1’, ‘European Farmhouse’ and ‘Mixed origin’ groups showed the highest copy numbers *MAL11, MAL31, IMA2, IMA3,* and *IMA5* (Supplementary Figure S12).

**Figure 7.**
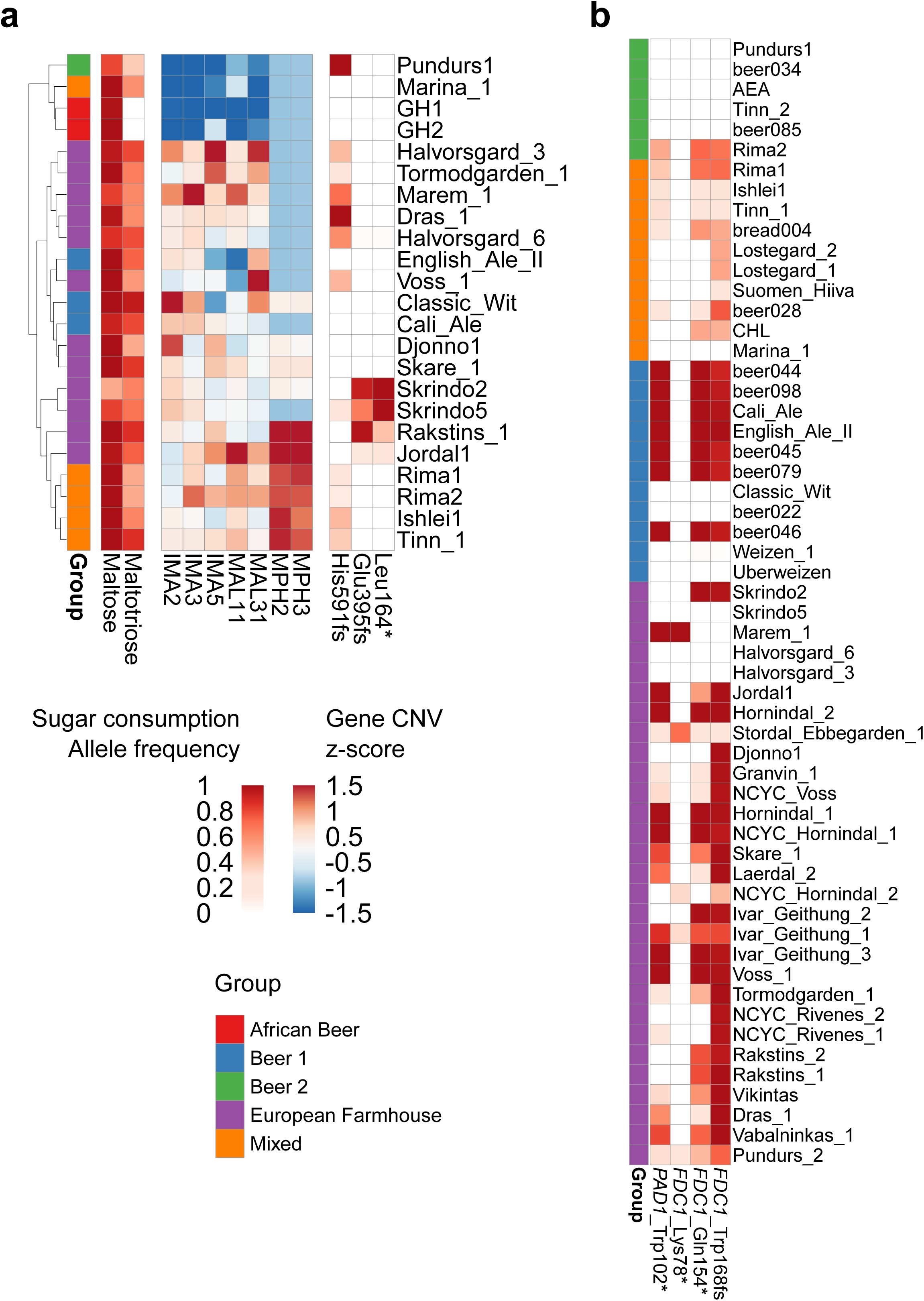
**(A)** The heatmap depicts sugar consumption, copy numbers of genes involved in maltose and maltotriose transport and metabolism, and allele frequencies of inactivating mutations in *MAL11* among the phenotyped strains. **(B)** The allele frequencies of inactivating mutations in *PAD1* and *FDC1* among the strains belonging to the ‘Beer 1’, ‘Beer 2’, ‘Mixed origin’ and ‘European Farmhouse’ populations. Strains are ordered as in Figure 1B.

The landrace strains also showed significant variation in volatile aroma production, with strains both among the top and bottom producers of higher alcohols and esters. In regards to acetate ester production, the top producers included Marem 1, Ishlei 1, Tinn 1, Skare 1, Hornindal 1, and Marina (Figure 6C and Supplementary Figure 11). Of these strains, those belonging to the Mixed population contained a heterozygous frameshift mutation in *FAS2* (Lys1857fs). Mutations in *FAS2* have previously been linked to altered ester formation (Aritomi et al. 2004; De Carvalho et al. 2017). Interestingly, the *FAS2*^BTC.1D^ allele described in De Carvalho et al. (2017), which was shown to result in enhanced acetate ester formation, was also found as heterozygous in Marem 1. In addition, a homozygous loss of stop codon was observed in Marem 1 for *ATF1*, coding for the main enzyme involved in acetate ester biosynthesis. The mutation results in an amino acid sequence that extends for an additional 6 amino acids. This same mutation was also heterozygous in three other landrace strains (Rakstins 1, Skare 1, and Voss 1), while homozygous in the Hefeweizen strains Weizen 1 and Uberweizen. The mutation was also present in the three Hefeweizen strains (Beer072, Beer074, and Beer093) included in the study by Gallone et al. (2016), and only in a single ‘Mosaic Region 3’ strain from the 1,011 yeast genomes set (Peter et al. 2018). While Weizen 1 and Uberweizen were not included in the small-scale fermentations performed here, Hefeweizen strains are known for high acetate ester formation (Schneiderbanger et al. 2016; Meier-Dörnberg et al. 2017). In the study by Gallone et al. (2016), for example, Beer093 was among the top producers of acetate esters. The effect of this extended *ATF1* enzyme on acetate ester formation should therefore be tested in subsequent studies.

The inability to produce 4-vinyl guaiacol from ferulic acid (phenolic off-flavour negative; POF-) is typically considered a domestication marker in *S. cerevisiae* (Gallone et al. 2016; Gonçalves et al. 2016). This trait is common particularly among strains in the ‘Beer 1’ population, as 4-vinyl guaiacol is undesired in many beer styles. The POF-phenotype is due to loss-of-function mutations in either *PAD1* or *FDC1*. While the 4-vinyl guaiacol concentrations were not here analysed from the fermentations, the majority of the studied landrace strains contained loss-of-function mutations in either of these genes (Figure 7B). The POF-landrace strains were mainly found in the ‘European Farmhouse’ population. However, not all ‘European Farmhouse’ strains were POF-, with the two Halvorsgard strains and Skrindo_5 lacking inactivating mutations. Among the strains in the Mixed population, inactivating mutations were common, but heterozygous, indicating the strains were not POF-. Outside the European brewing strains, the POF-phenotype was observed among the ‘African Beer’ (AEN) and ‘Mosaic Region 3’ strains (AEE, AEL, ARA, and BTB) as well. However, no inactivating mutations were observed in the two Ghanaian strains GH1 and GH2 isolated here. Marem 1, Stordal Ebbegarden 1, Ivar_Geithung_1, NCYC_Hornindal_2, and Pundurs 2 contained loss-of-function mutations distinct from those in the ‘Beer 1’ and ‘Mixed origin’ populations.

## Discussion

Investigating the potential application of diverse yeasts is important for the brewing industry to meet an evolving market and shifting consumer preferences. Our study provides new insights into the genetic diversity and characteristics of landrace yeasts, highlighting their potential to offer new combinations of flavour and fermentation traits relevant to modern brewing practices. Here, we extend prior work on discovering, documenting, and characterizing landrace brewing yeasts and brewing practices from traditional farmhouse brewers.

Of the landrace yeasts studied here, the Norwegian ‘kveik’ strains have received the most interest in the brewing industry, with several commercial yeast suppliers now offering such strains in their selection. When these strains were first phenotypically and genetically characterized, it was revealed that they possess many traits desirable for beer production (e.g. maltose and maltotriose use, flocculation and lack of phenolic off-flavour formation), yet they were genetically distinct from other brewing strains (Preiss et al. 2018; Foster et al. 2022). Here, we show that genetically distinct farmhouse brewing yeasts are not restricted only to Norway, but rather, related strains were also isolated from farmhouse breweries in the Baltic region. Hence, we propose to refer to this group of yeast strains as ‘European Farmhouse’ brewing yeast. Inside this group, geographical origin of the strains could also be distinguished based on genome sequence, with strains from western Norway, eastern Norway, and the Baltic region clustering separately based on genome-wide SNPs. The ‘European Farmhouse’ group is closely related to the ‘Beer 1’ group of brewing strains, and these groups have likely branched off and evolved in parallel before the predicted emergence of the ‘Beer 1’ branch in the 16th century (Gallone et al. 2019). In comparison with the ‘Beer 1’ strains, the ‘European Farmhouse’ strains have not been used industrially, nor undergone any pure culture isolations to our knowledge.

While the ‘European Farmhouse’ strains are closely related to ‘Beer 1’ strains, they also show more signs of admixture. In early work, it was hypothesized that the ‘kveik’ strains have an admixed ancestry between a ‘Beer 1’ strain and one possibly of Asian origin. Here, we provide further evidence to support this hypothesis. A recent study, however, argues that the ‘Kveik’ population, rather than being closely related to the ‘Beer 1’ population, forms a sister group to other domesticated populations closer to the root of the *S. cerevisiae* tree (Dondrup et al. 2023). In our study, we observed a similar phenomenon of the ‘kveik’ strains grouping closer to ‘Mosaic Region 3’ and Asian fermentation strains when constructing a SNP-based tree (Figure 1B). However, we believe this is an artifact of the relatively high levels of heterozygosity in, and admixed history of, these strains, which places the ‘European Farmhouse’ branch intermediate of the ‘Mixed origin’-‘Beer 1’ and ‘Asian Fermentation’-‘Sake’ branches (Kopelman et al. 2012; Meirmans et al. 2018; Huang et al. 2022). Firstly, in the assembly-based trees (Figure 2 and Supplementary Figure S4) based on the haploid major allele sequence, the ‘European Farmhouse’ strains form a sister group to the ‘Beer 1’ strains. Secondly, using different algorithms, admixture and migration events from the Asian fermentation strains into a ‘Beer 1’ ancestor was predicted. Thirdly, long-read phasing of the SNPs suggests that the studied ‘European Farmhouse’ strains show lowest genetic distance to the ‘Beer 1’ and ‘Mixed origin’ strains out of the major populations identified in Peter et al. (2018). But, compared to the ‘Beer 1’ and ‘Mixed origin’ strains, the ‘European Farmhouse’ strains also show lower distance to the Asian fermentation strains supporting admixture with the latter. Recent studies on ale yeast, have revealed that the ‘Beer 1’ ancestor in itself was also derived from admixture between close relatives of European wine and Asian fermentations strains (Fay et al. 2019; Abou Saada et al. 2022). The ‘European Farmhouse’ strains, in comparison, show an even higher contribution of Asian fermentation alleles. It is unclear why gene flow from the Asian fermentation strains is beneficial in the beer wort environment; however, many domesticated Asian strains show duplications of genes related to maltose metabolism (Duan et al. 2018) and carry the *RTM1* cluster which confers resistance to molasses toxicity (Borneman et al. 2016; Pontes et al. 2020). High copy number of the *MAL* genes is also characteristic of the ‘Beer 1’ and ‘European Farmhouse’ populations, while the *RTM1* cluster is also present in the ‘Beer 1’, ‘Mixed origin’, and ‘European Farmhouse’ strains (Supplementary Figure S13 and S14).

These results led us to wonder how the repeated admixture of Asian fermentation genetics into brewing strain groups have happened historically. While purely speculative, there has been influence of beer brewing in the Middle East and Georgia, eliminated by Islam some time in the 12th century (Haider 2013). Brewing continued in Georgia, and it is possible that yeast may have spread north of the Caucasus into Russia and beyond either before or shortly after 700 CE. Known viking activity in present-day Russia and Ukraine between 700 CE until at least 1100 CE (Roesdahl 2016; Jarman 2021) could have influenced Scandinavian brewing culture, methods, and beer yeast. Additionally, we note that the Rakstins_1 and Dras_1 strains appear to be genetically ‘European Farmhouse’ yeasts, despite being isolated from sources in the Baltic region. Similarly, the Norway East strains appear to form their own sub-group within the ‘European Farmhouse’ population that appears to span both the north and south regions. The two yeast types referred to as ‘gong’ and ‘berm’ thus appear to be a single group. Unsurprisingly, the Norway East strains clustered next to the kveik strains from Norway West, suggesting that the strains from Norway East form an outgroup of kveik. Further investigation is required to analyze more yeast sources and isolates from traditional brewers in the Baltic region in order to understand whether the occurrence of yeasts from the ‘European Farmhouse’ group is common among Baltic farmhouse brewing yeasts. It is also likely that similar strains have in the past been used for farmhouse brewing in Finland, Sweden, and Denmark. However, in Finland, for example, the farmhouse beer sahti is still commonly brewed, but brewers transitioned to commercial baker’s yeast during the first half of the 20th century (Räsänen 1975). The same transition took place in the brewing of the Swedish farmhouse ale gotlandsdricke (Salomonsson 1979).

Limited work has so far been carried out on the phenotypic diversity of farmhouse yeasts. Prior work on a relatively small subset of ‘kveik’ isolates has revealed that they tend to ferment wort rapidly, have high thermotolerance, and can produce beers ranging from clean and neutral to strongly aromatic (Preiss et al. 2018; Foster et al. 2022; Kawa-Rygielska et al. 2022; Habschied et al. 2022; Dippel et al. 2022). Here, we reveal that farmhouse yeast, including kveik, show significant phenotypic diversity, both in regards to fermentation dynamics and flavour formation. In our screen of farmhouse strains and industrial controls, we observed both the highest and lowest maximum fermentation rates from strains belonging to the ‘European Farmhouse’ population. So, while kveik yeast are generally considered rapid fermenters, we here reveal that this is not always the case. Many of the landrace strains belonging to the ‘Mixed’ population also showed rapid fermentation. This is anticipated, as the ‘Mixed’ population consists of strains isolated from breweries, bakeries and distilleries, and tend to have the ability to consume maltotriose (Gallone et al. 2016; Peter et al. 2018). The landrace strains showing poor fermentation were unable to completely use the maltose and maltotriose in the wort naturally limiting ethanol formation, and this appeared linked to lower copy numbers of or non-functional α-glucoside permeases. While the brewing industry has traditionally been interested in fast-fermenting strains with high ethanol yields, there is also an increasing interest towards strains with limited maltose and maltotriose use, as the consumer demand for beer with low and no alcohol content continues to rise (Bellut and Arendt 2019). Landrace strains therefore show potential for many different applications within the brewing industry.

In addition to showing variation in fermentation dynamics, the studied landrace strains showed considerable variation in aroma formation. Farmhouse ales can also have very different flavour profiles, with traditional European farmhouse ales usually being ester-rich and sometimes phenolic, while some kveik strains are known for their ability to produce ‘clean’ flavour profiles even at high fermentation temperatures. Similarly to the fermentation dynamics, we here show that strains from the ‘Kveik’ population were both among the top and bottom producers of higher alcohols and esters, meaning they have potential for producing both cleaner ‘lager-style’ and fruitier ‘pale ale-style’ beers. In contrast with previous work (Preiss et al. 2018), we here also reveal that not all strains in the ‘Kveik’ population are POF-, and that the non-kveik landrace strains tended to be POF+. The vast majority of the POF-strains in the ‘Beer 1’, ‘Mixed origin’ and ‘European Farmhouse’ populations carry the same three inactivating mutations in *PAD1* and *FDC1*, as also highlighted in other recent studies (Gallone et al. 2016; Gonçalves et al. 2016; Abou Saada et al. 2022). This supports that these mutations have been acquired by an early ancestor of the ‘Mixed origin’-‘European Farmhouse’-‘Beer 1’ branch. Interestingly, some ‘European Farmhouse’ strains carry a premature stop codon in *FDC1* (Lys78*) that is not found in any other ‘Beer 1’ or ‘Mixed origin’ strains, supporting a potential independent domestication for a sub-group of the European farmhouse strains. The unique inactivating mutations in the POF-‘African Beer’ strains similarly imply an independent domestication for the phenotype in these strains (Abou Saada et al. 2022).

In addition to the direct applied value of landrace brewing strains, they are also valuable from a strain development perspective. Compared to ‘Beer 1’ strains, which tend to be sterile (Gallone et al. 2016; De Chiara et al. 2022), the majority of the kveik strains that have been tested so far have been fertile (Preiss et al. 2018; Dippel et al. 2022). This means that kveik strains can be quite readily used for breeding. The fertility of the non-kveik landrace strains has not been studied, but they are also likely to be fertile, as strains from the ‘Mixed origin’, ‘Mosaic beer’, and ‘African beer’ populations typically are (De Chiara et al. 2022). Landrace strains also tend to be polyploid (here the majority were tetraploid), which makes them good candidates for strain improvement through adaptive evolution. Previous studies have revealed that polyploid strains tend to adapt faster and undergo larger genomic changes during adaptive evolution compared to diploid strains (Selmecki et al. 2015; Lu et al. 2016). Indeed, the technique has been used to enhance ethanol tolerance and sugar utilization of brewing yeast (Brickwedde et al. 2017; Krogerus et al. 2018a). Similar strategies could be applied to further improve the landrace strains; while many of them showed rapid fermentation, some showed incomplete maltose or maltotriose use as a result of mutations in *MAL11*. Finally, from a strain development perspective, the genomes of the landrace strains can provide new insights and targets for strain engineering. Several kveik strains, for example, have been shown to have enhanced thermotolerance compared to traditional brewing strains, which in turn appears to be linked to enhanced intracellular trehalose accumulation (Foster et al. 2022). The molecular mechanisms responsible for this phenotype have not yet been identified, but several potential causative single nucleotide polymorphisms in genes related to trehalose metabolism have been identified in kveik strains that could be tested. Similarly, targets for other brewing-relevant phenotypes, such as altered aroma formation, could be identified from the genomes of the landrace strains.

In summary, we found genetically distinct and phenotypically useful yeast strains among various landrace brewing yeast cultures used to ferment traditional farmhouse beers. This study helps understand the genetic history of kveik and related landrace yeasts by suggesting that ‘European Farmhouse’ is a sister group to ‘Beer 1’. We further found evidence for increased Asian admixture in the ‘European Farmhouse’ yeast lineage. It remains unknown exactly how modern beer yeasts as well as landrace yeasts such as kveik have evolved over time. Further studies may explore a larger sample set of landrace yeast isolates or populations from the >60 farmhouse yeast cultures already identified (Garshol 2020c). The unique genetics of such yeasts may yield new technological benefits to the brewing, baking, or biofuel industries including traits such as stress resistance, alcohol tolerance, and flavour/metabolite production. Newly developing and growing industries such as synthetic biology and cellular agriculture may also benefit from the knowledge to develop more robust and metabolically diverse yeasts.

## Declarations

### Funding

Research at University of Guelph was funded by Natural Sciences and Engineering Research Council of Canada Discovery (NSERC #264792-400922), Alliance-GLAAIR Product Development (UG-GLPD-2021-101214), and Ontario Regional Priorities Partnership program (ON-RP3 06) grants. Research at VTT was funded by the Research Council of Finland (Academy Research Fellow 355120).

### Conflicts of interest

Richard Preiss and Eugene Fletcher were employed by Escarpment Laboratories Inc. Kristoffer Krogerus was employed by VTT Technical Research Centre of Finland Ltd. The funders had no role in study design, data collection and analysis, decision to publish, or preparation of the manuscript.

### Availability of data and material

Illumina and basecalled ONT sequencing reads have been deposited in NCBI-SRA under BioProject number PRJNA1018716 (https://www.ncbi.nlm.nih.gov/bioproject/).

### Authors’ contributions

RP: Conceived the study, designed experiments, wrote the manuscript

EF: Conceived the study, designed experiments, performed experiments

LG: Provided yeast strains, wrote the manuscript

BF: Performed data analysis

EO: Performed data analysis

ML: Performed data analysis

GM: Designed experiments, edited the manuscript

KK: Conceived the study, designed experiments, performed experiments, performed data analysis, wrote the manuscript.

All authors read and approved the final manuscript.

## Supplementary Figure and Table legends

**Supplementary Figure S1.** Visualisation of phased haplotigs detected by nPhase in the 17 phased strains.

**Supplementary Figure S2.** Median coverage in 10 kbp windows across the genome of the 105 *S. cerevisiae* strains included in the genomic analysis.

**Supplementary Figure S3.** Allele frequency distributions across the genome of the 105 *S. cerevisiae* strains included in the genomic analysis.

**Supplementary Figure S4.** A neighbour-joining tree generated based on MinHash distances in the 142 assemblies of ScRAP (O’Donnell et al. 2023) and long-read assemblies of 9 landrace and 6 control strains generated here. The branches containing the ‘Mixed origin’, ‘European farmhouse’ and ‘Beer 1’/’Ale beer’ populations are highlighted.

**Supplementary Figure S5.** Maximum likelihood phylogenetic tree based on SNPs at 310688 sites in 1,011 yeast genome strains (Peter et al. 2018), the 35 landrace and five brewing control strains sequenced here, and the five sequenced NCYC ‘kveik’ strains. Strain names of the landrace and control strains are coloured red and blue, respectively. The branch containing the ‘Mixed origin’, ‘European farmhouse’ and ‘Beer 1’/’Ale beer’ populations is highlighted in grey.

**Supplementary Figure S6.** Population structure of a merged set of 35 *S. cerevisiae* landrace brewing strains and the 1,011 yeast genomes strains estimated with ADMIXTURE based on SNPs at 31766 sites. Each strain along the x-axis is represented by a vertical bar partitioned into colors based on estimated membership fractions to the resolved populations for K = 8 to 20 assumed ancestral populations.

**Supplementary Figure S7.** Heatmaps visualising *D, f_4_*-ratio and *f*-branch statistics between populations as calculated with DSuite. (**A**) The *f*-branch statistics between population tree tips and nodes. A warmer colour represents a higher estimated f-branch value, which indicates the proportion of the genome affected by admixture. Grey shading indicates tests that could not be made. The maximum (**B**) *D* and (**C**) *f_4_*-ratio statistics between pairs of populations. A red color indicates a higher value, while a blue color a lower value.

**Supplementary Figure S8.** The network graph topology with highest posterior probability as derived by AdmixtureBayes from the merged data set of 1,011 yeast genome strains (Peter et al. 2018), the 35 landrace and five brewing control strains sequenced here, and the five sequenced NCYC ‘kveik’ strains. The data set converged to and was best described by 20 admixture events. Red squares indicate admixture events, while blue ovals indicate populations.

**Supplementary Figure S9.** Change in specific gravity over time in the wort fermentations with landrace and control brewing strains. Error bars show standard deviation of three replicates.

**Supplementary Figure S10.** Concentration of isoamyl, isobutyl and phenylethyl acetate in the beers fermented with landrace and control brewing strains. Error bars show standard deviation of three replicates. Different letters above bars indicate significant differences by one-way ANOVA (*p* < 0.05).

**Supplementary Figure S11.** Relative total ester formation by group among the landrace and control brewing strains during wort fermentations. Different letters above bars indicate significant differences by one-way ANOVA (*p* < 0.05).

**Supplementary Figure S12.** Average copy number of genes related to maltose and maltotriose transport and metabolism among the landrace strains sequenced here and the 1,011 yeast genome strains (Peter et al. 2018) grouped by population.

**Supplementary Figure S13.** Copy number of *RTM1* among the 1,011 yeast genomes strains (Peter et al. 2018) grouped by population.

**Supplementary Figure S14.** The *RTM1* cluster in the long-read assemblies of the landrace and control brewing strains. *RTM1* is highlighted in red, while *SUC2* is highlighted in blue.

**Supplementary Table S1.** A list of strains included in the genomic analysis, the number of homozygous and heterozygous SNPs detected, and their accession numbers.

**Supplementary Table S2.** Assembly statistics for the 15 long-read assemblies generated here.

## Footnotes

1 Registry accessible online at https://www.garshol.priv.no/download/farmhouse/kveik.html

## Notes

### Competing Interest Statement

The authors have declared no competing interest.

