## Supplementary Figure S1 for "European Farmhouse Brewing Yeasts Form a Distinct Genetic Group"

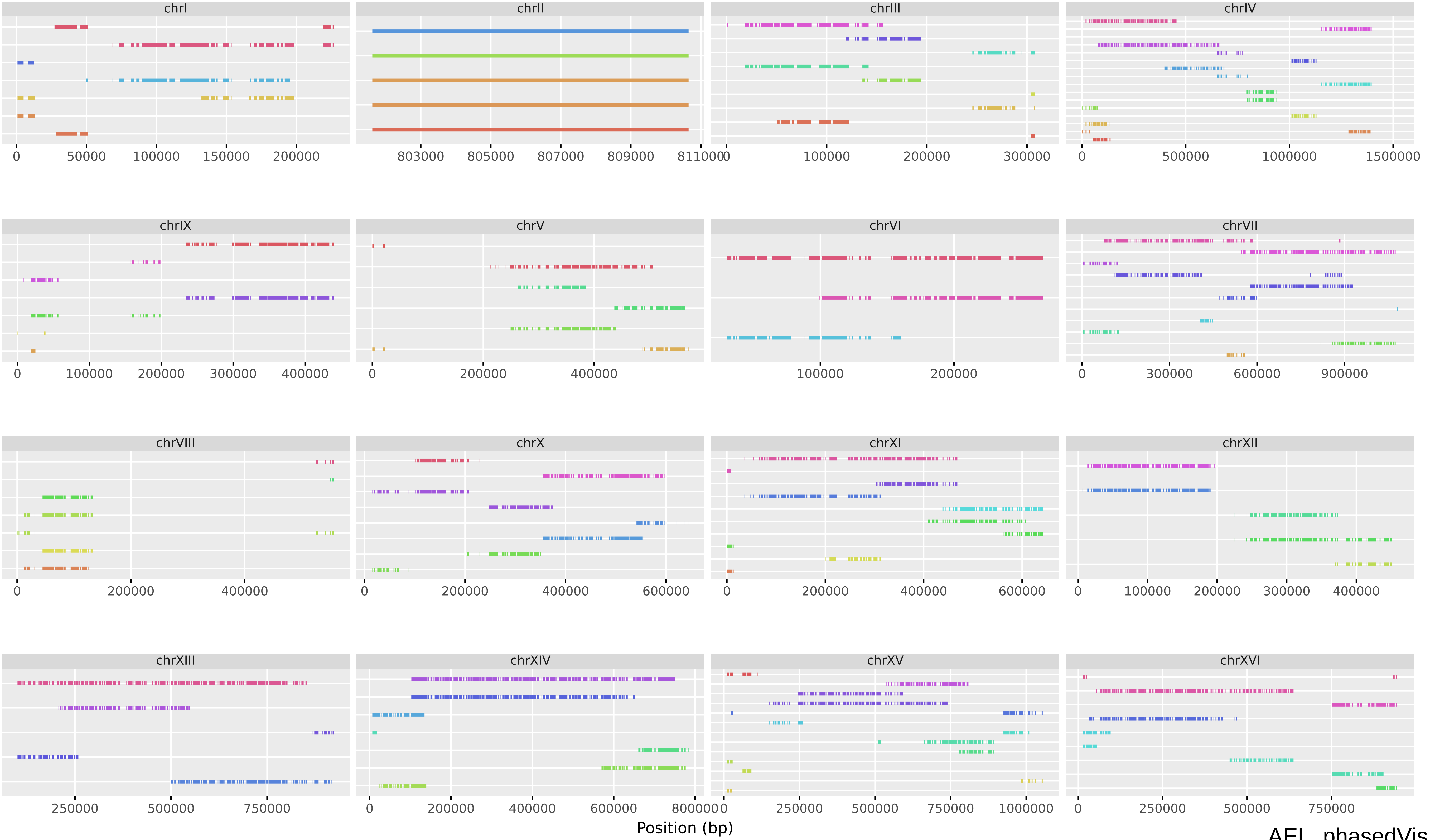


AEL_phasedVis

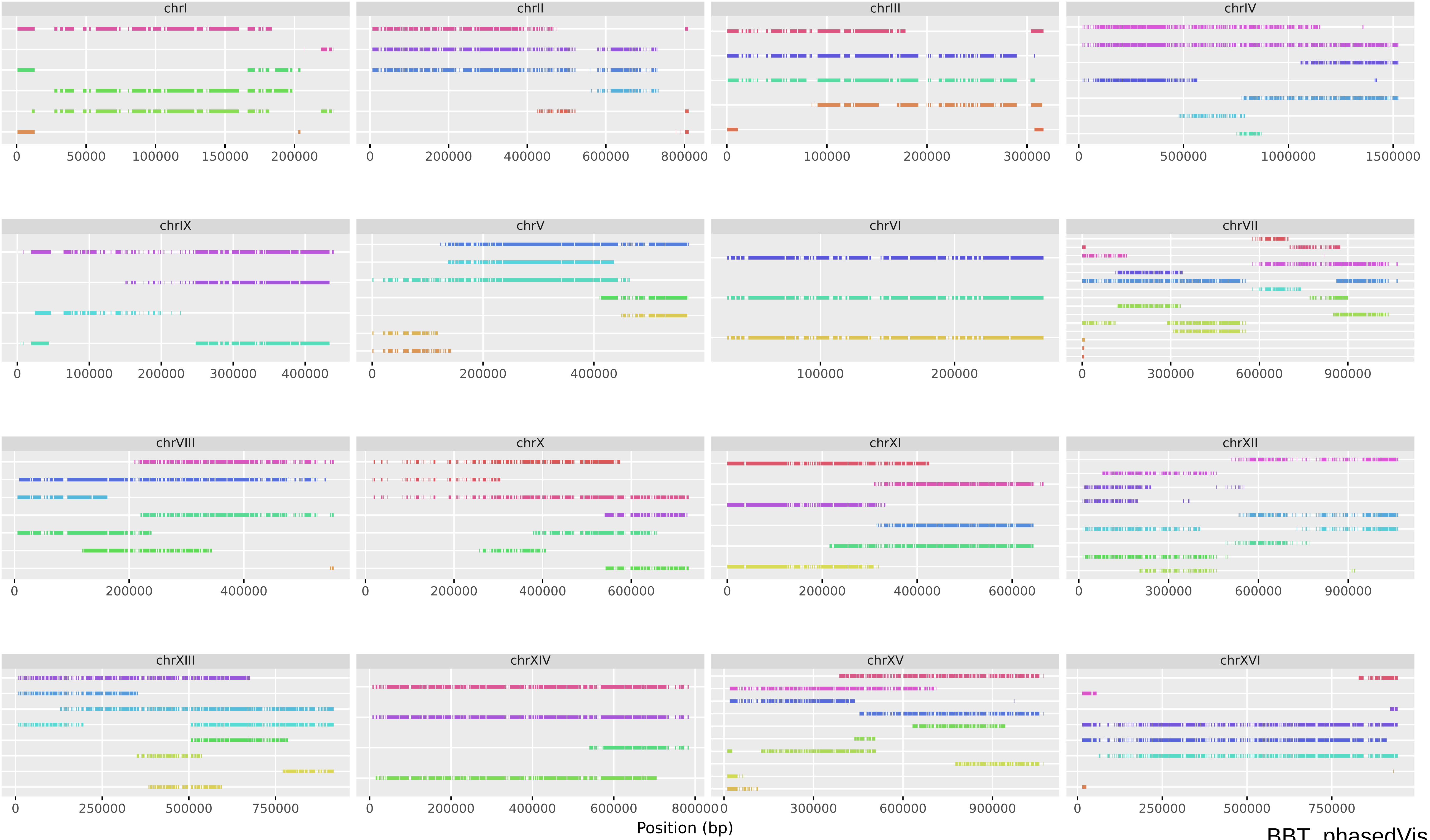


BBT_phasedVis

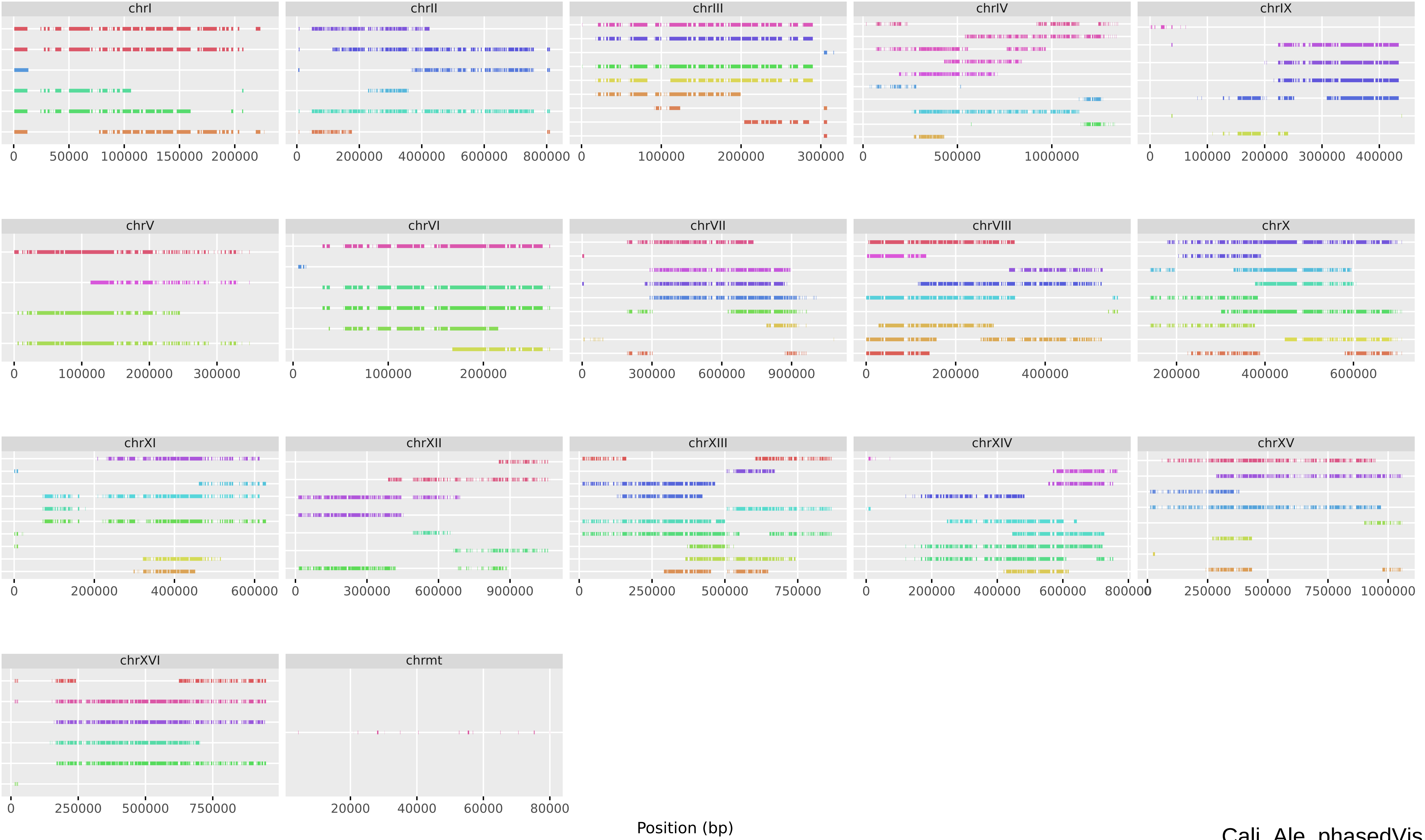


Cali_Ale_phasedVis

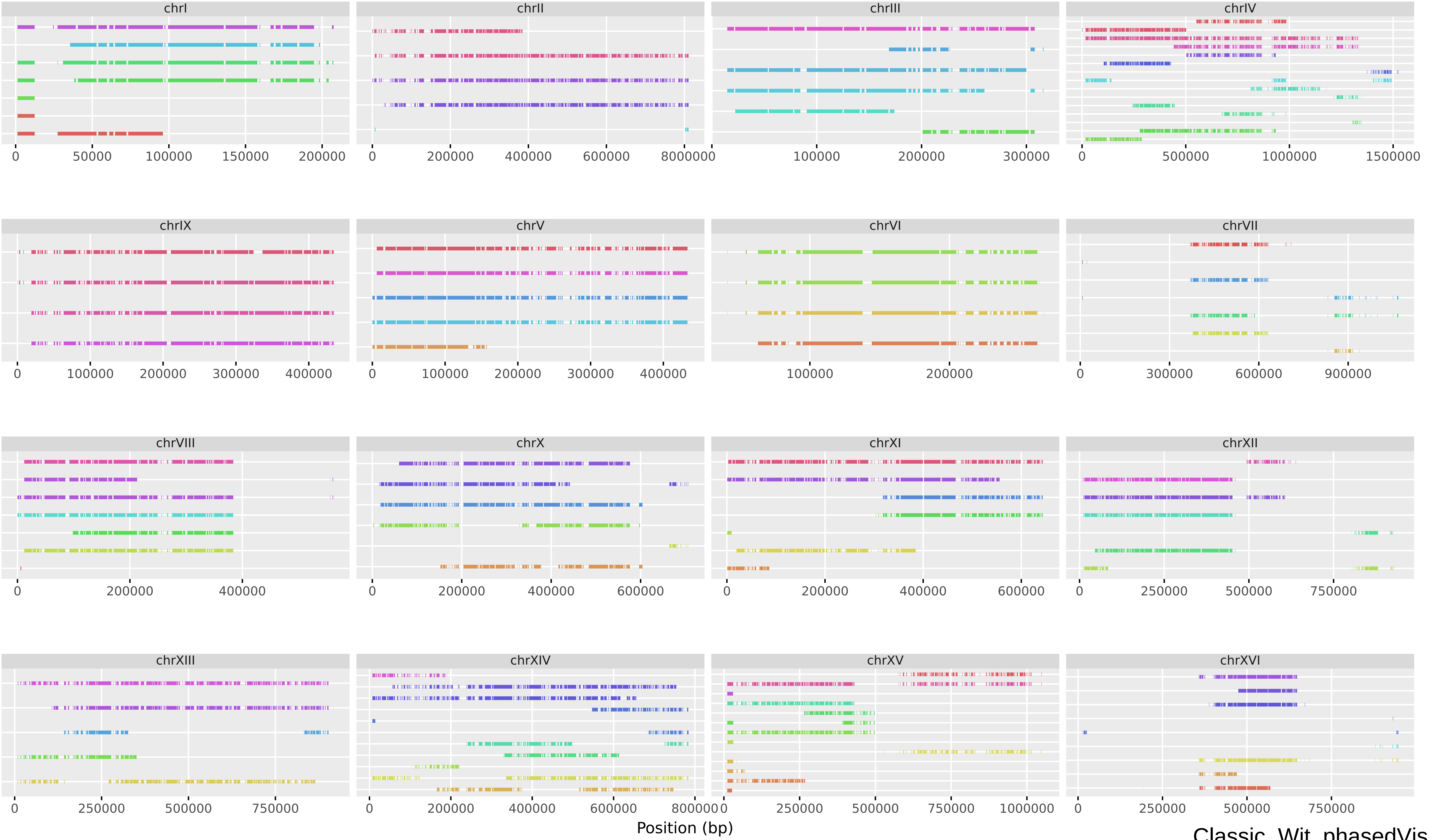

Classic_Wit_phasedVis

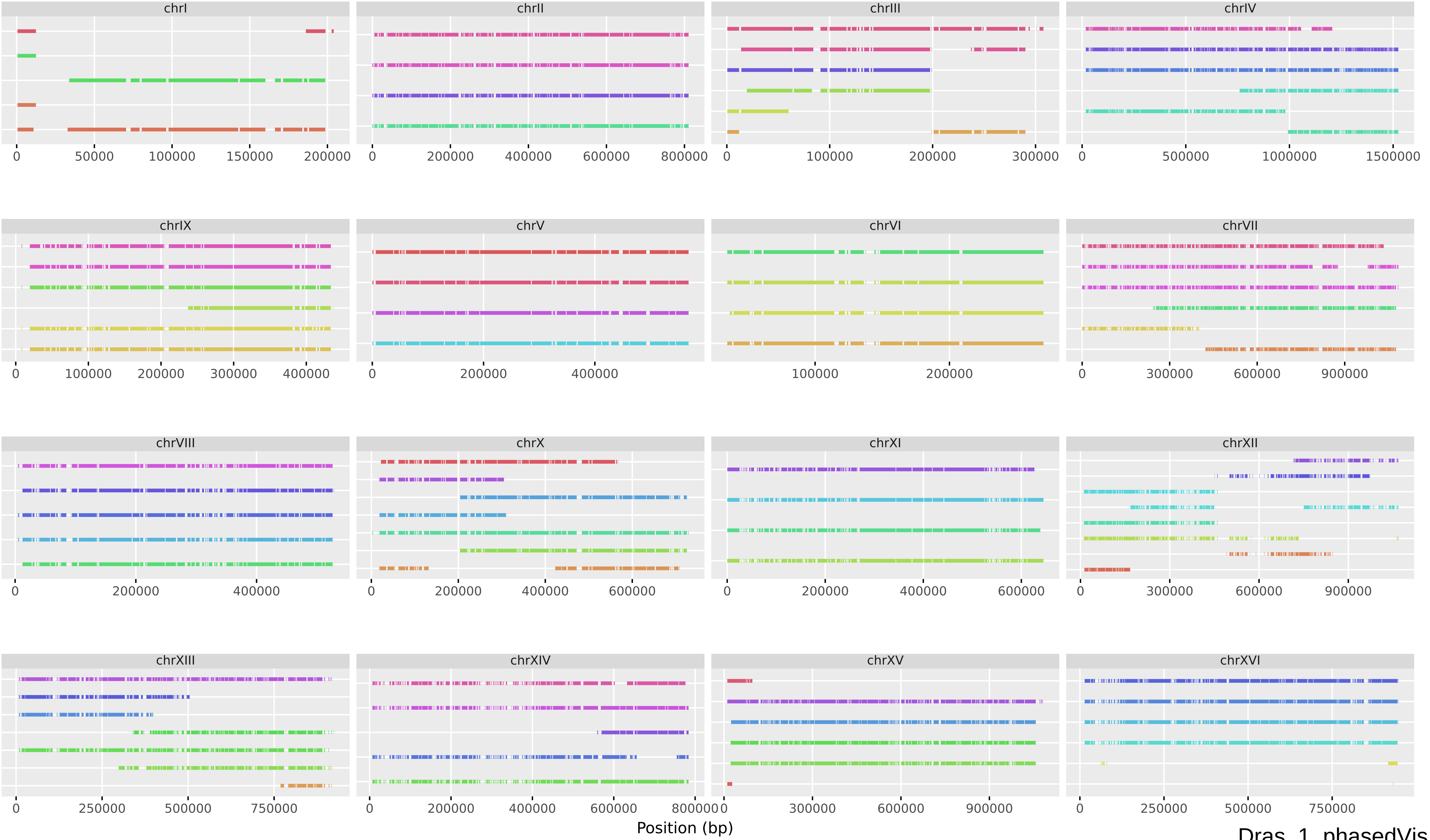


Dras_1_phasedVis

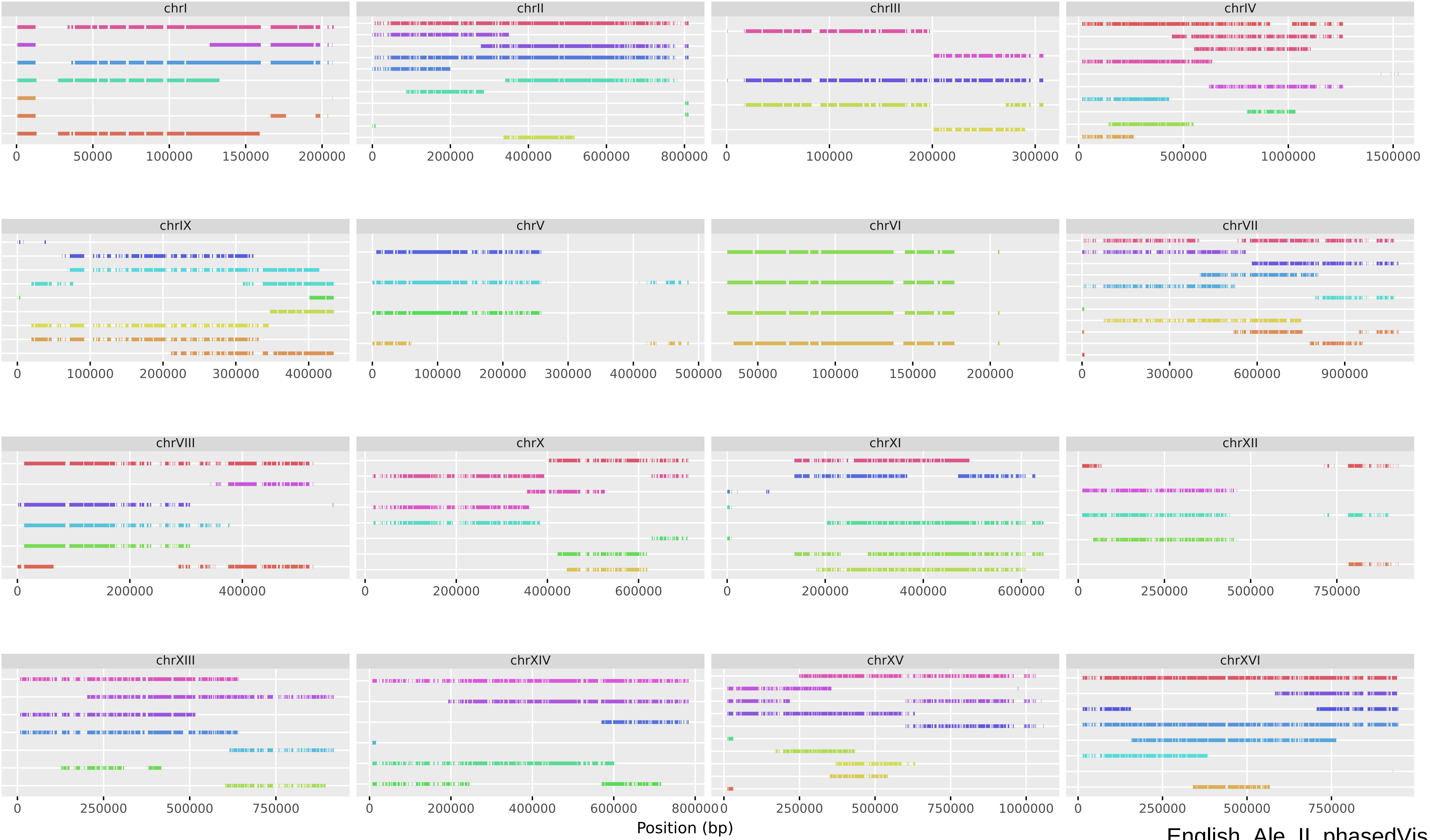


English_Ale_II_phasedVis

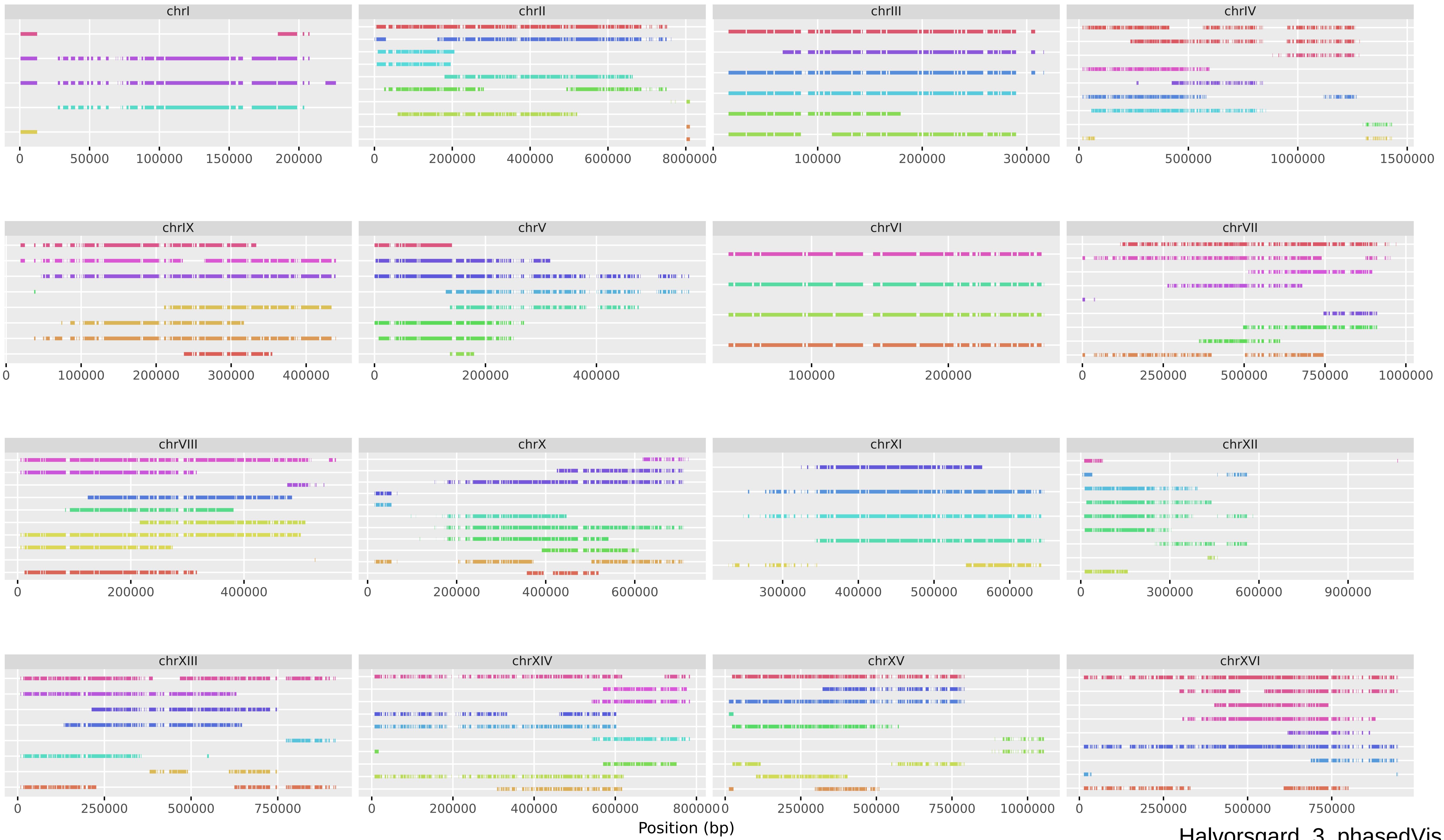


Halvorsgard_3_phasedVis

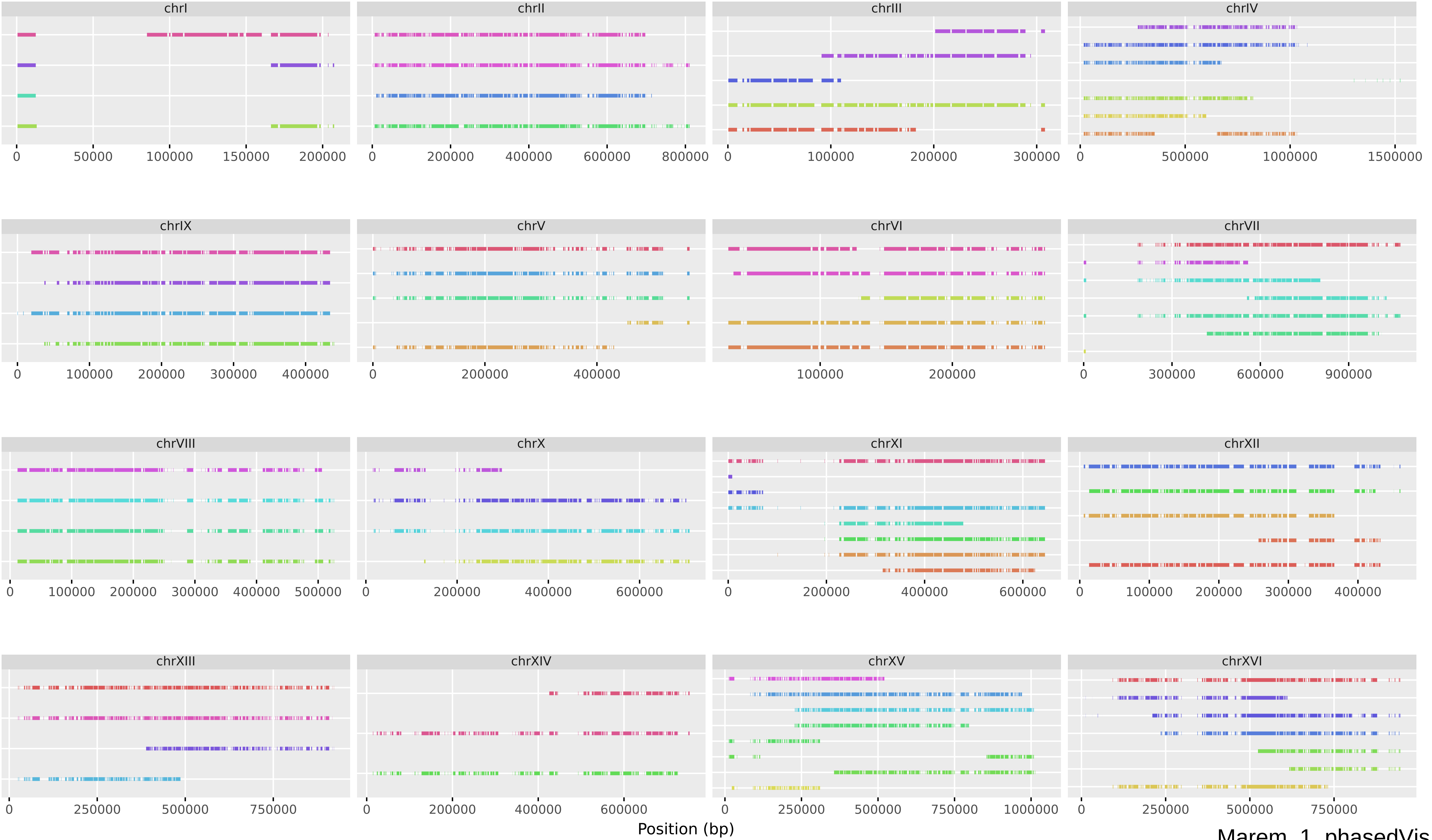


Marem 1 phasedVis

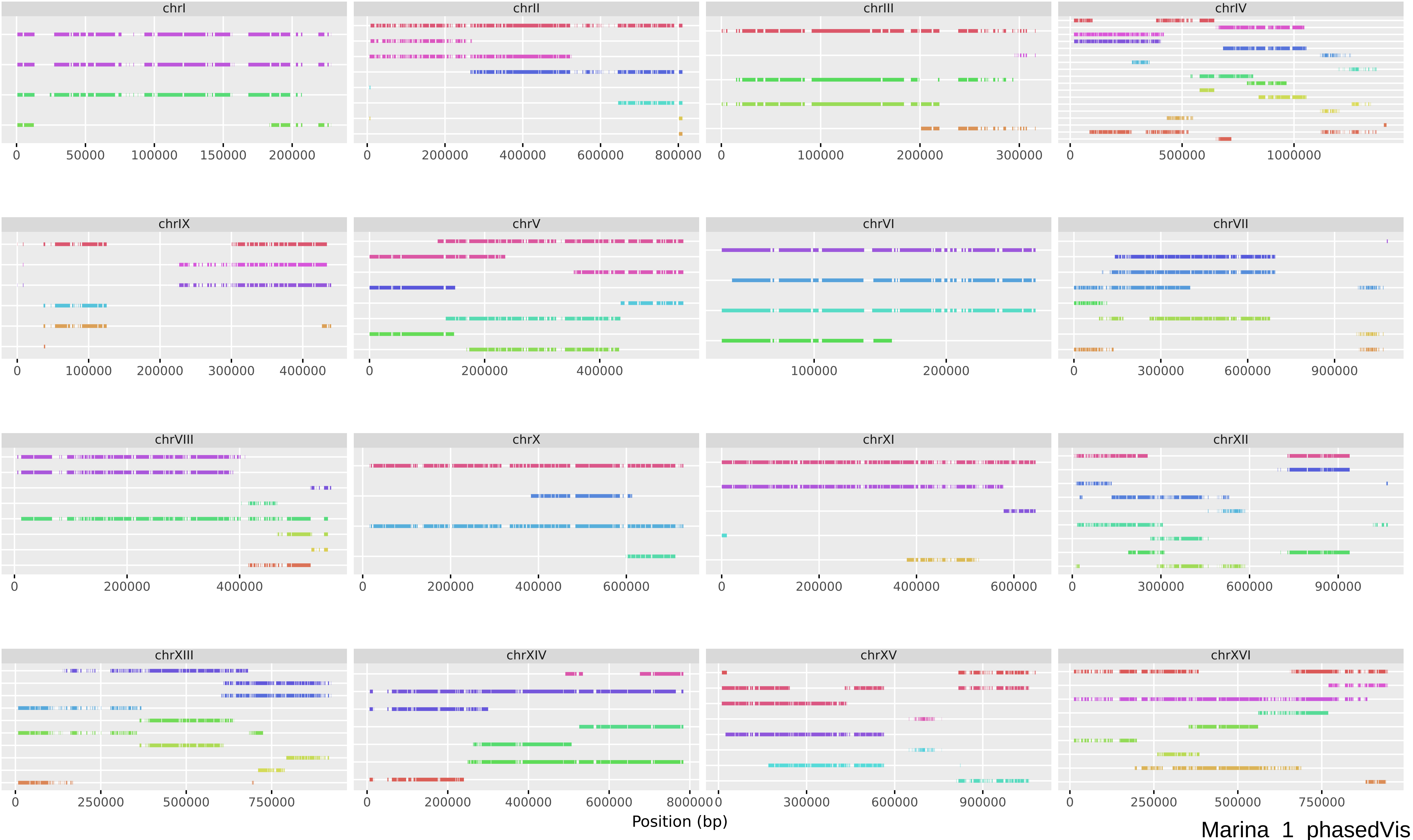

chrI
0 50000 100000 150000 200000
chrII
0 200000 400000 600000 800000
chrIII
0 100000 200000 300000
chrIV
0 500000 1000000
chrIX
0 100000 200000 300000 400000
chrV
0 200000 400000
chrVI
100000 200000
chrVII
0 300000 600000 900000
chrVIII
0 200000 400000
chrX
0 200000 400000 600000
chrXI
0 200000 400000 600000
chrXII
0 300000 600000 900000
chrXIII
0 250000 500000 750000
chrXIV
0 200000 400000 600000 800000
chrXV
0 300000 600000 900000
chrXVI
0 250000 500000 750000
Position (bp)
Marina 1 phasedVis

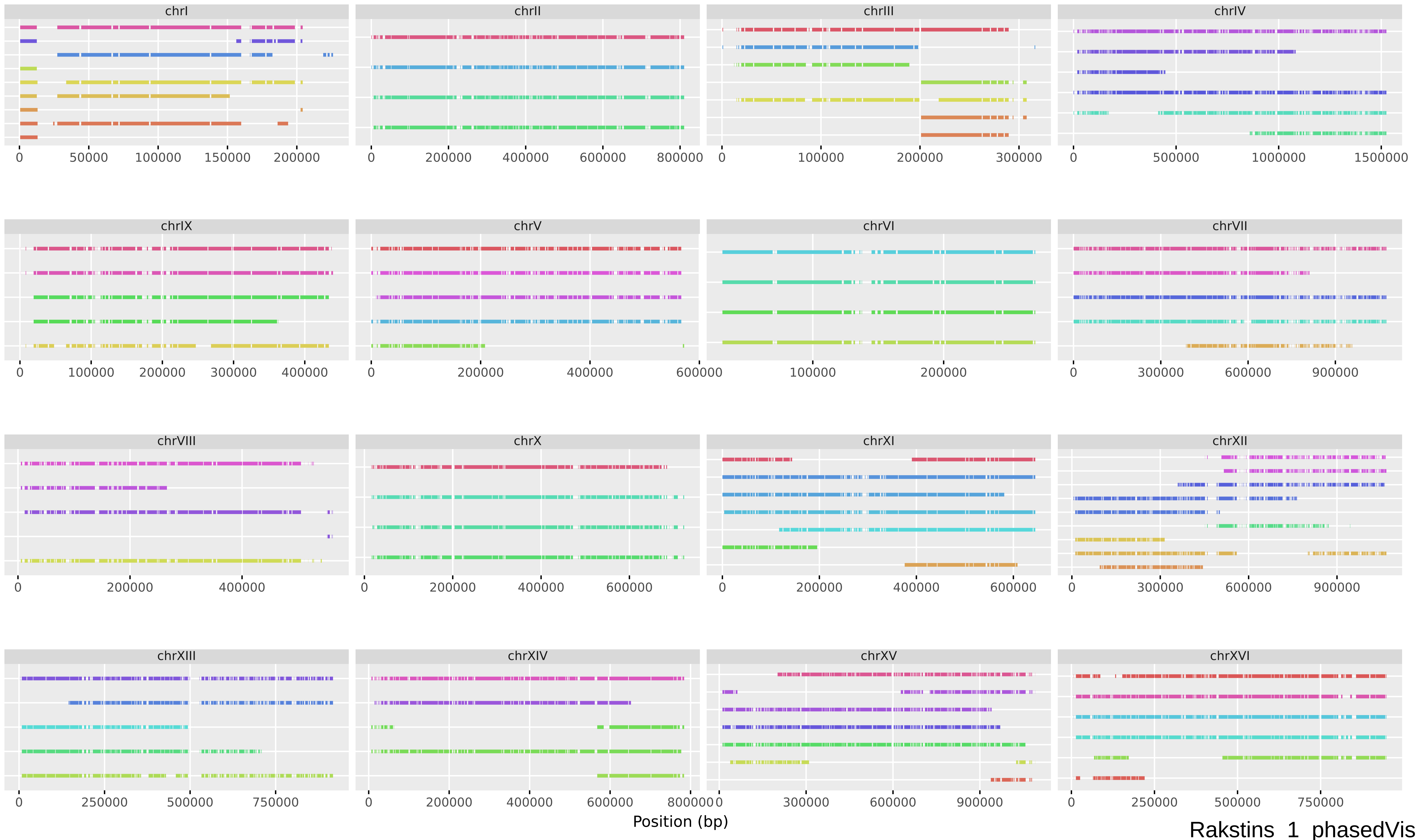
chrI
chrII
chrIII
chrIV
chrIX
chrV
chrVI
chrVII
chrVIII
chrX
chrXI
chrXII
chrXIII
chrXIV
chrXV
chrXVI
Position (bp)
Rakstins_1_phasedVis

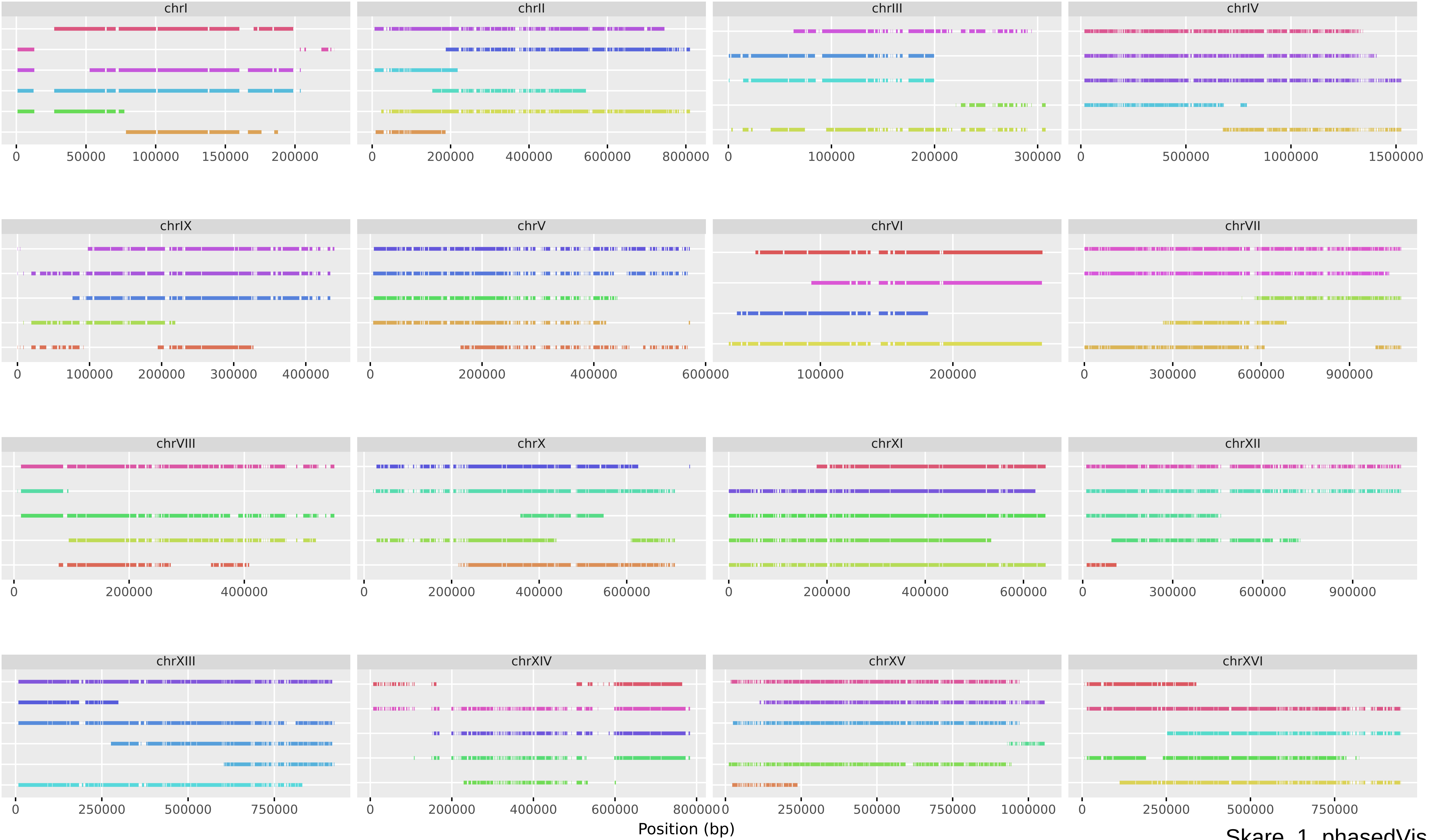


Skare_1_phasedVis

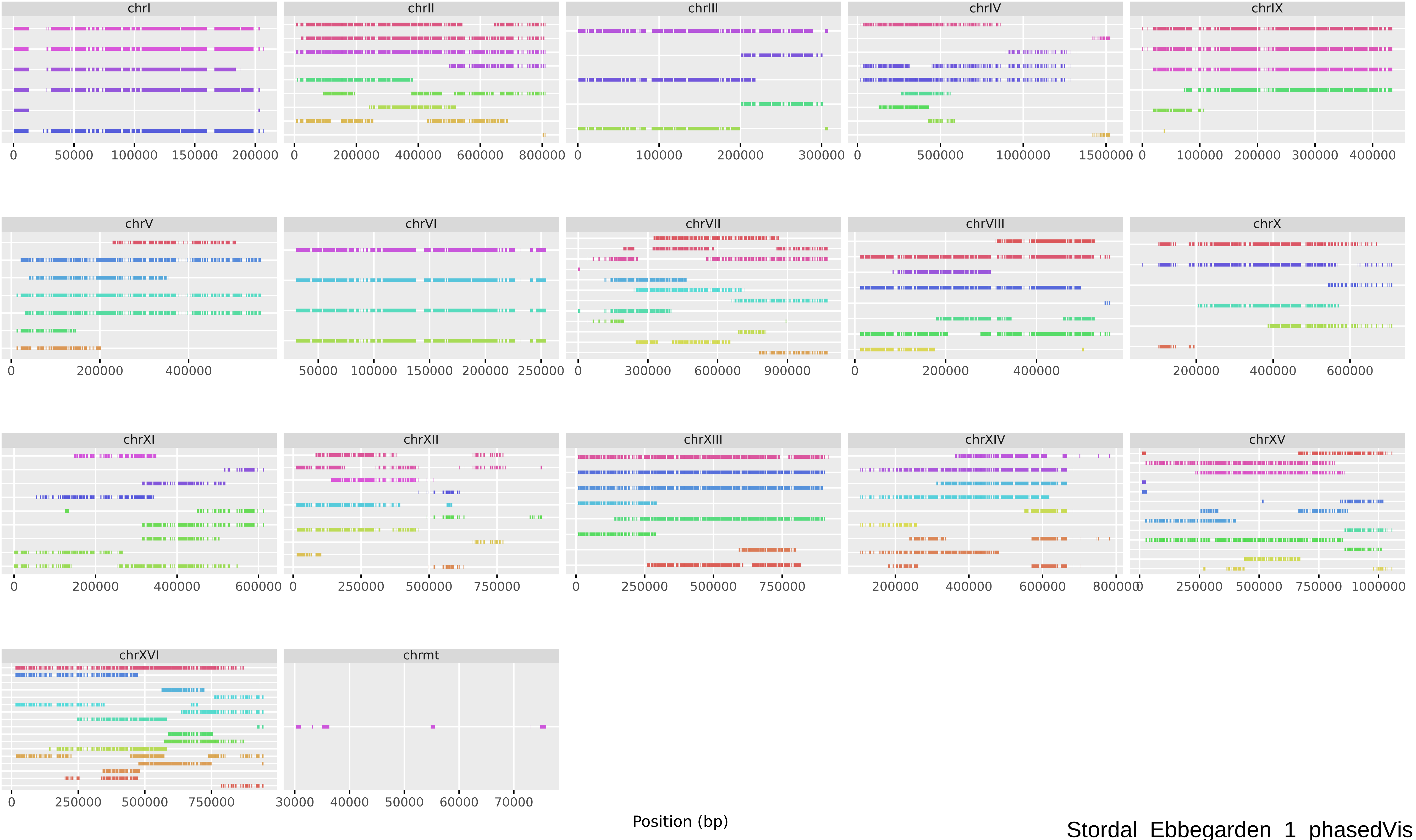
chrI
chrII
chrIII
chrIV
chrIX
chrV
chrVI
chrVII
chrVIII
chrX
chrXI
chrXII
chrXIII
chrXIV
chrXV
chrXVI
chrmt
Position (bp)
Stordal_Ebbegarden_1_phasedVis

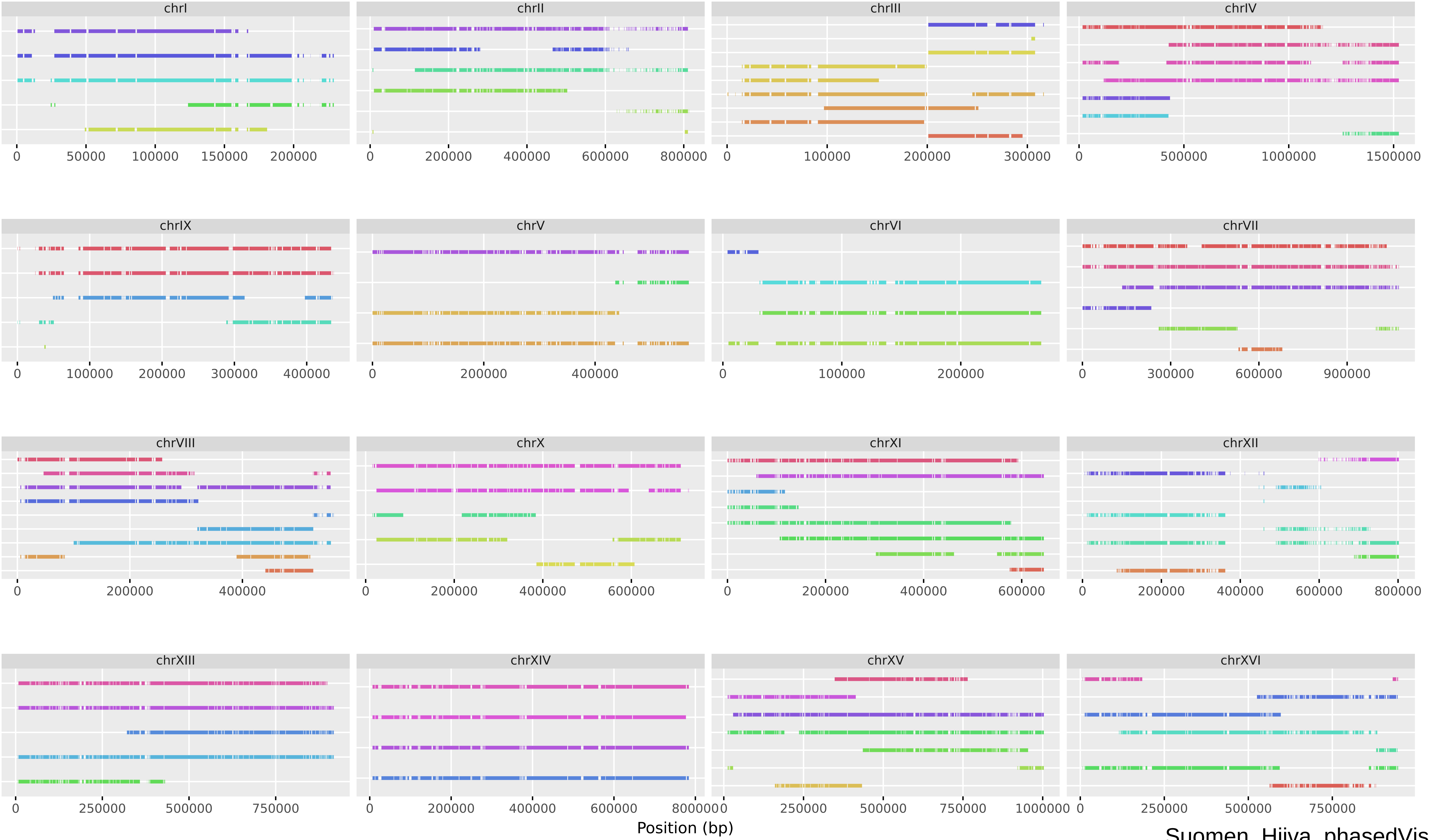


Suomen_Hiiva_phasedVis

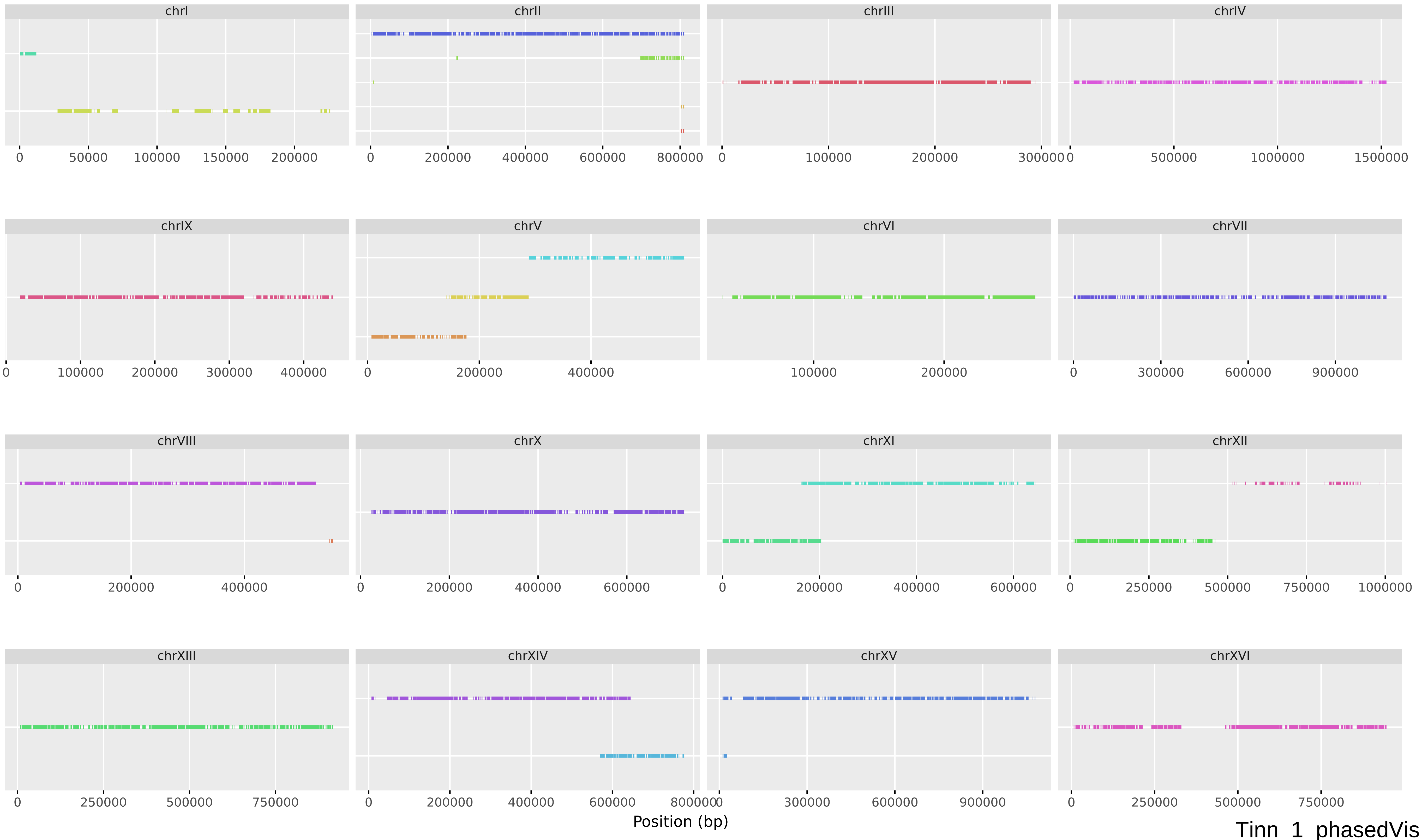
chrI
chrII
chrIII
chrIV
chrIX
chrV
chrVI
chrVII
chrVIII
chrX
chrXI
chrXII
chrXIII
chrXIV
chrXV
chrXVI
Position (bp)
Tinn_1_phasedVis

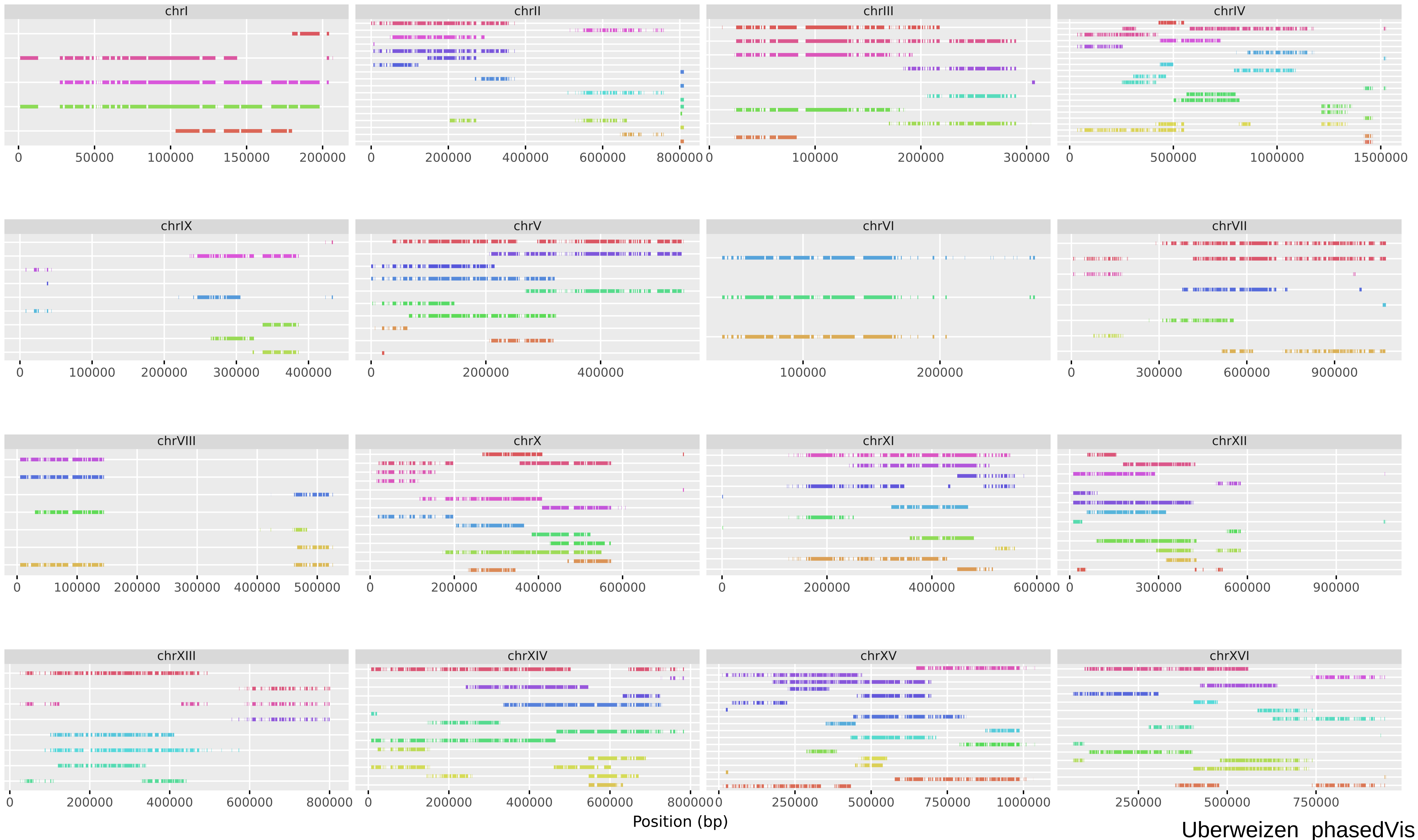
chrI
chrII
chrIII
chrIV
chrIX
chrV
chrVI
chrVII
chrVIII
chrX
chrXI
chrXII
chrXIII
chrXIV
chrXV
chrXVI
Position (bp)
Uberweizen_phasedVis

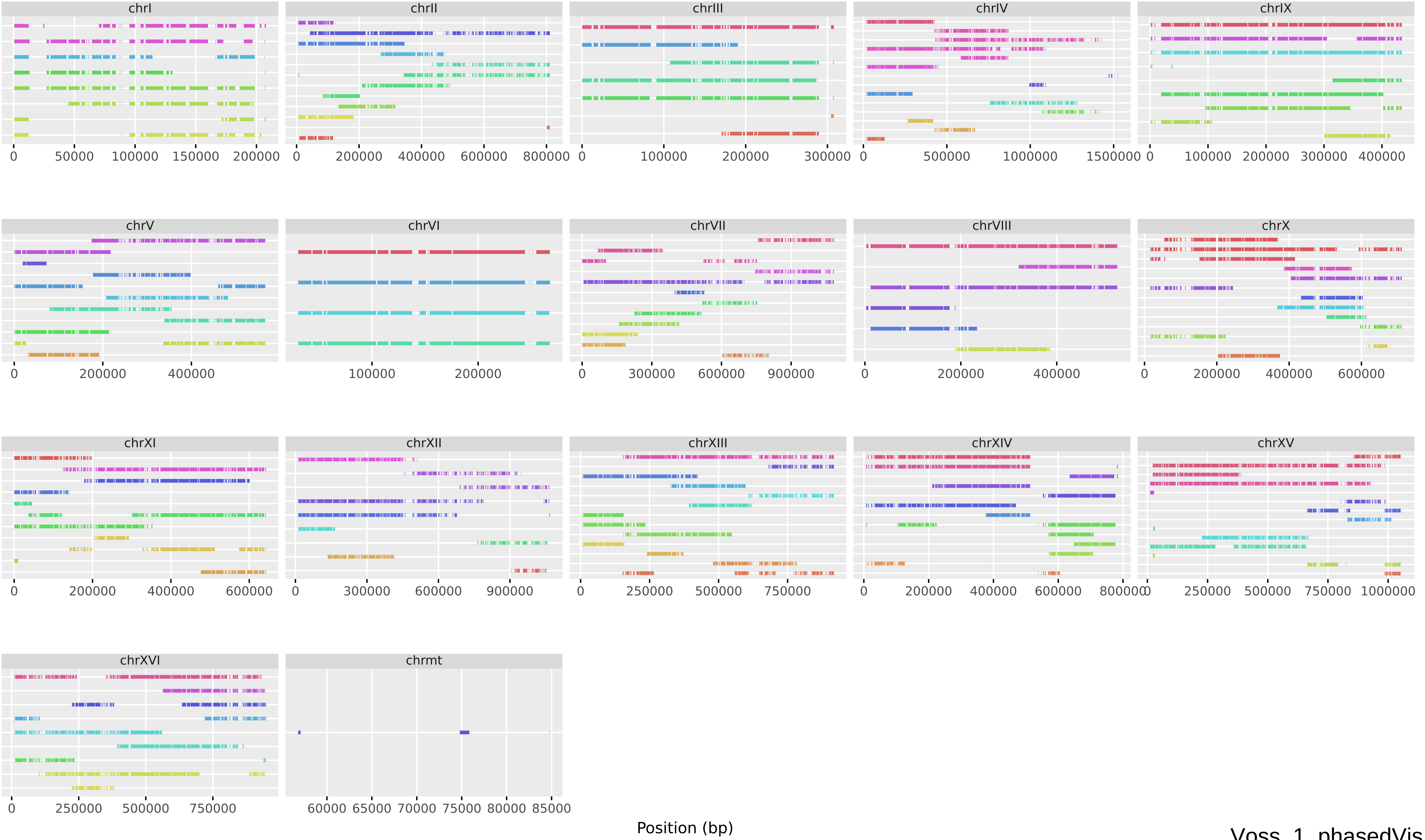


Voss_1_phasedVis

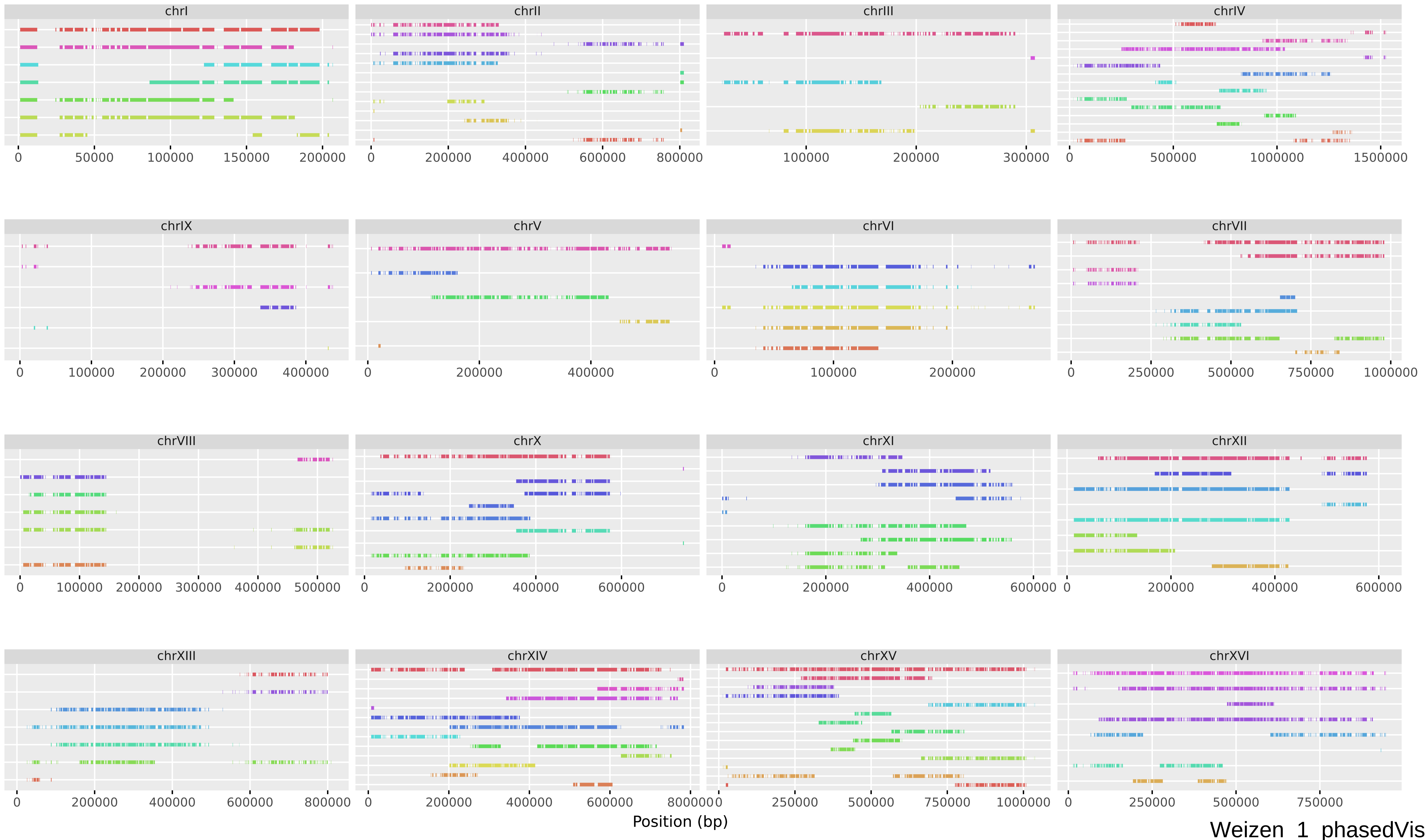
chrI
chrII
chrIII
chrIV
chrIX
chrV
chrVI
chrVII
chrVIII
chrX
chrXI
chrXII
chrXIII
chrXIV
chrXV
chrXVI
Position (bp)
Weizen_1_phasedVis
