## Supplementary Figure S3 for "European Farmhouse Brewing Yeasts Form a Distinct Genetic Group"

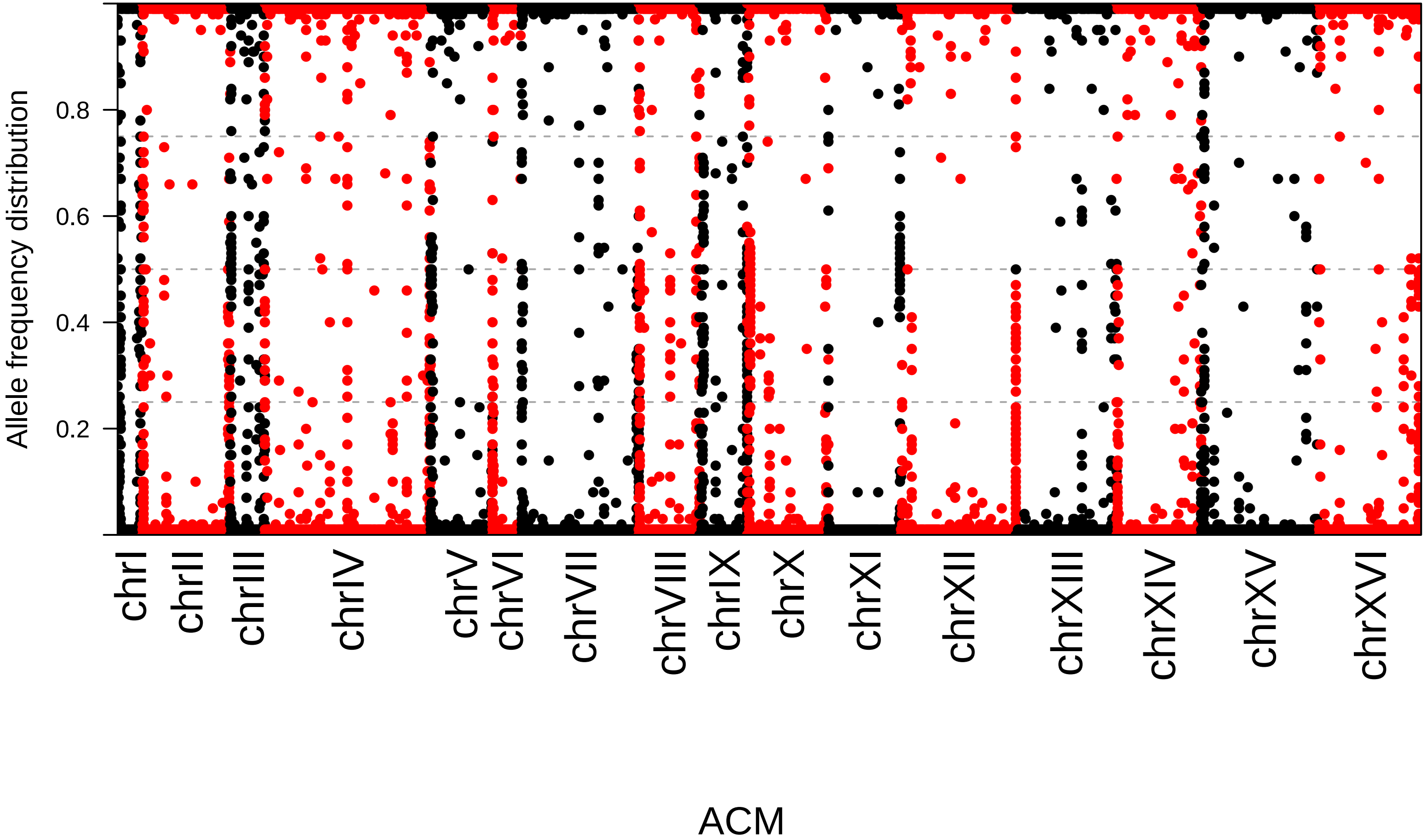
Allele frequency distribution
0.2
0.4
0.6
0.8
chrI
chrII
chrIII
chrIV
chrV
chrVI
chrVII
chrVIII
chrIX
chrX
chrXI
chrXII
chrXIII
chrXIV
chrXV
chrXVI
ACM

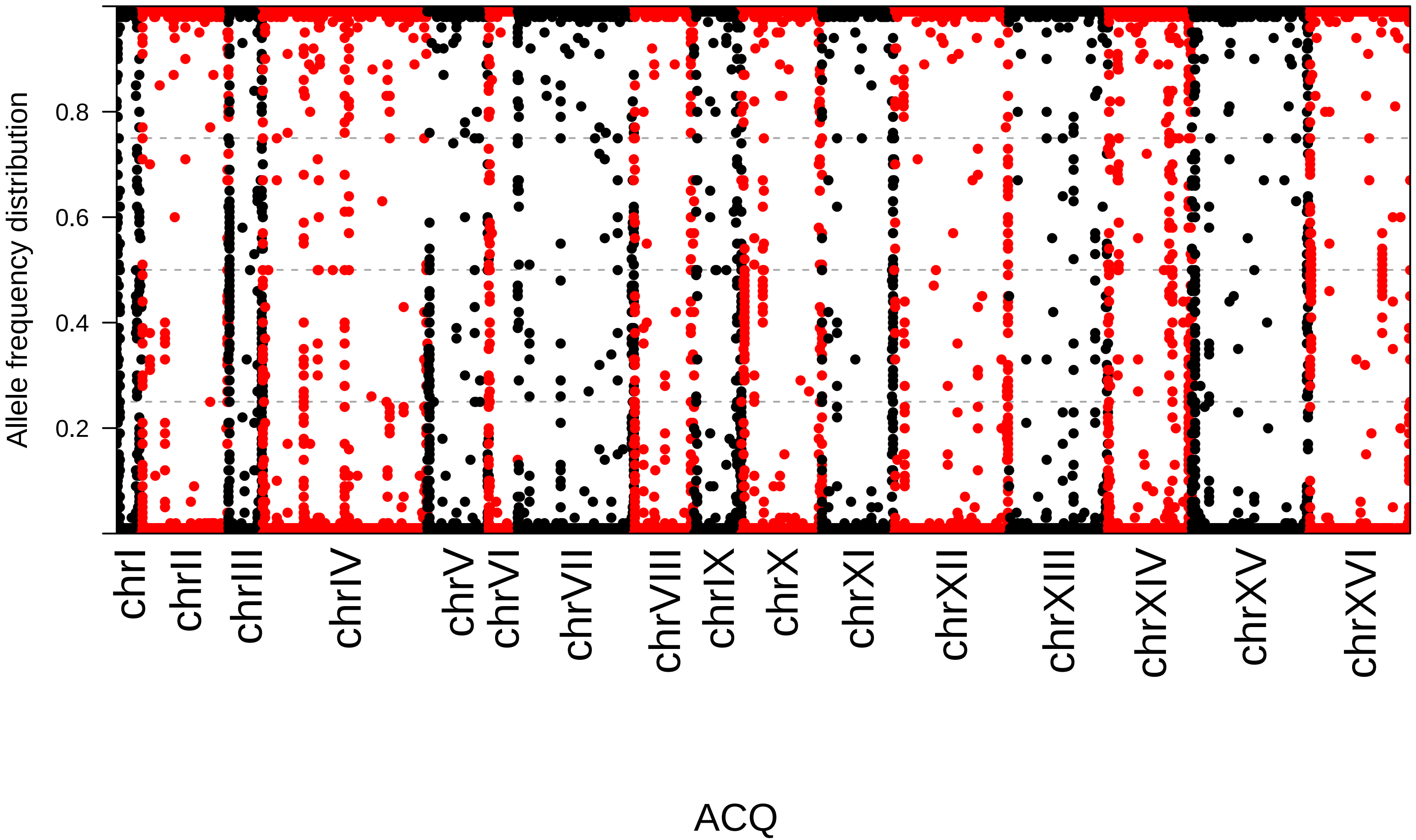
Allele frequency distribution
0.2
0.4
0.6
0.8
chrI
chrII
chrIII
chrIV
chrV
chrVI
chrVII
chrVIII
chrIX
chrX
chrXI
chrXII
chrXIII
chrXIV
chrXV
chrXVI
ACQ

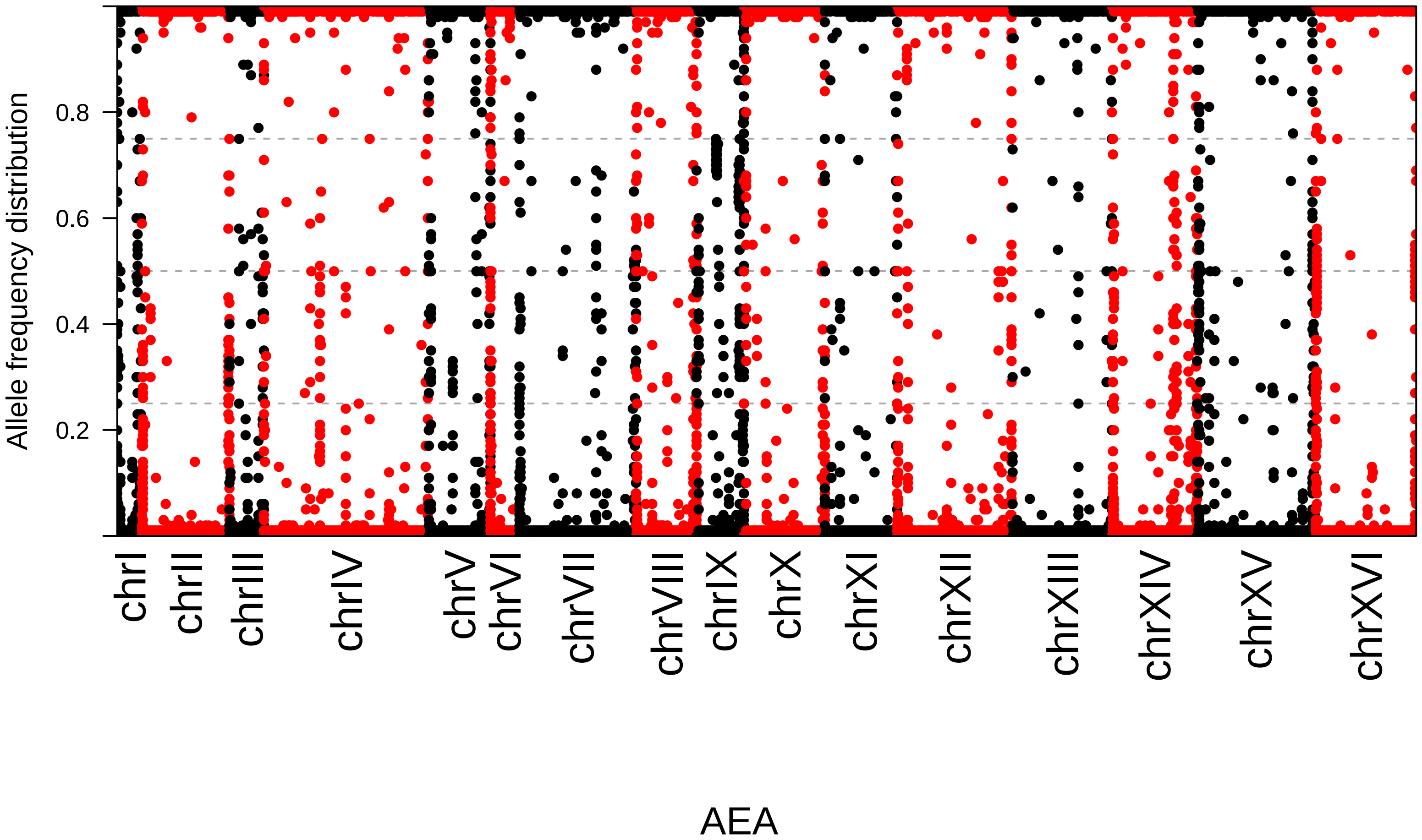

Allele frequency distribution
0.2
0.4
0.6
0.8
chrI
chrII
chrIII
chrIV
chrV
chrVI
chrVII
chrVIII
chrIX
chrX
chrXI
chrXII
chrXIII
chrXIV
chrXV
chrXVI
AEA

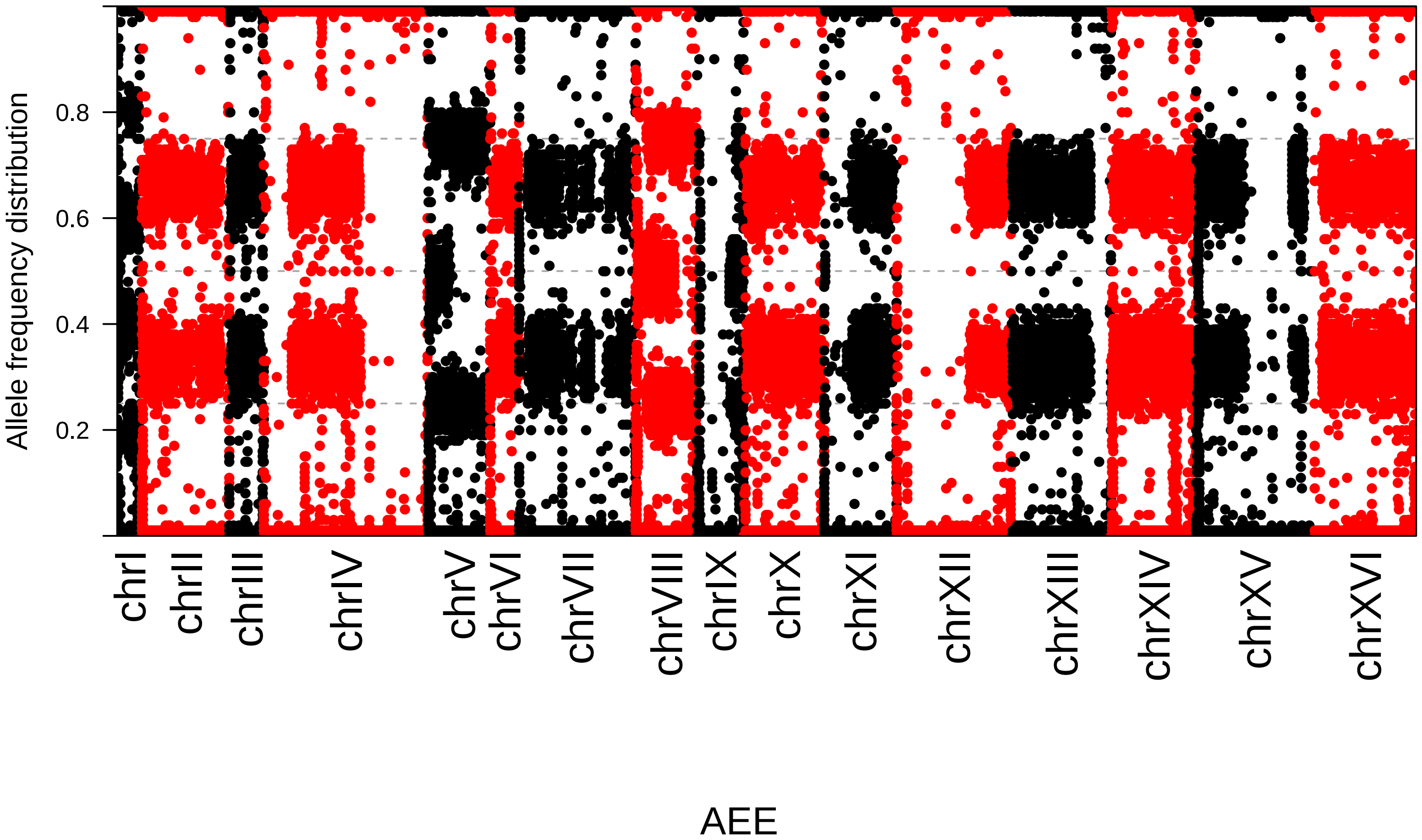

Allele frequency distribution
0.2
0.4
0.6
0.8
chrI
chrII
chrIII
chrIV
chrV
chrVI
chrVII
chrVIII
chrIX
chrX
chrXI
chrXII
chrXIII
chrXIV
chrXV
chrXVI
AEE

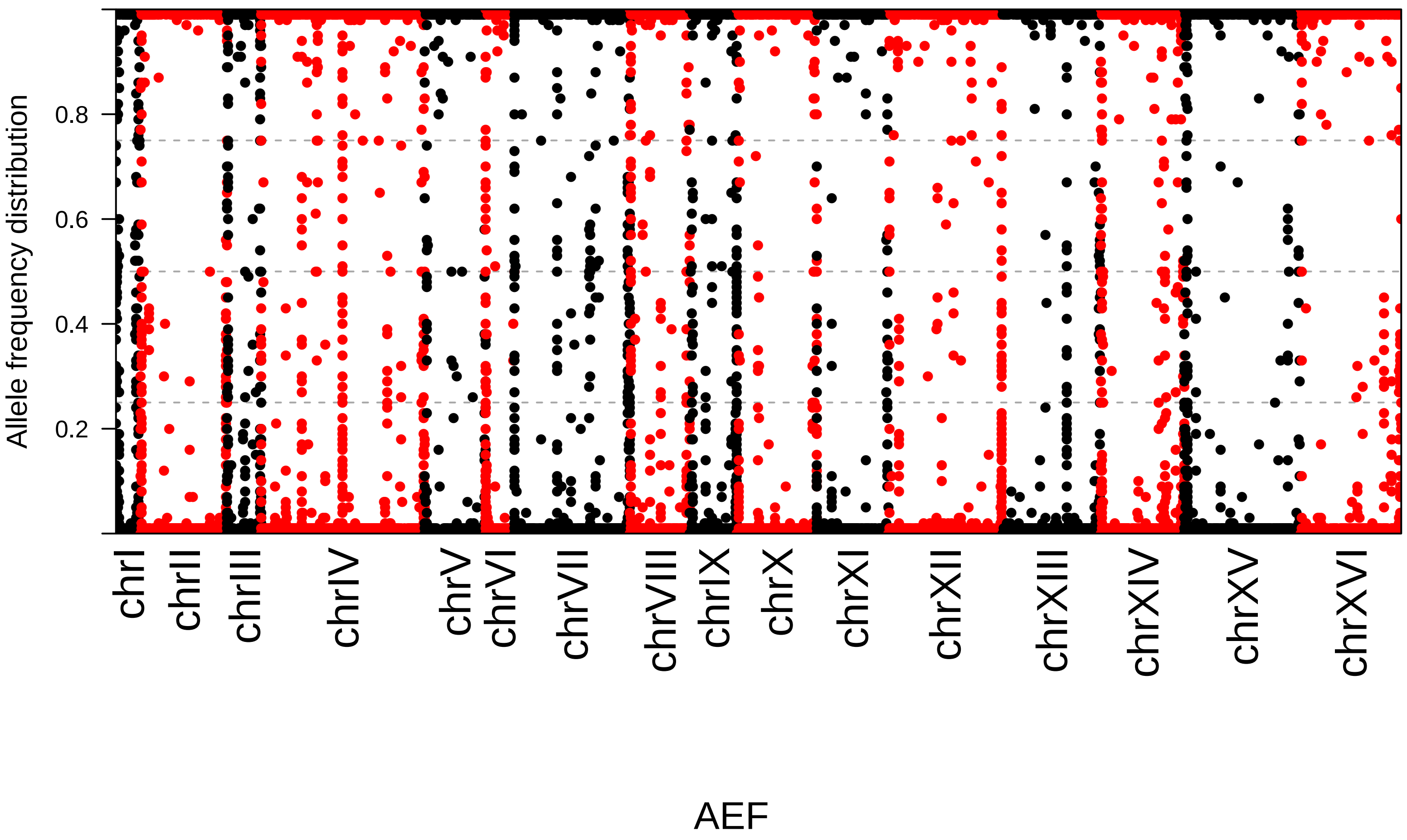
Allele frequency distribution
0.2
0.4
0.6
0.8
chrI
chrII
chrIII
chrIV
chrV
chrVI
chrVII
chrVIII
chrIX
chrX
chrXI
chrXII
chrXIII
chrXIV
chrXV
chrXVI
AEF

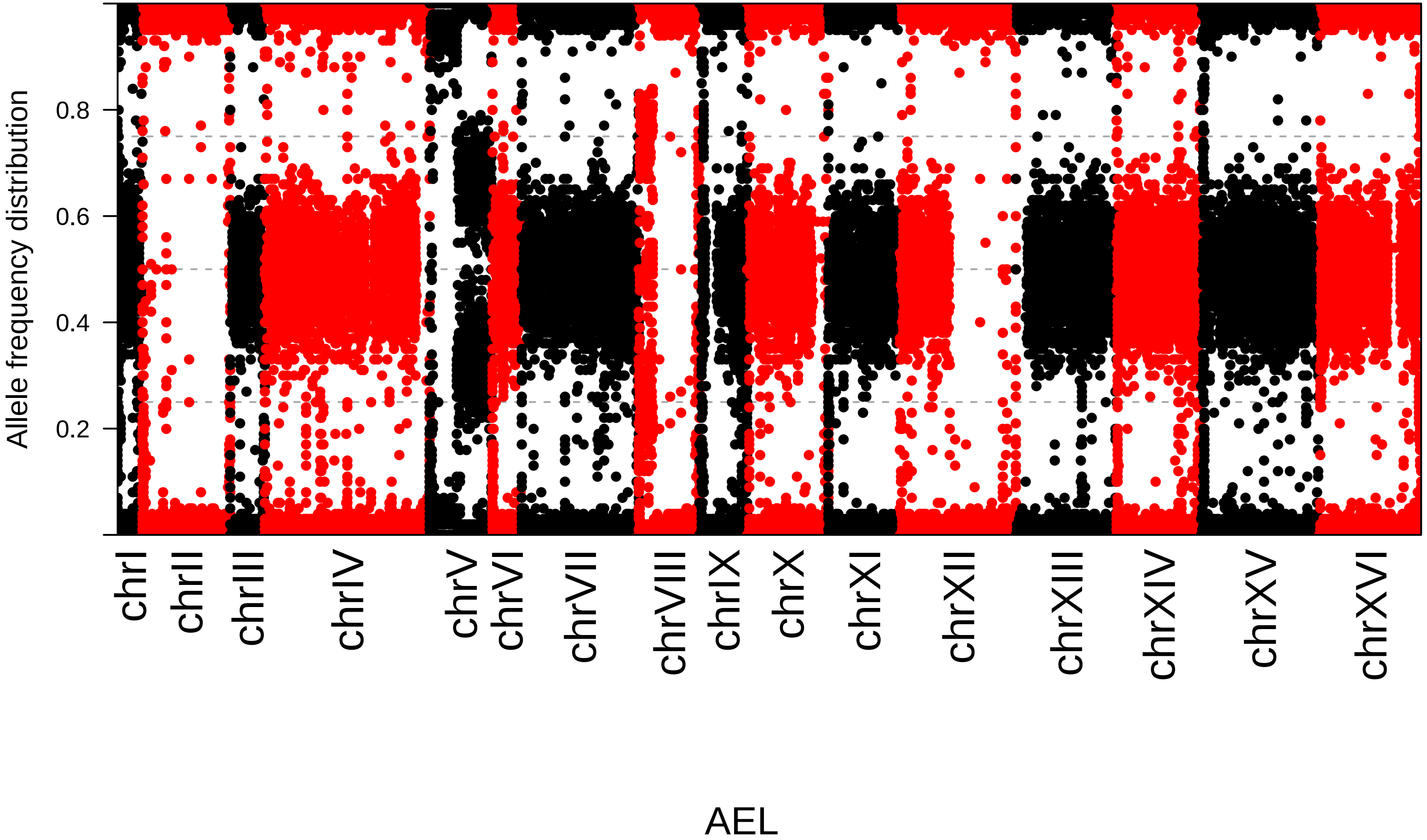

Allele frequency distribution
0.2
0.4
0.6
0.8
chrI
chrII
chrIII
chrIV
chrV
chrVI
chrVII
chrVIII
chrIX
chrX
chrXI
chrXII
chrXIII
chrXIV
chrXV
chrXVI
AEL

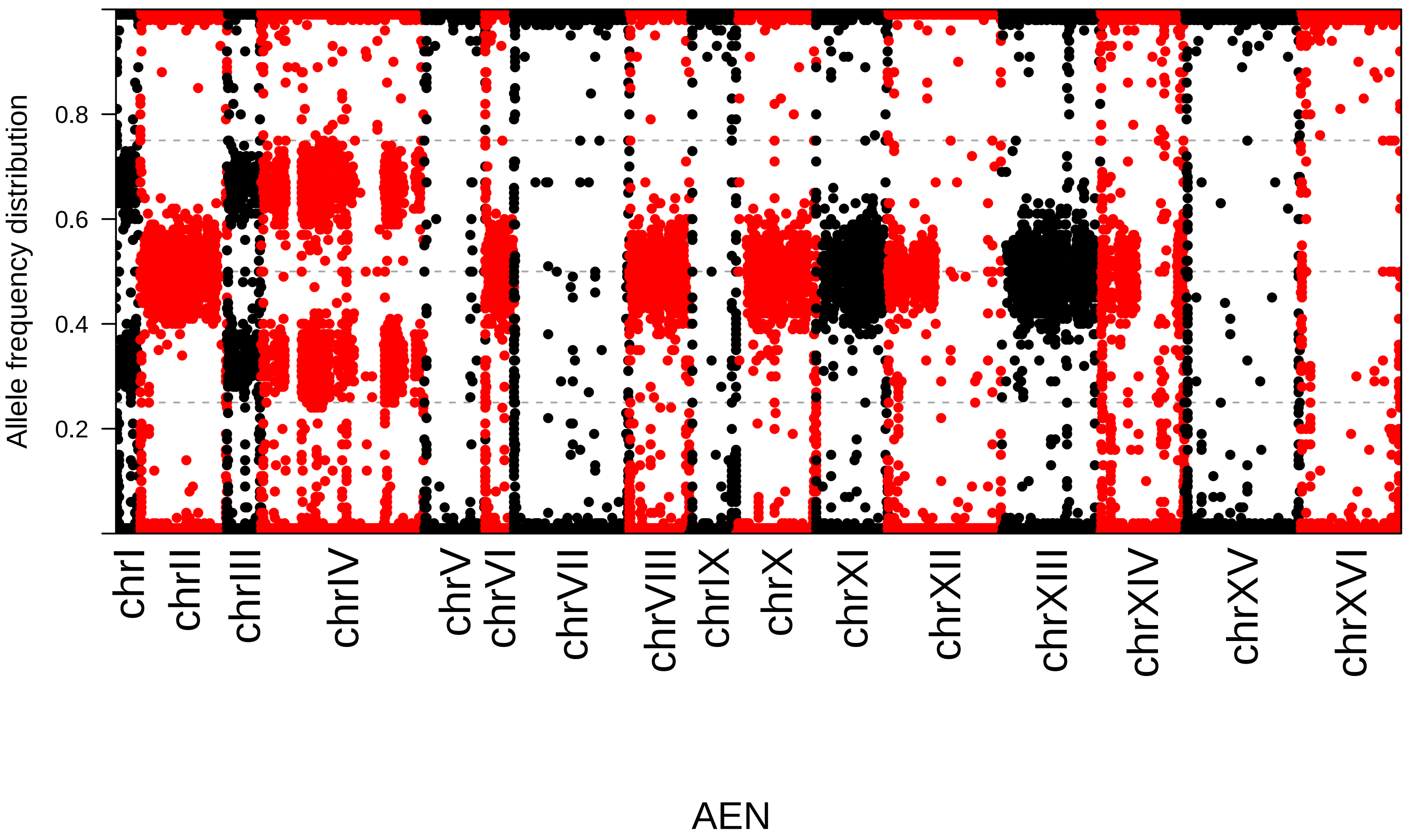

Allele frequency distribution
0.2
0.4
0.6
0.8
chrI
chrII
chrIII
chrIV
chrV
chrVI
chrVII
chrVIII
chrIX
chrX
chrXI
chrXII
chrXIII
chrXIV
chrXV
chrXVI
AEN

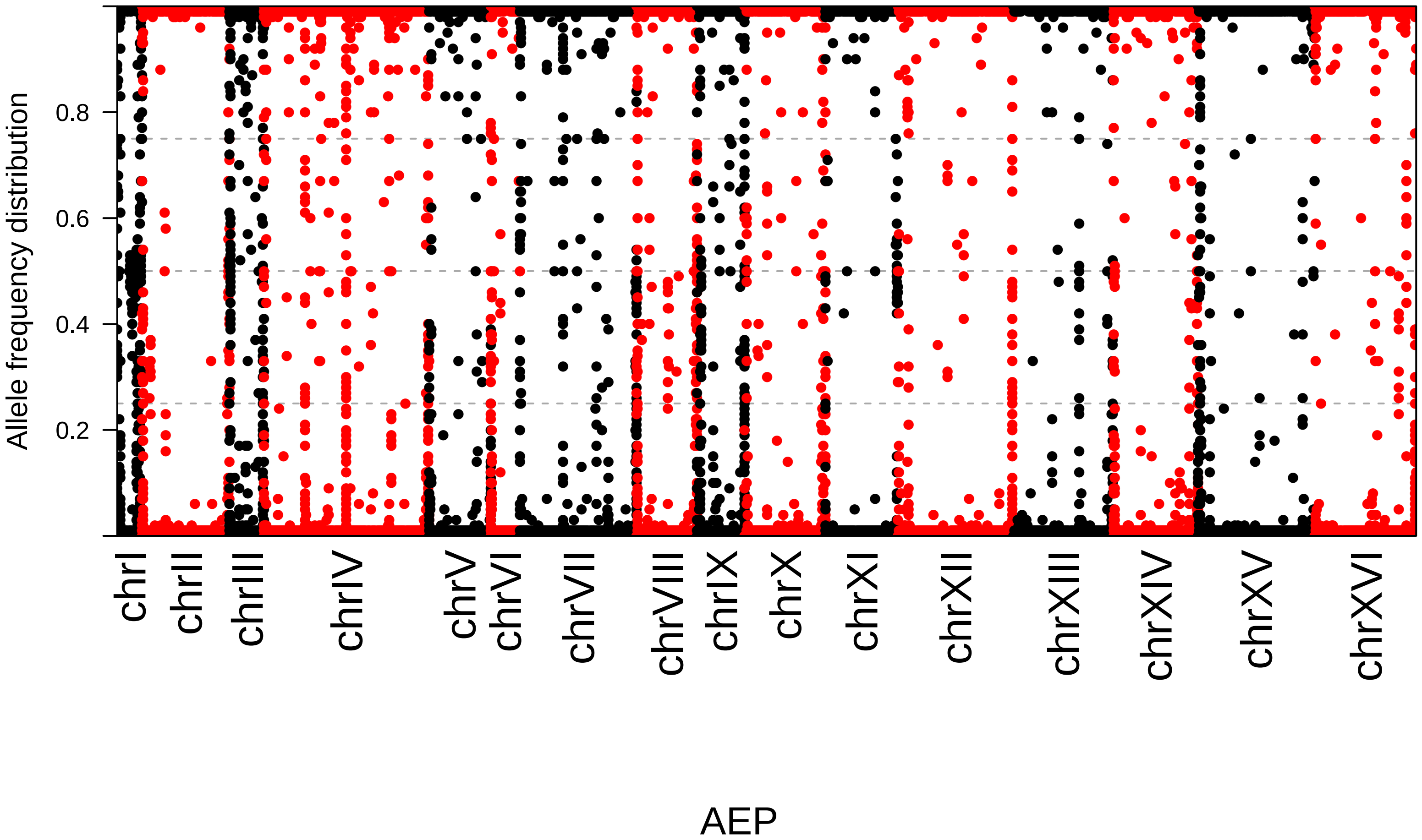

Allele frequency distribution
0.2
0.4
0.6
0.8
chrI
chrII
chrIII
chrIV
chrV
chrVI
chrVII
chrVIII
chrIX
chrX
chrXI
chrXII
chrXIII
chrXIV
chrXV
chrXVI
AEP

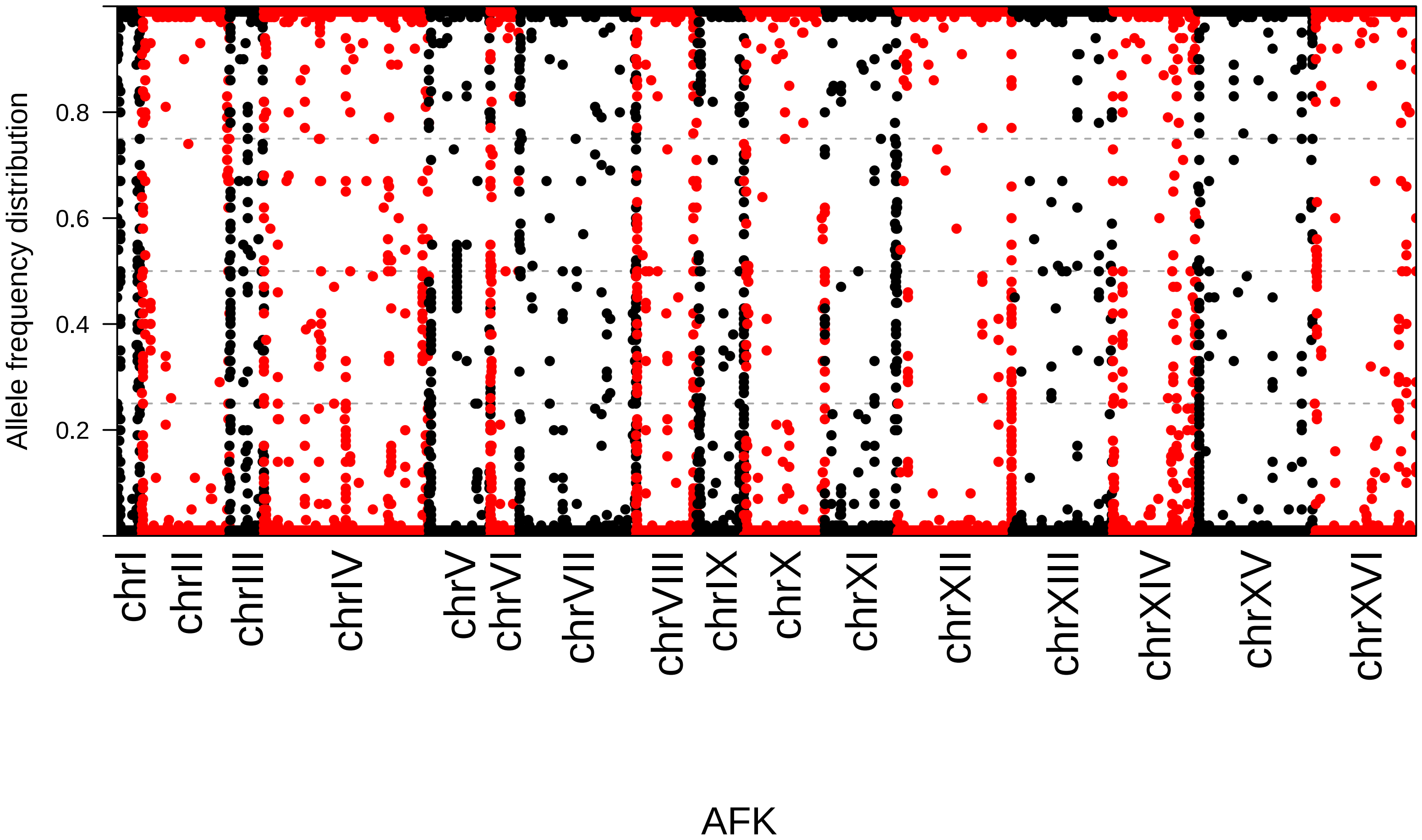
Allele frequency distribution
0.2
0.4
0.6
0.8
chrI
chrII
chrIII
chrIV
chrV
chrVI
chrVII
chrVIII
chrIX
chrX
chrXI
chrXII
chrXIII
chrXIV
chrXV
chrXVI
AFK

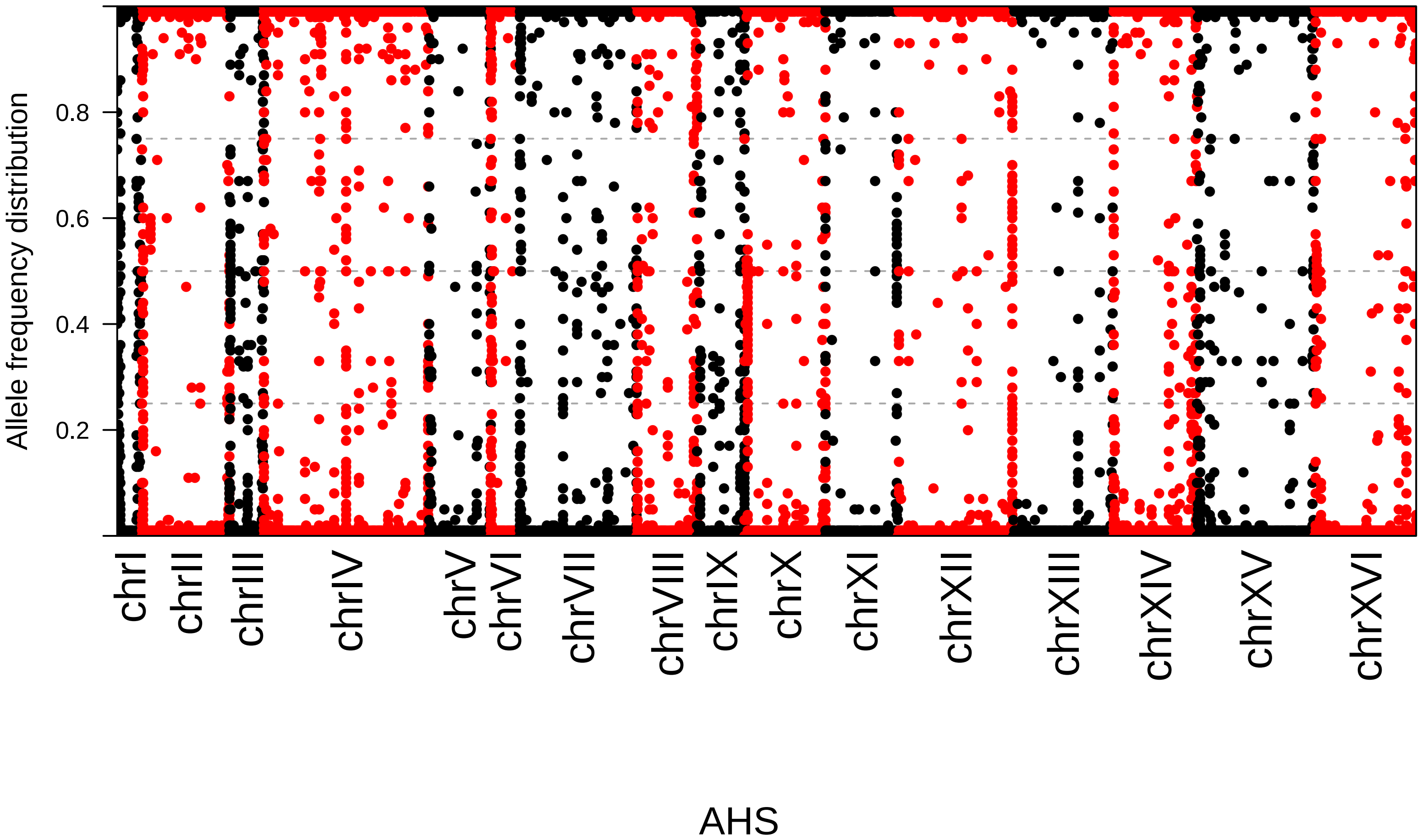

Allele frequency distribution
0.8
0.6
0.4
0.2
chrI
chrII
chrIII
chrIV
chrV
chrVI
chrVII
chrVIII
chrIX
chrX
chrXI
chrXII
chrXIII
chrXIV
chrXV
chrXVI
AHS

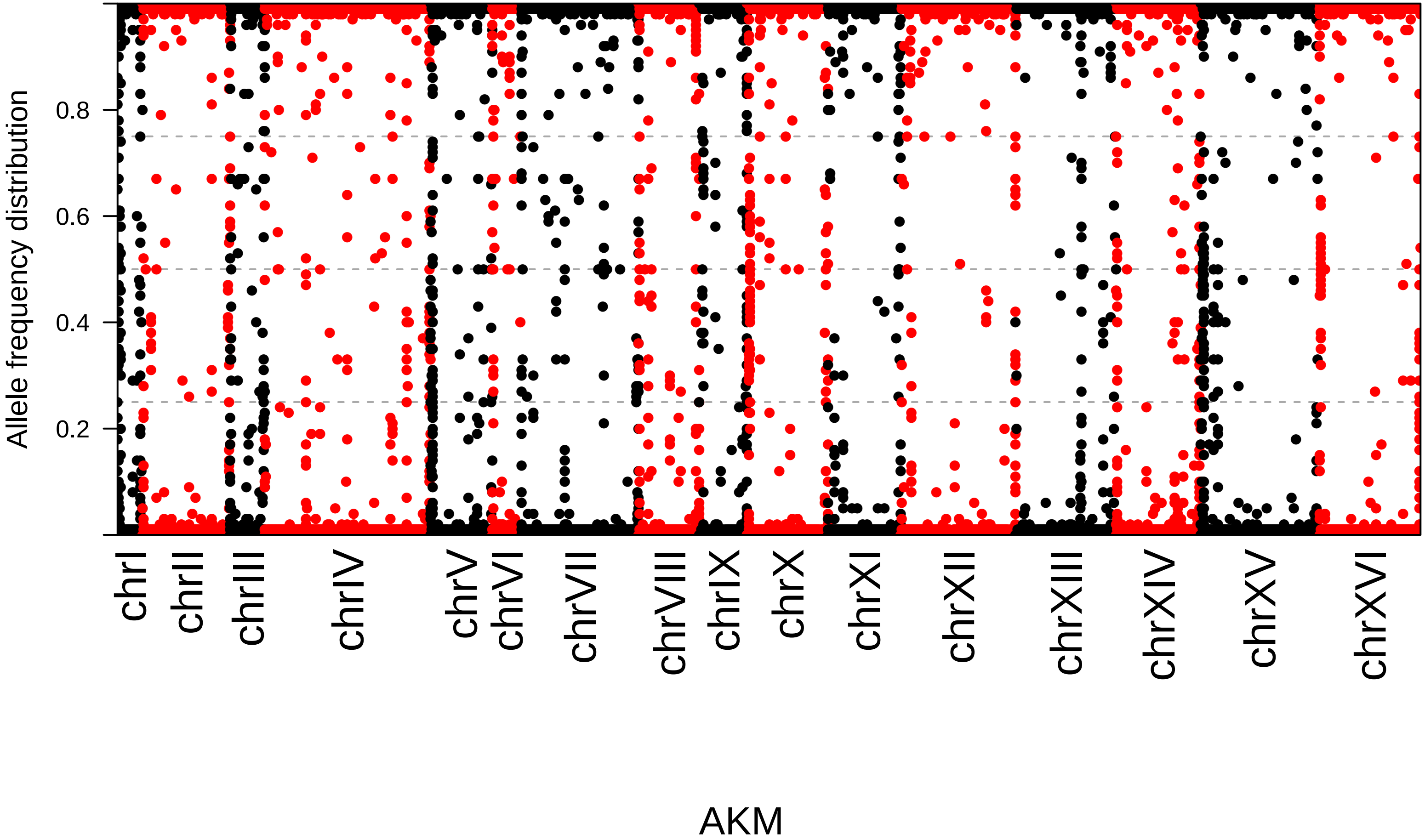

Allele frequency distribution
0.2
0.4
0.6
0.8
chrI
chrII
chrIII
chrIV
chrV
chrVI
chrVII
chrVIII
chrIX
chrX
chrXI
chrXII
chrXIII
chrXIV
chrXV
chrXVI
AKM

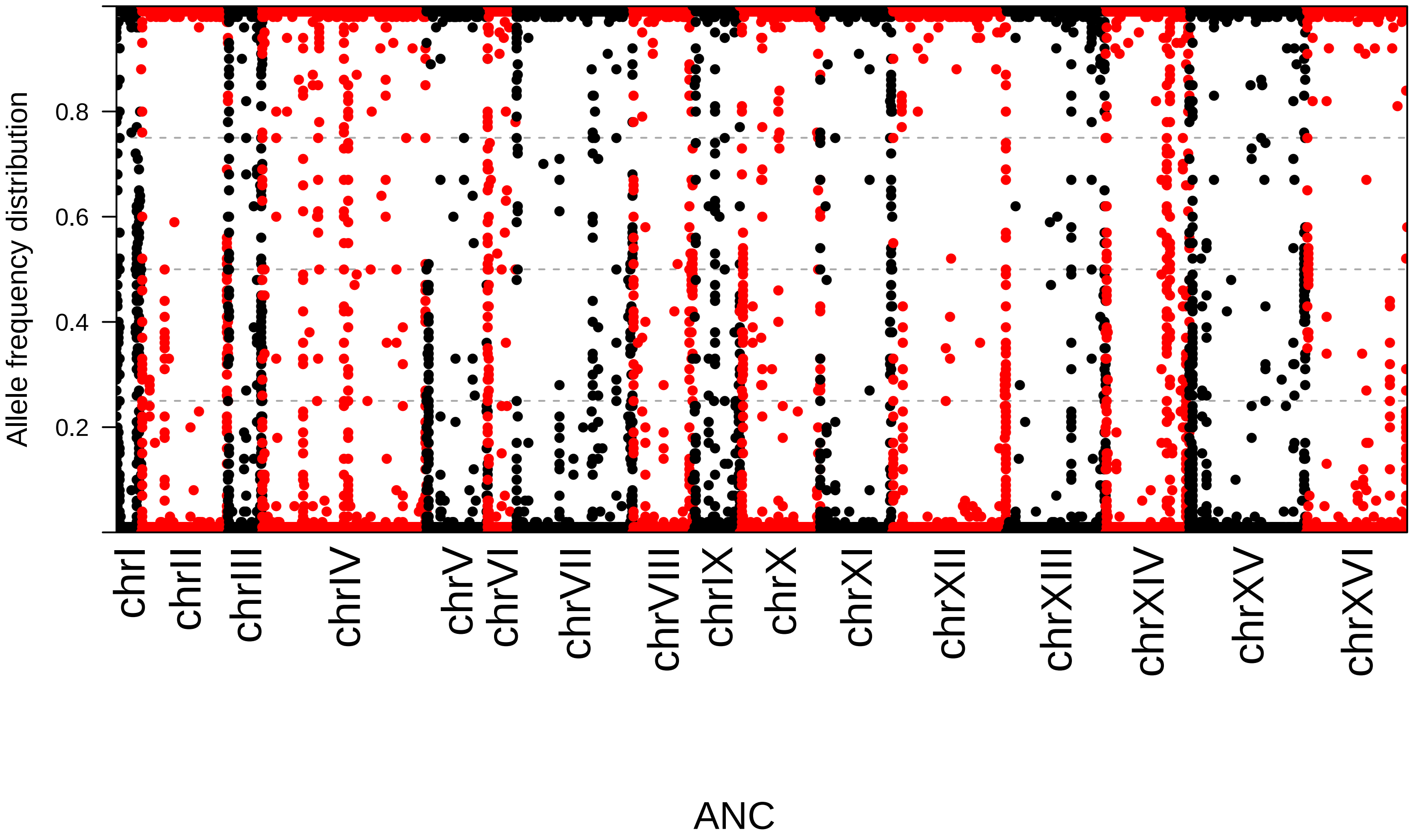

Allele frequency distribution
0.2
0.4
0.6
0.8
chrI
chrII
chrIII
chrIV
chrV
chrVI
chrVII
chrVIII
chrIX
chrX
chrXI
chrXII
chrXIII
chrXIV
chrXV
chrXVI
ANC

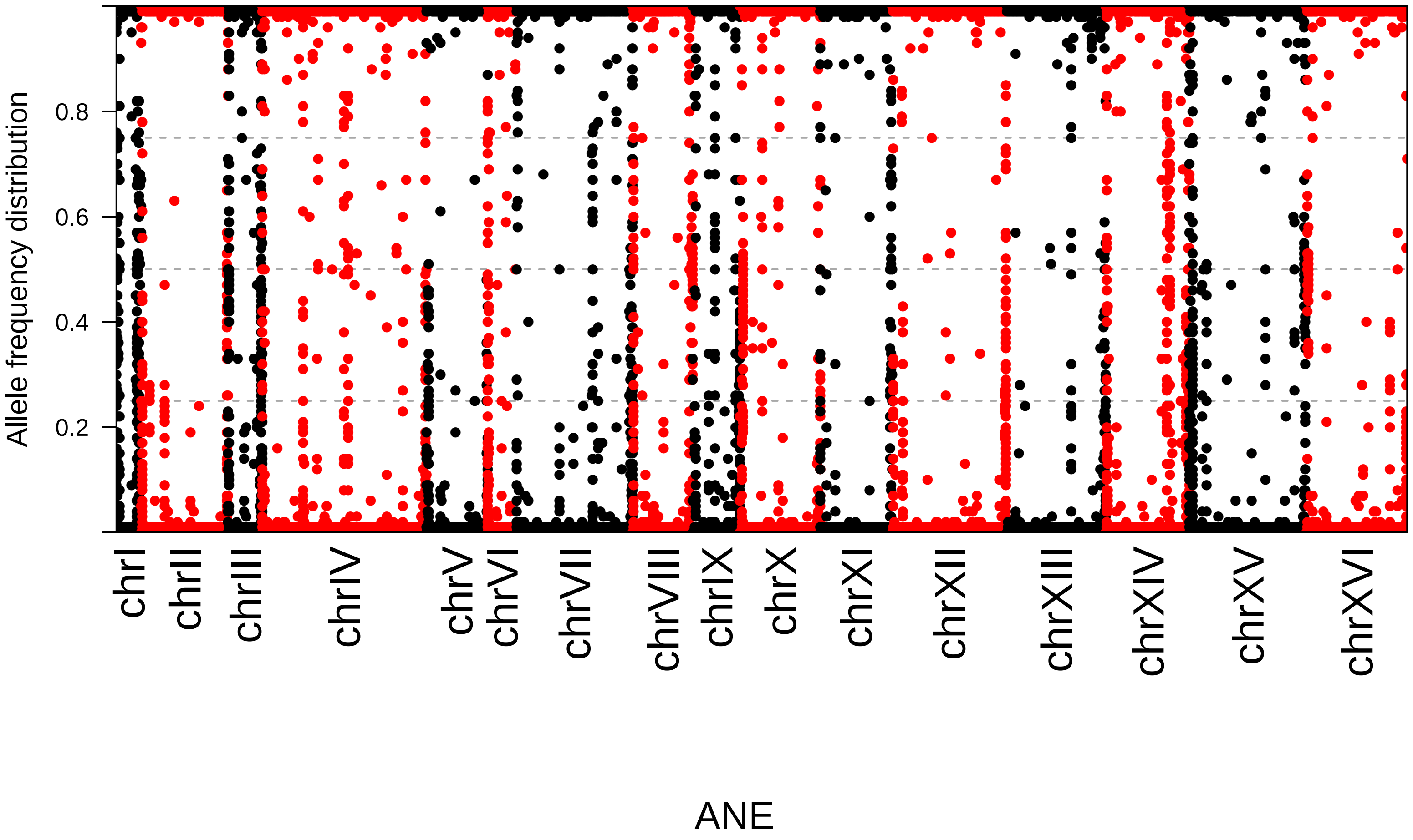

Allele frequency distribution
0.2
0.4
0.6
0.8
chrI
chrII
chrIII
chrIV
chrV
chrVI
chrVII
chrVIII
chrIX
chrX
chrXI
chrXII
chrXIII
chrXIV
chrXV
chrXVI
ANE

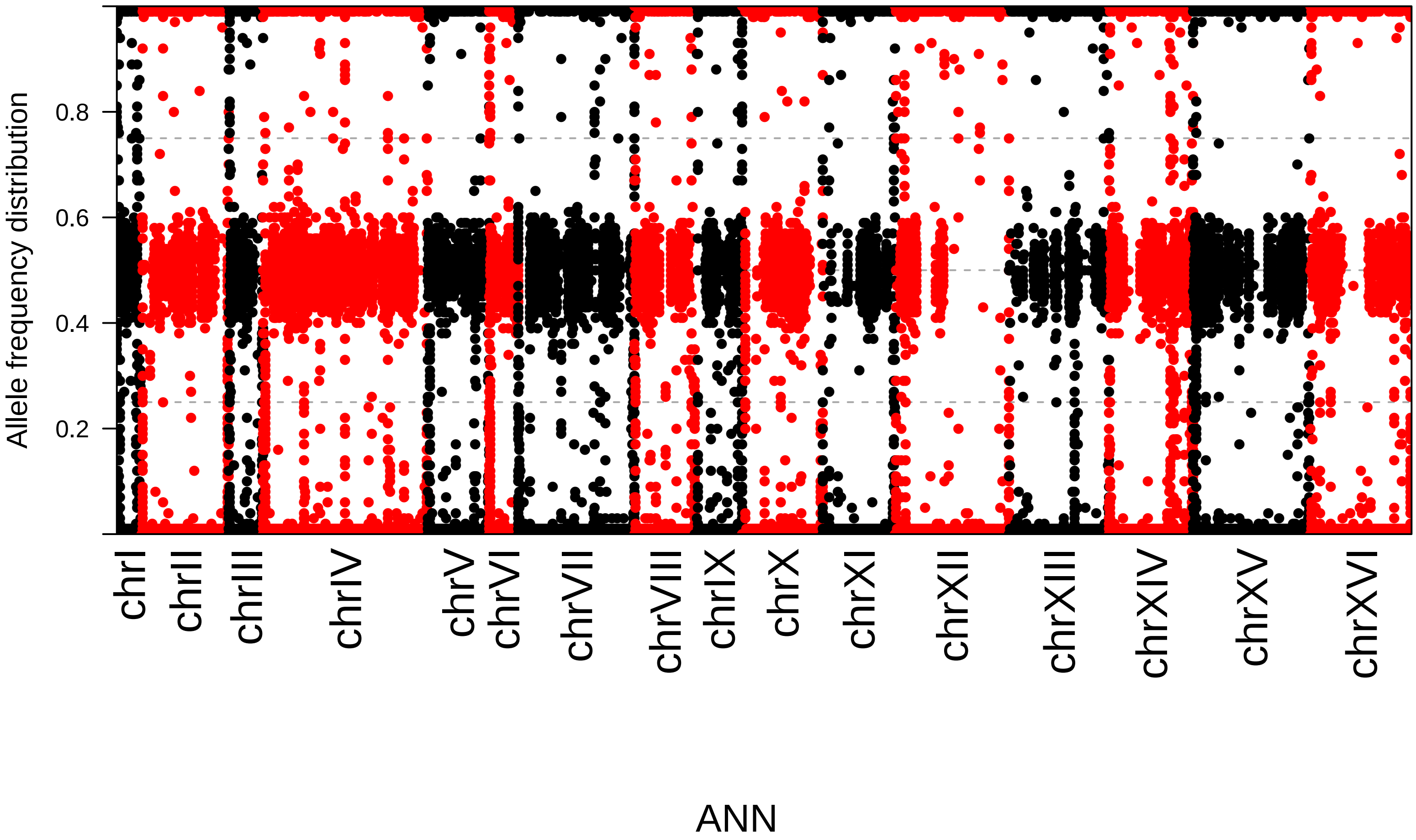

Allele frequency distribution
0.8
0.6
0.4
0.2
chrI
chrII
chrIII
chrIV
chrV
chrVI
chrVII
chrVIII
chrIX
chrX
chrXI
chrXII
chrXIII
chrXIV
chrXV
chrXVI
ANN

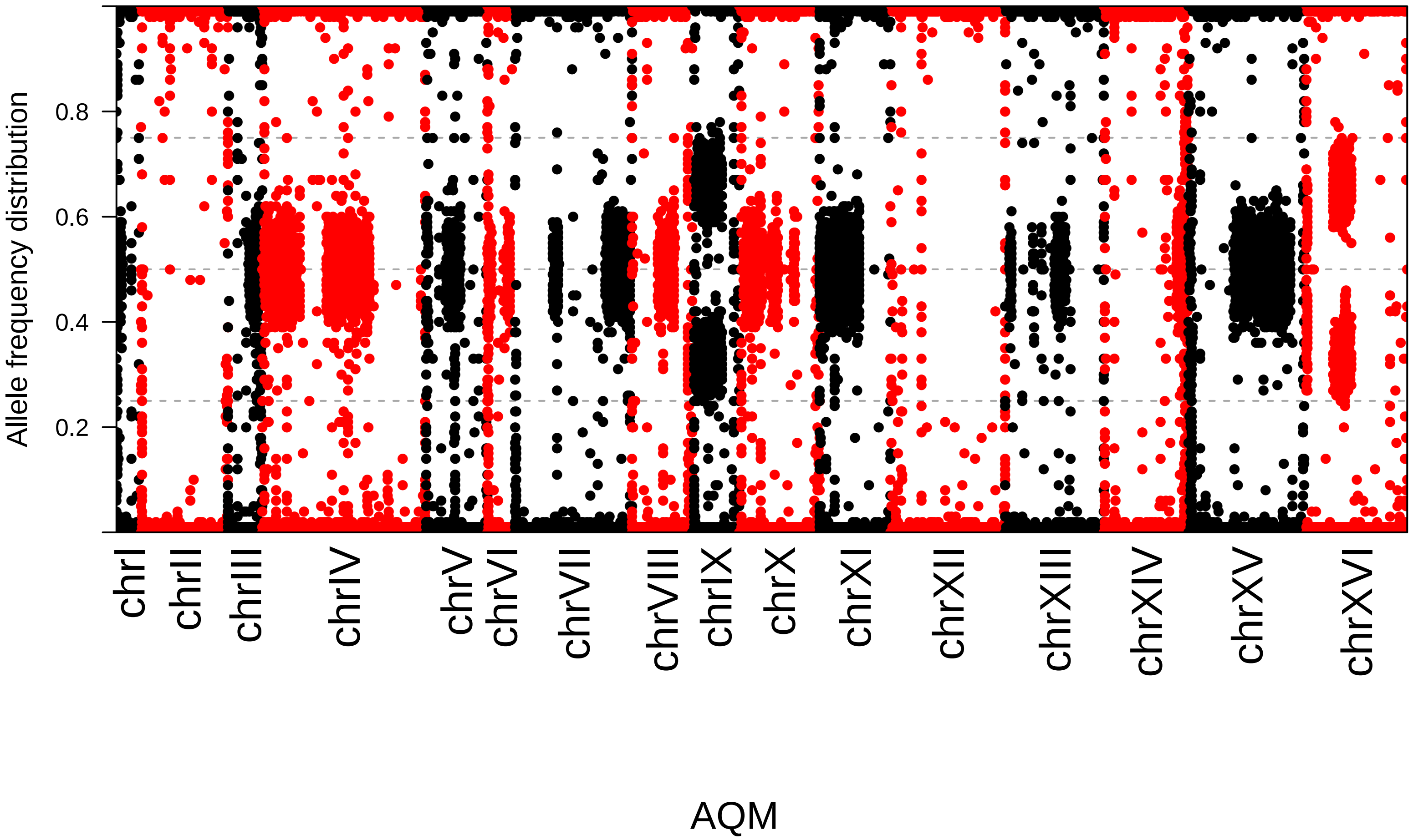
Allele frequency distribution
0.2
0.4
0.6
0.8
chrI
chrII
chrIII
chrIV
chrV
chrVI
chrVII
chrVIII
chrIX
chrX
chrXI
chrXII
chrXIII
chrXIV
chrXV
chrXVI
AQM

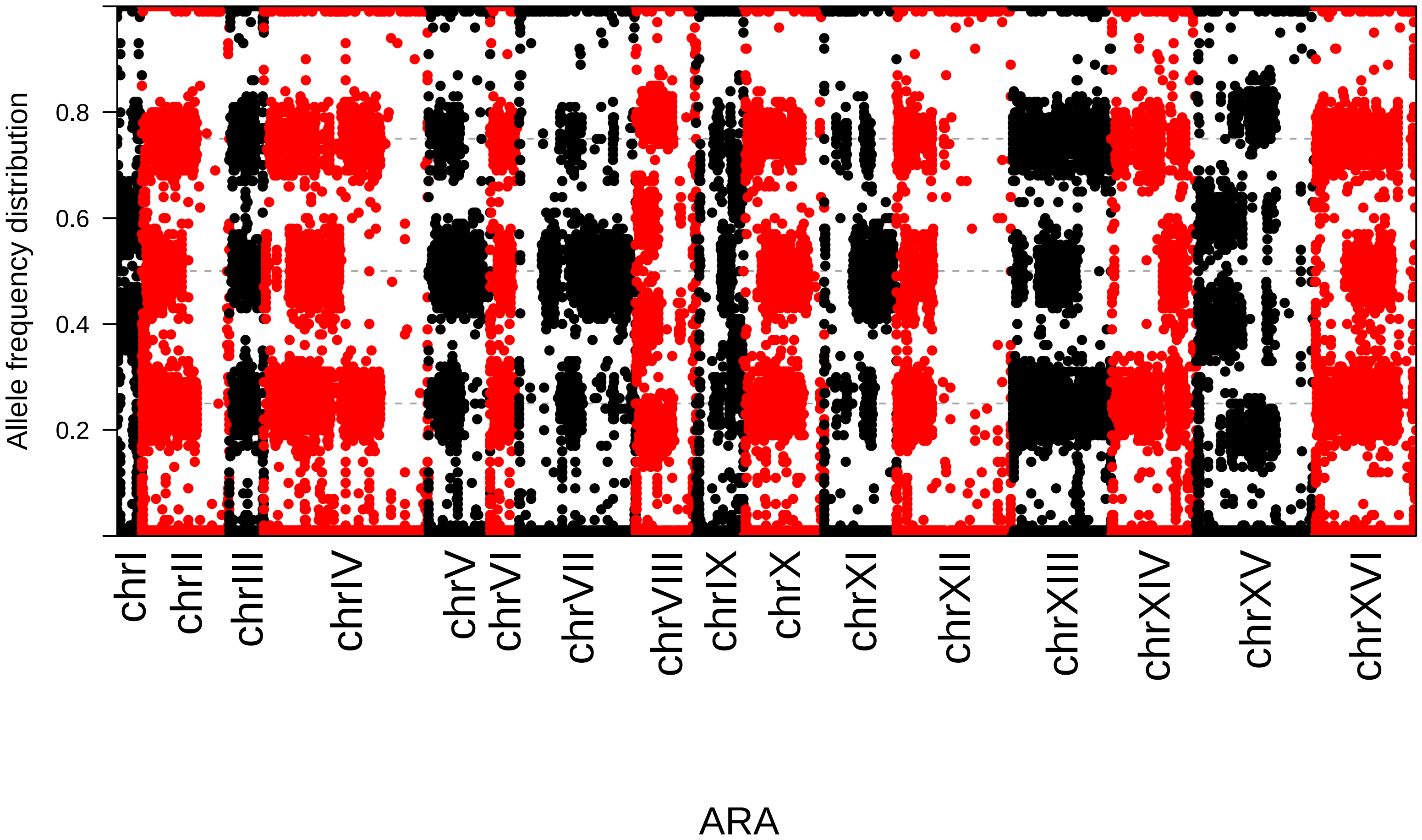

Allele frequency distribution
0.2
0.4
0.6
0.8
chrI
chrII
chrIII
chrIV
chrV
chrVI
chrVII
chrVIII
chrIX
chrX
chrXI
chrXII
chrXIII
chrXIV
chrXV
chrXVI
ARA

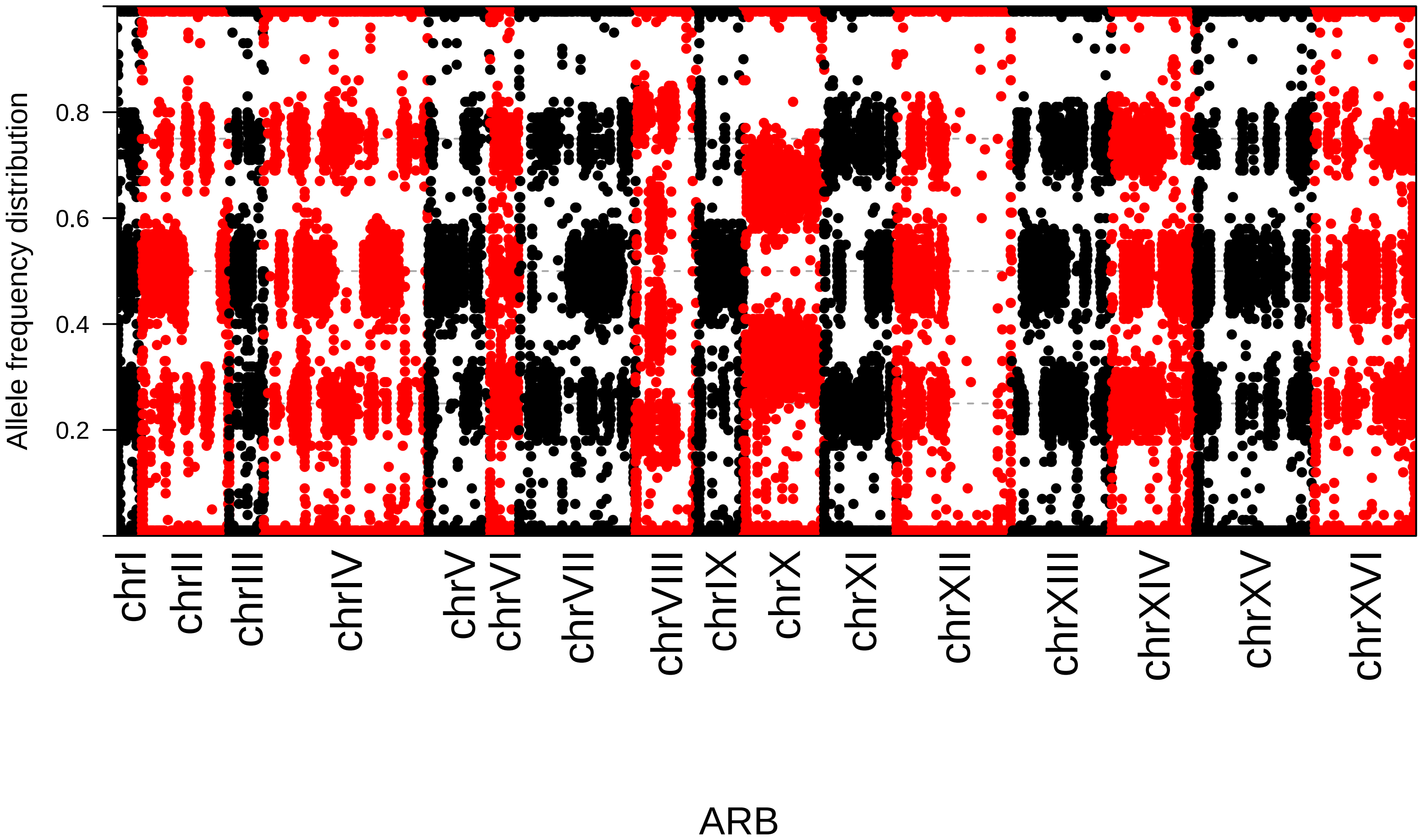

Allele frequency distribution
0.2
0.4
0.6
0.8
chrI
chrII
chrIII
chrIV
chrV
chrVI
chrVII
chrVIII
chrIX
chrX
chrXI
chrXII
chrXIII
chrXIV
chrXV
chrXVI
ARB

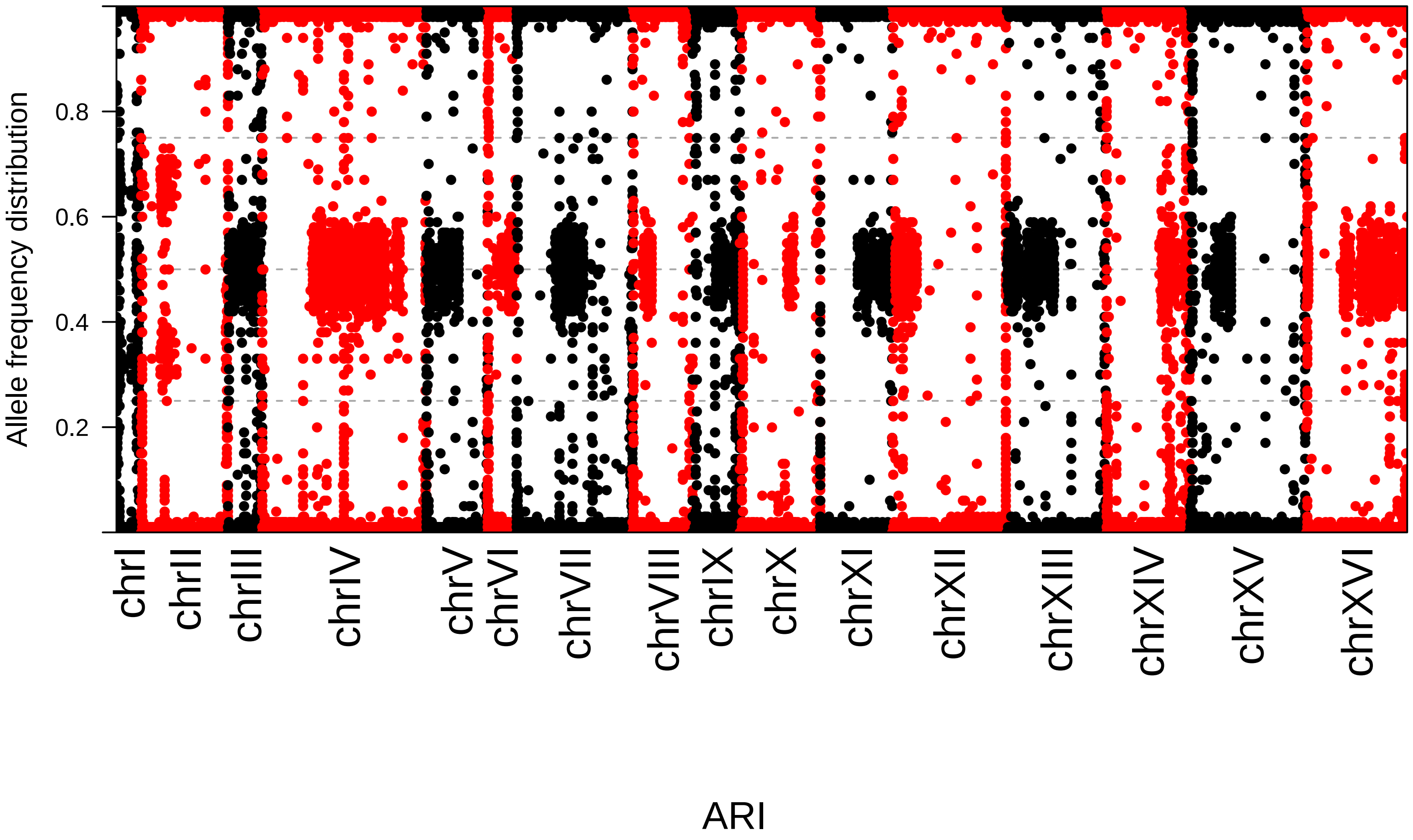
Allele frequency distribution
0.2
0.4
0.6
0.8
chrI
chrII
chrIII
chrIV
chrV
chrVI
chrVII
chrVIII
chrIX
chrX
chrXI
chrXII
chrXIII
chrXIV
chrXV
chrXVI
ARI

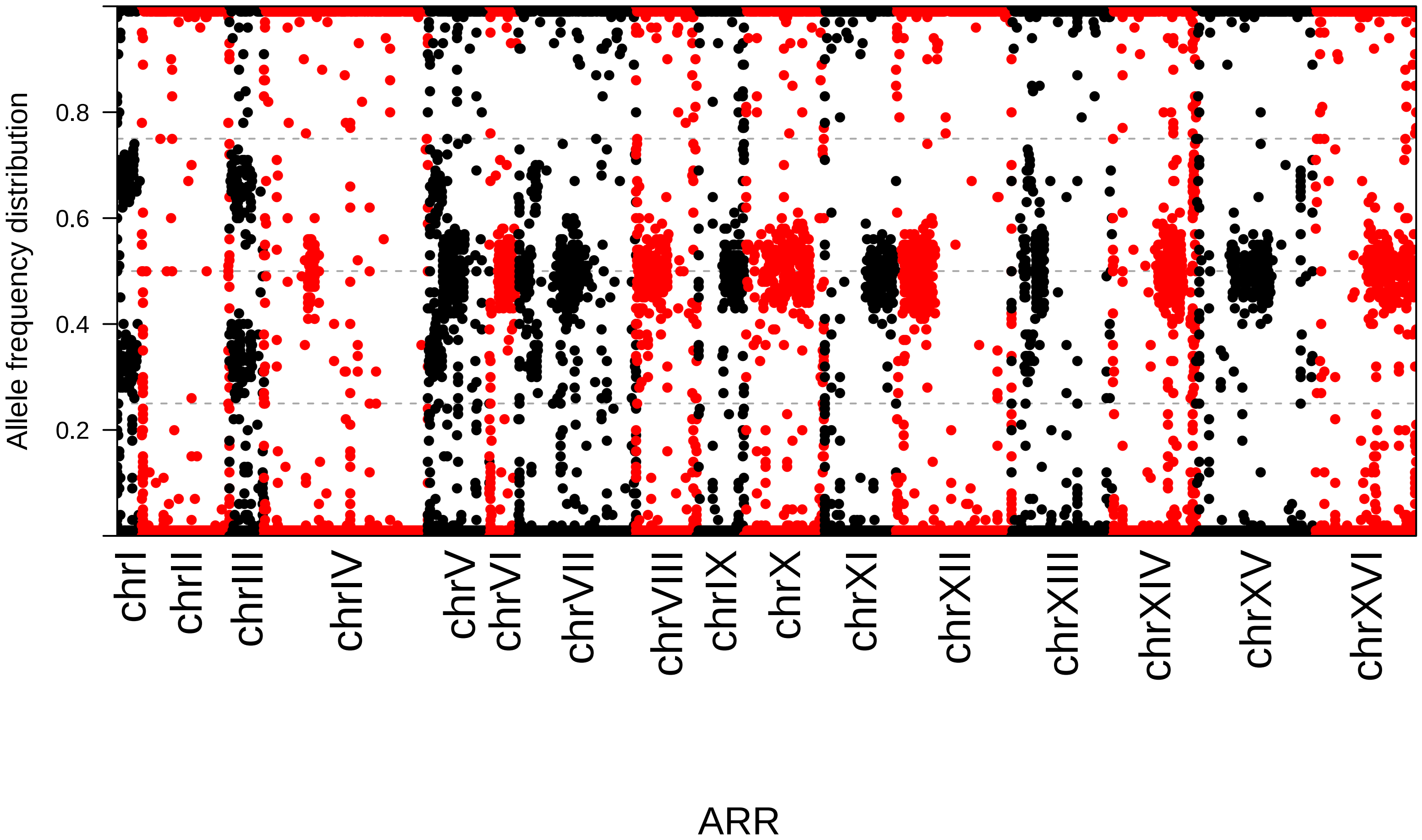

Allele frequency distribution
0.2
0.4
0.6
0.8
chrI
chrII
chrIII
chrIV
chrV
chrVI
chrVII
chrVIII
chrIX
chrX
chrXI
chrXII
chrXIII
chrXIV
chrXV
chrXVI
ARR

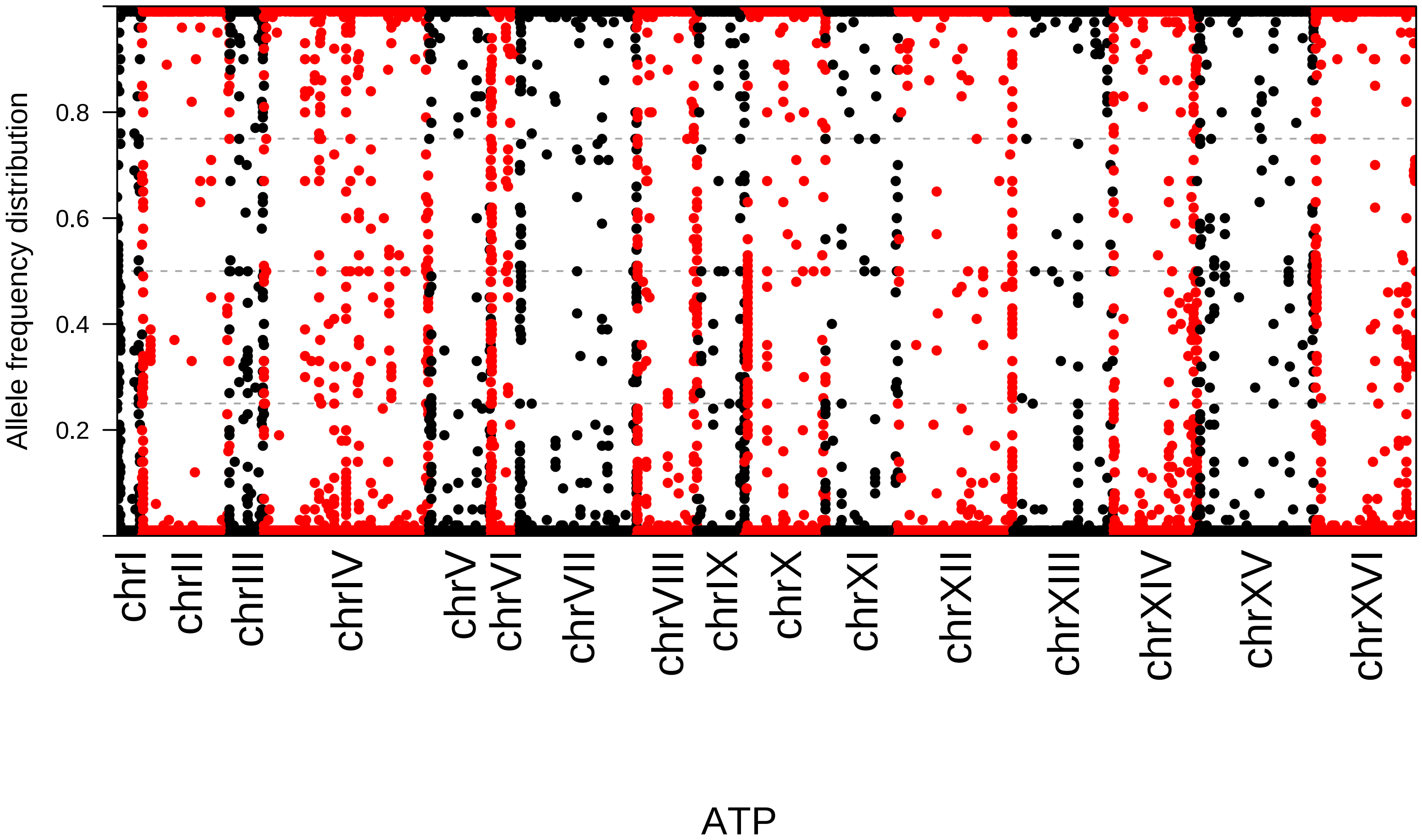
Allele frequency distribution
0.2
0.4
0.6
0.8
chrI
chrII
chrIII
chrIV
chrV
chrVI
chrVII
chrVIII
chrIX
chrX
chrXI
chrXII
chrXIII
chrXIV
chrXV
chrXVI
ATP

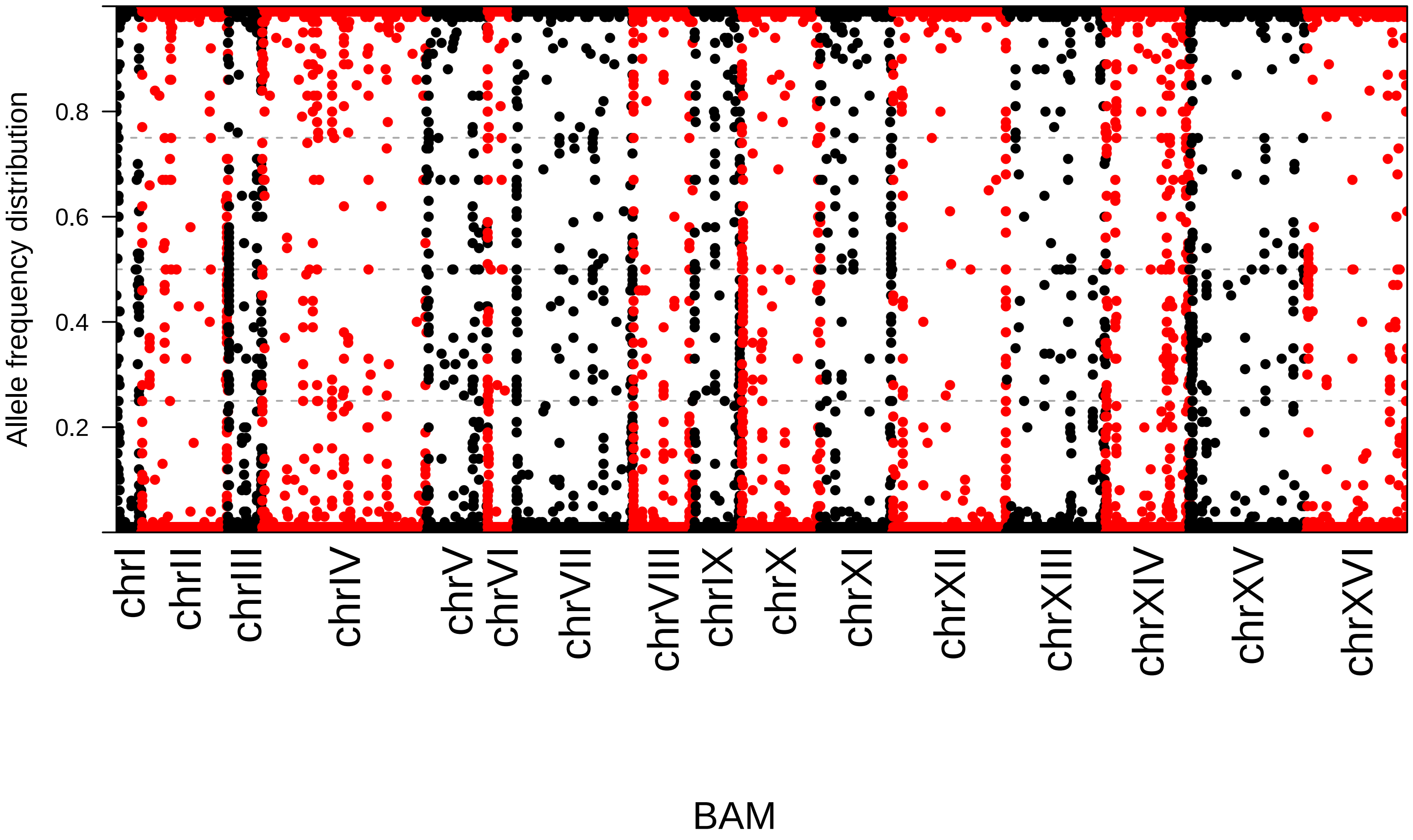

Allele frequency distribution
0.2
0.4
0.6
0.8
chrI
chrII
chrIII
chrIV
chrV
chrVI
chrVII
chrVIII
chrIX
chrX
chrXI
chrXII
chrXIII
chrXIV
chrXV
chrXVI
BAM

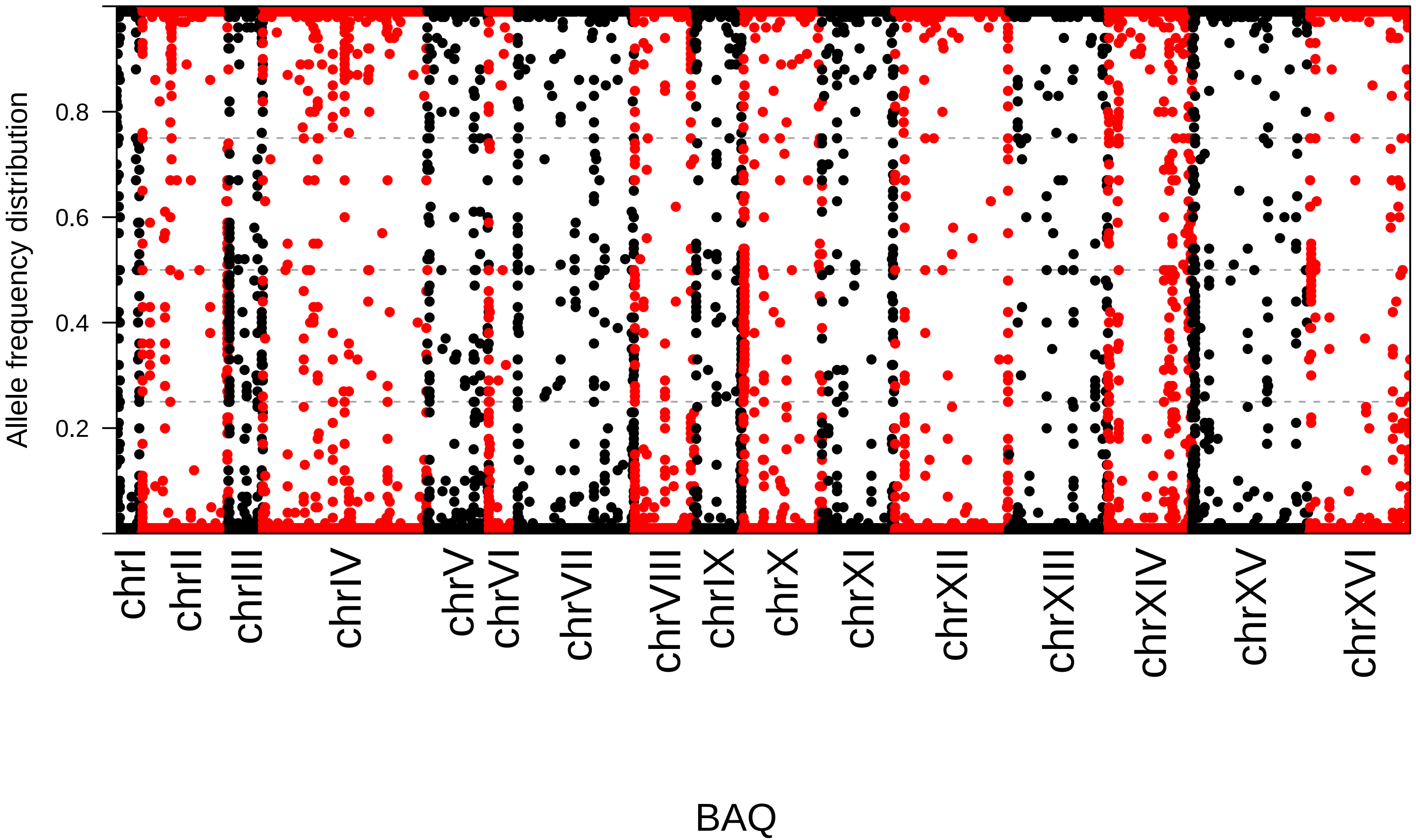
Allele frequency distribution
0.2
0.4
0.6
0.8
chrI
chrII
chrIII
chrIV
chrV
chrVI
chrVII
chrVIII
chrIX
chrX
chrXI
chrXII
chrXIII
chrXIV
chrXV
chrXVI
BAQ

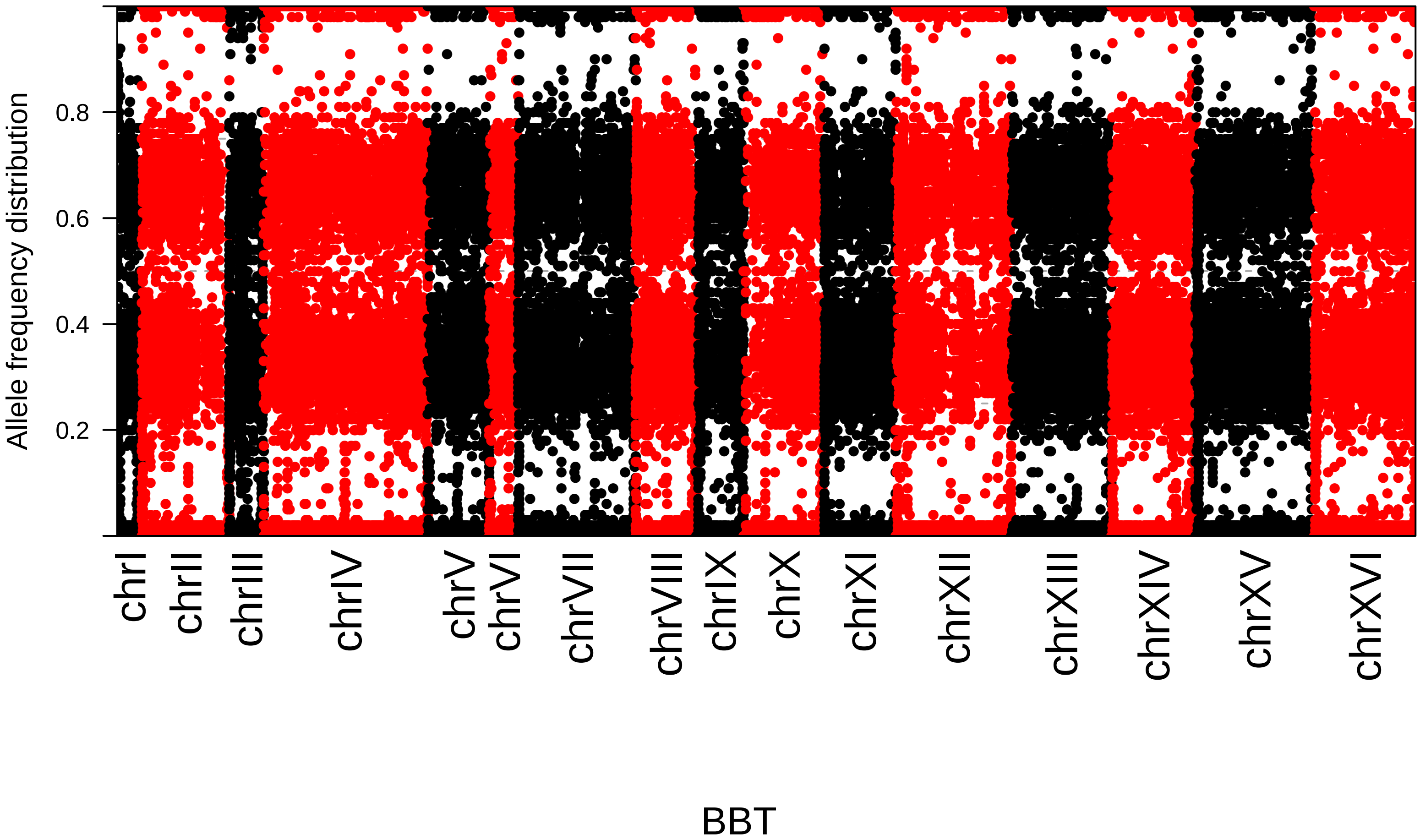
Allele frequency distribution
0.2
0.4
0.6
0.8
chrI
chrII
chrIII
chrIV
chrV
chrVI
chrVII
chrVIII
chrIX
chrX
chrXI
chrXII
chrXIII
chrXIV
chrXV
chrXVI
BBT

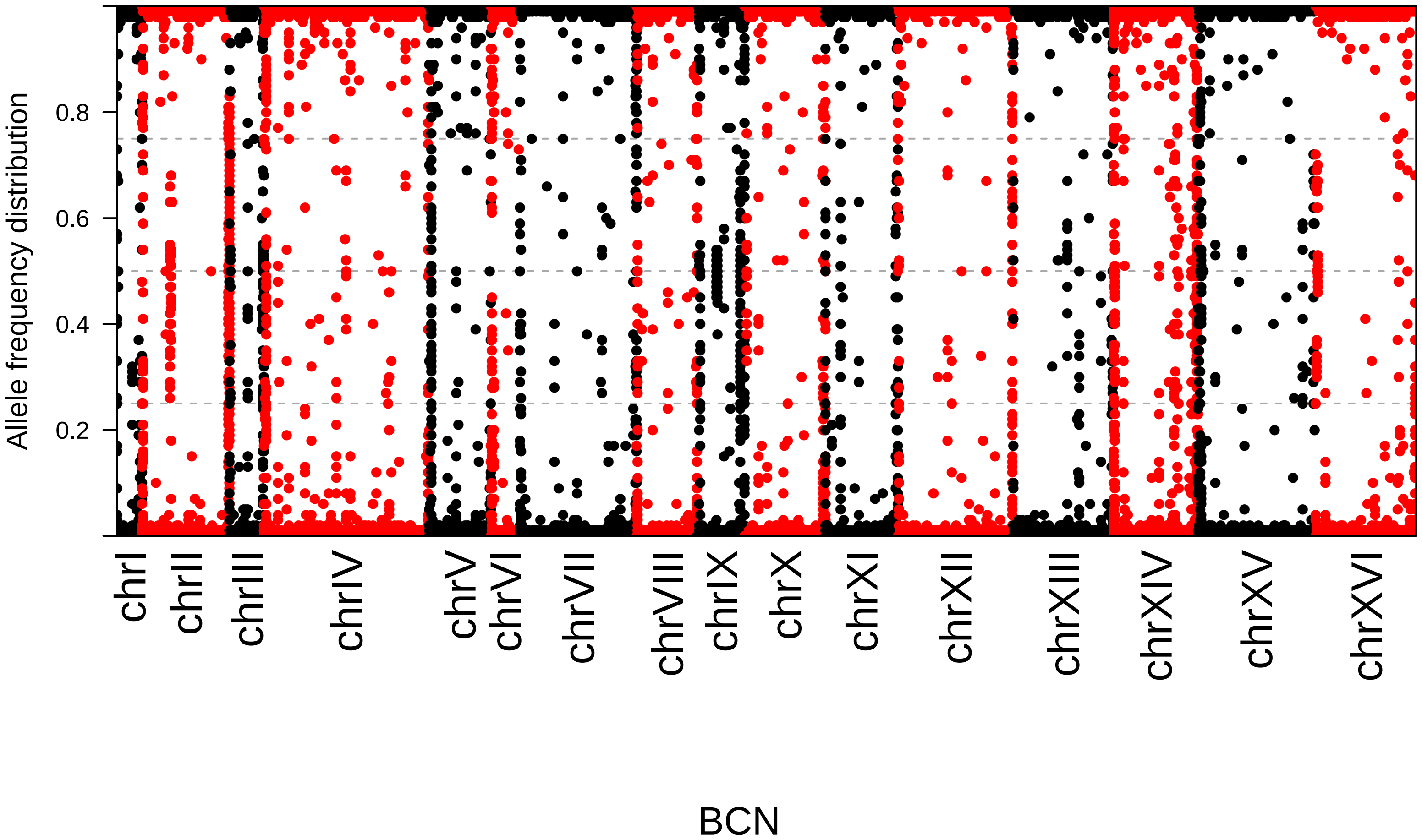

Allele frequency distribution
0.2
0.4
0.6
0.8
chrI
chrII
chrIII
chrIV
chrV
chrVI
chrVII
chrVIII
chrIX
chrX
chrXI
chrXII
chrXIII
chrXIV
chrXV
chrXVI
BCN

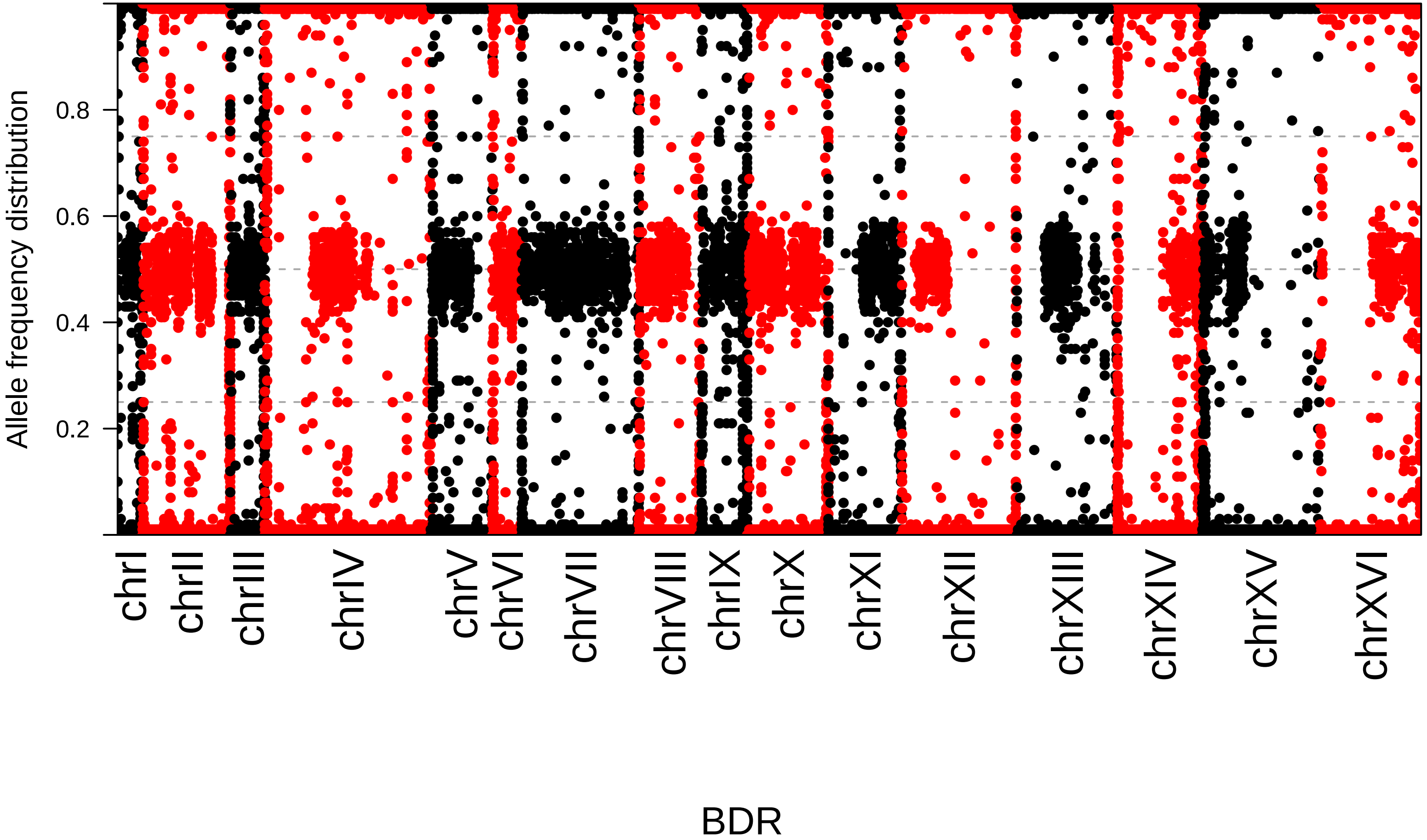

Allele frequency distribution
0.2
0.4
0.6
0.8
chrI
chrII
chrIII
chrIV
chrV
chrVI
chrVII
chrVIII
chrIX
chrX
chrXI
chrXII
chrXIII
chrXIV
chrXV
chrXVI
BDR

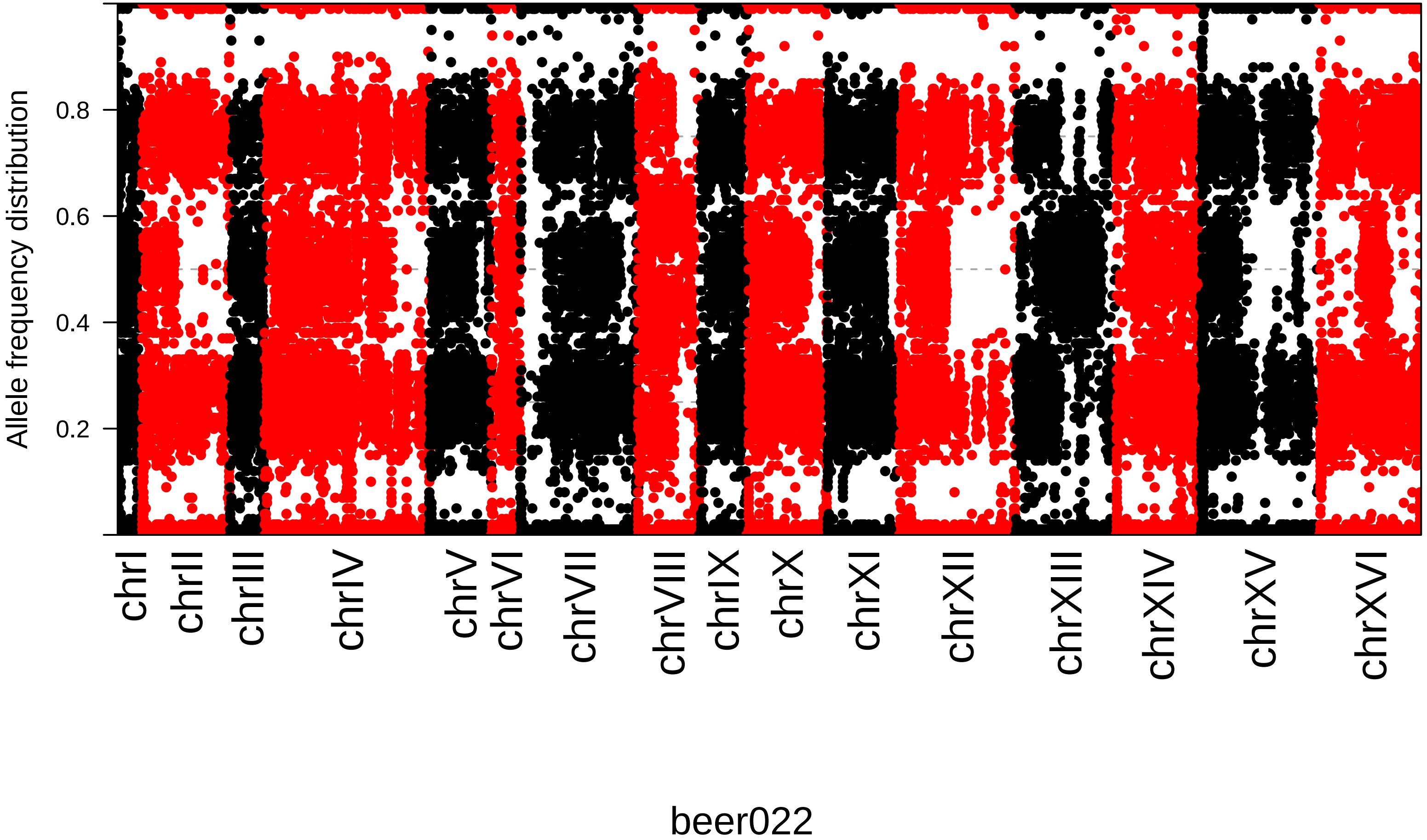

Allele frequency distribution
0.2
0.4
0.6
0.8
chrI
chrII
chrIII
chrIV
chrV
chrVI
chrVII
chrVIII
chrIX
chrX
chrXI
chrXII
chrXIII
chrXIV
chrXV
chrXVI
beer022

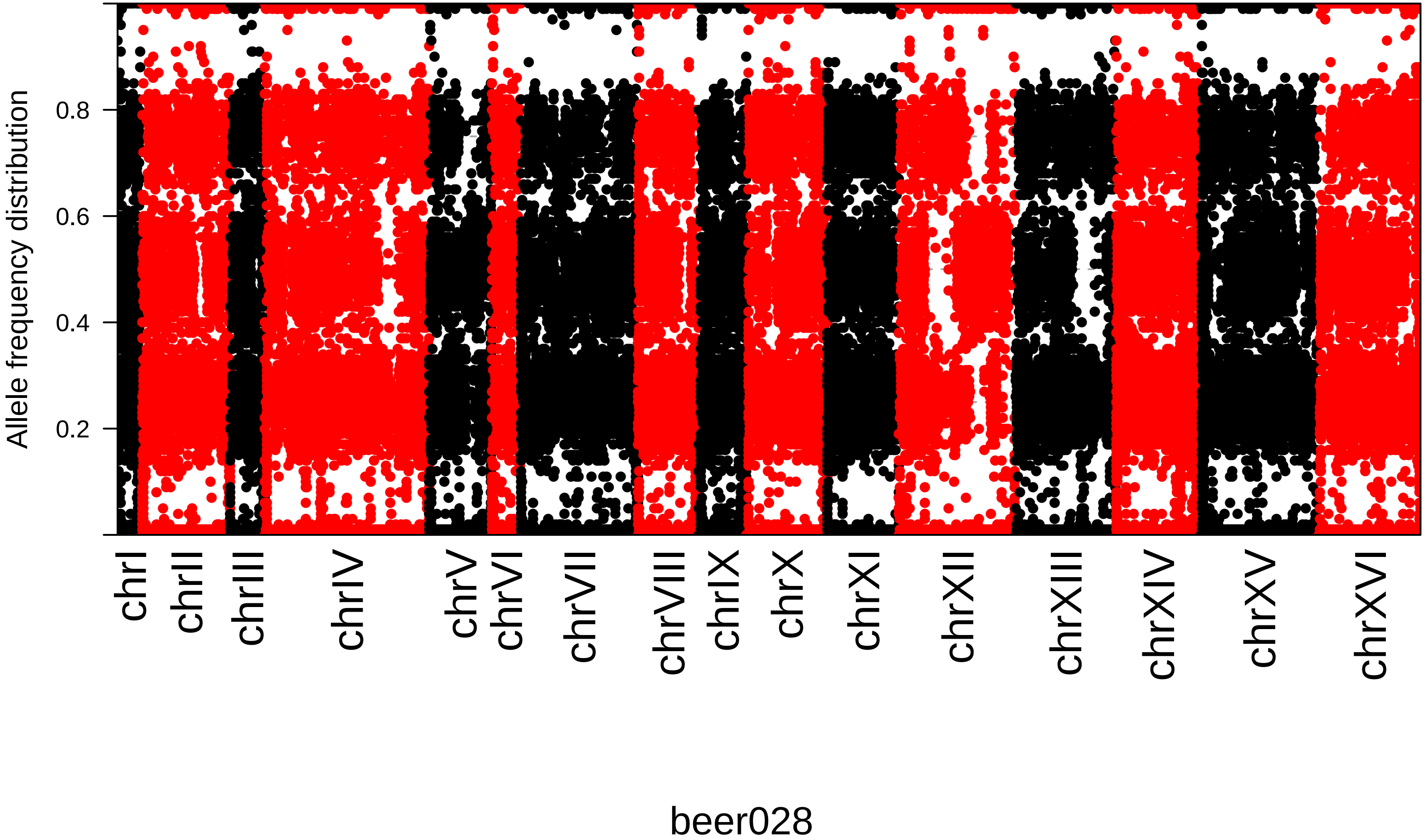
Allele frequency distribution
0.2
0.4
0.6
0.8
chrI
chrII
chrIII
chrIV
chrV
chrVI
chrVII
chrVIII
chrIX
chrX
chrXI
chrXII
chrXIII
chrXIV
chrXV
chrXVI
beer028

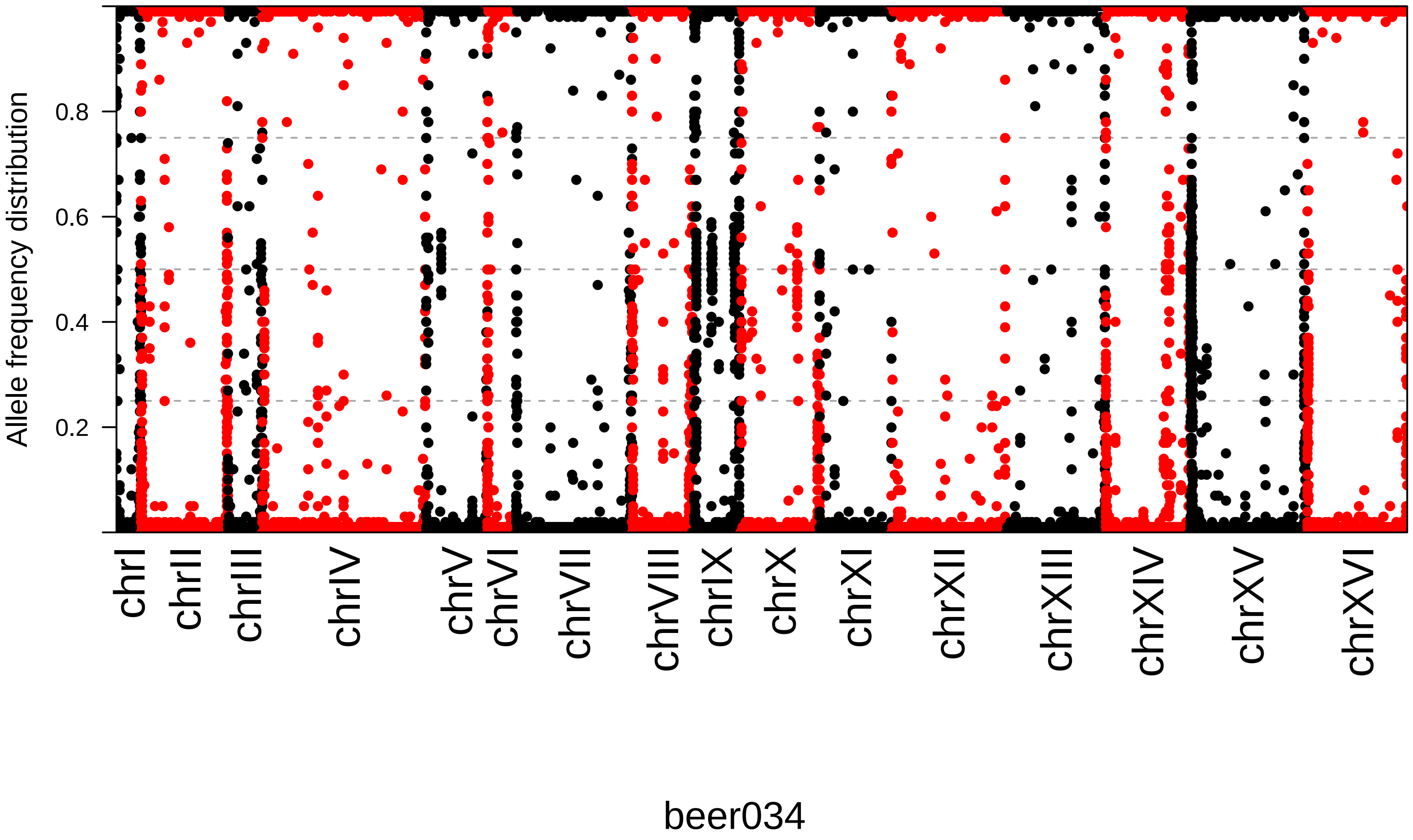
Allele frequency distribution
0.2
0.4
0.6
0.8
chrI
chrII
chrIII
chrIV
chrV
chrVI
chrVII
chrVIII
chrIX
chrX
chrXI
chrXII
chrXIII
chrXIV
chrXV
chrXVI
beer034

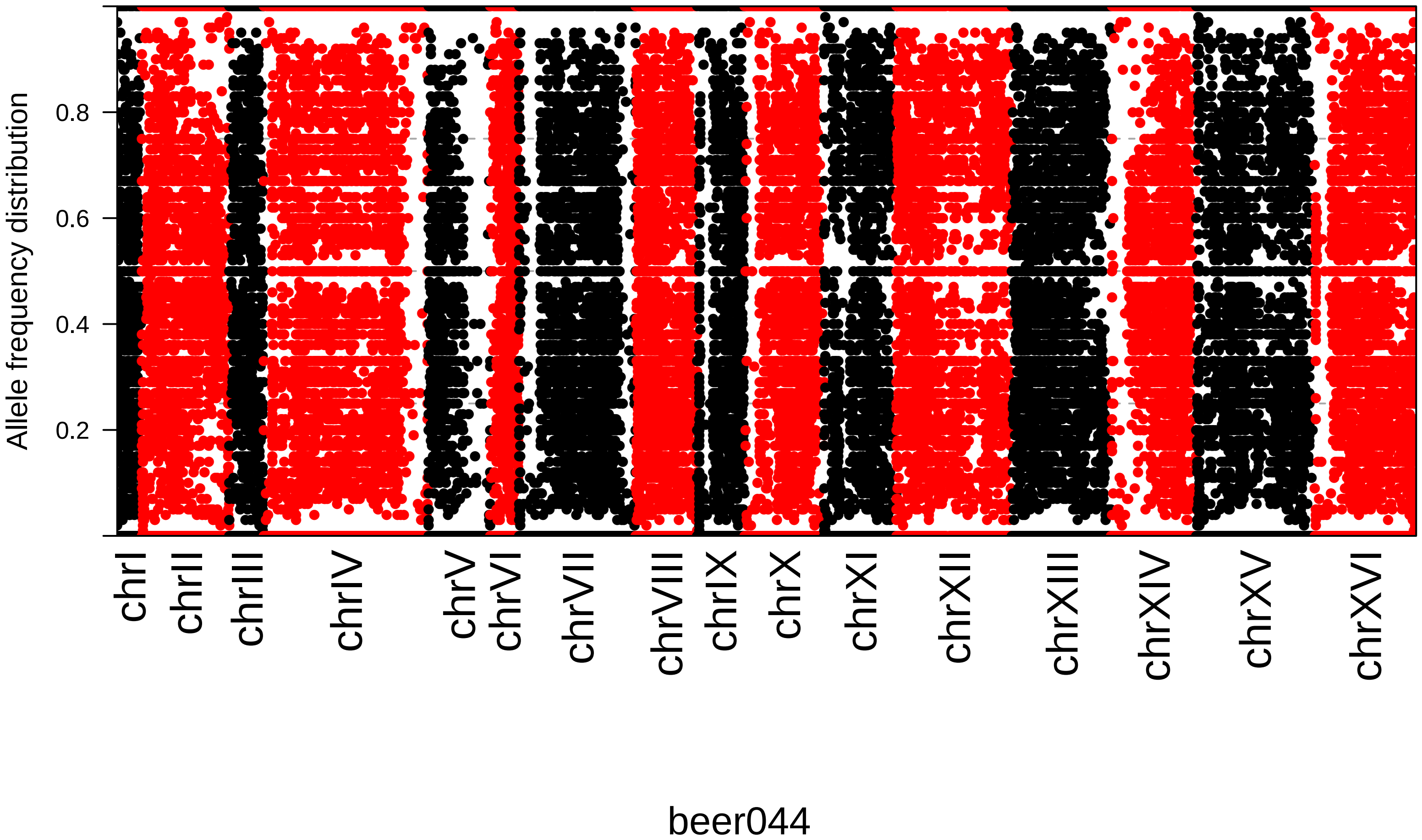

Allele frequency distribution
0.2
0.4
0.6
0.8
chrI
chrII
chrIII
chrIV
chrV
chrVI
chrVII
chrVIII
chrIX
chrX
chrXI
chrXII
chrXIII
chrXIV
chrXV
chrXVI
beer044

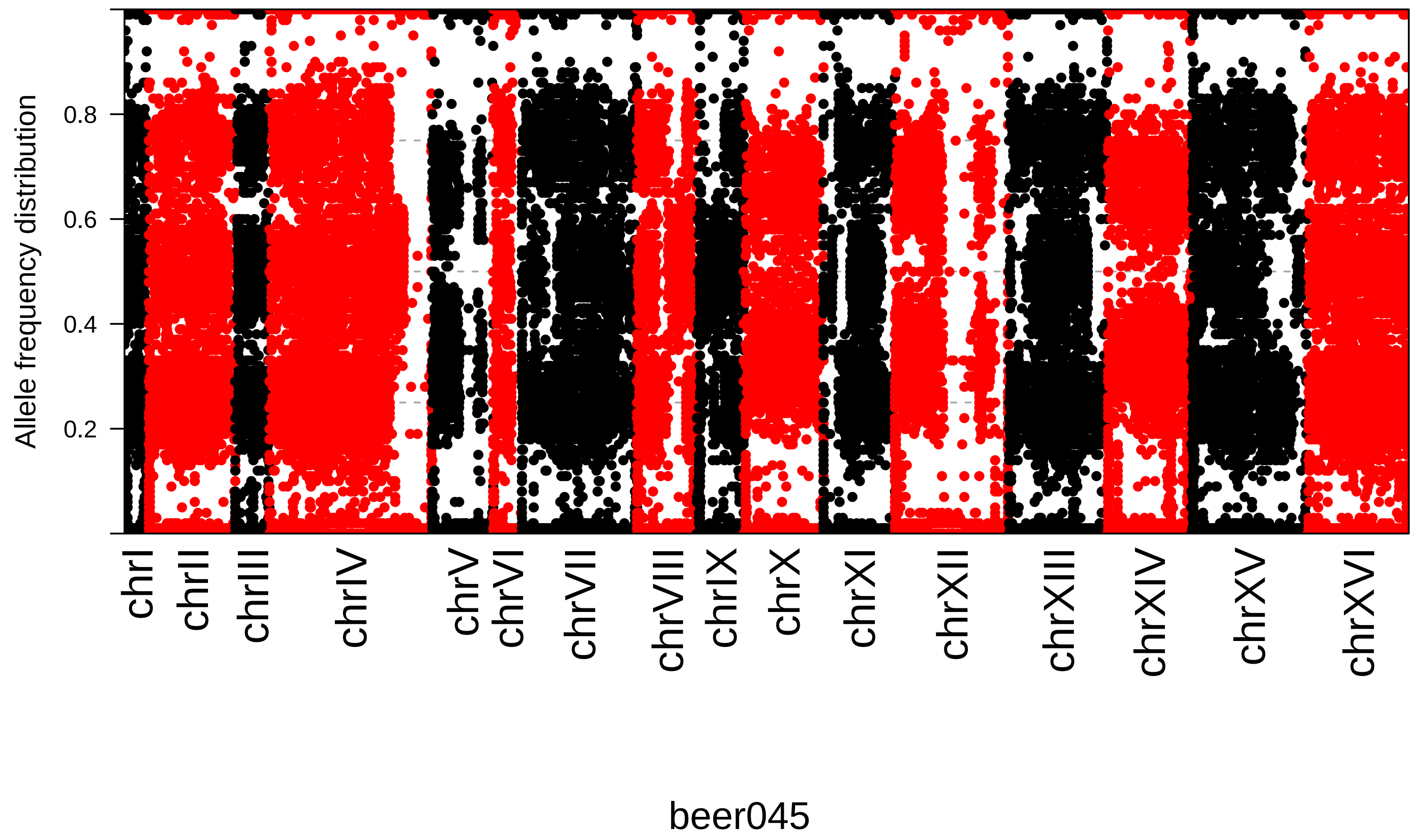

Allele frequency distribution
0.2
0.4
0.6
0.8
chrI
chrII
chrIII
chrIV
chrV
chrVI
chrVII
chrVIII
chrIX
chrX
chrXI
chrXII
chrXIII
chrXIV
chrXV
chrXVI
beer045

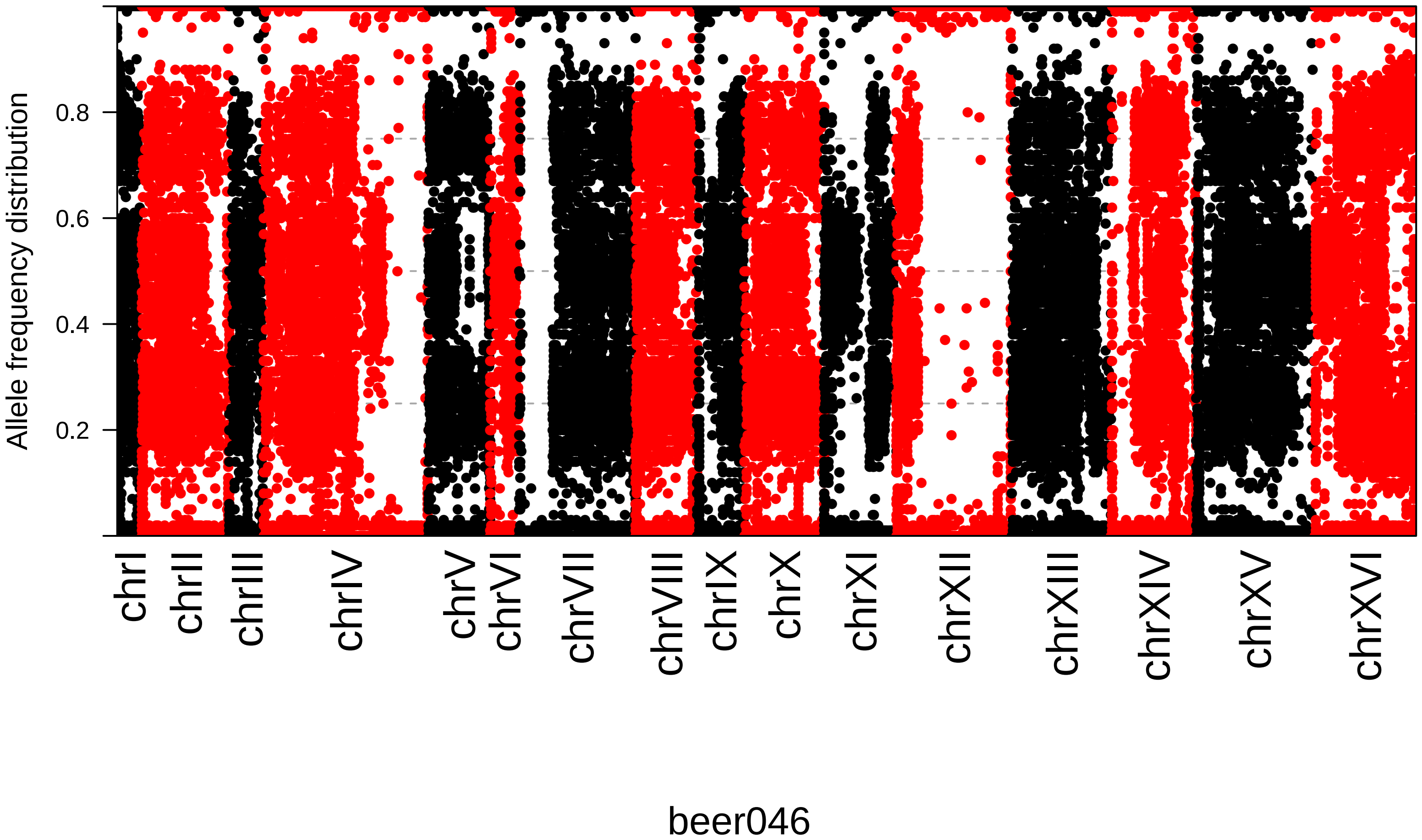
Allele frequency distribution
0.8
0.6
0.4
0.2
chrI
chrII
chrIII
chrIV
chrV
chrVI
chrVII
chrVIII
chrIX
chrX
chrXI
chrXII
chrXIII
chrXIV
chrXV
chrXVI
beer046

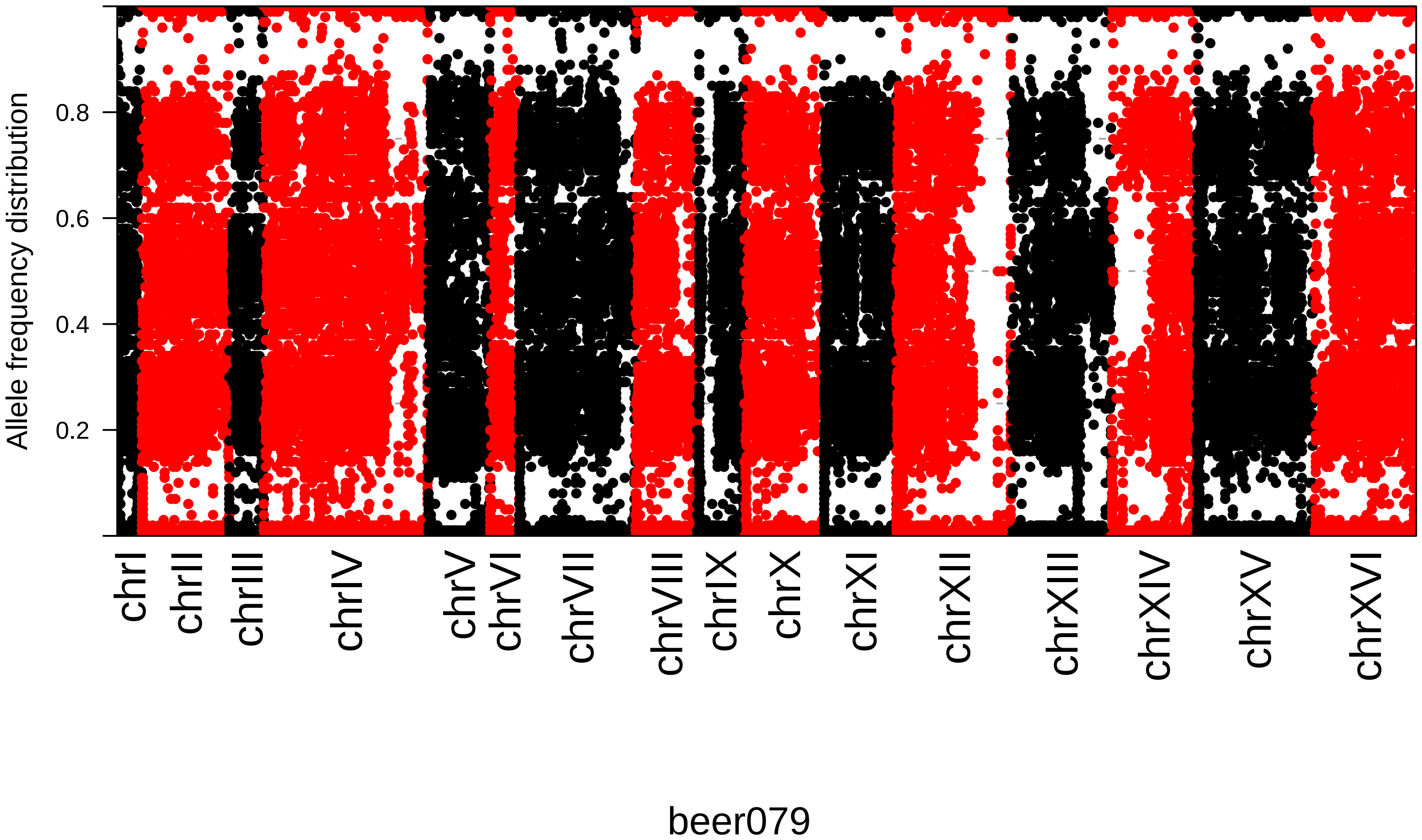

Allele frequency distribution
0.2
0.4
0.6
0.8
chrI
chrII
chrIII
chrIV
chrV
chrVI
chrVII
chrVIII
chrIX
chrX
chrXI
chrXII
chrXIII
chrXIV
chrXV
chrXVI
beer079

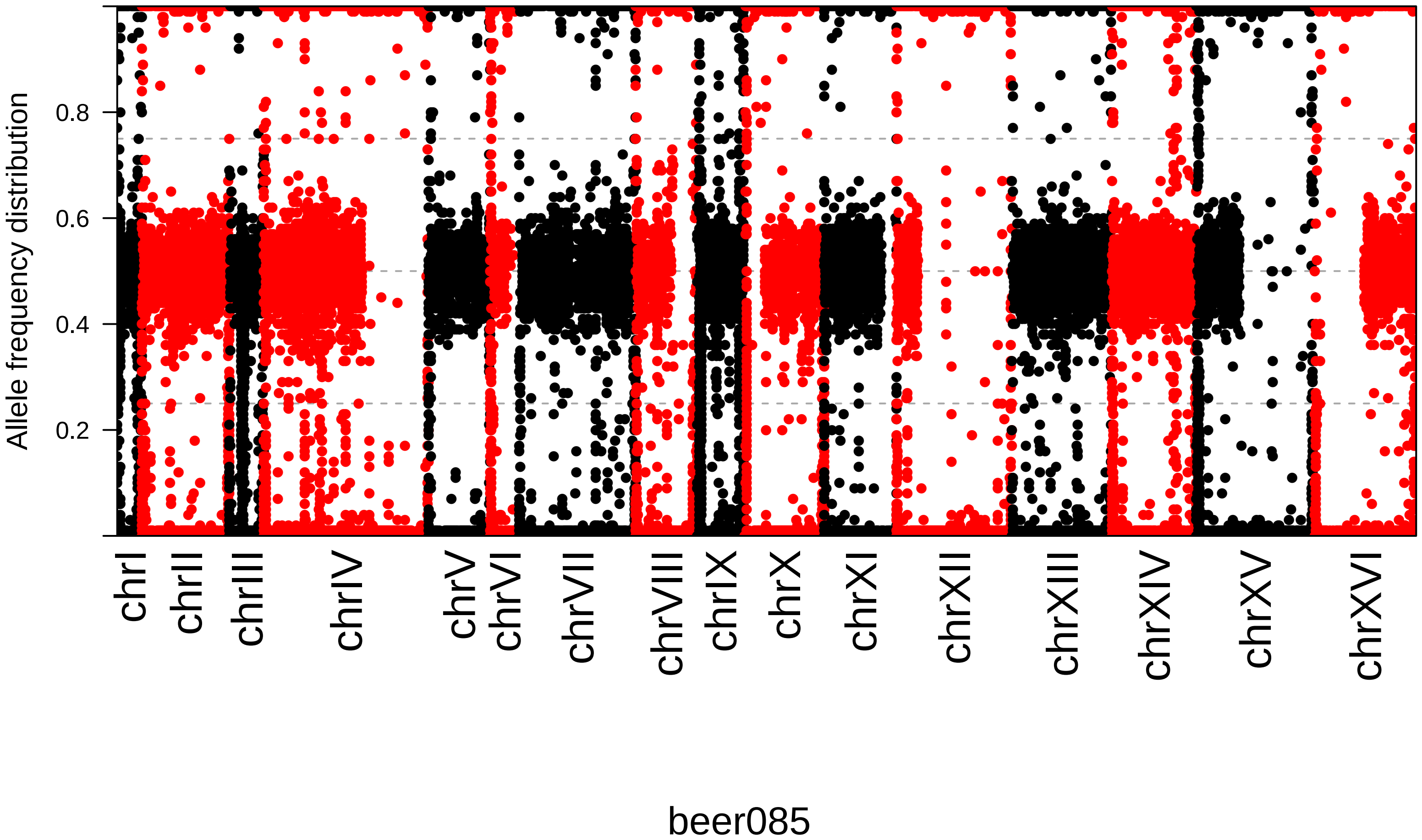

Allele frequency distribution
0.8
0.6
0.4
0.2
chrI
chrII
chrIII
chrIV
chrV
chrVI
chrVII
chrVIII
chrIX
chrX
chrXI
chrXII
chrXIII
chrXIV
chrXV
chrXVI
beer085

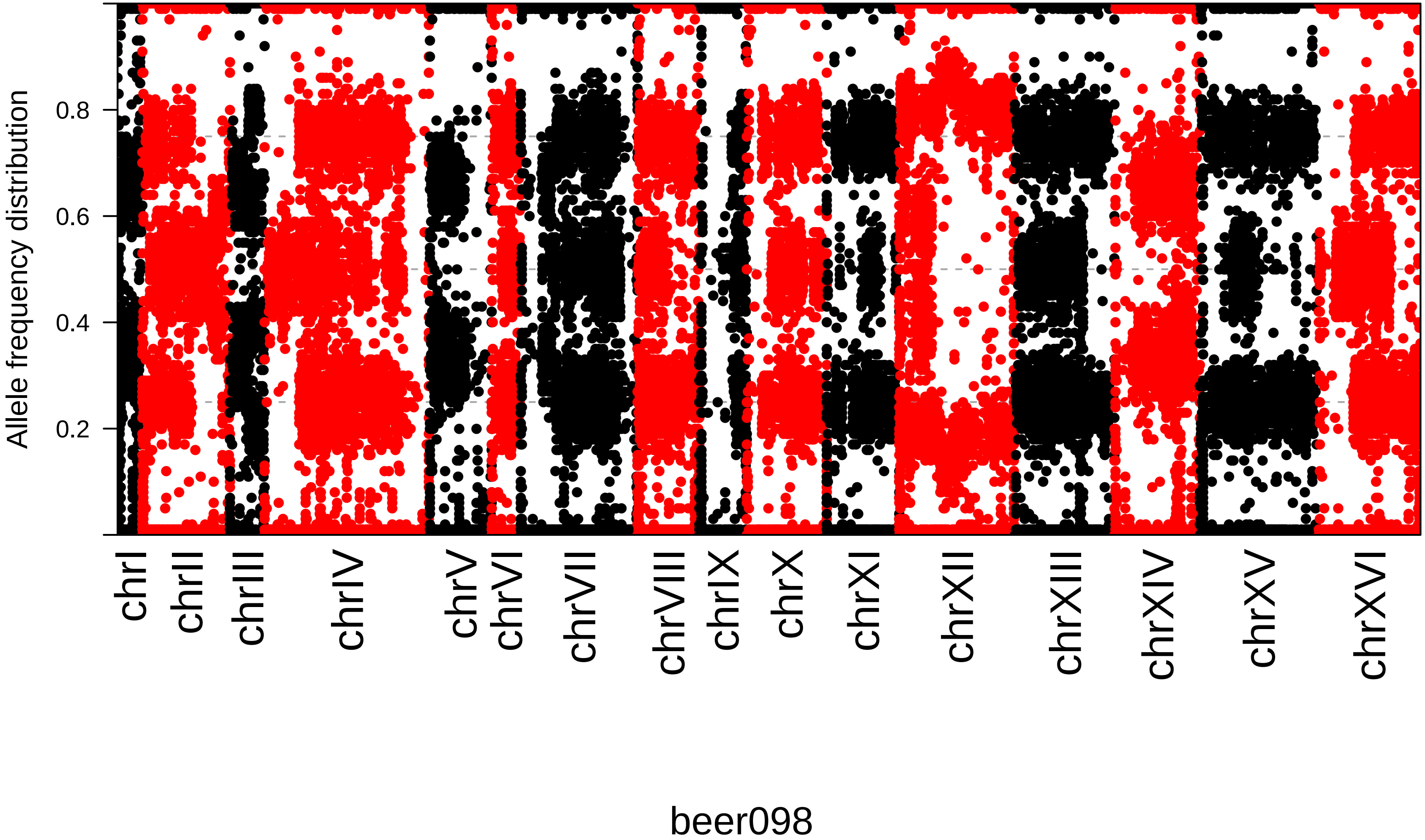
Allele frequency distribution
0.2
0.4
0.6
0.8
chrI
chrII
chrIII
chrIV
chrV
chrVI
chrVII
chrVIII
chrIX
chrX
chrXI
chrXII
chrXIII
chrXIV
chrXV
chrXVI
beer098

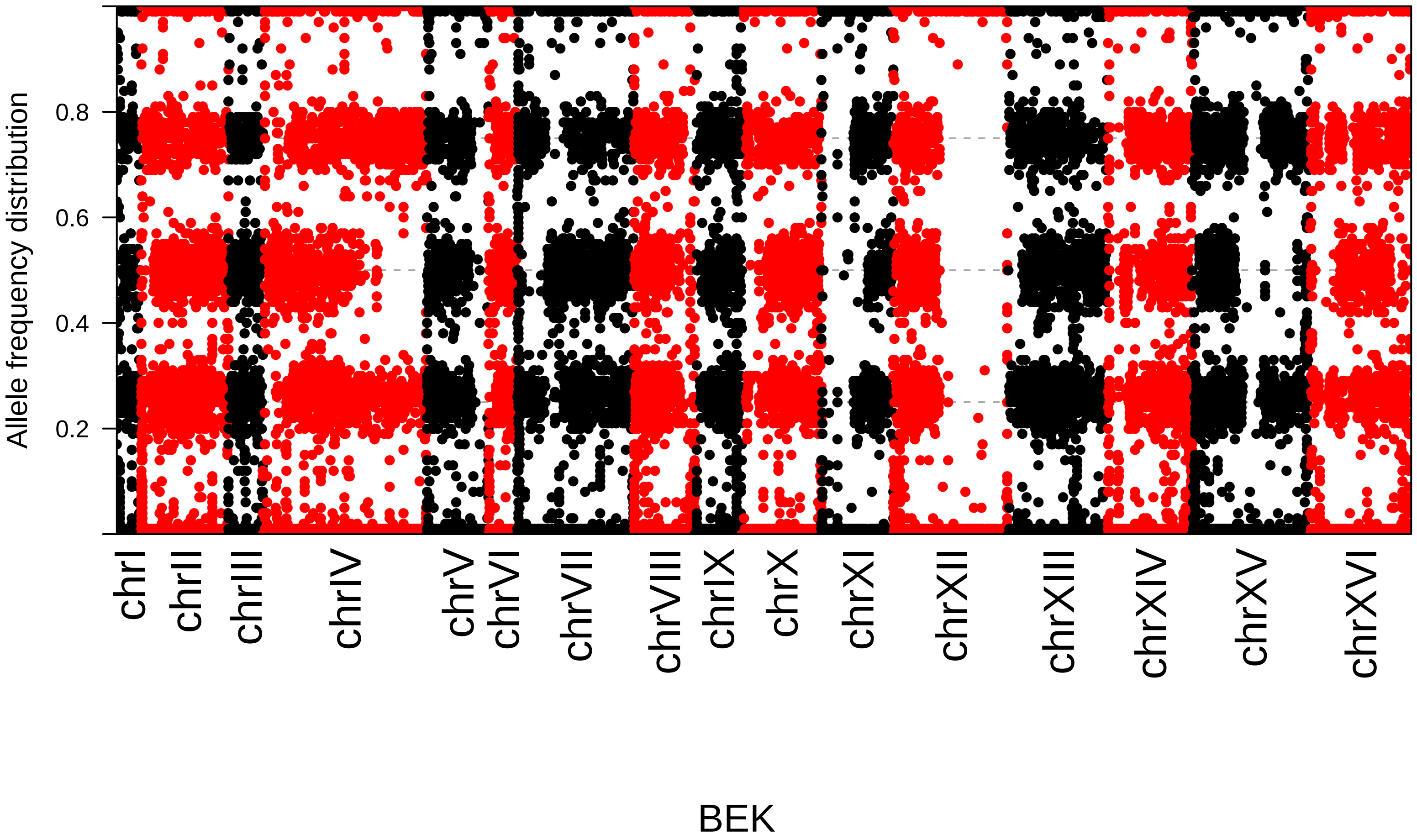

Allele frequency distribution
0.2
0.4
0.6
0.8
chrI
chrII
chrIII
chrIV
chrV
chrVI
chrVII
chrVIII
chrIX
chrX
chrXI
chrXII
chrXIII
chrXIV
chrXV
chrXVI
BEK

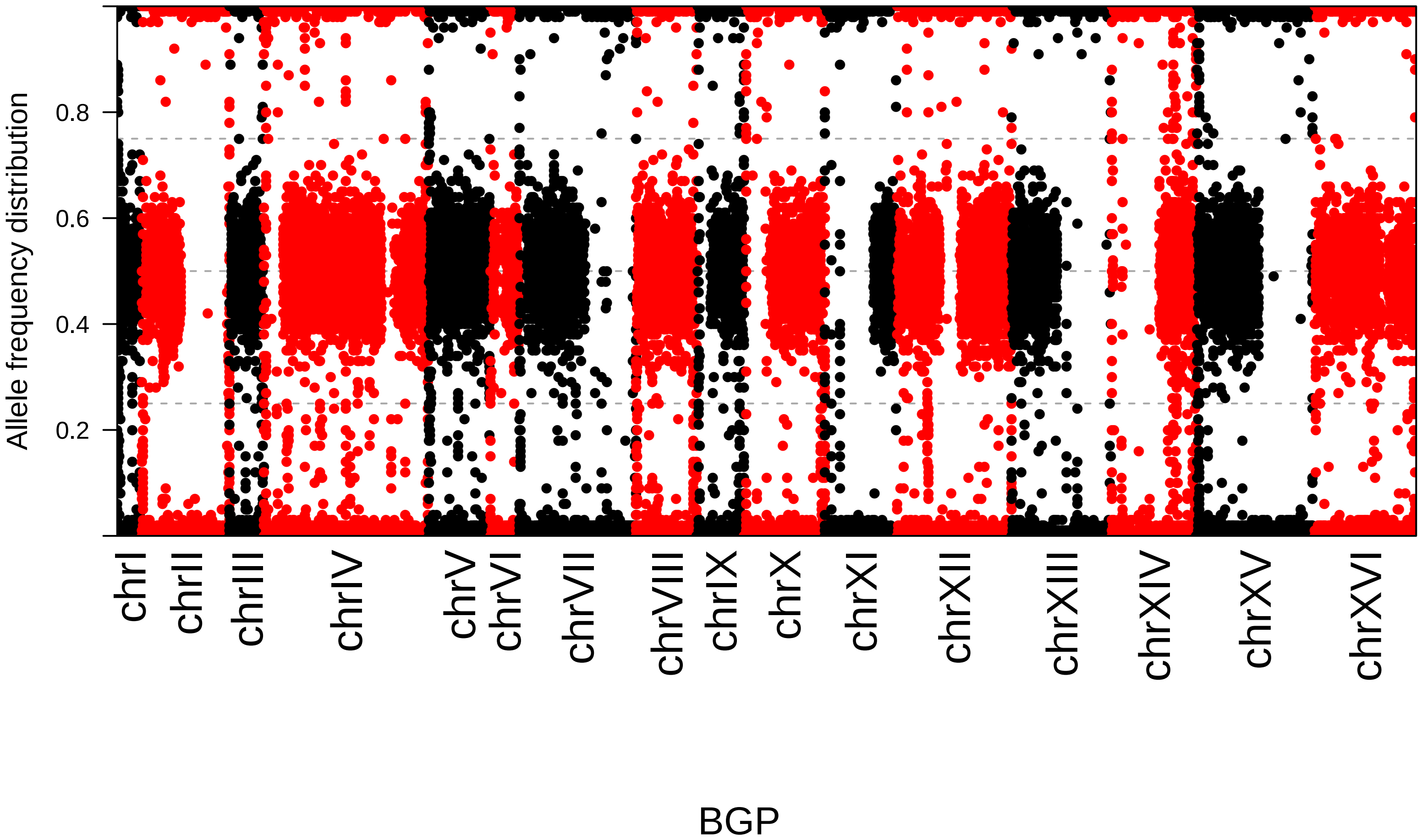

Allele frequency distribution
0.2
0.4
0.6
0.8
chrI
chrII
chrIII
chrIV
chrV
chrVI
chrVII
chrVIII
chrIX
chrX
chrXI
chrXII
chrXIII
chrXIV
chrXV
chrXVI
BGP

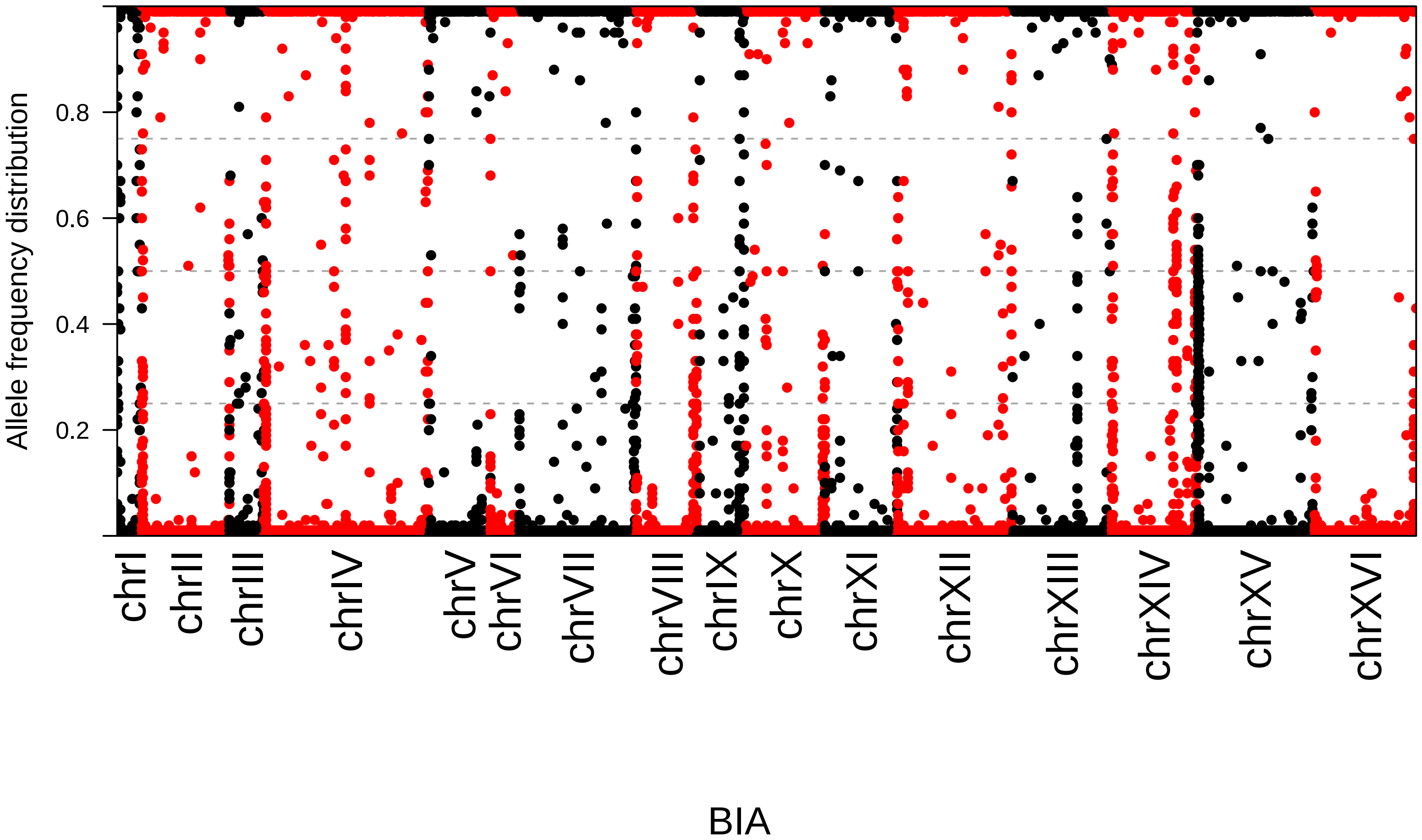
Allele frequency distribution
0.2
0.4
0.6
0.8
chrI
chrII
chrIII
chrIV
chrV
chrVI
chrVII
chrVIII
chrIX
chrX
chrXI
chrXII
chrXIII
chrXIV
chrXV
chrXVI
BIA

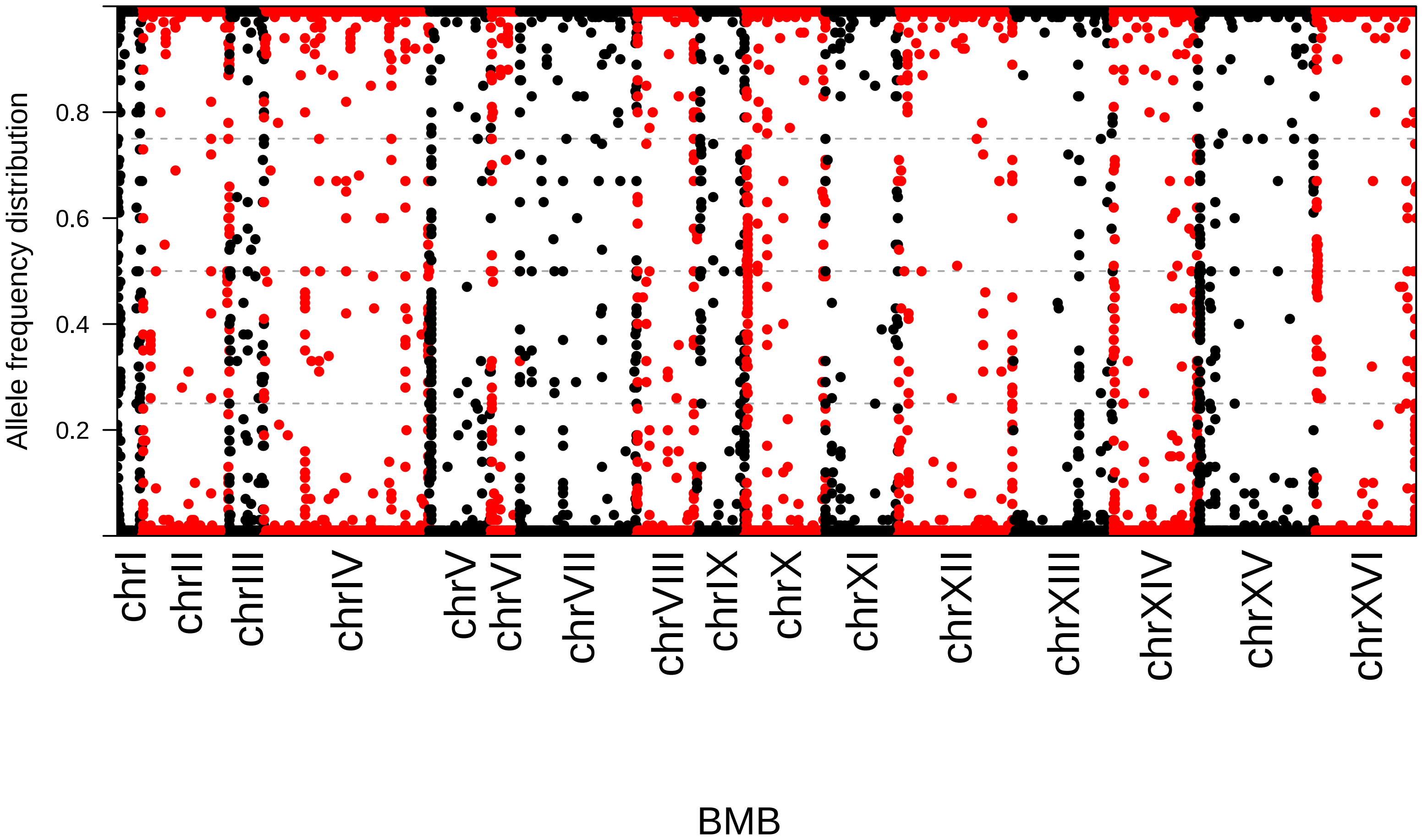

Allele frequency distribution
0.2
0.4
0.6
0.8
chrI
chrII
chrIII
chrIV
chrV
chrVI
chrVII
chrVIII
chrIX
chrX
chrXI
chrXII
chrXIII
chrXIV
chrXV
chrXVI
BMB

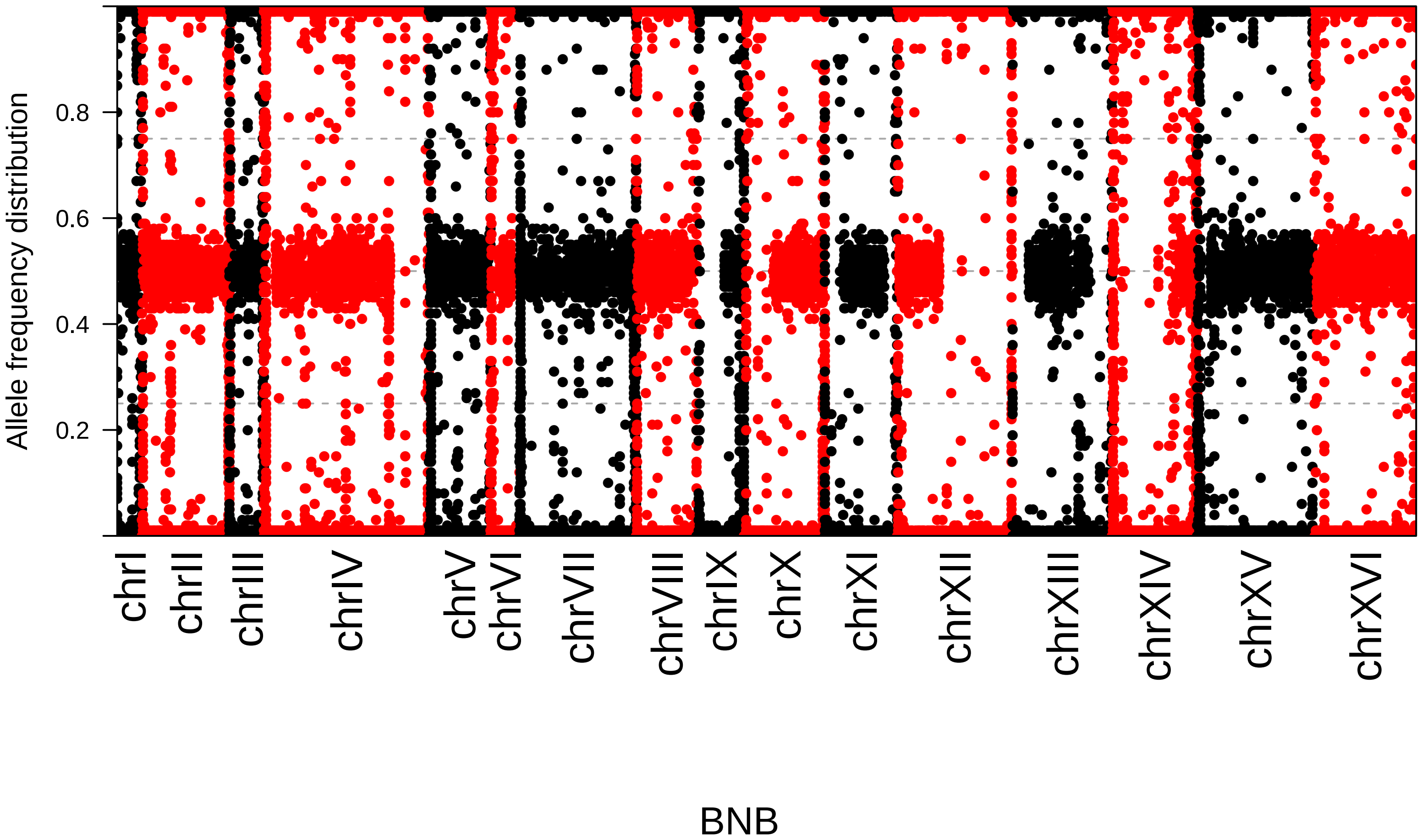

Allele frequency distribution
0.2
0.4
0.6
0.8
chrI
chrII
chrIII
chrIV
chrV
chrVI
chrVII
chrVIII
chrIX
chrX
chrXI
chrXII
chrXIII
chrXIV
chrXV
chrXVI
BNB

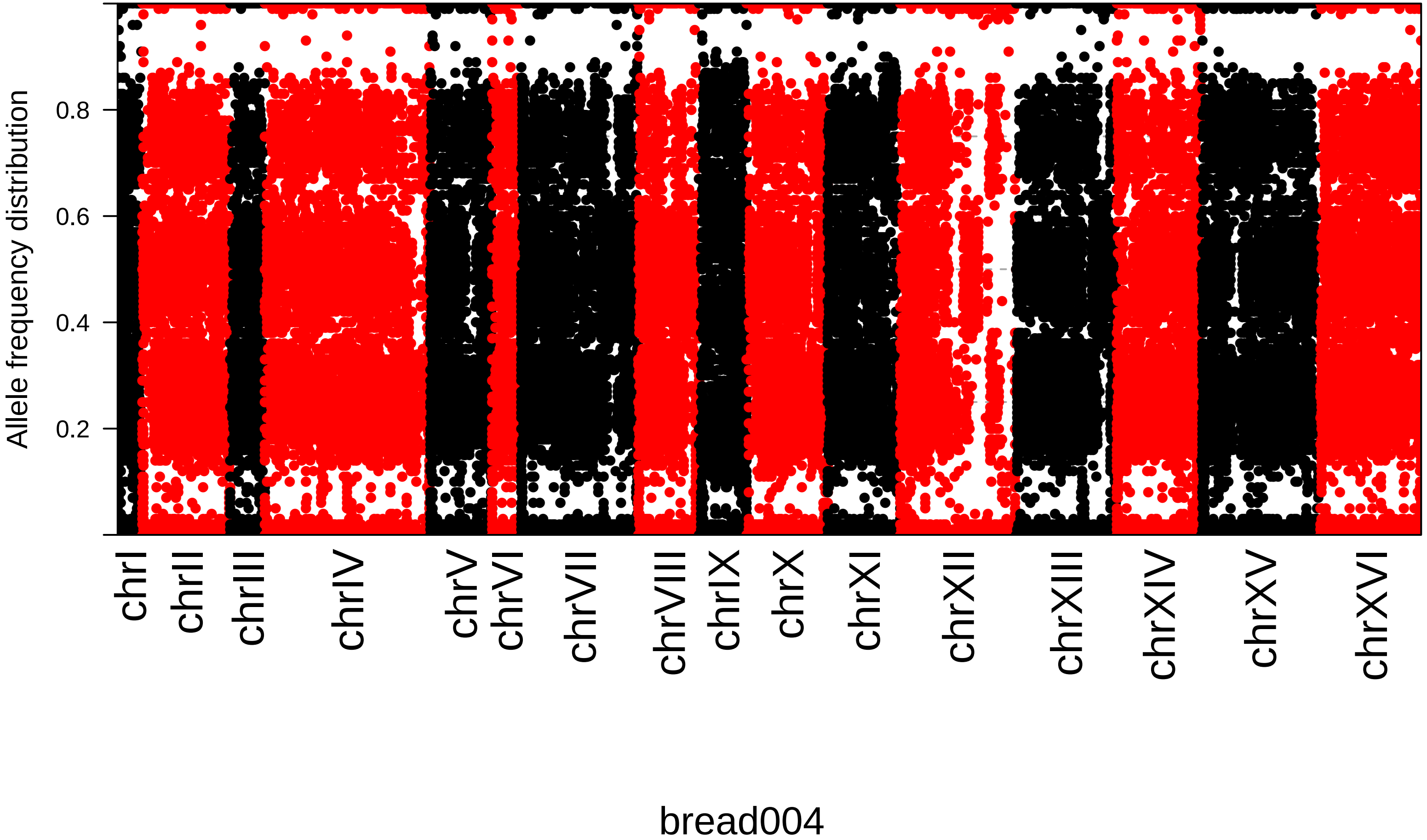
Allele frequency distribution
0.2
0.4
0.6
0.8
chrI
chrII
chrIII
chrIV
chrV
chrVI
chrVII
chrVIII
chrIX
chrX
chrXI
chrXII
chrXIII
chrXIV
chrXV
chrXVI
bread004

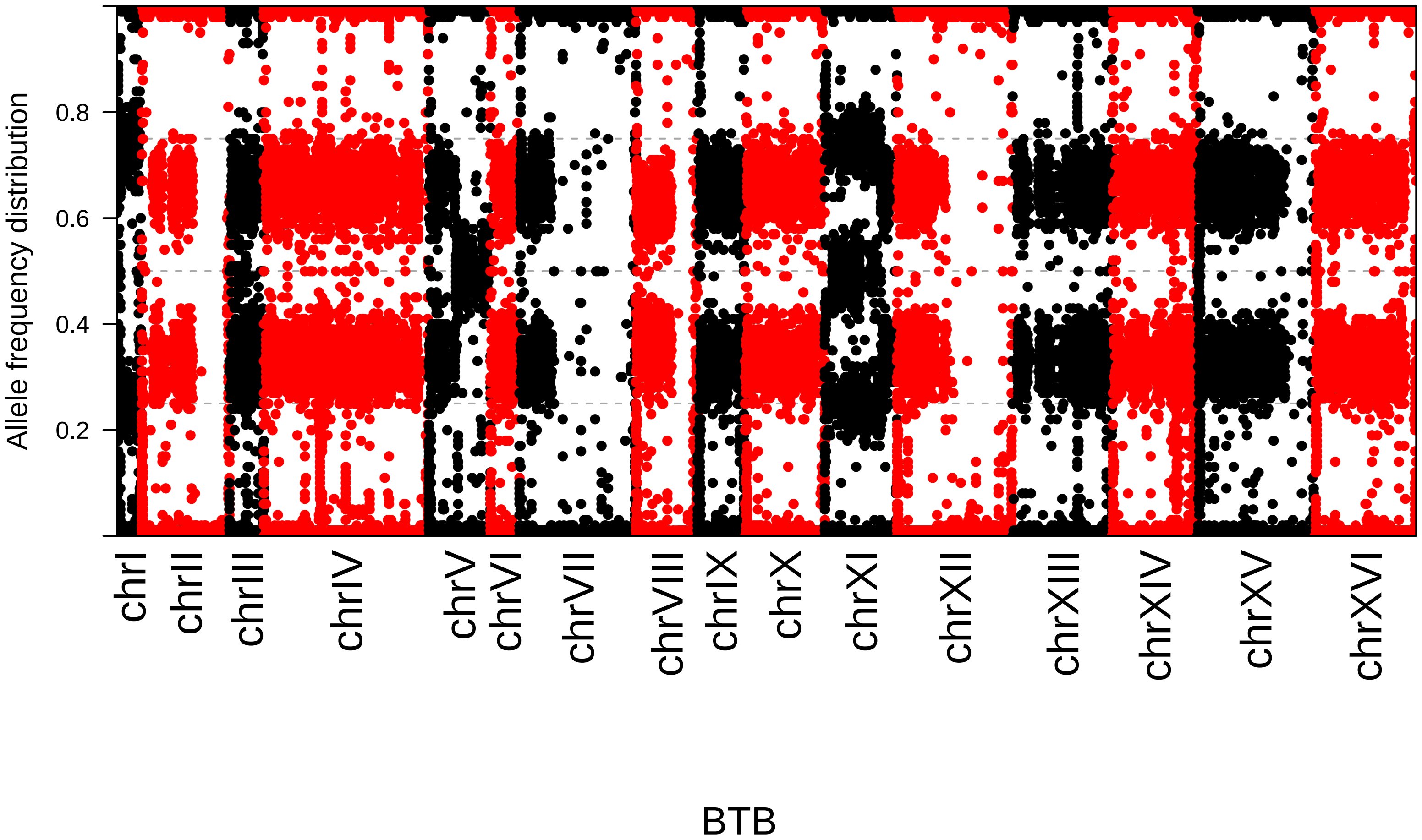

Allele frequency distribution
0.2
0.4
0.6
0.8
chrI
chrII
chrIII
chrIV
chrV
chrVI
chrVII
chrVIII
chrIX
chrX
chrXI
chrXII
chrXIII
chrXIV
chrXV
chrXVI
BTB

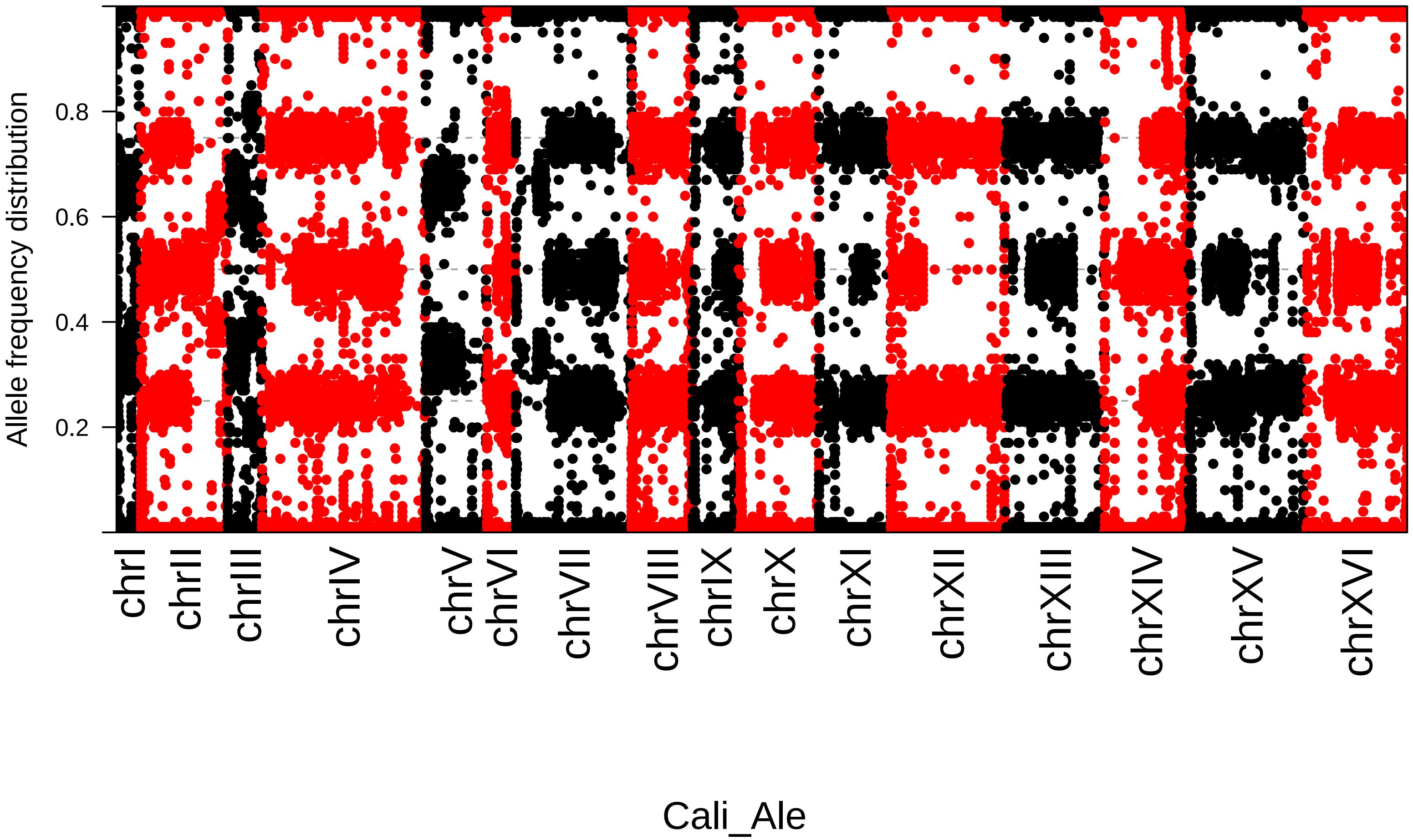

Allele frequency distribution
0.2
0.4
0.6
0.8
chrI
chrII
chrIII
chrIV
chrV
chrVI
chrVII
chrVIII
chrIX
chrX
chrXI
chrXII
chrXIII
chrXIV
chrXV
chrXVI
Cali_Ale

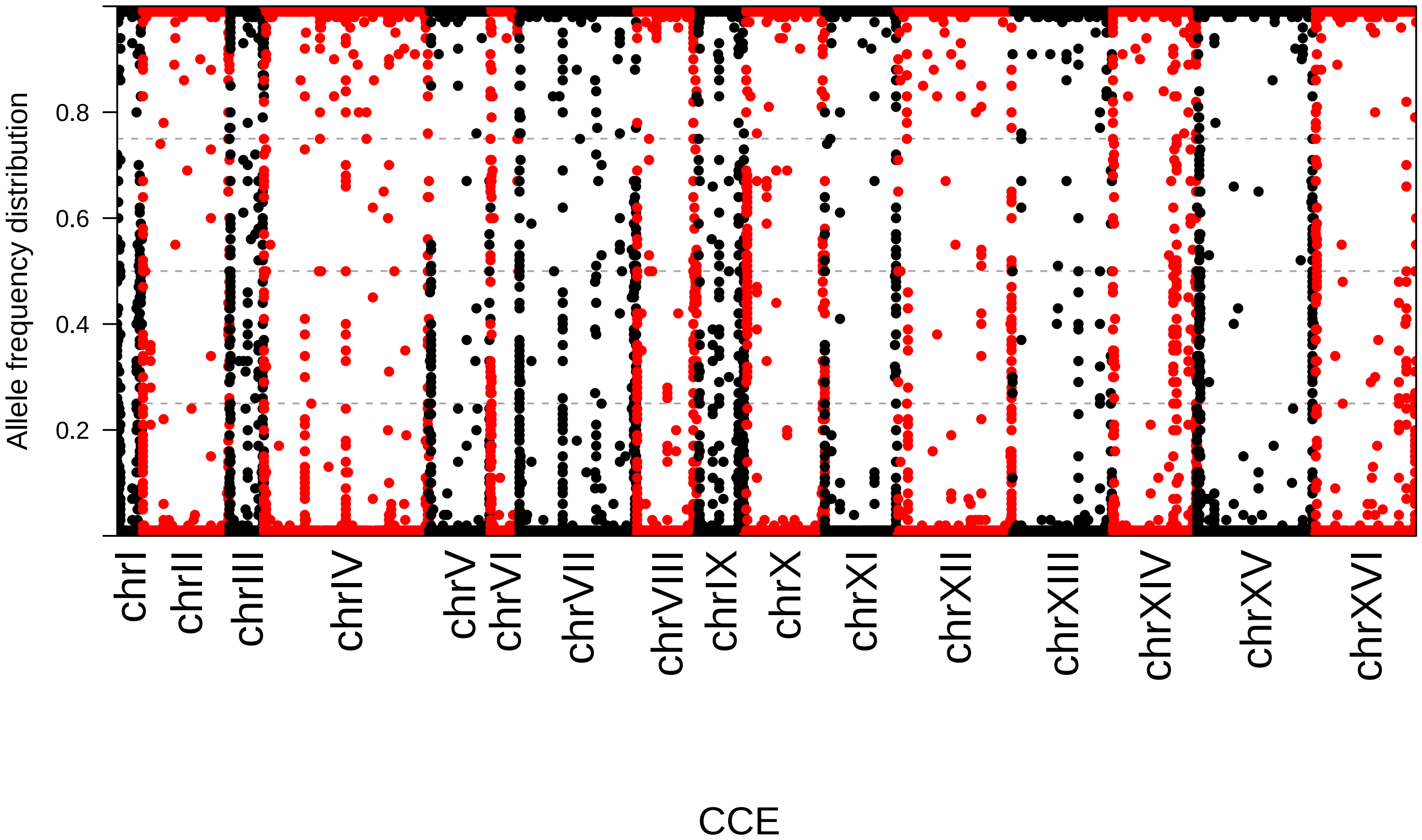

Allele frequency distribution
0.2
0.4
0.6
0.8
chrI
chrII
chrIII
chrIV
chrV
chrVI
chrVII
chrVIII
chrIX
chrX
chrXI
chrXII
chrXIII
chrXIV
chrXV
chrXVI
CCE

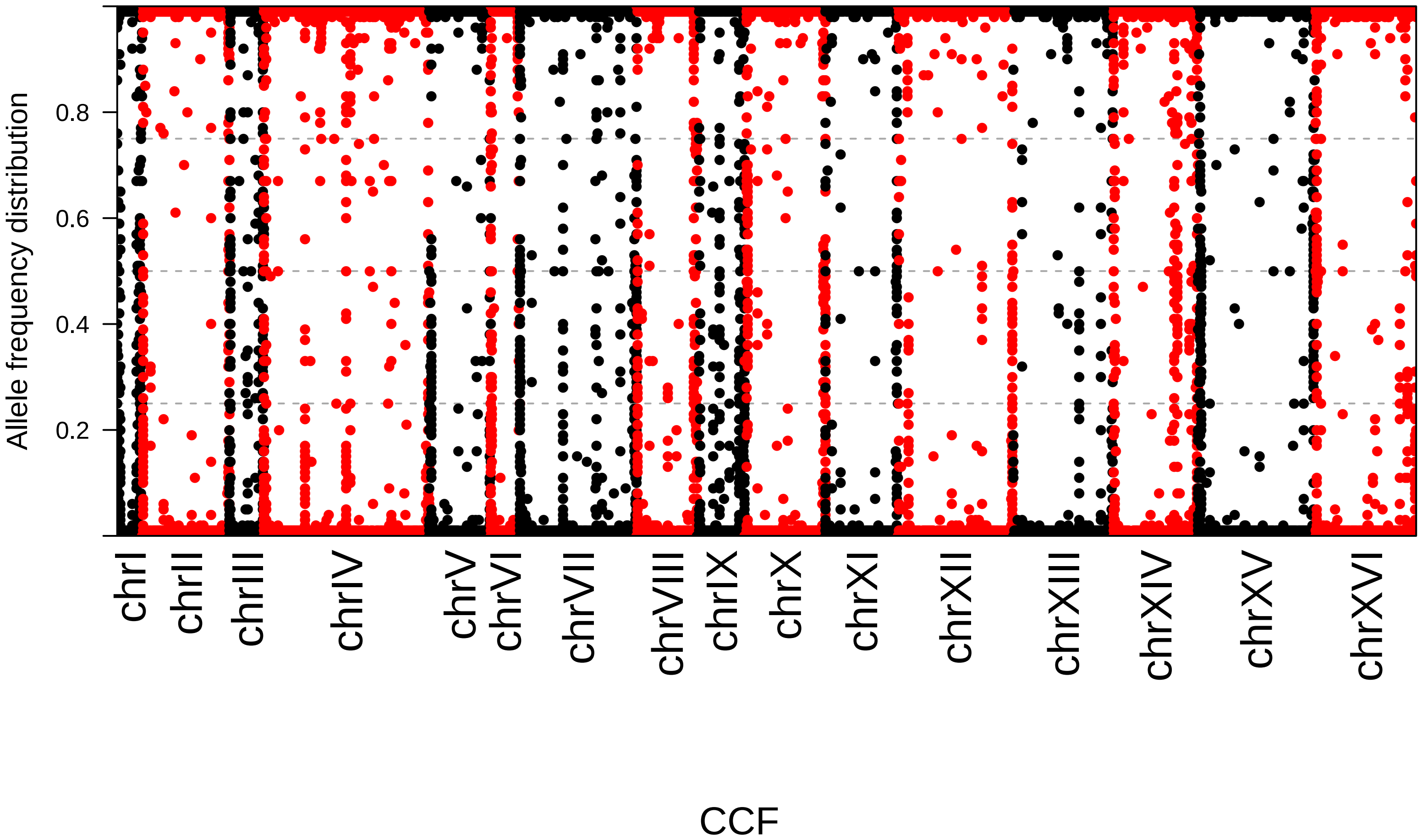

Allele frequency distribution
0.2
0.4
0.6
0.8
chrI
chrII
chrIII
chrIV
chrV
chrVI
chrVII
chrVIII
chrIX
chrX
chrXI
chrXII
chrXIII
chrXIV
chrXV
chrXVI
CCF

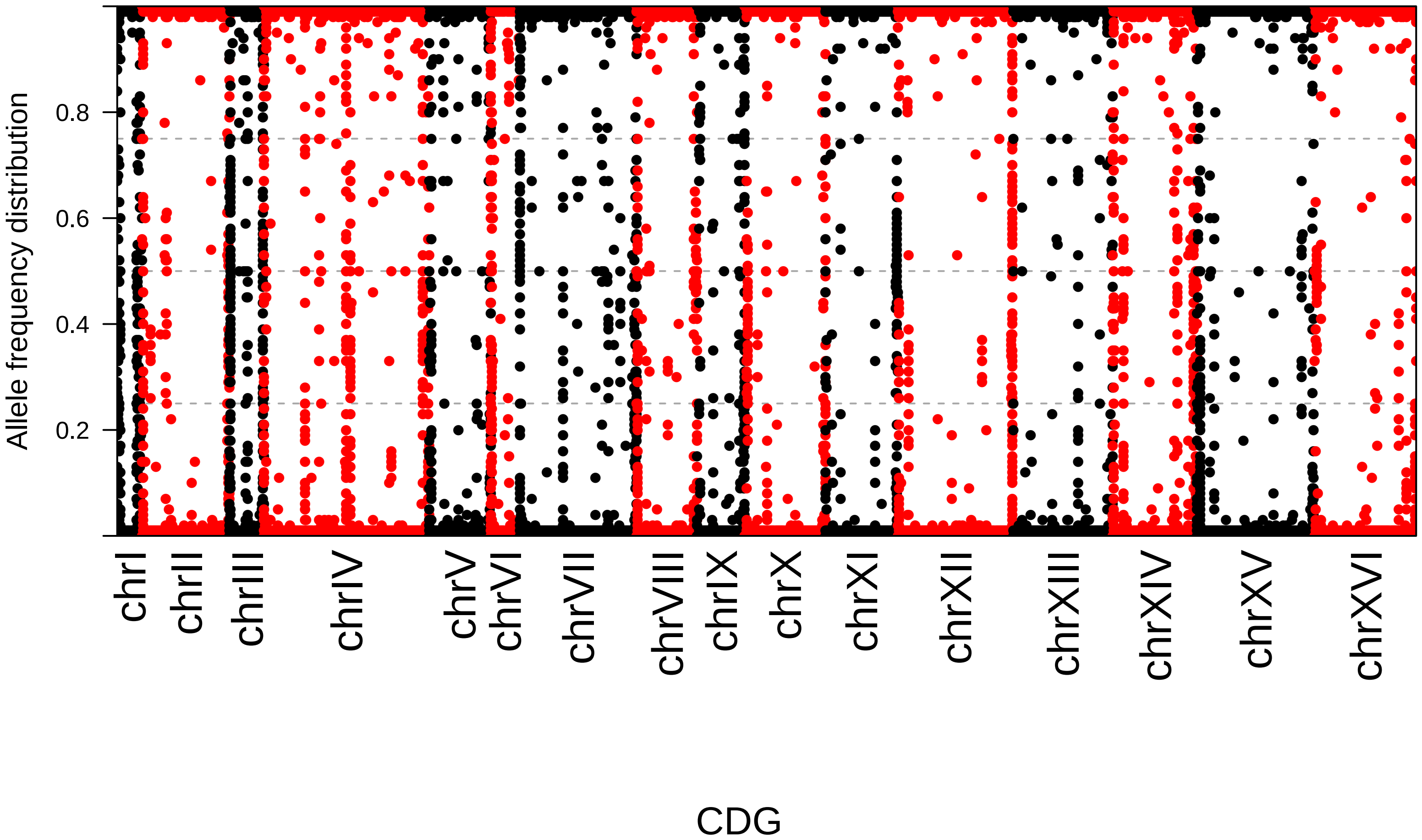
Allele frequency distribution
0.8
0.6
0.4
0.2
chrI
chrII
chrIII
chrIV
chrV
chrVI
chrVII
chrVIII
chrIX
chrX
chrXI
chrXII
chrXIII
chrXIV
chrXV
chrXVI
CDG

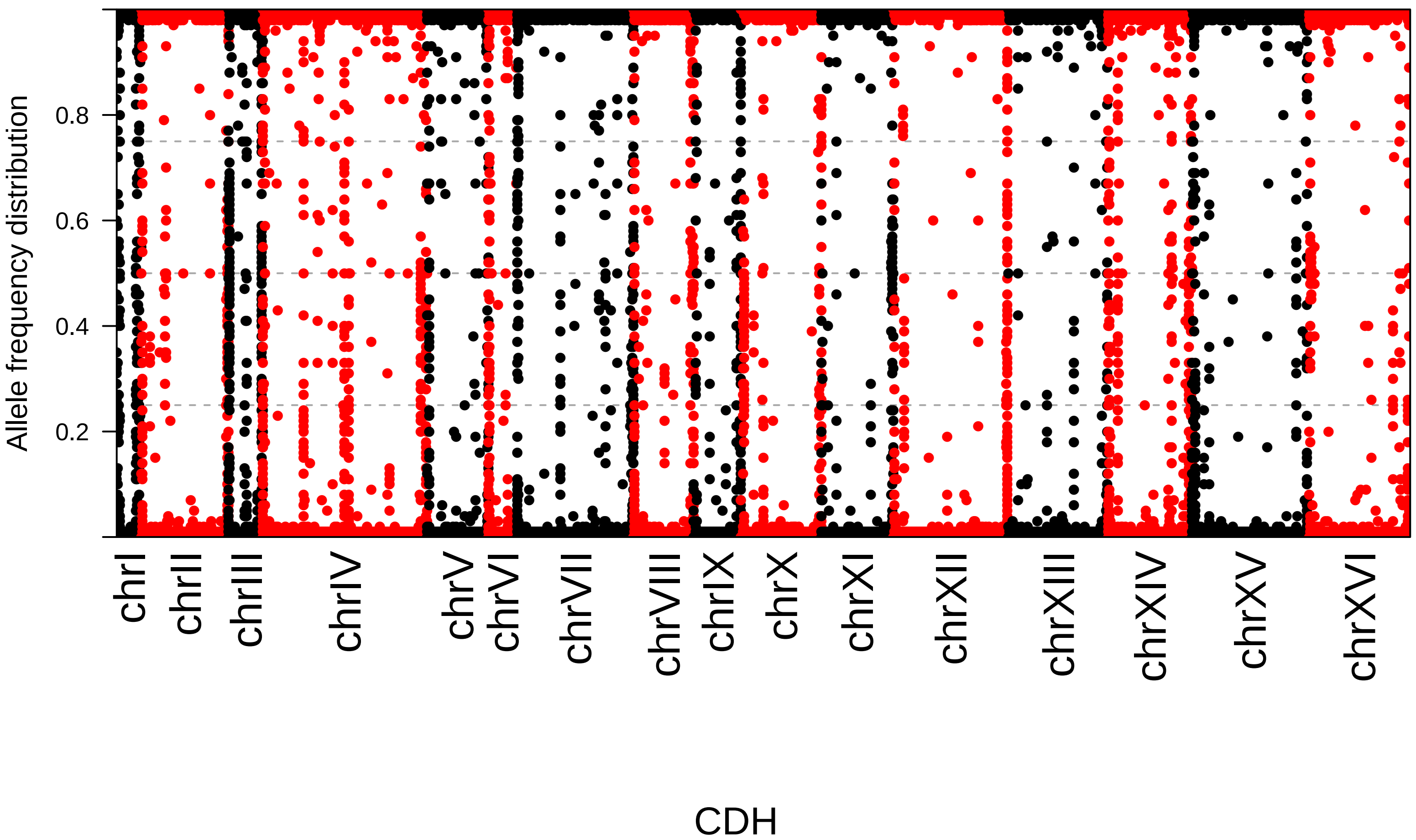
Allele frequency distribution
0.2
0.4
0.6
0.8
chrI
chrII
chrIII
chrIV
chrV
chrVI
chrVII
chrVIII
chrIX
chrX
chrXI
chrXII
chrXIII
chrXIV
chrXV
chrXVI
CDH

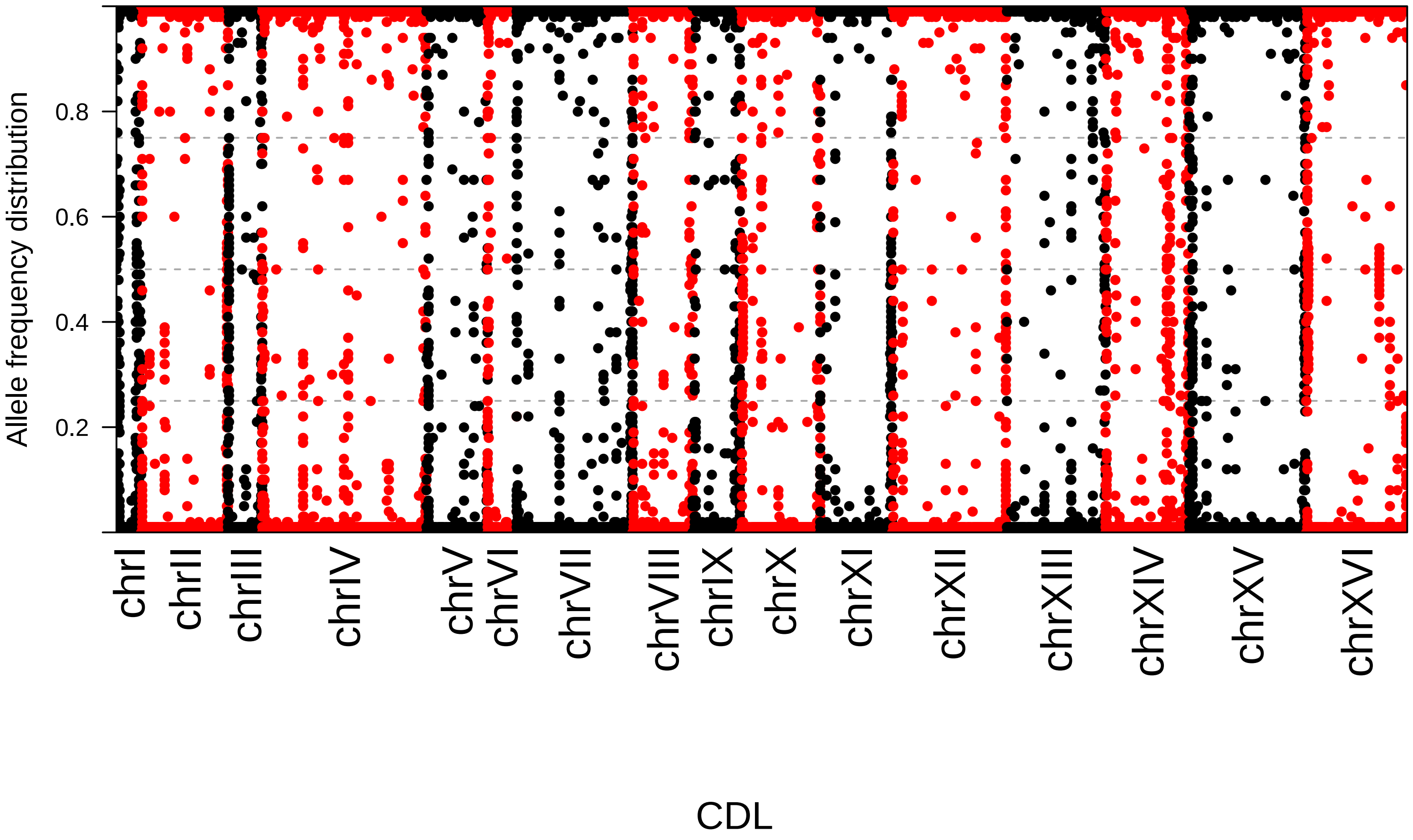

Allele frequency distribution
0.2
0.4
0.6
0.8
chrI
chrII
chrIII
chrIV
chrV
chrVI
chrVII
chrVIII
chrIX
chrX
chrXI
chrXII
chrXIII
chrXIV
chrXV
chrXVI
CDL

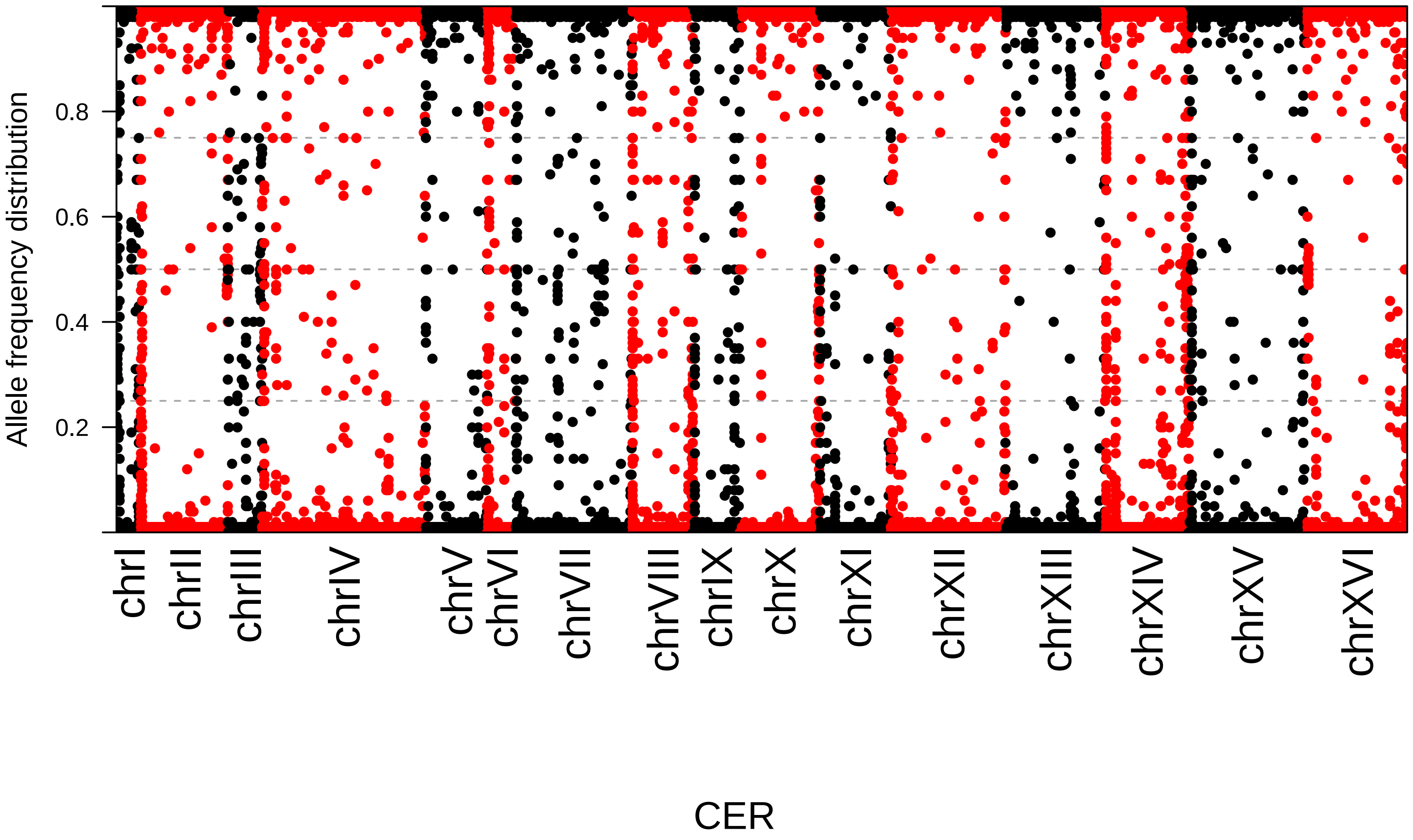

Allele frequency distribution
0.2
0.4
0.6
0.8
chrI
chrII
chrIII
chrIV
chrV
chrVI
chrVII
chrVIII
chrIX
chrX
chrXI
chrXII
chrXIII
chrXIV
chrXV
chrXVI
CER

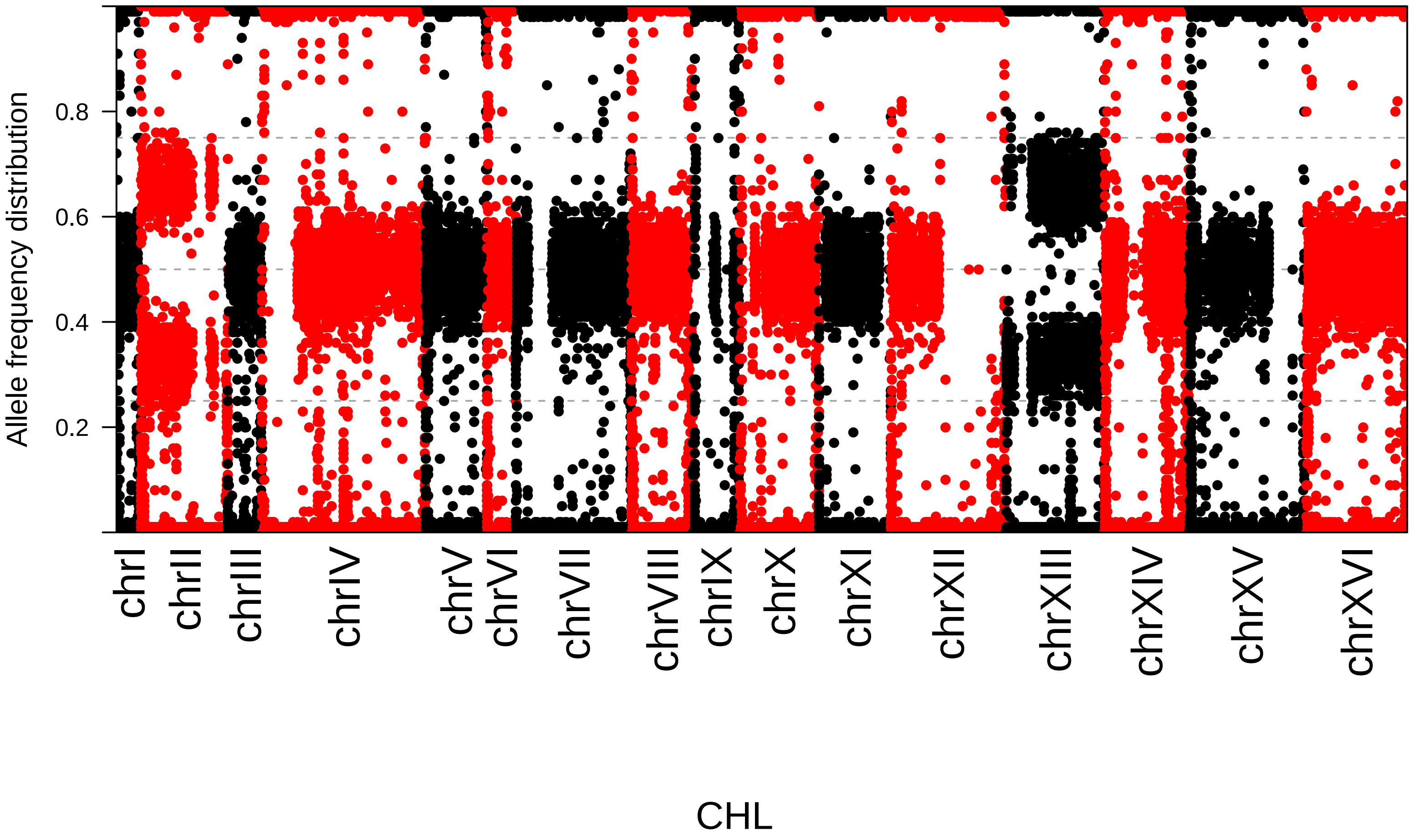
Allele frequency distribution
0.2
0.4
0.6
0.8
chrI
chrII
chrIII
chrIV
chrV
chrVI
chrVII
chrVIII
chrIX
chrX
chrXI
chrXII
chrXIII
chrXIV
chrXV
chrXVI
CHL

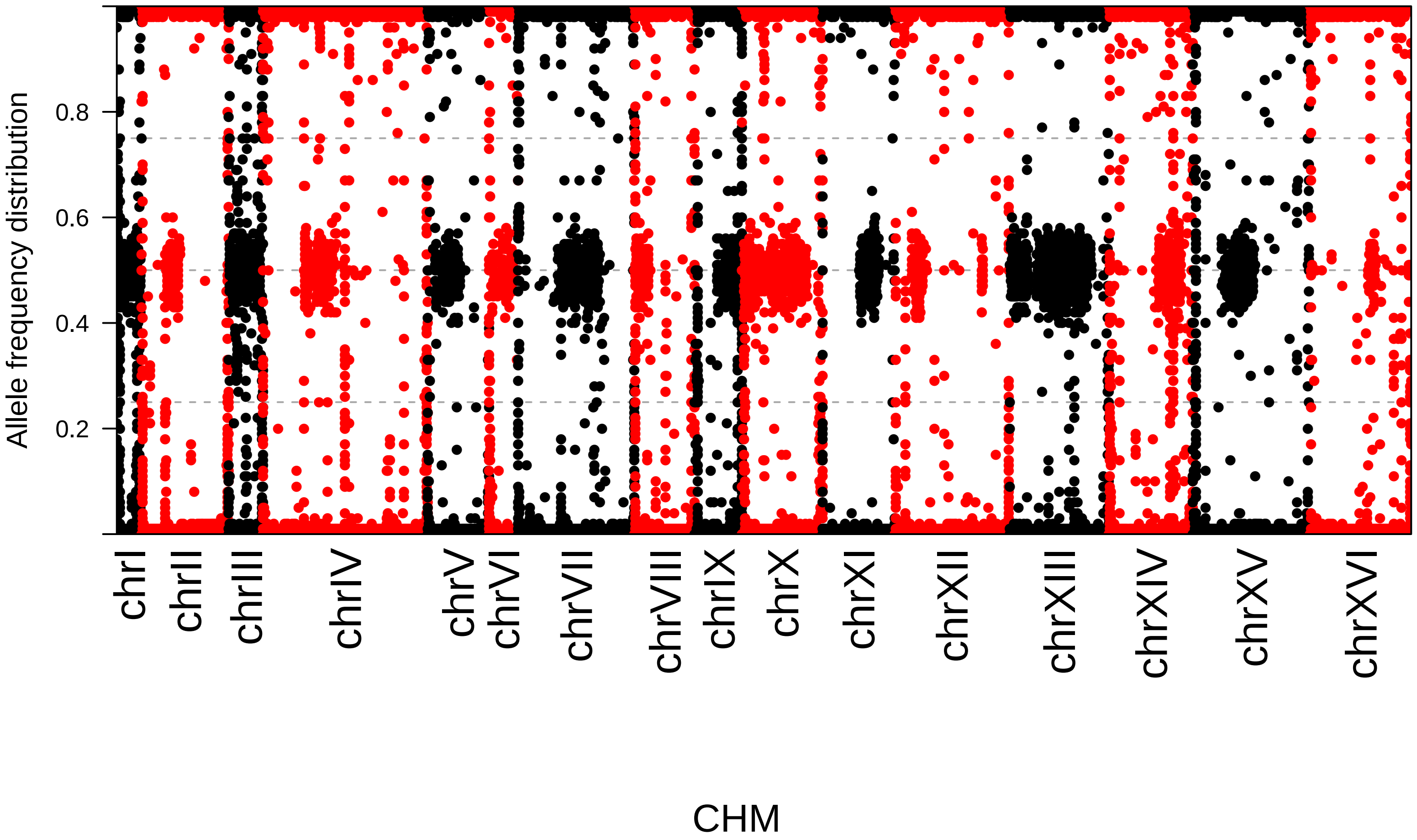

Allele frequency distribution
0.2
0.4
0.6
0.8
chrI
chrII
chrIII
chrIV
chrV
chrVI
chrVII
chrVIII
chrIX
chrX
chrXI
chrXII
chrXIII
chrXIV
chrXV
chrXVI
CHM

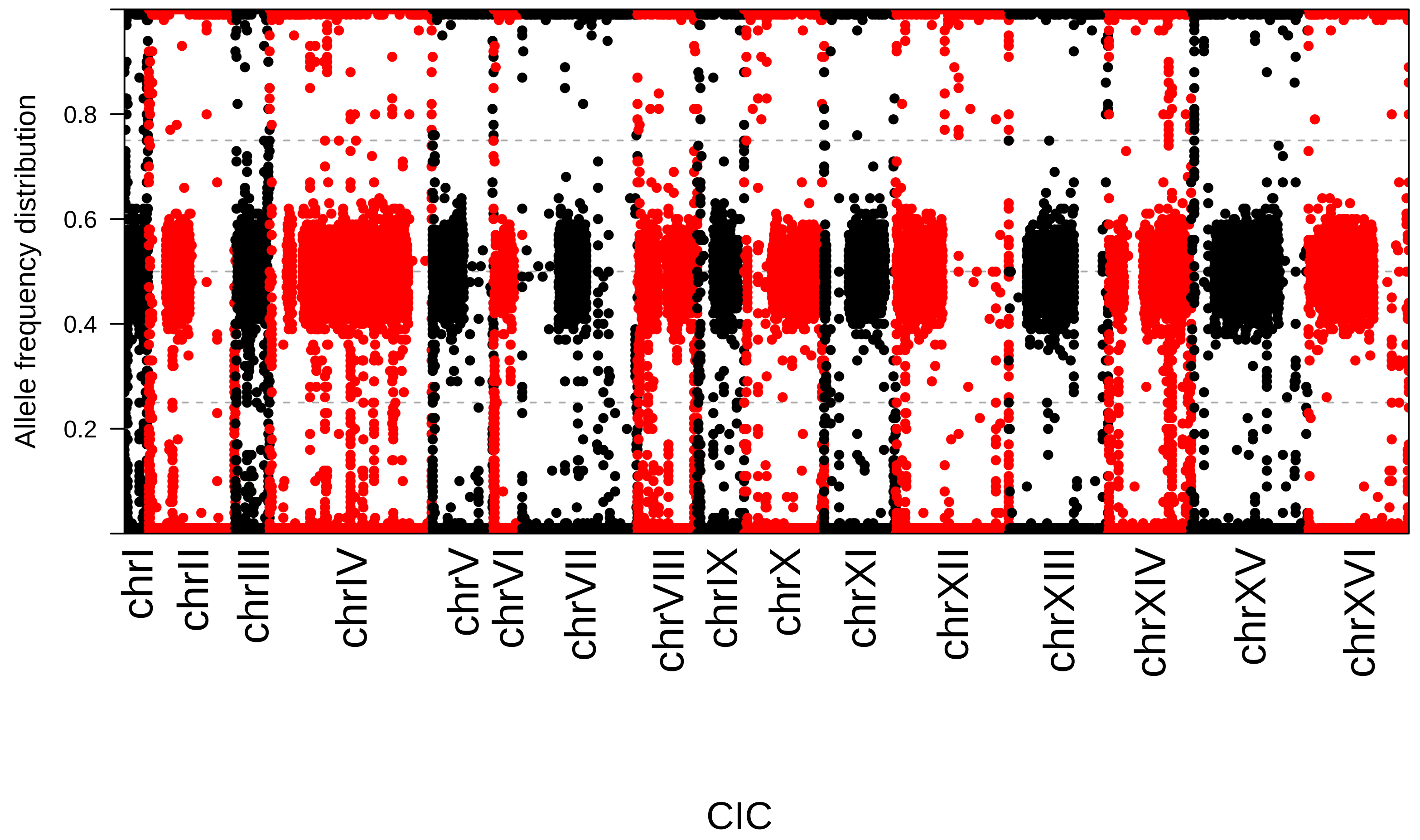

Allele frequency distribution
0.2
0.4
0.6
0.8
chrI
chrII
chrIII
chrIV
chrV
chrVI
chrVII
chrVIII
chrIX
chrX
chrXI
chrXII
chrXIII
chrXIV
chrXV
chrXVI
CIC

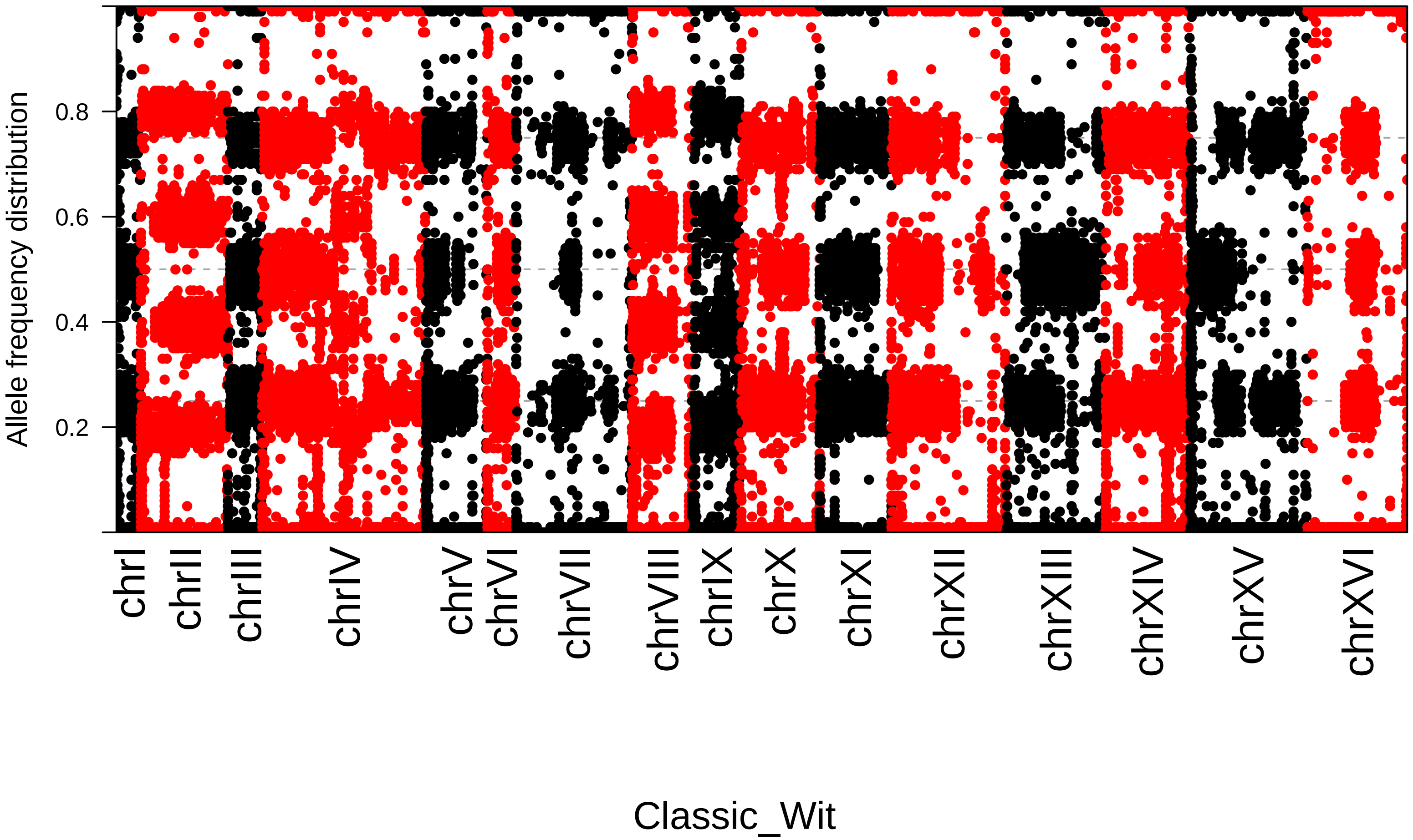

Allele frequency distribution
0.2
0.4
0.6
0.8
chrI
chrII
chrIII
chrIV
chrV
chrVI
chrVII
chrVIII
chrIX
chrX
chrXI
chrXII
chrXIII
chrXIV
chrXV
chrXVI
Classic_Wit

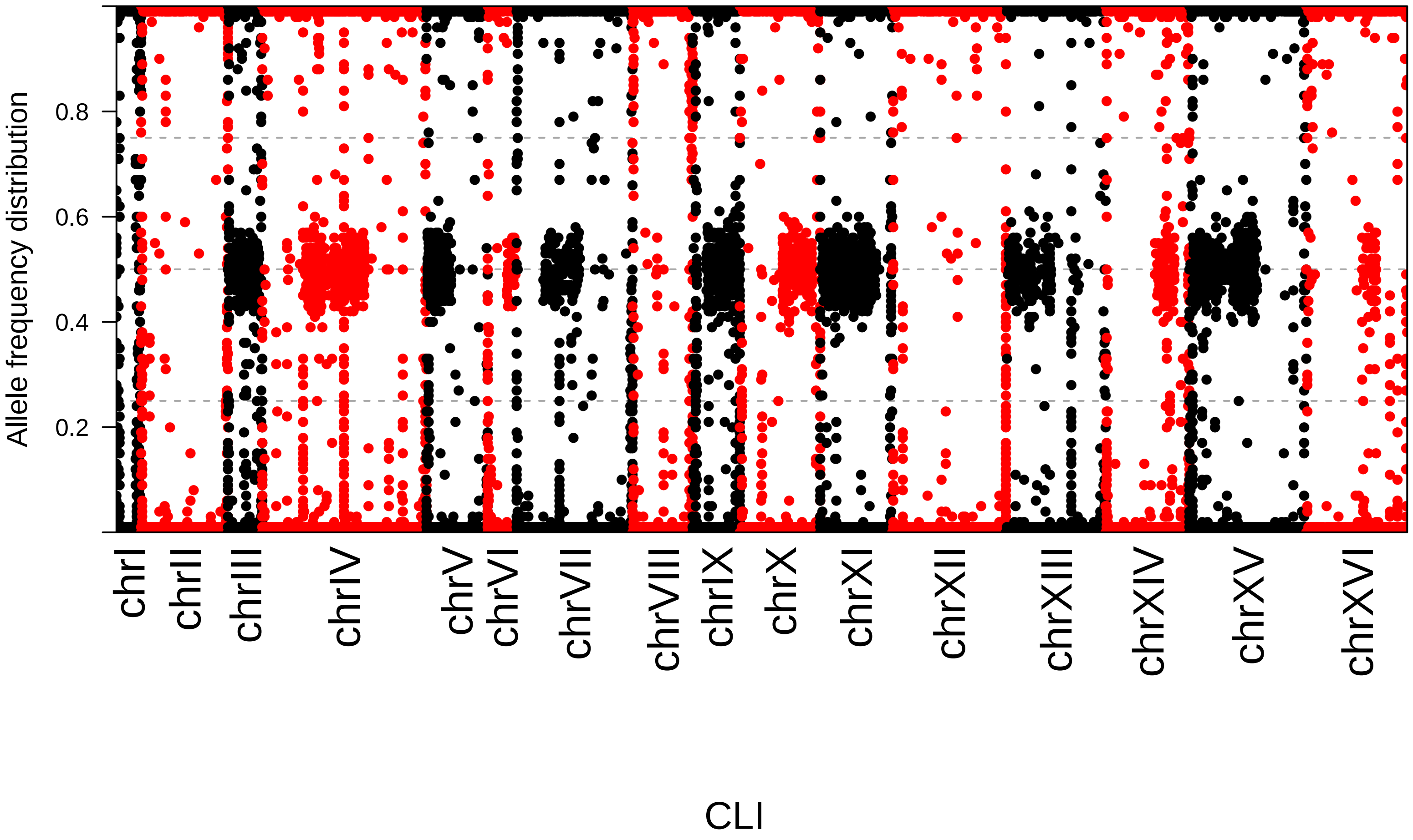

Allele frequency distribution
0.2
0.4
0.6
0.8
chrI
chrII
chrIII
chrIV
chrV
chrVI
chrVII
chrVIII
chrIX
chrX
chrXI
chrXII
chrXIII
chrXIV
chrXV
chrXVI
CLI

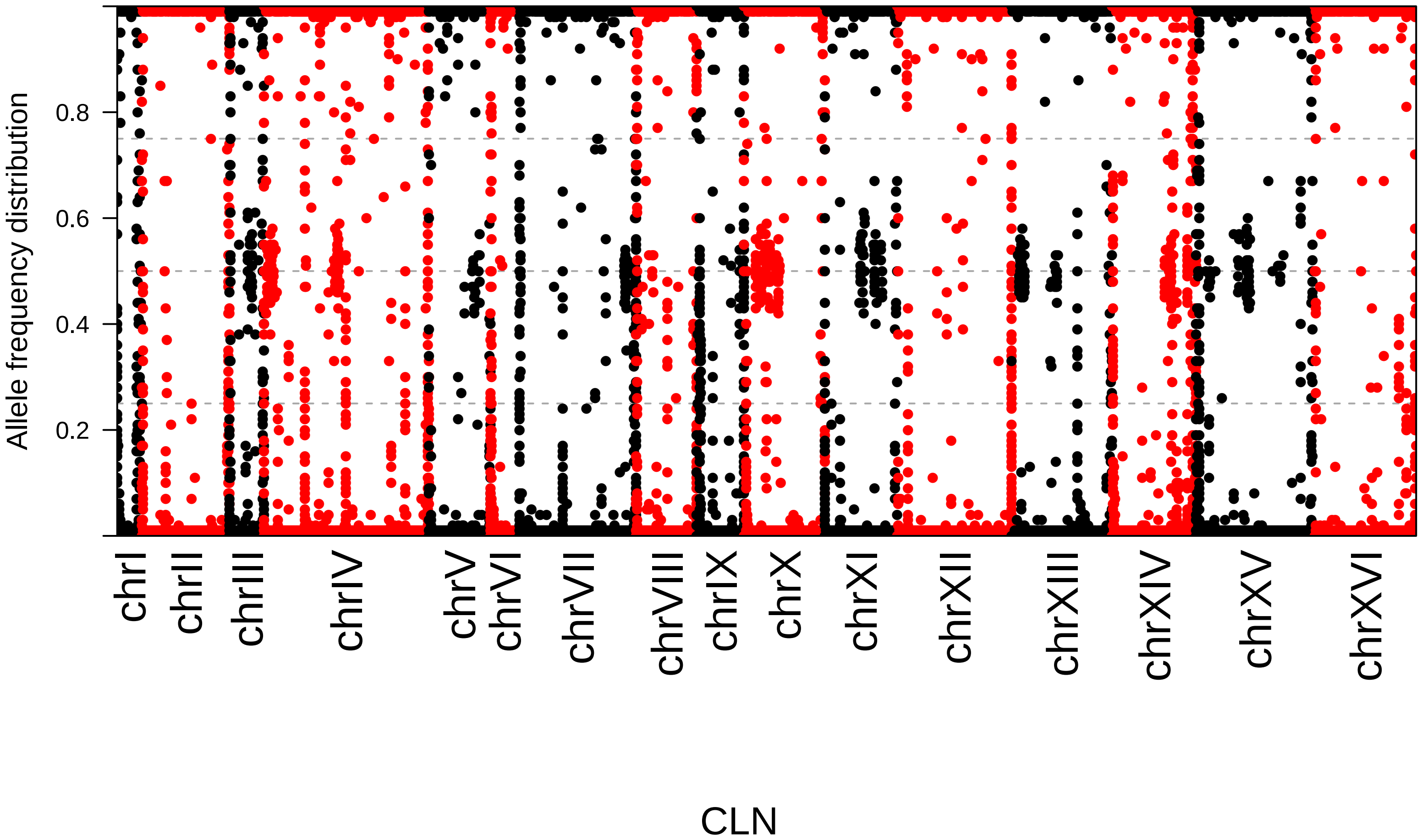

Allele frequency distribution
0.2
0.4
0.6
0.8
chrI
chrII
chrIII
chrIV
chrV
chrVI
chrVII
chrVIII
chrIX
chrX
chrXI
chrXII
chrXIII
chrXIV
chrXV
chrXVI
CLN

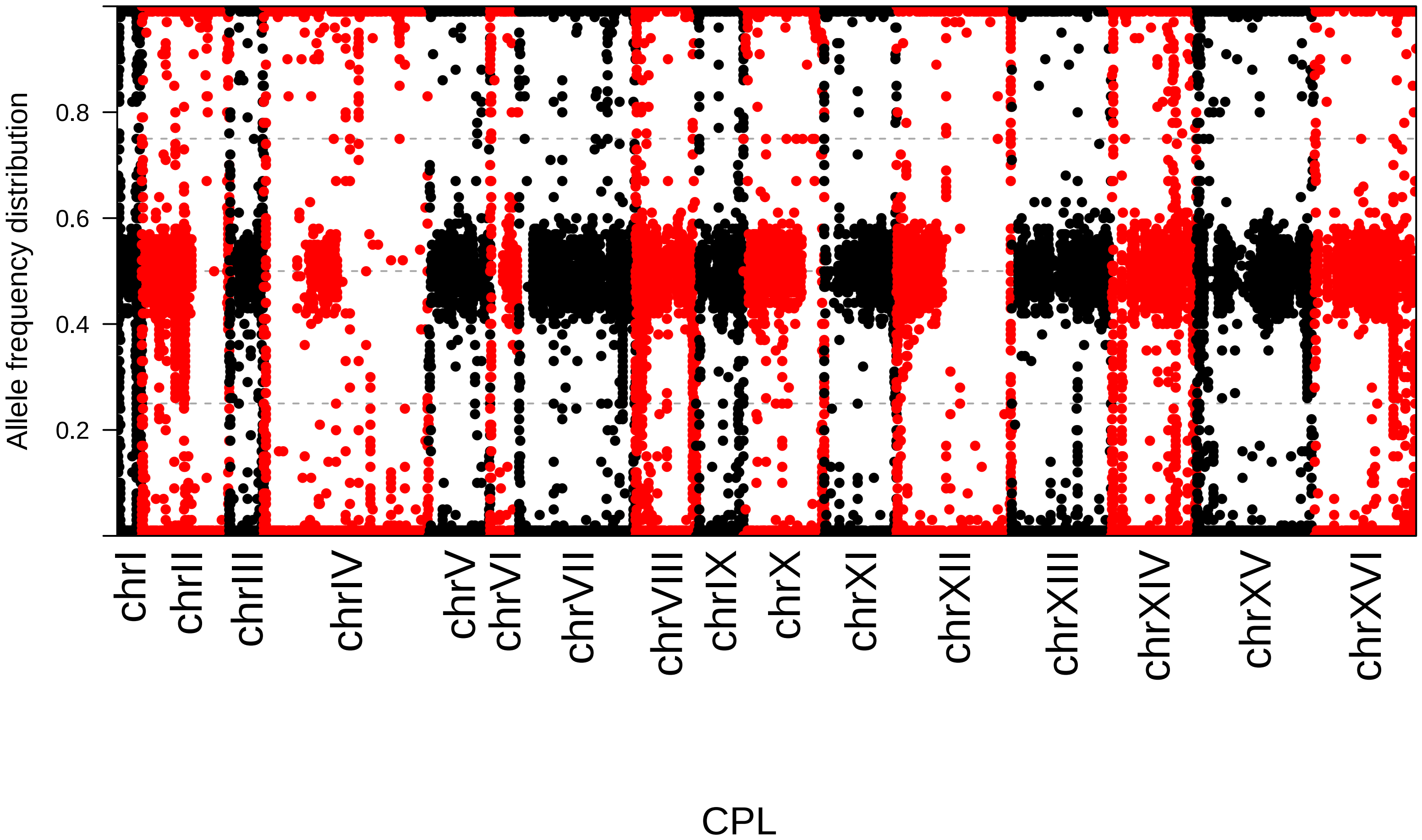

Allele frequency distribution
0.2
0.4
0.6
0.8
chrI
chrII
chrIII
chrIV
chrV
chrVI
chrVII
chrVIII
chrIX
chrX
chrXI
chrXII
chrXIII
chrXIV
chrXV
chrXVI
CPL

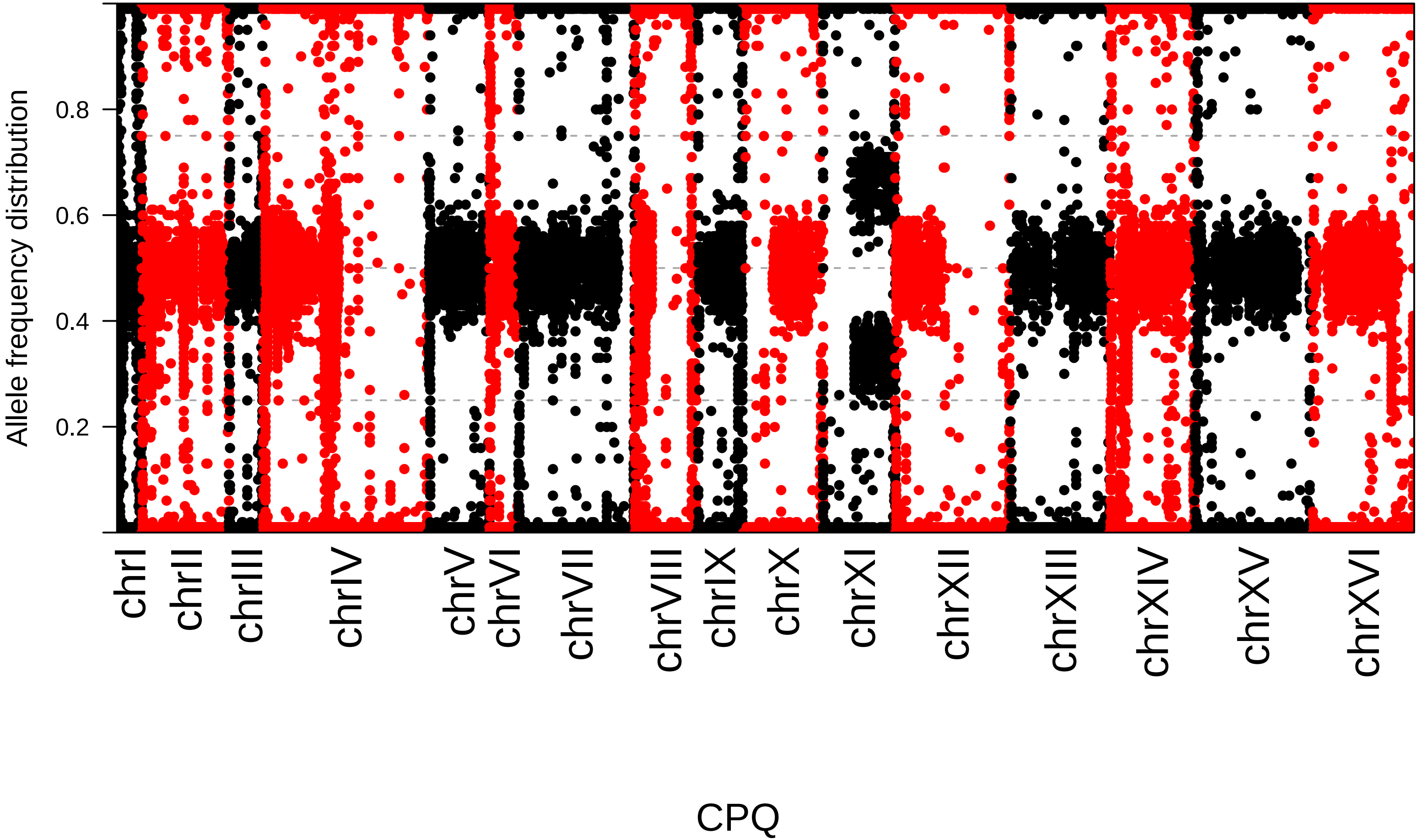
Allele frequency distribution
0.2
0.4
0.6
0.8
chrI
chrII
chrIII
chrIV
chrV
chrVI
chrVII
chrVIII
chrIX
chrX
chrXI
chrXII
chrXIII
chrXIV
chrXV
chrXVI
CPQ

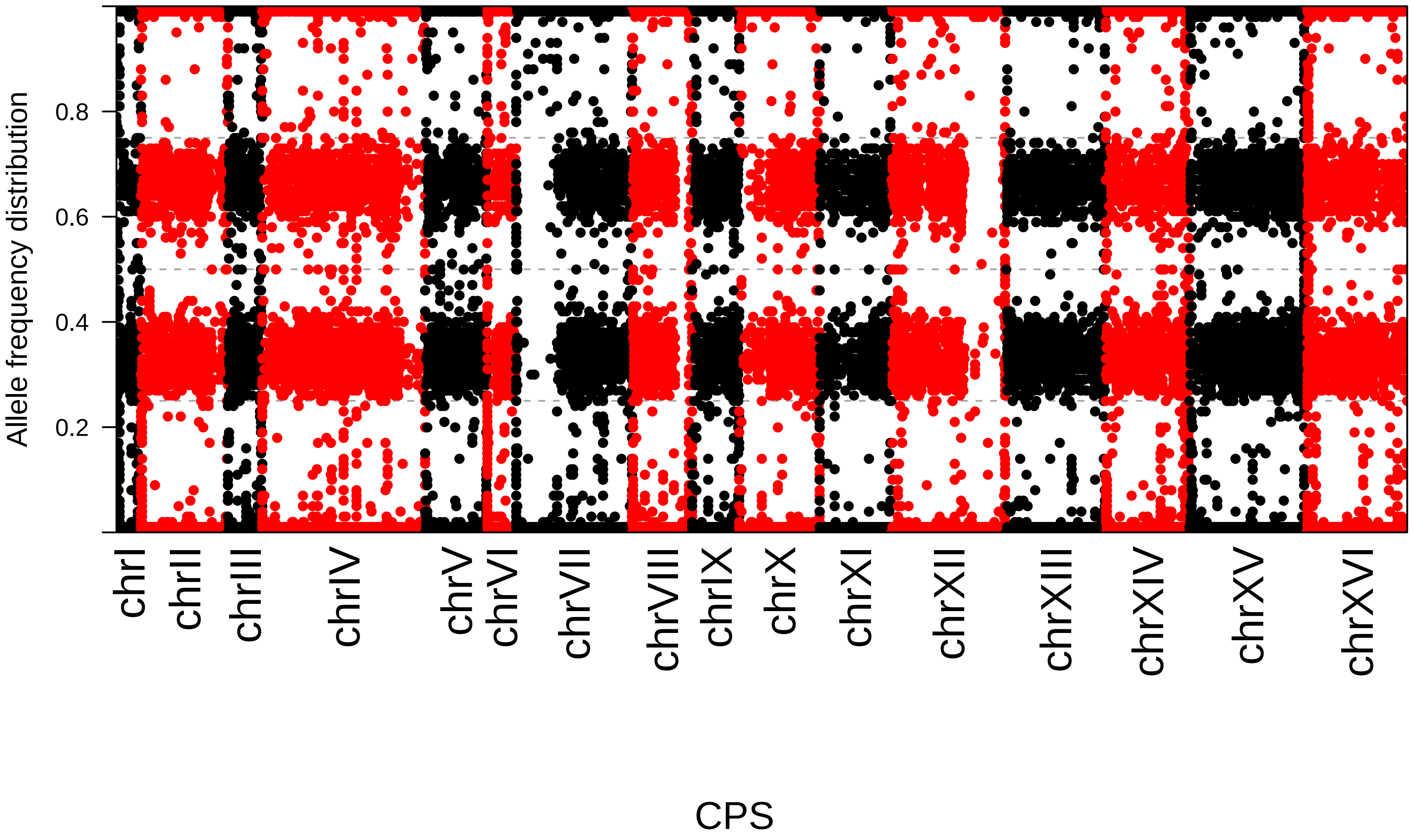
Allele frequency distribution
0.2
0.4
0.6
0.8
chrI
chrII
chrIII
chrIV
chrV
chrVI
chrVII
chrVIII
chrIX
chrX
chrXI
chrXII
chrXIII
chrXIV
chrXV
chrXVI
CPS

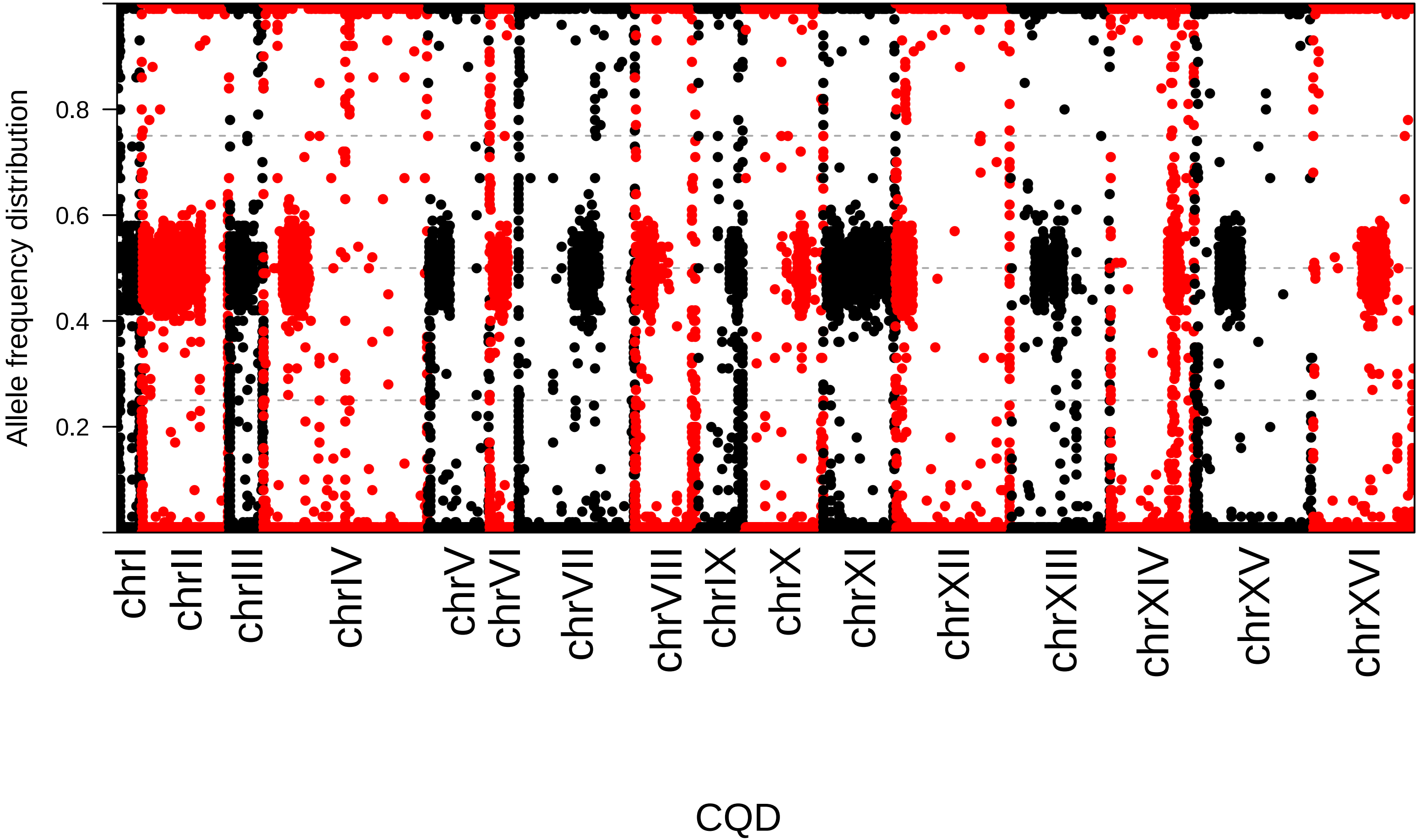
Allele frequency distribution
0.2
0.4
0.6
0.8
chrI
chrII
chrIII
chrIV
chrV
chrVI
chrVII
chrVIII
chrIX
chrX
chrXI
chrXII
chrXIII
chrXIV
chrXV
chrXVI
CQD

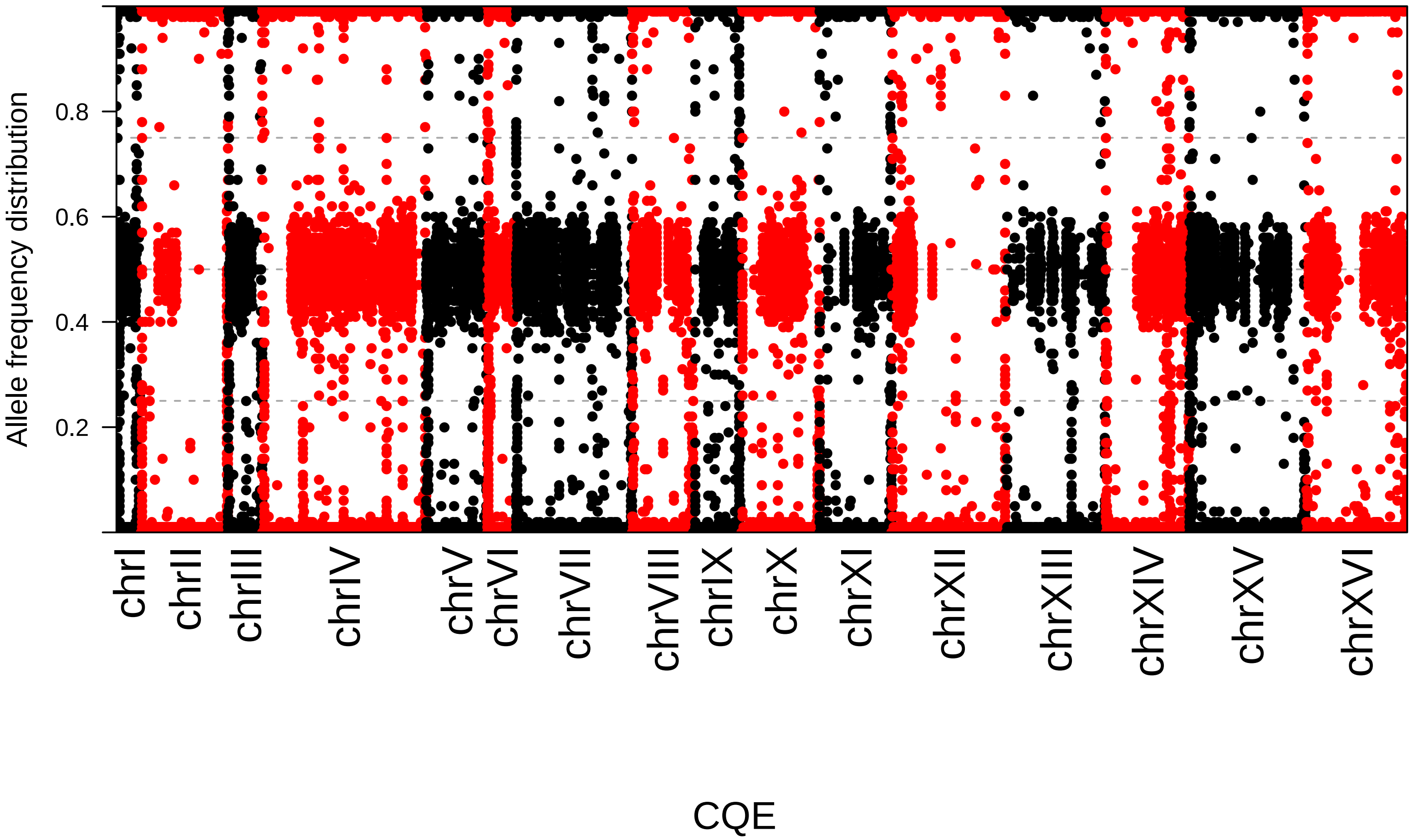

Allele frequency distribution
0.2
0.4
0.6
0.8
chrI
chrII
chrIII
chrIV
chrV
chrVI
chrVII
chrVIII
chrIX
chrX
chrXI
chrXII
chrXIII
chrXIV
chrXV
chrXVI
CQE

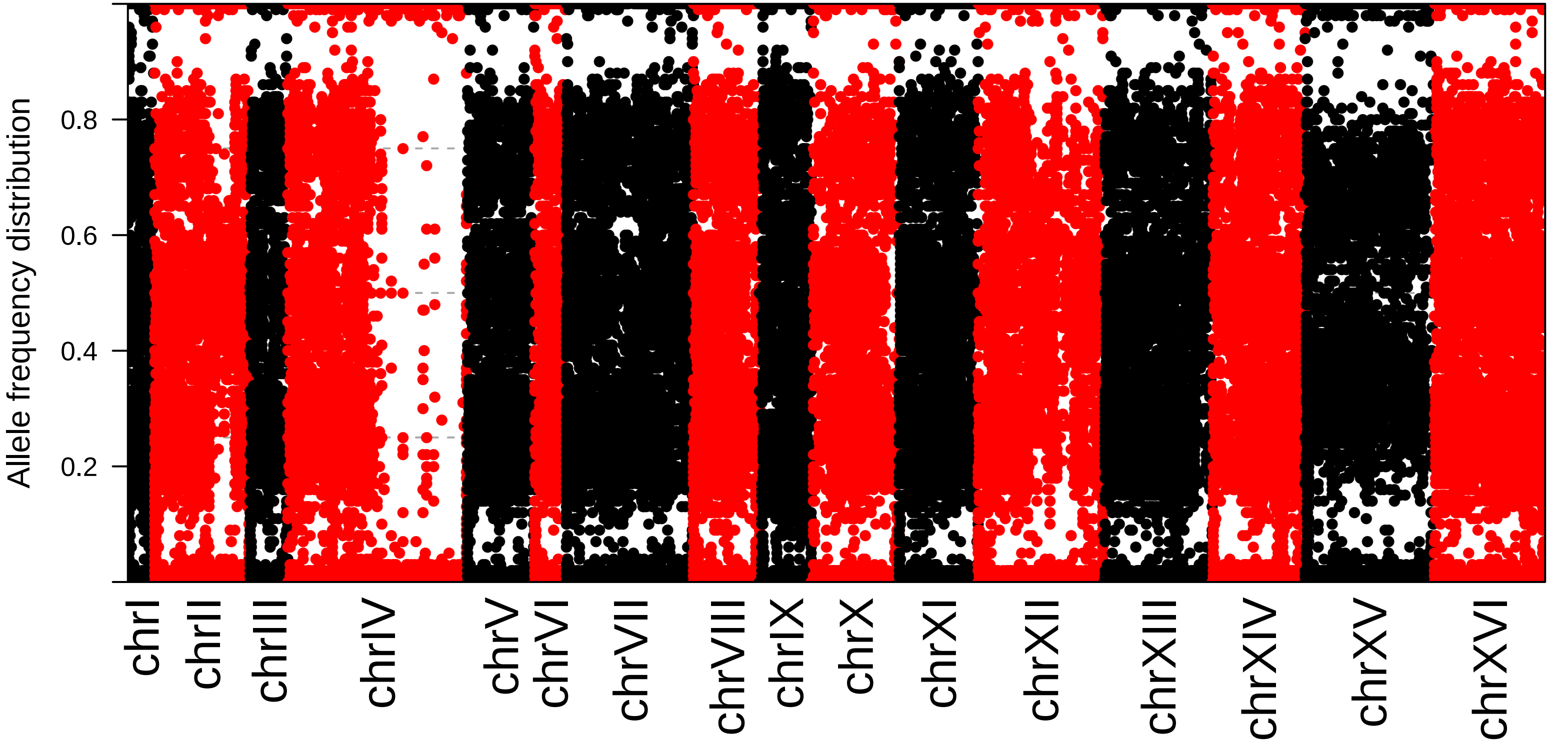
Allele frequency distribution
0.2
0.4
0.6
0.8
chrI
chrII
chrIII
chrIV
chrV
chrVI
chrVII
chrVIII
chrIX
chrX
chrXI
chrXII
chrXIII
chrXIV
chrXV
chrXVI

Djonno1

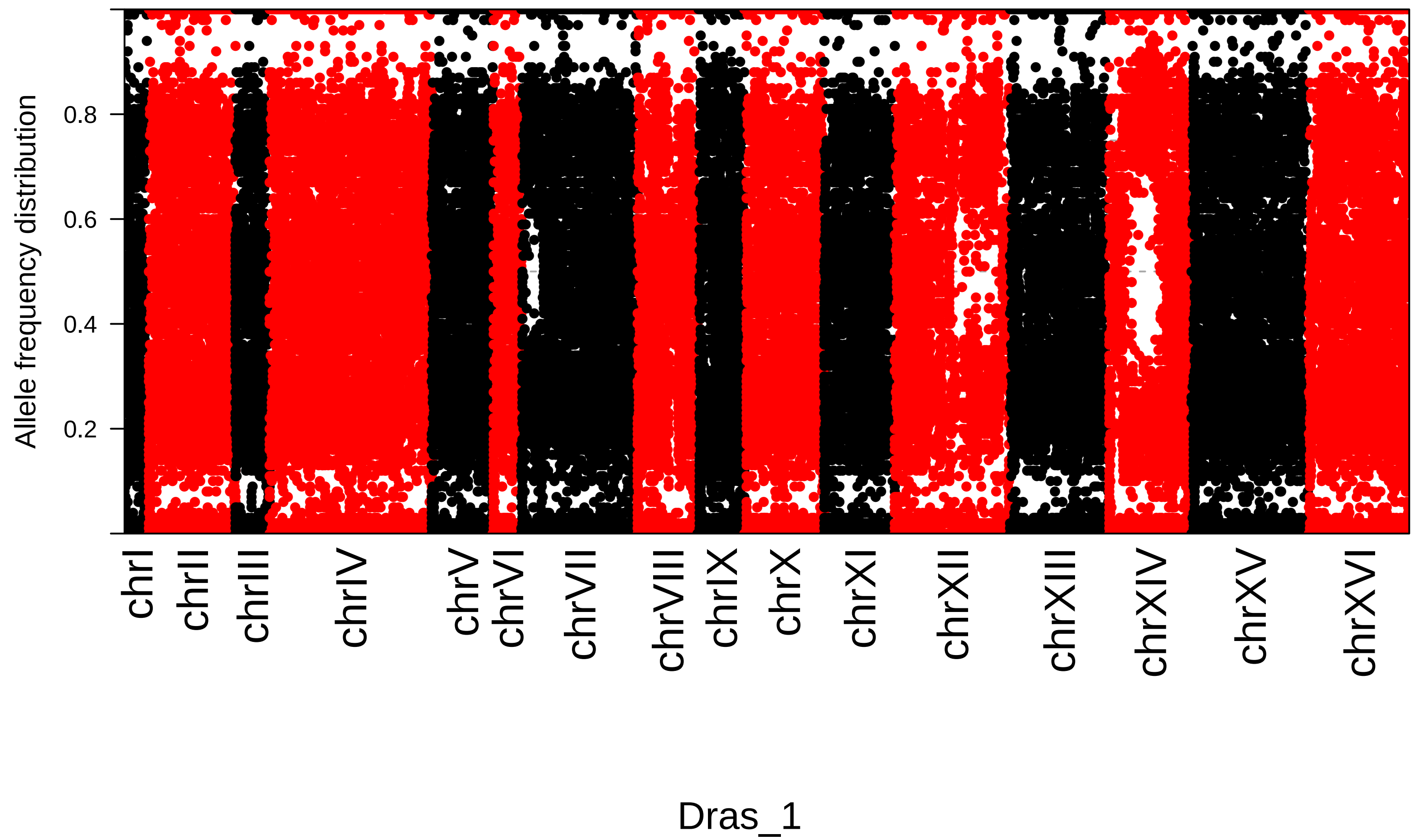

Allele frequency distribution
0.8
0.6
0.4
0.2
chrI
chrII
chrIII
chrIV
chrV
chrVI
chrVII
chrVIII
chrIX
chrX
chrXI
chrXII
chrXIII
chrXIV
chrXV
chrXVI
Dras_1

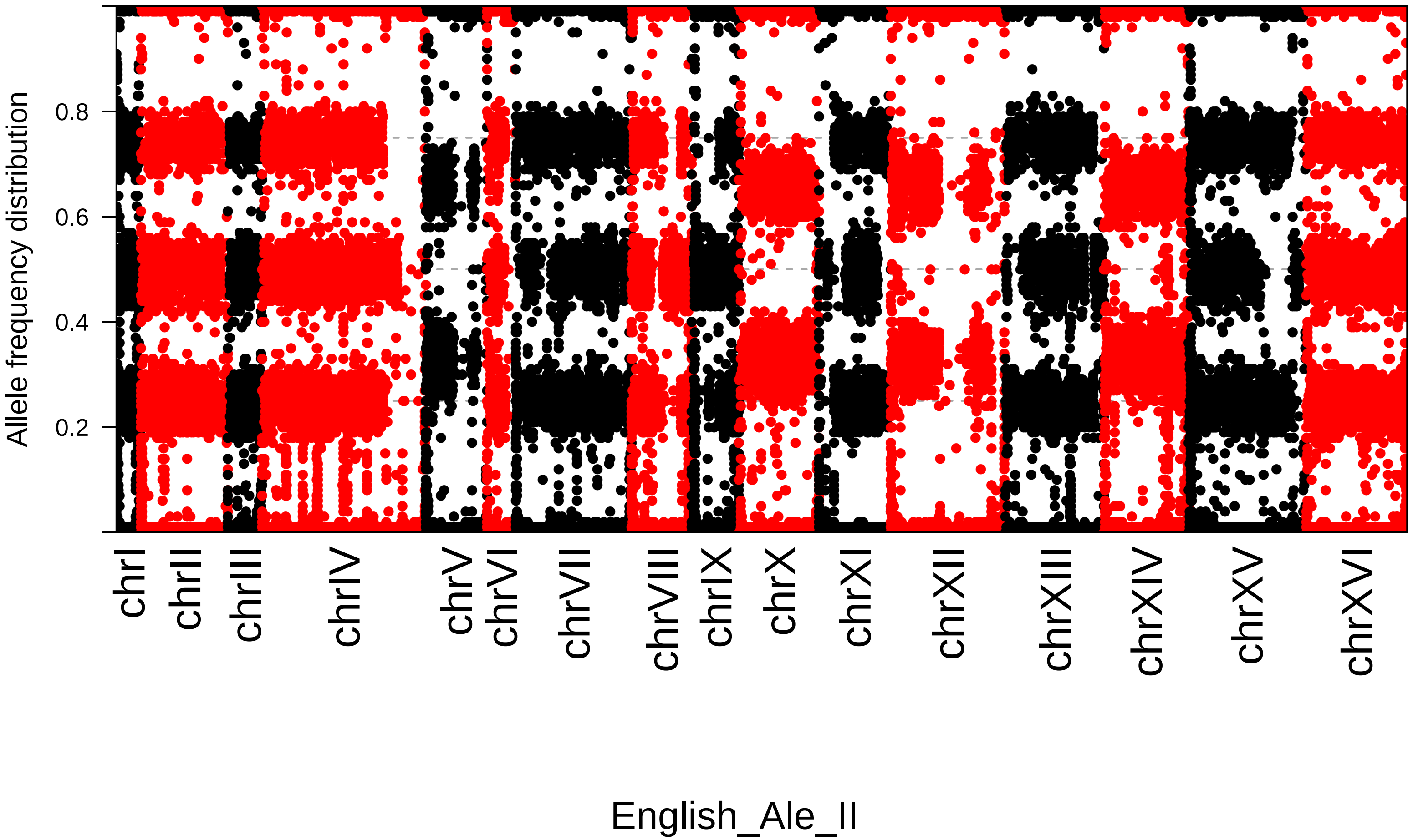

Allele frequency distribution
0.2
0.4
0.6
0.8
chrI
chrII
chrIII
chrIV
chrV
chrVI
chrVII
chrVIII
chrIX
chrX
chrXI
chrXII
chrXIII
chrXIV
chrXV
chrXVI
English_Ale_II

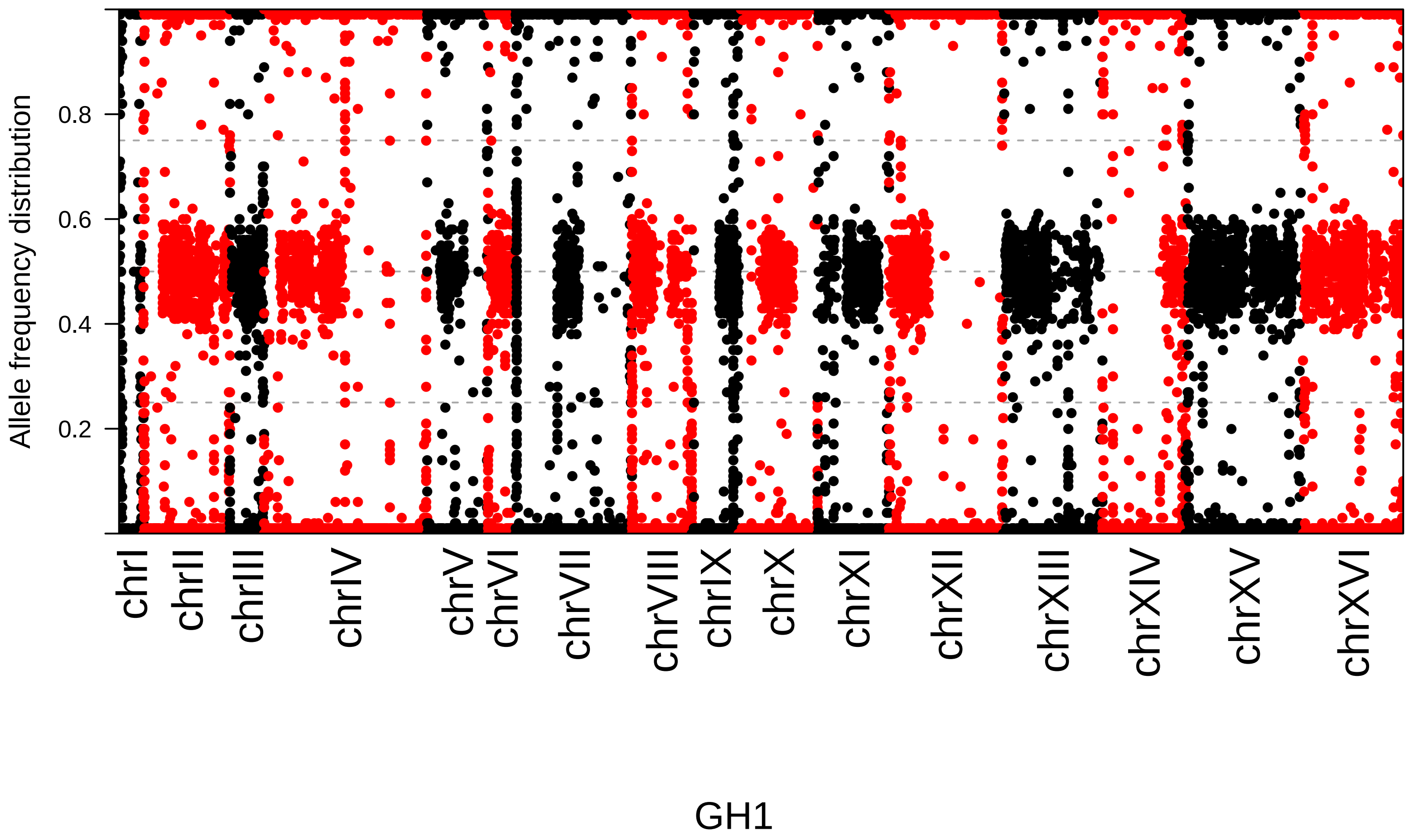

Allele frequency distribution
0.2
0.4
0.6
0.8
chrI
chrII
chrIII
chrIV
chrV
chrVI
chrVII
chrVIII
chrIX
chrX
chrXI
chrXII
chrXIII
chrXIV
chrXV
chrXVI
GH1

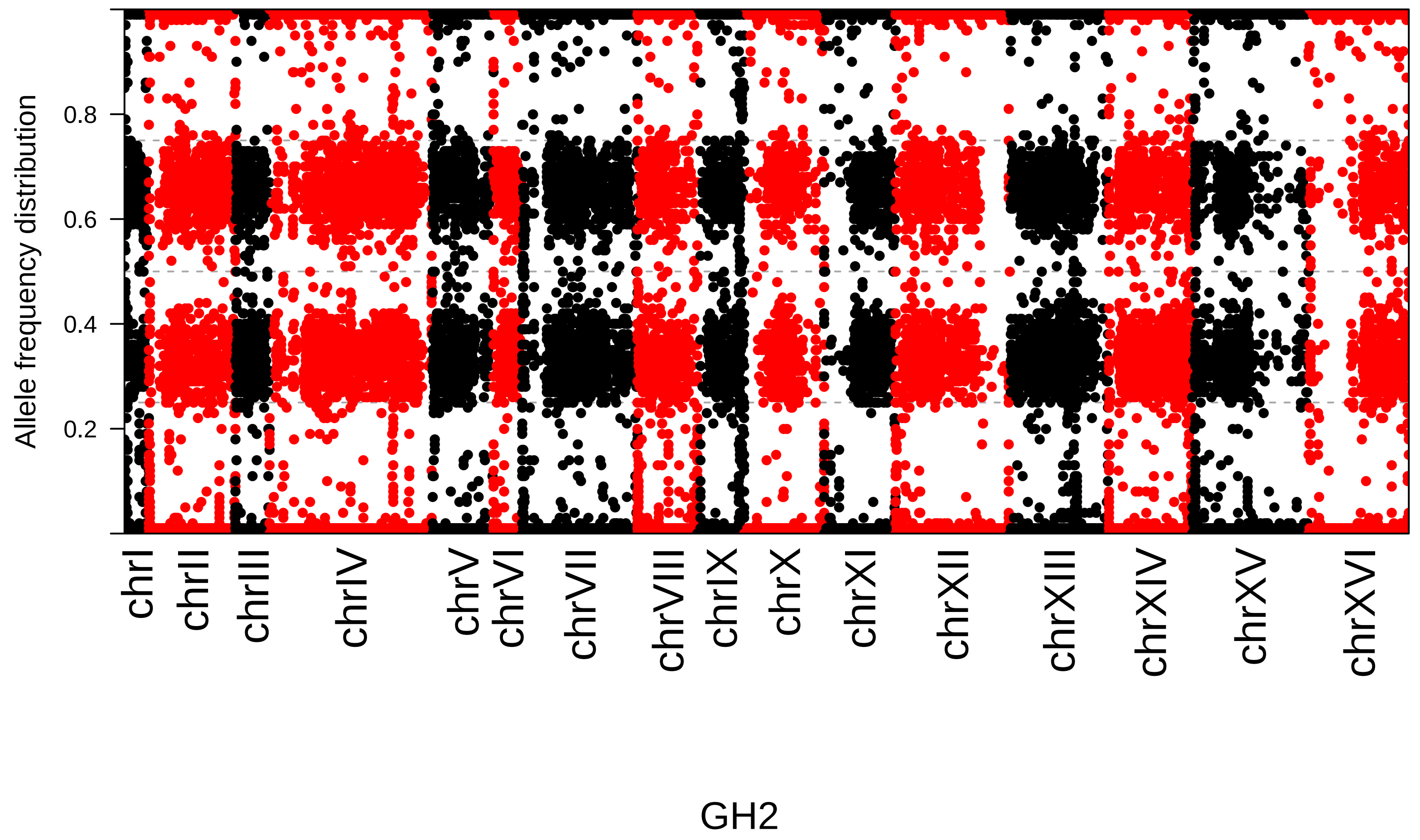

Allele frequency distribution
0.8
0.6
0.4
0.2
chrI
chrII
chrIII
chrIV
chrV
chrVI
chrVII
chrVIII
chrIX
chrX
chrXI
chrXII
chrXIII
chrXIV
chrXV
chrXVI
GH2

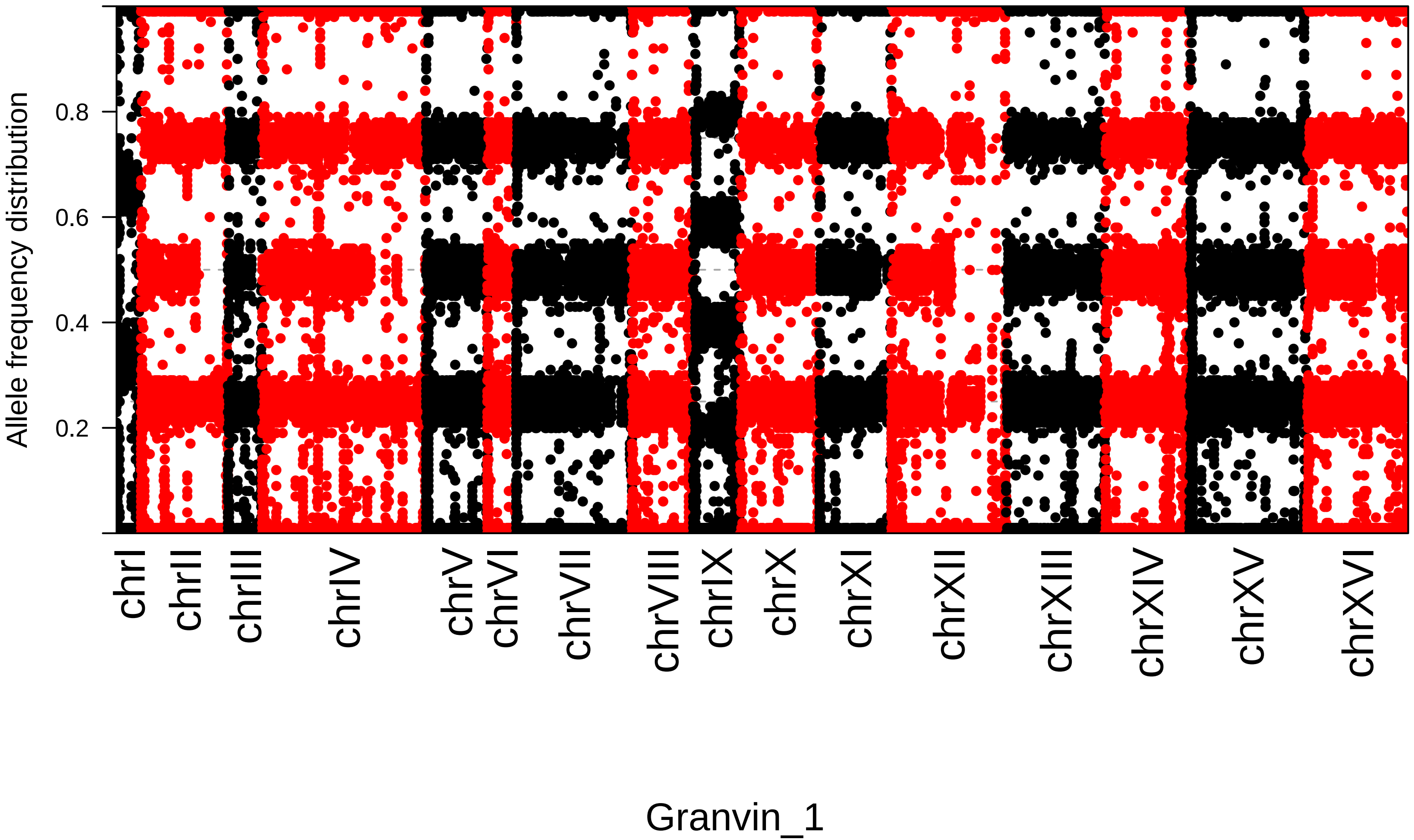
Allele frequency distribution
0.2
0.4
0.6
0.8
chrI
chrII
chrIII
chrIV
chrV
chrVI
chrVII
chrVIII
chrIX
chrX
chrXI
chrXII
chrXIII
chrXIV
chrXV
chrXVI
Granvin_1

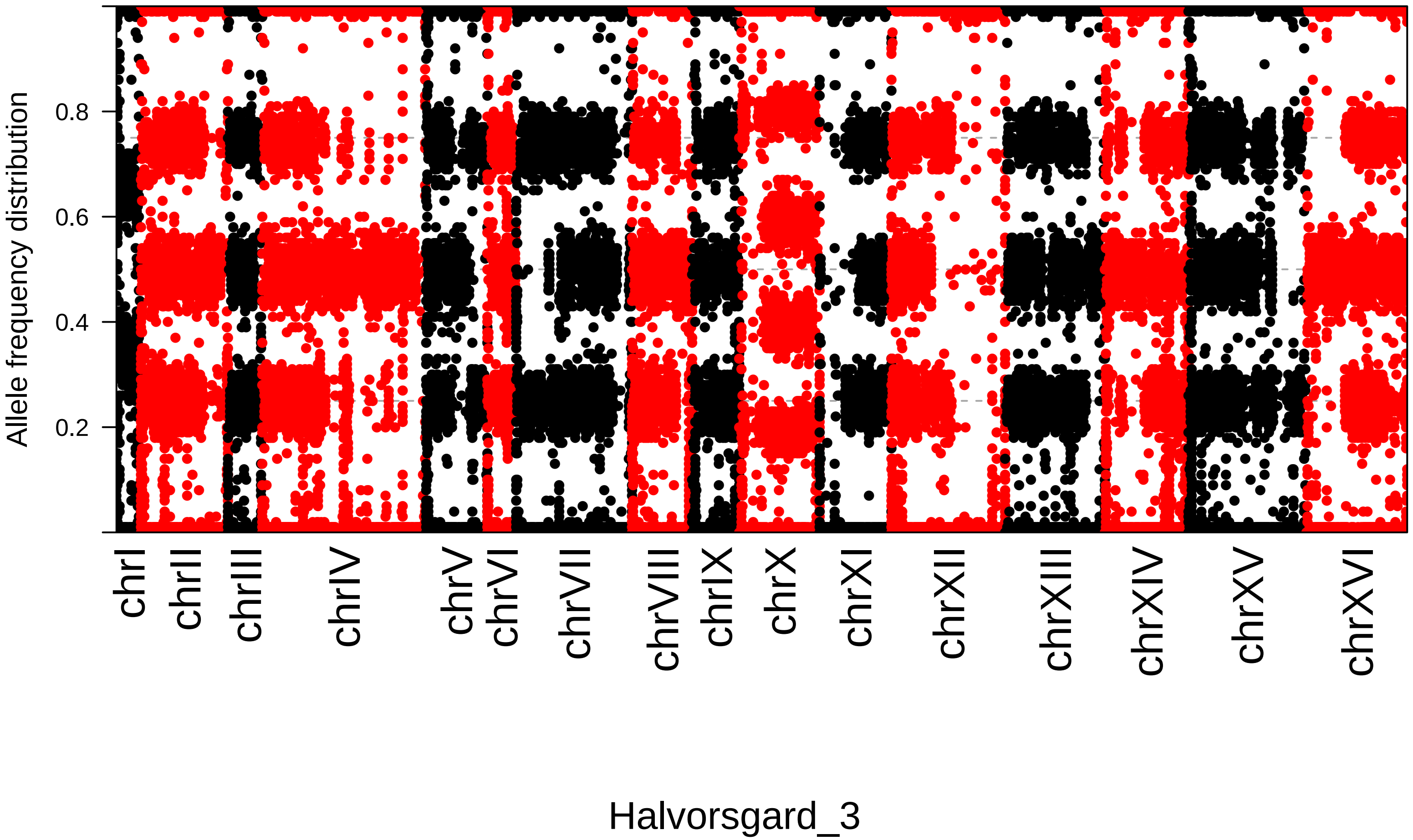

Allele frequency distribution
0.2
0.4
0.6
0.8
chrI
chrII
chrIII
chrIV
chrV
chrVI
chrVII
chrVIII
chrIX
chrX
chrXI
chrXII
chrXIII
chrXIV
chrXV
chrXVI
Halvorsgard_3

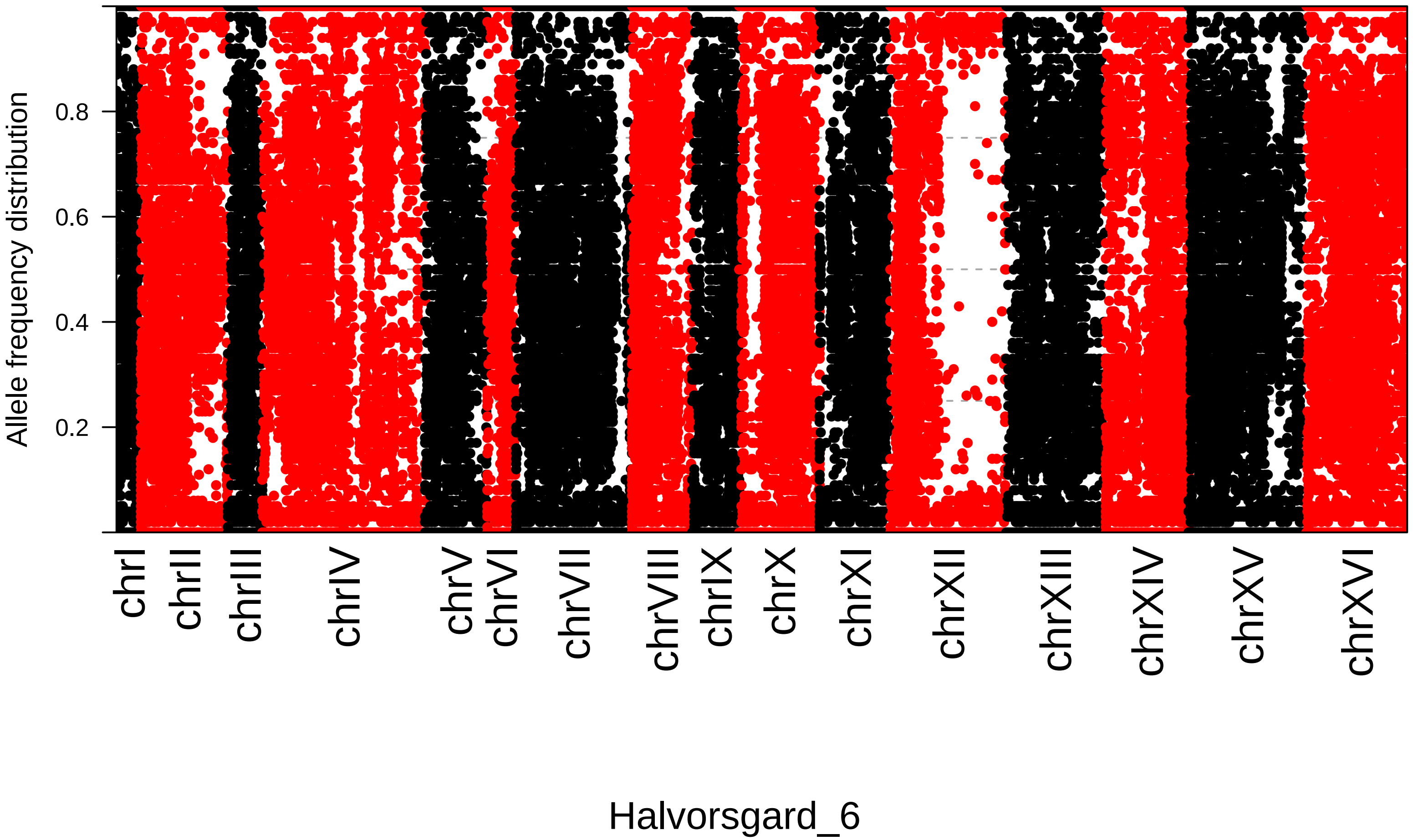

Allele frequency distribution
0.2
0.4
0.6
0.8
chrI
chrII
chrIII
chrIV
chrV
chrVI
chrVII
chrVIII
chrIX
chrX
chrXI
chrXII
chrXIII
chrXIV
chrXV
chrXVI
Halvorsgard_6

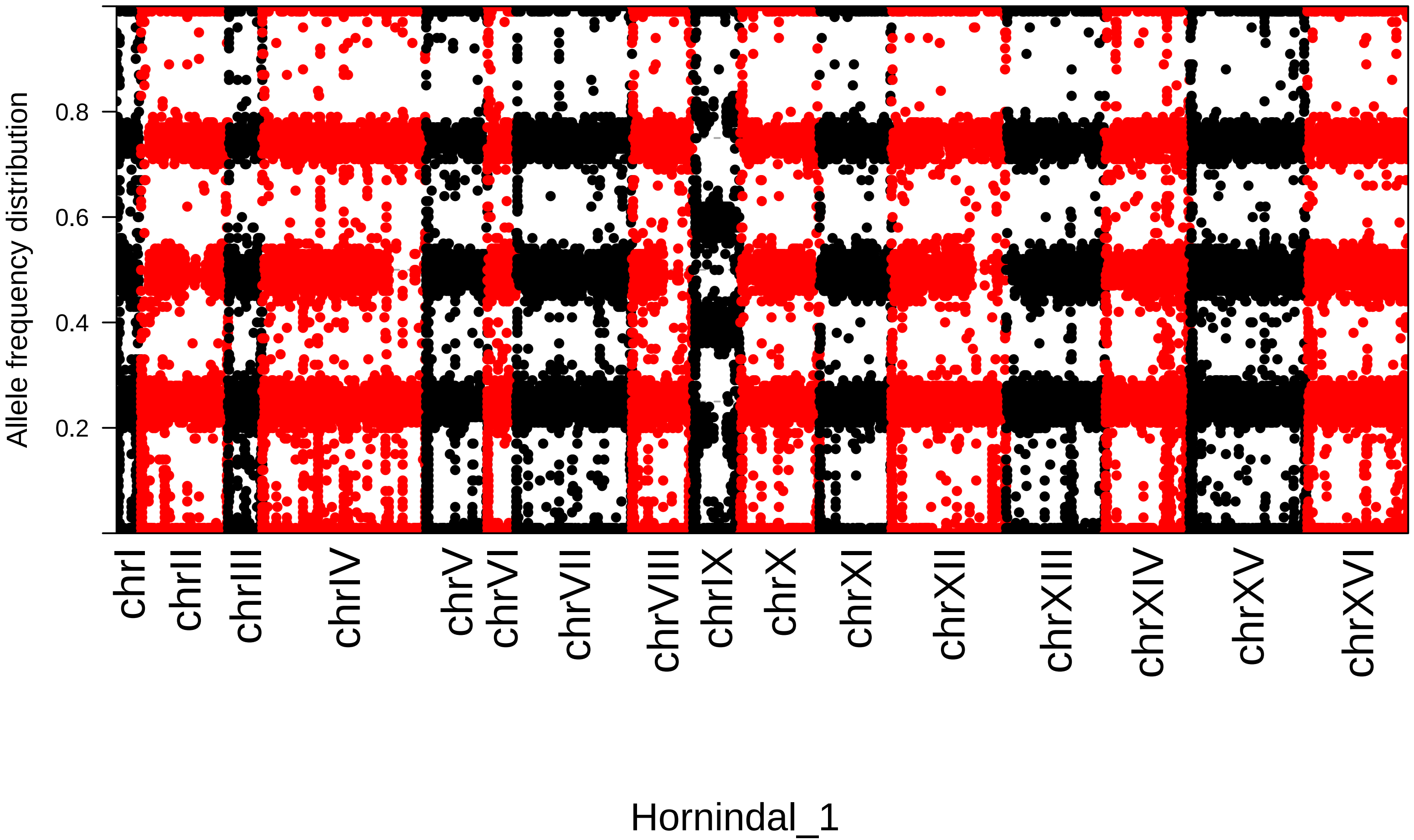

Allele frequency distribution
0.2
0.4
0.6
0.8
chrI
chrII
chrIII
chrIV
chrV
chrVI
chrVII
chrVIII
chrIX
chrX
chrXI
chrXII
chrXIII
chrXIV
chrXV
chrXVI
Hornindal_1

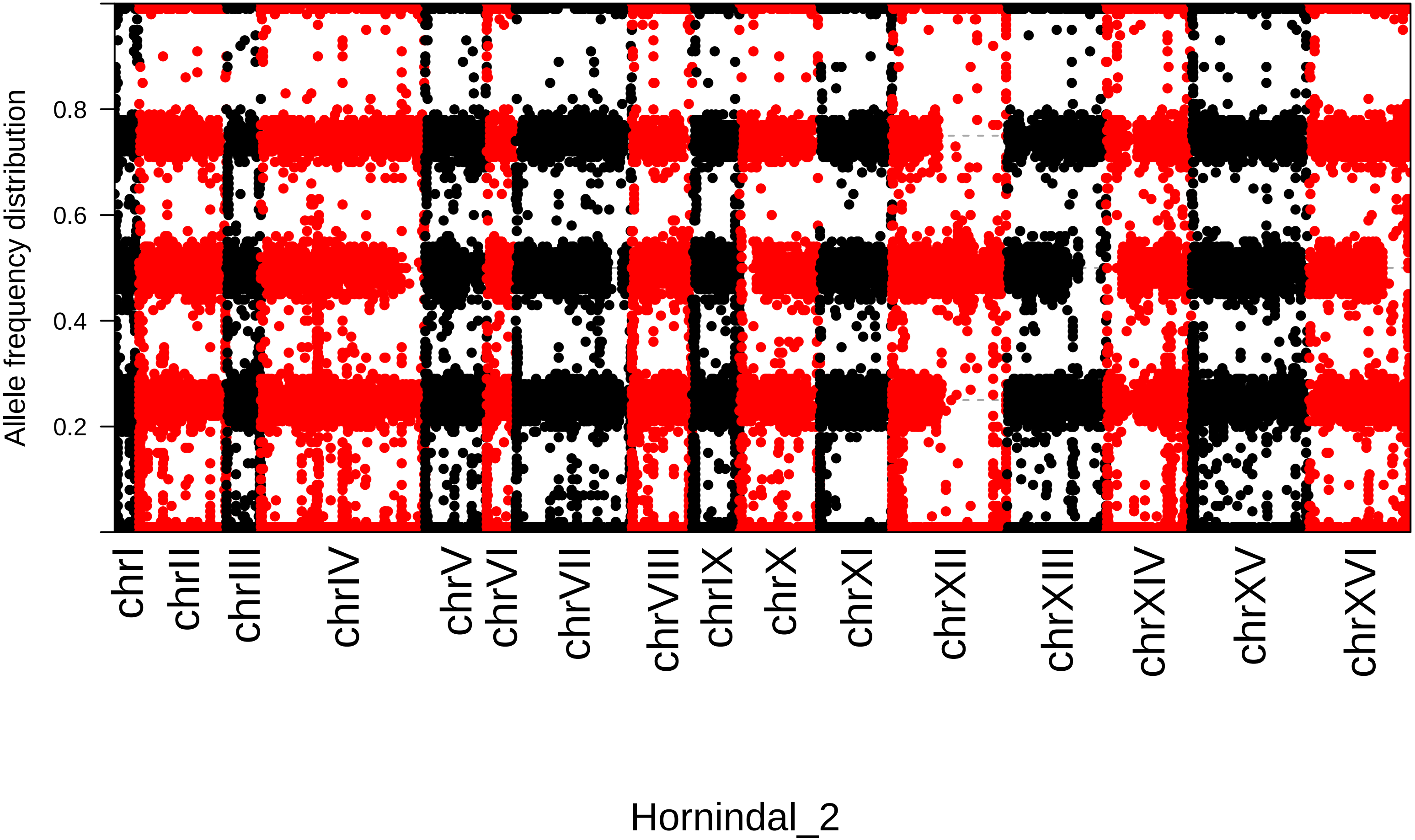

Allele frequency distribution
0.2
0.4
0.6
0.8
chrI
chrII
chrIII
chrIV
chrV
chrVI
chrVII
chrVIII
chrIX
chrX
chrXI
chrXII
chrXIII
chrXIV
chrXV
chrXVI
Hornindal_2

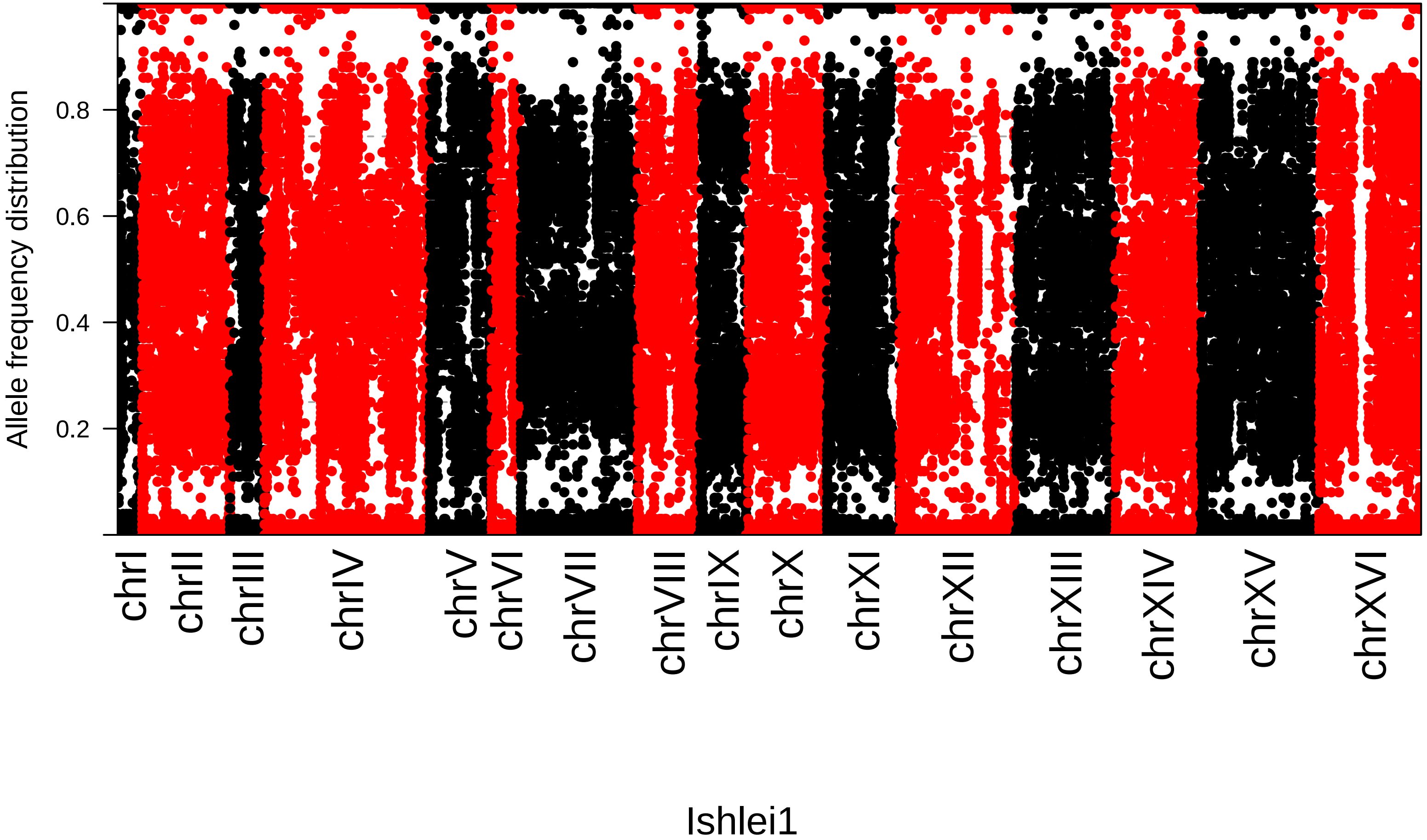
Allele frequency distribution
0.8
0.6
0.4
0.2
chrI
chrII
chrIII
chrIV
chrV
chrVI
chrVII
chrVIII
chrIX
chrX
chrXI
chrXII
chrXIII
chrXIV
chrXV
chrXVI
Ishlei1

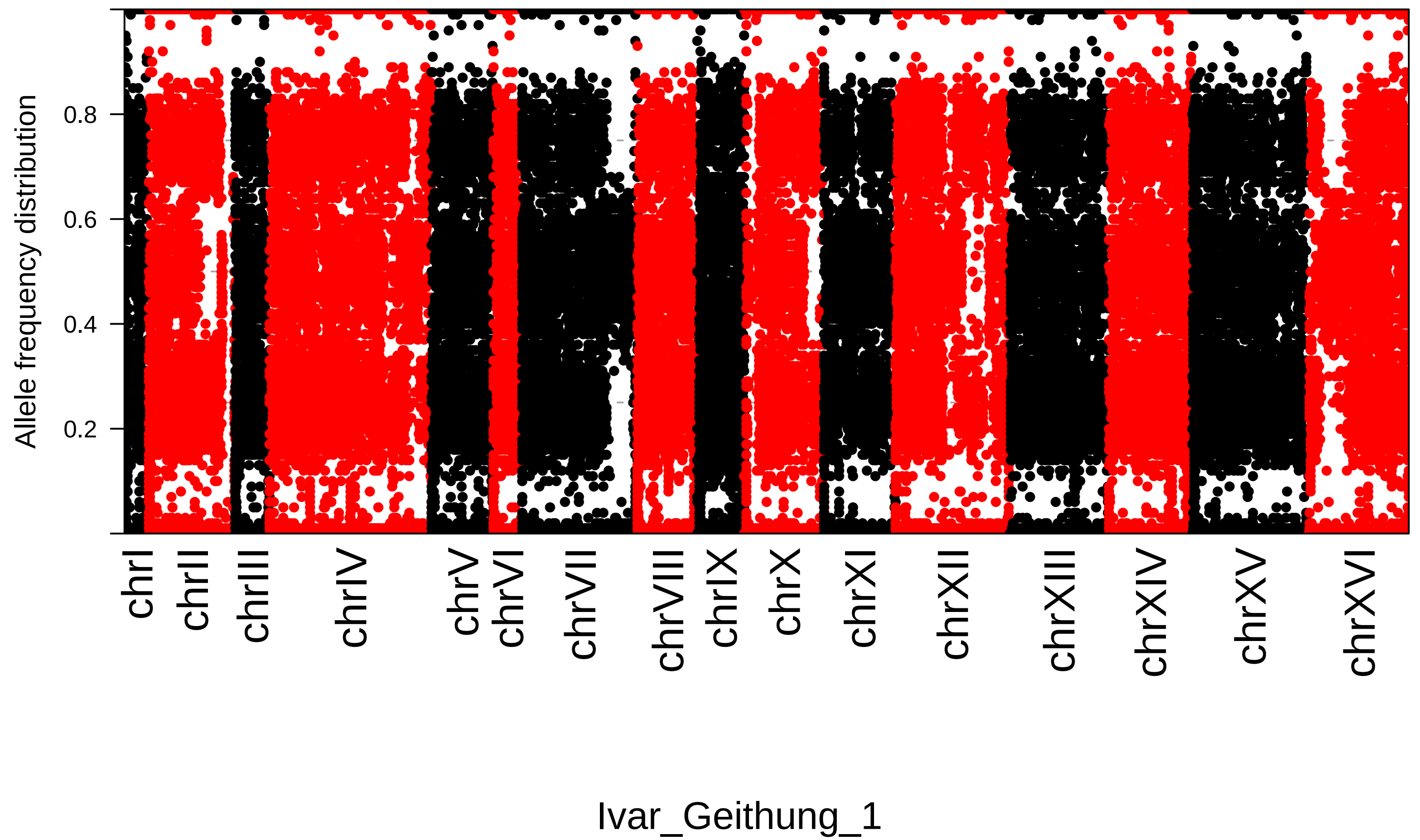
Allele frequency distribution
0.2
0.4
0.6
0.8
chrI
chrII
chrIII
chrIV
chrV
chrVI
chrVII
chrVIII
chrIX
chrX
chrXI
chrXII
chrXIII
chrXIV
chrXV
chrXVI
Ivar_Geithung_1

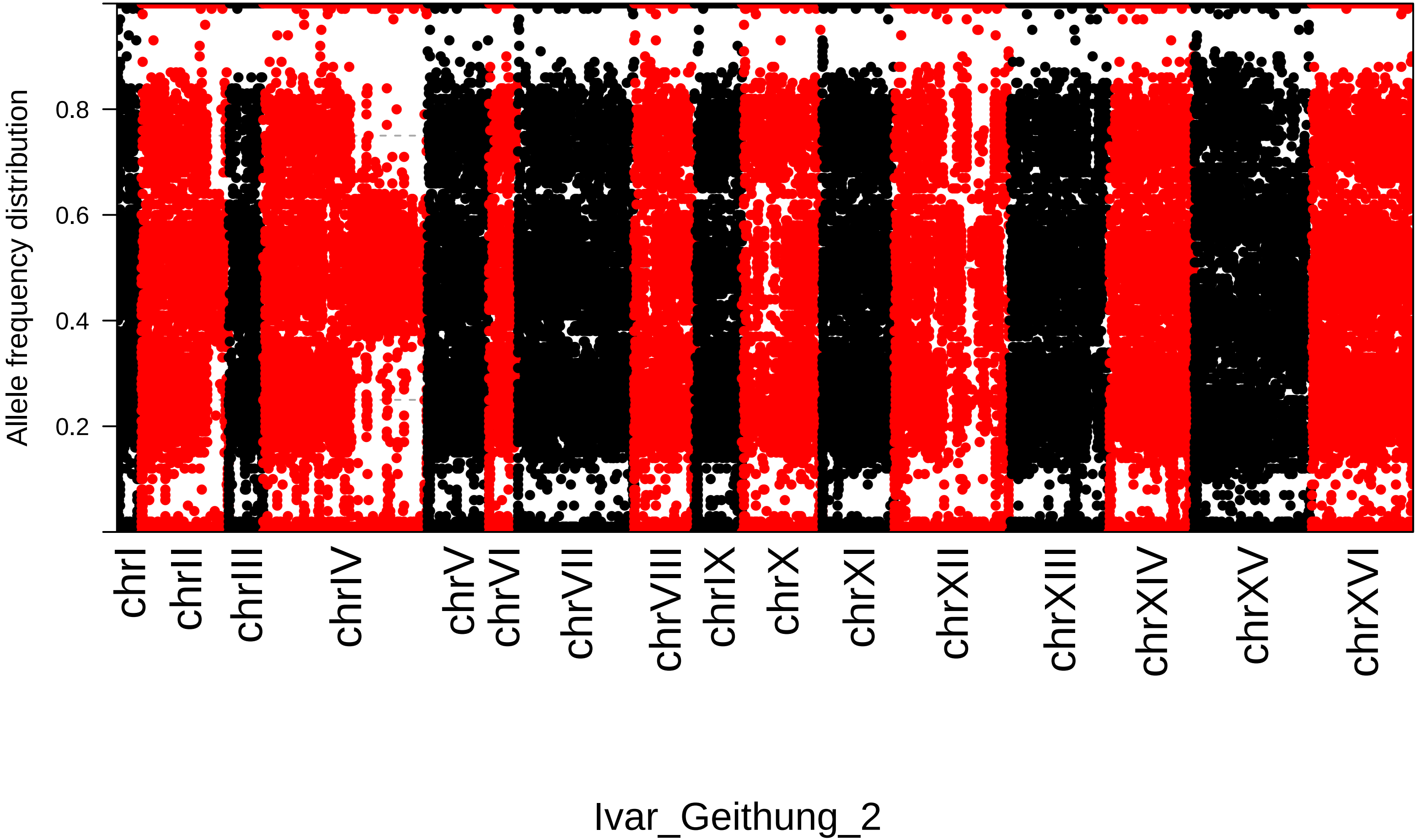
Allele frequency distribution
0.2
0.4
0.6
0.8
chrI
chrII
chrIII
chrIV
chrV
chrVI
chrVII
chrVIII
chrIX
chrX
chrXI
chrXII
chrXIII
chrXIV
chrXV
chrXVI
Ivar_Geithung_2

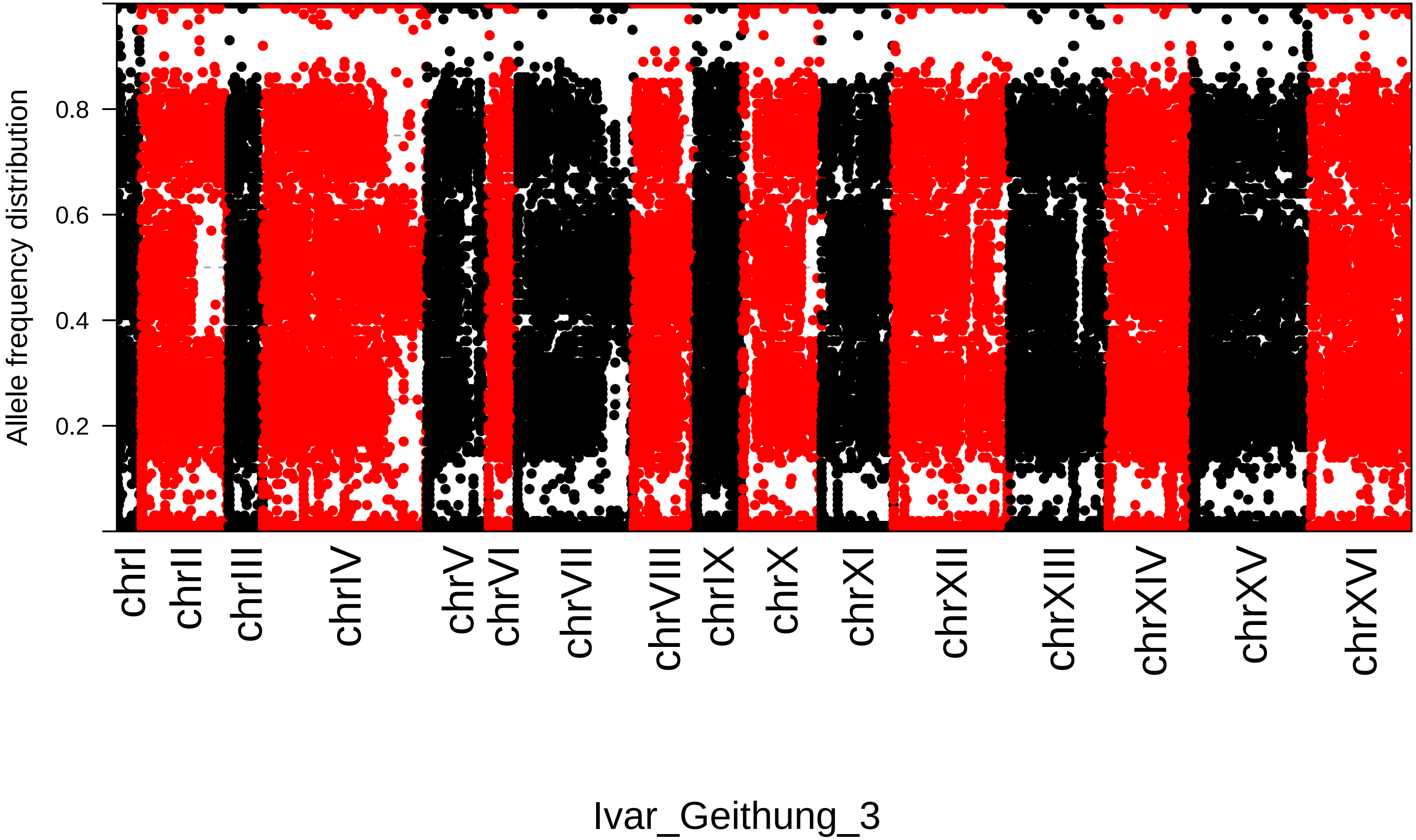

Allele frequency distribution
0.2
0.4
0.6
0.8
chrI
chrII
chrIII
chrIV
chrV
chrVI
chrVII
chrVIII
chrIX
chrX
chrXI
chrXII
chrXIII
chrXIV
chrXV
chrXVI
Ivar_Geithung_3

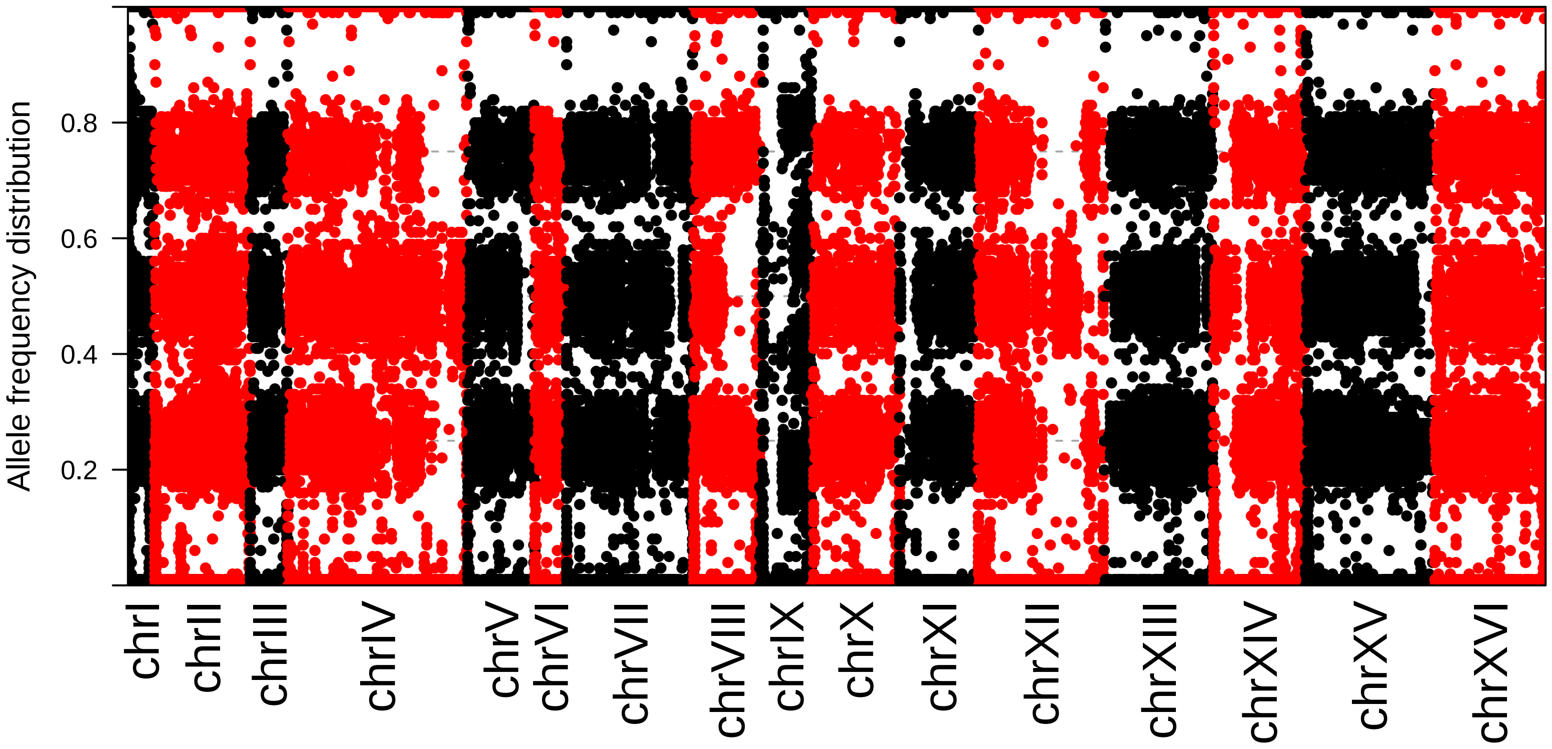
Allele frequency distribution
0.8
0.6
0.4
0.2
chrI
chrII
chrIII
chrIV
chrV
chrVI
chrVII
chrVIII
chrIX
chrX
chrXI
chrXII
chrXIII
chrXIV
chrXV
chrXVI

Jordal1

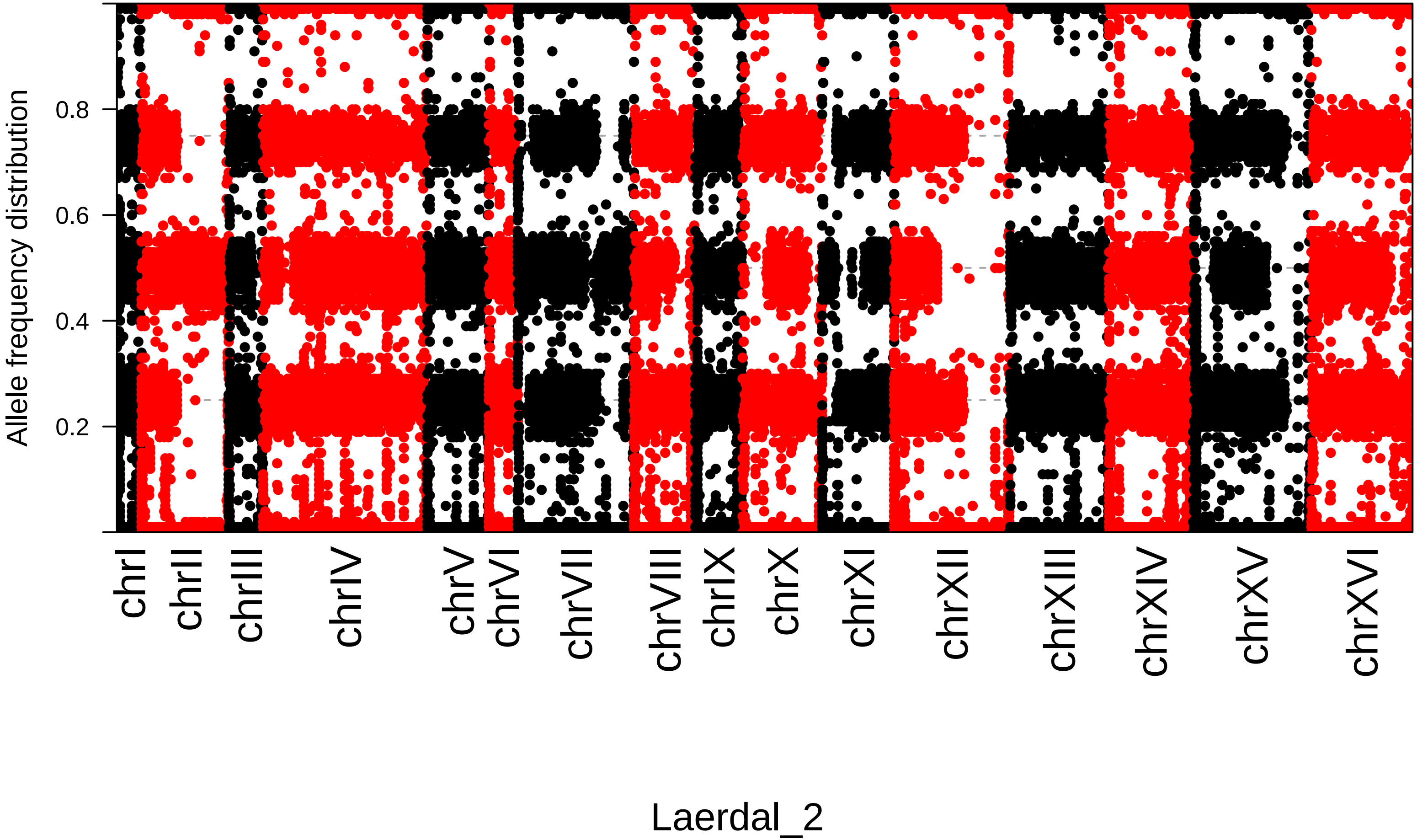
Allele frequency distribution
0.2
0.4
0.6
0.8
chrI
chrII
chrIII
chrIV
chrV
chrVI
chrVII
chrVIII
chrIX
chrX
chrXI
chrXII
chrXIII
chrXIV
chrXV
chrXVI
Laerdal_2

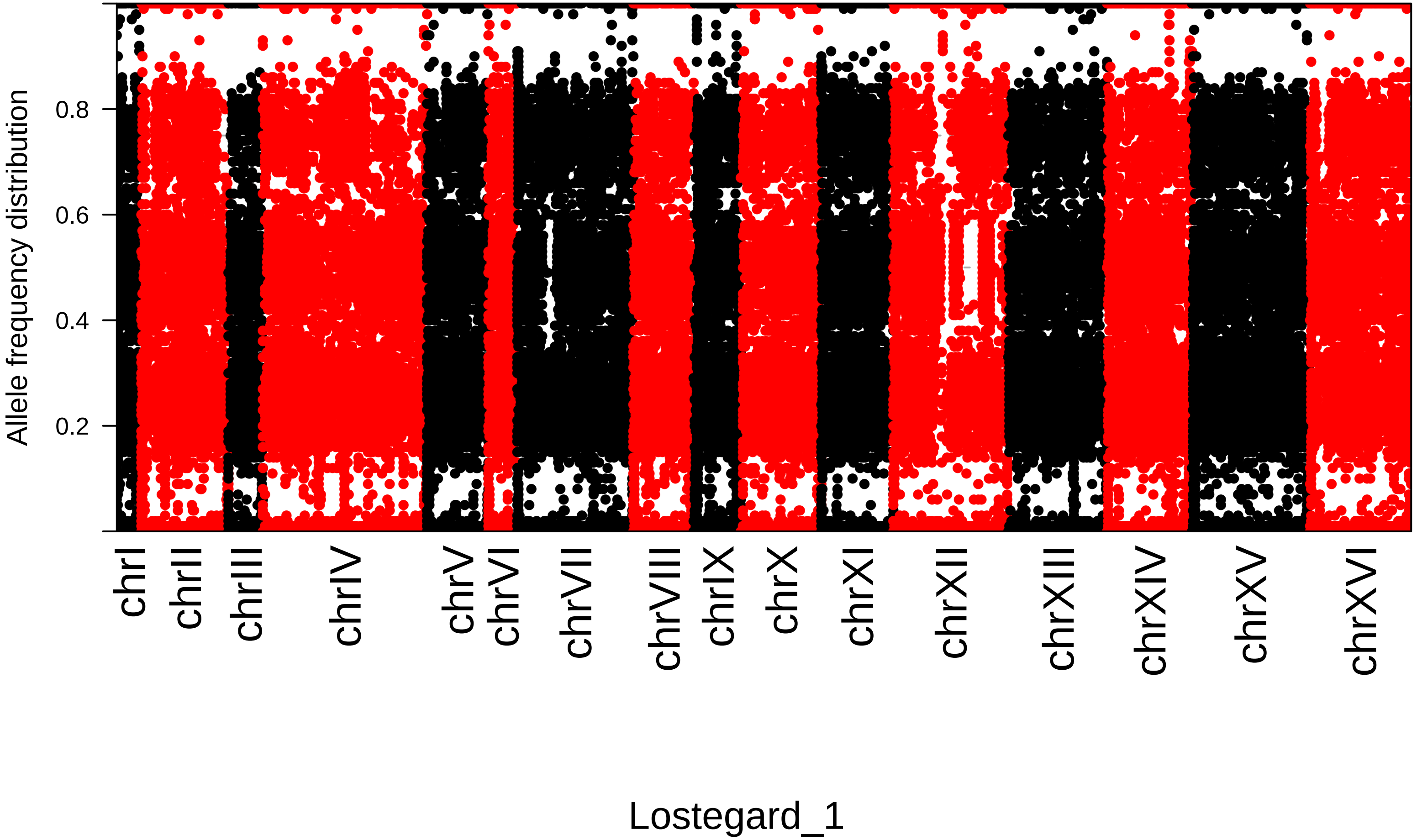
Allele frequency distribution
0.2
0.4
0.6
0.8
chrI
chrII
chrIII
chrIV
chrV
chrVI
chrVII
chrVIII
chrIX
chrX
chrXI
chrXII
chrXIII
chrXIV
chrXV
chrXVI
Lostegard_1

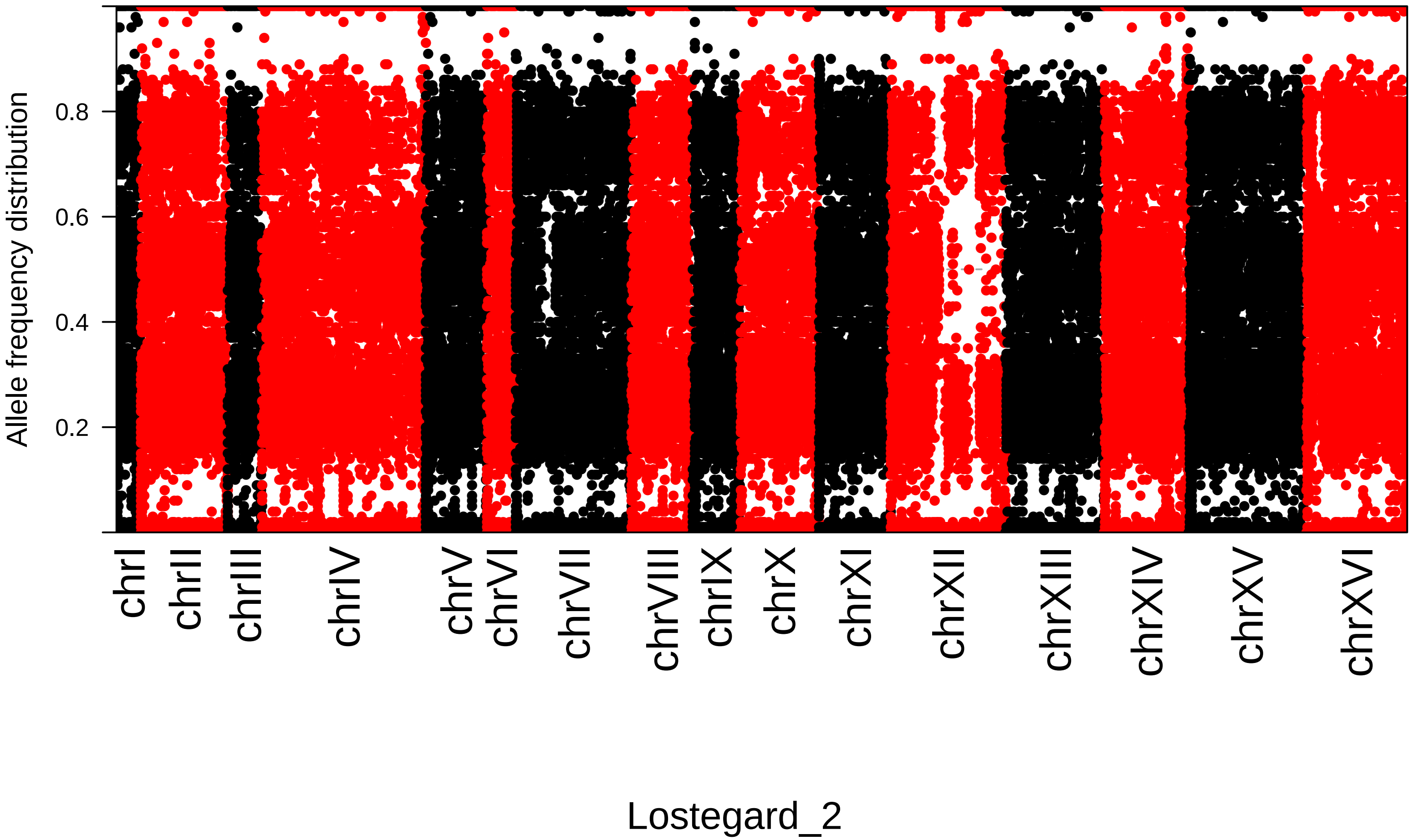
Allele frequency distribution
0.2
0.4
0.6
0.8
chrI
chrII
chrIII
chrIV
chrV
chrVI
chrVII
chrVIII
chrIX
chrX
chrXI
chrXII
chrXIII
chrXIV
chrXV
chrXVI
Lostegard_2

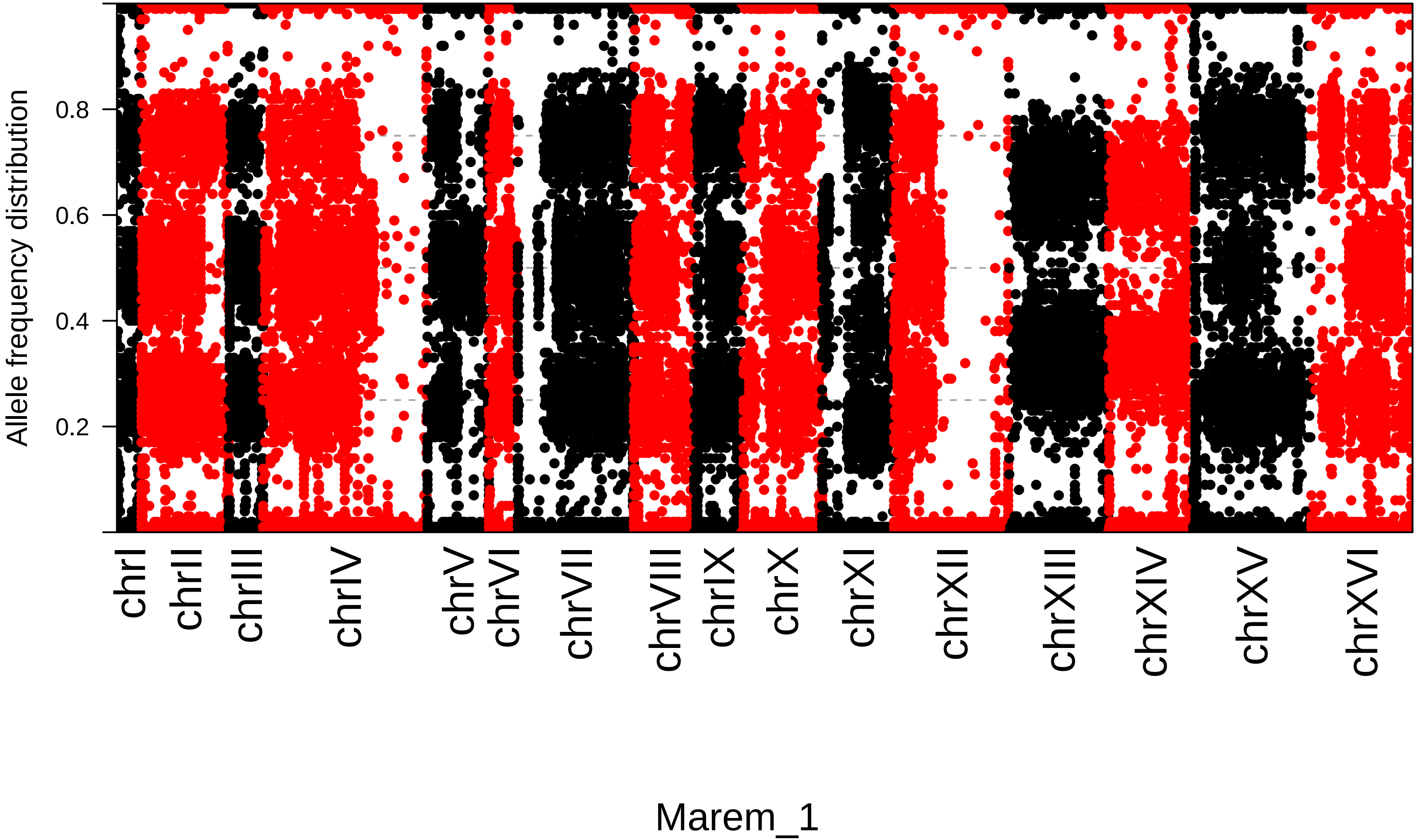
Allele frequency distribution
0.2
0.4
0.6
0.8
chrI
chrII
chrIII
chrIV
chrV
chrVI
chrVII
chrVIII
chrIX
chrX
chrXI
chrXII
chrXIII
chrXIV
chrXV
chrXVI
Marem_1

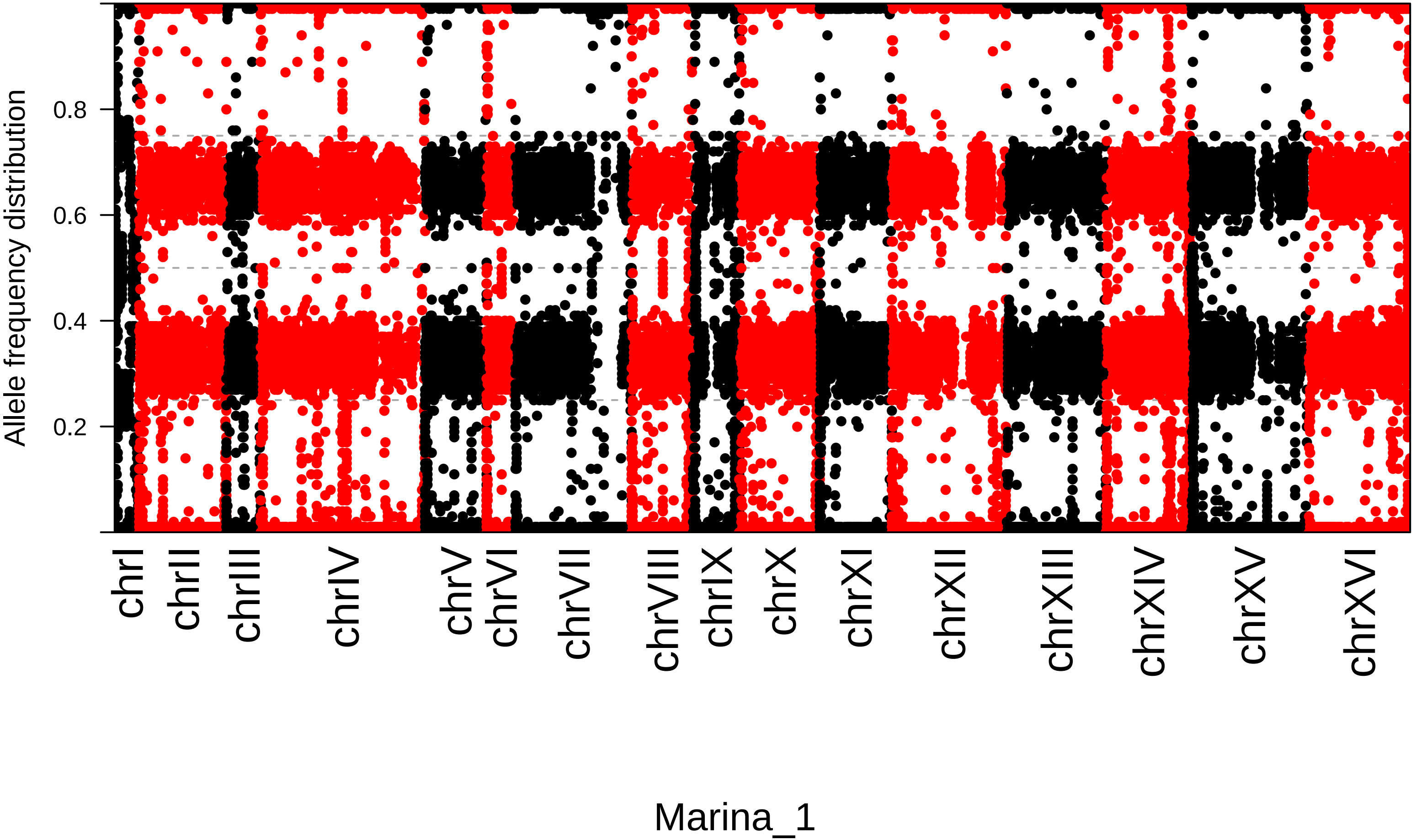
Allele frequency distribution
0.2
0.4
0.6
0.8
chrI
chrII
chrIII
chrIV
chrV
chrVI
chrVII
chrVIII
chrIX
chrX
chrXI
chrXII
chrXIII
chrXIV
chrXV
chrXVI
Marina_1

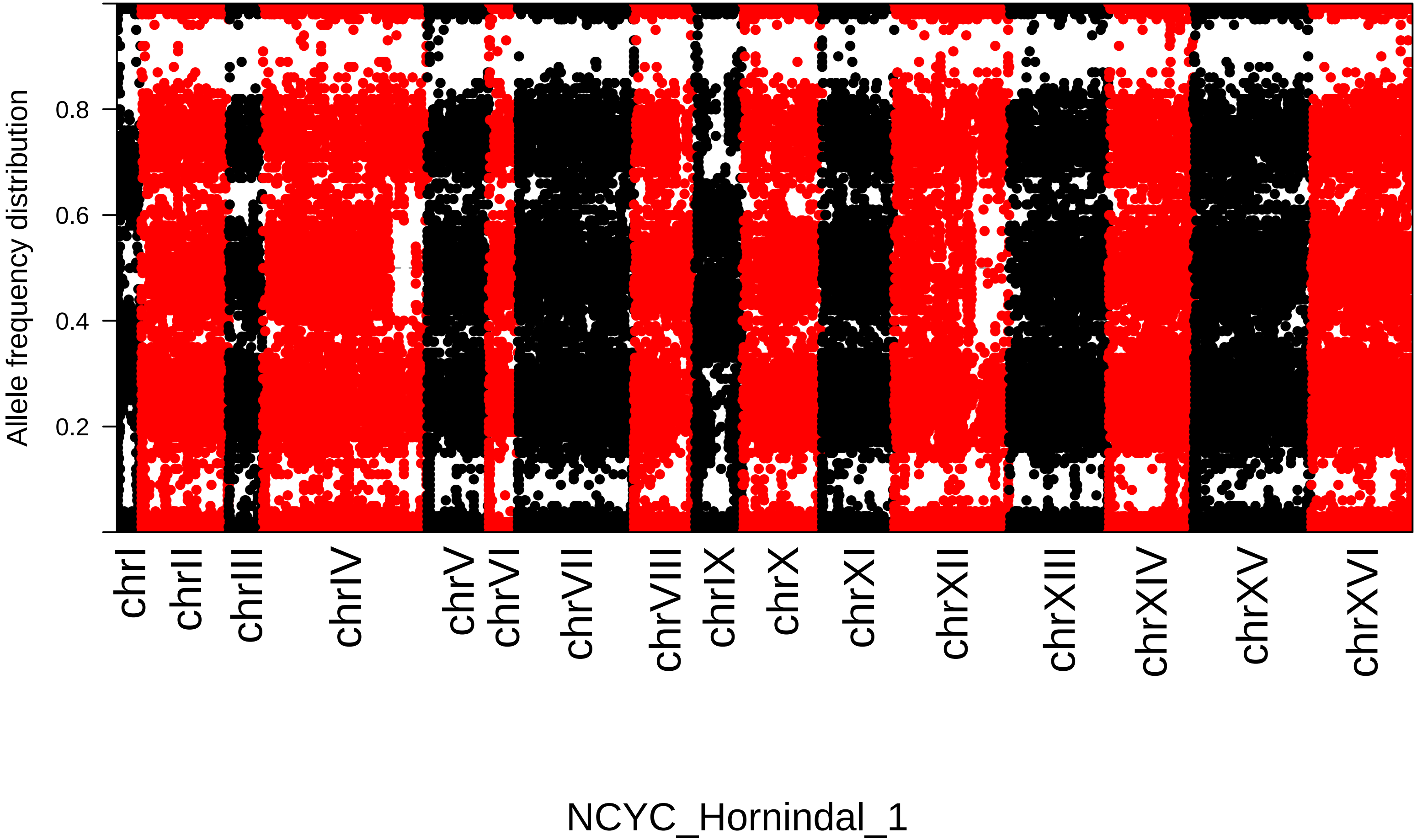

Allele frequency distribution
0.2
0.4
0.6
0.8
chrI
chrII
chrIII
chrIV
chrV
chrVI
chrVII
chrVIII
chrIX
chrX
chrXI
chrXII
chrXIII
chrXIV
chrXV
chrXVI
NCYC_Hornindal_1

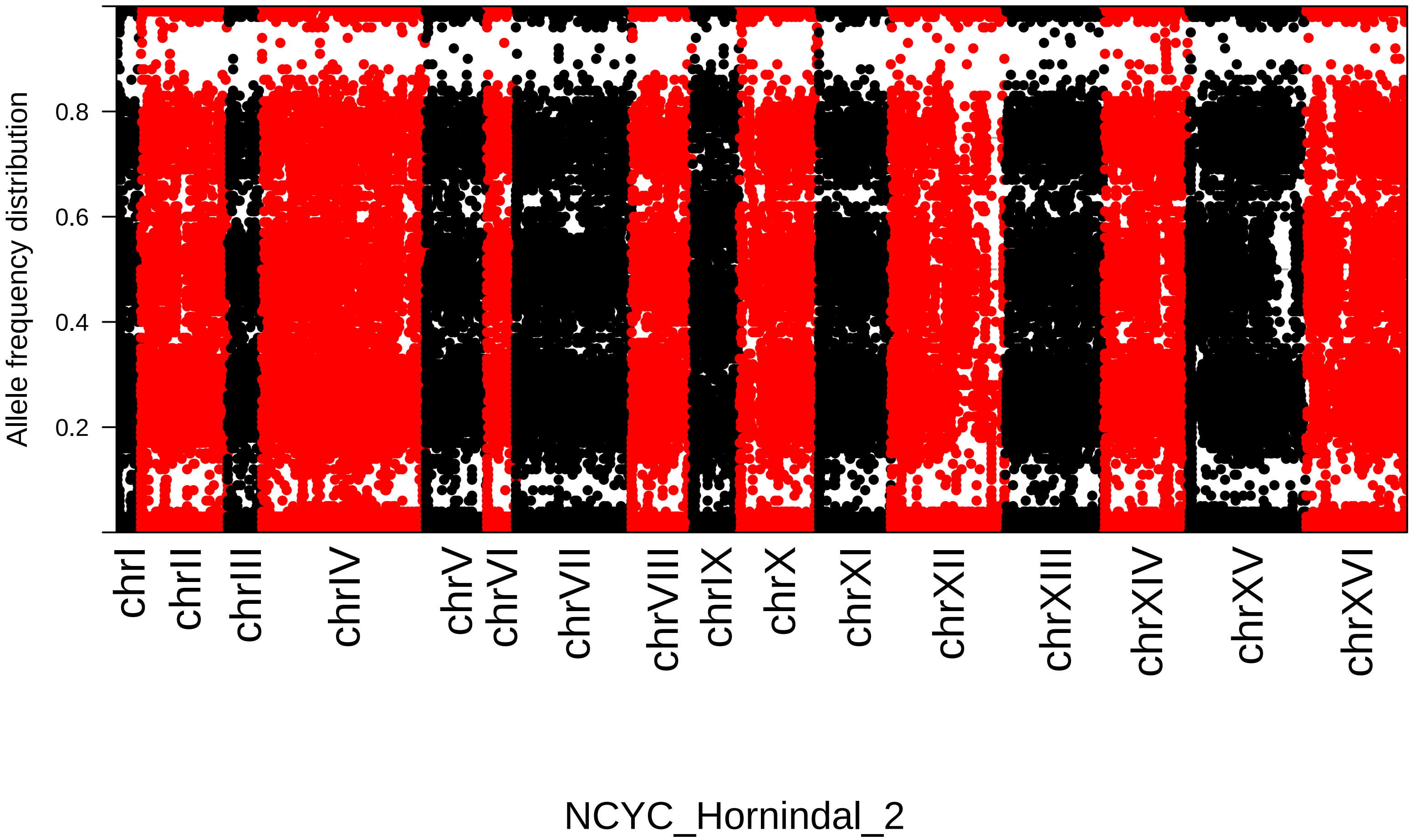

Allele frequency distribution
0.8
0.6
0.4
0.2
chrI
chrII
chrIII
chrIV
chrV
chrVI
chrVII
chrVIII
chrIX
chrX
chrXI
chrXII
chrXIII
chrXIV
chrXV
chrXVI
NCYC_Hornindal_2

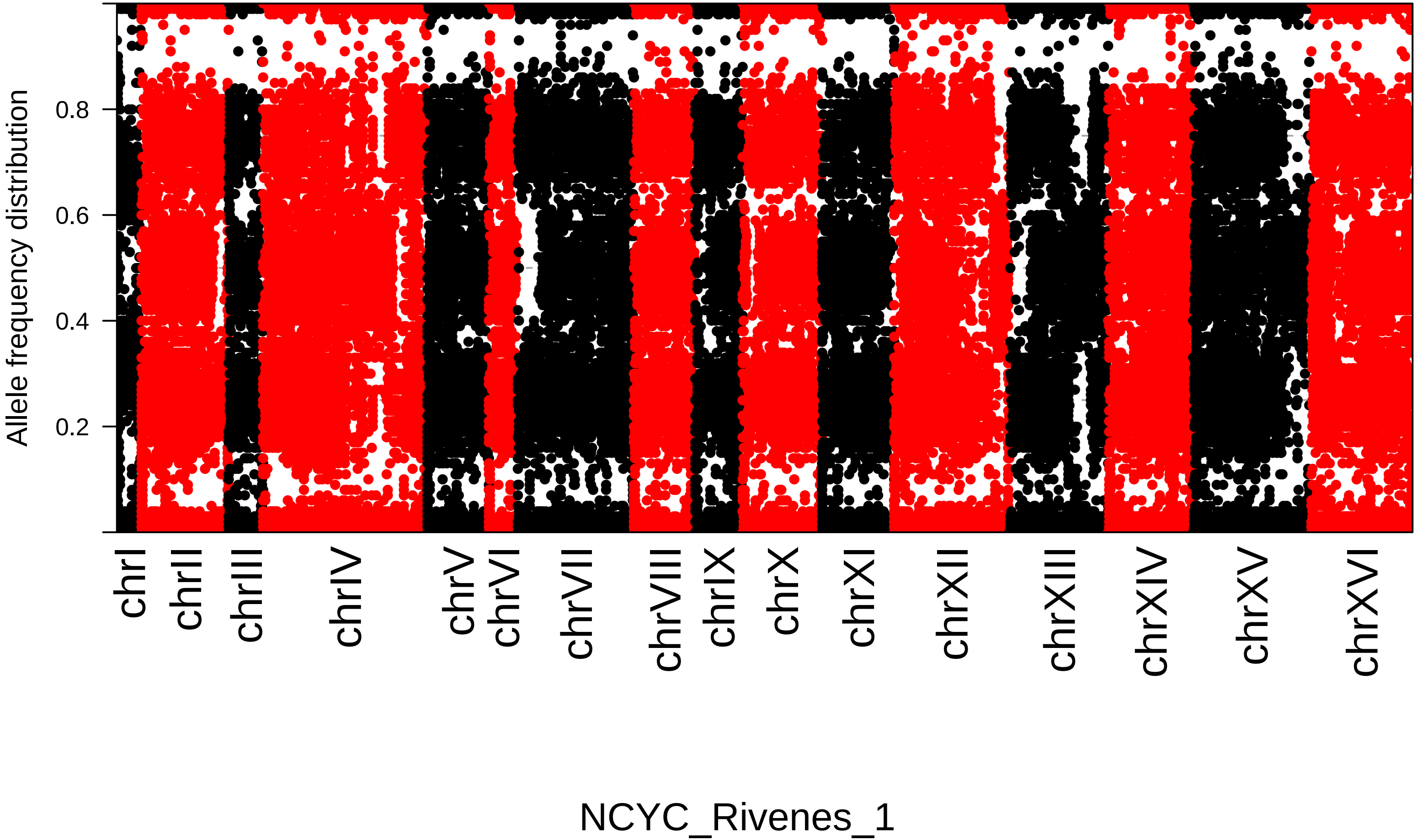
Allele frequency distribution
0.2
0.4
0.6
0.8
chrI
chrII
chrIII
chrIV
chrV
chrVI
chrVII
chrVIII
chrIX
chrX
chrXI
chrXII
chrXIII
chrXIV
chrXV
chrXVI
NCYC_Rivenes_1

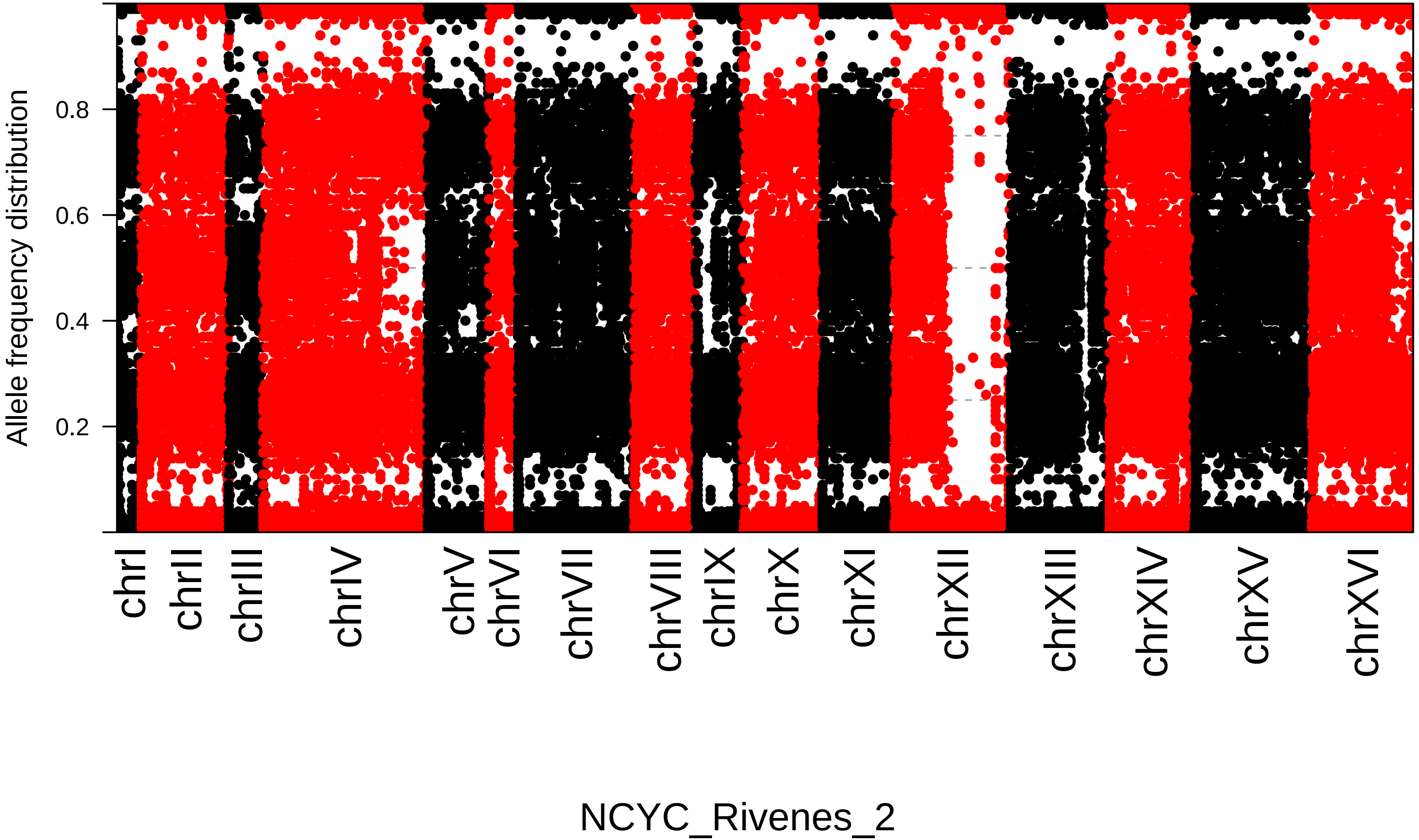
Allele frequency distribution
0.2
0.4
0.6
0.8
chrI
chrII
chrIII
chrIV
chrV
chrVI
chrVII
chrVIII
chrIX
chrX
chrXI
chrXII
chrXIII
chrXIV
chrXV
chrXVI
NCYC_Rivenes_2

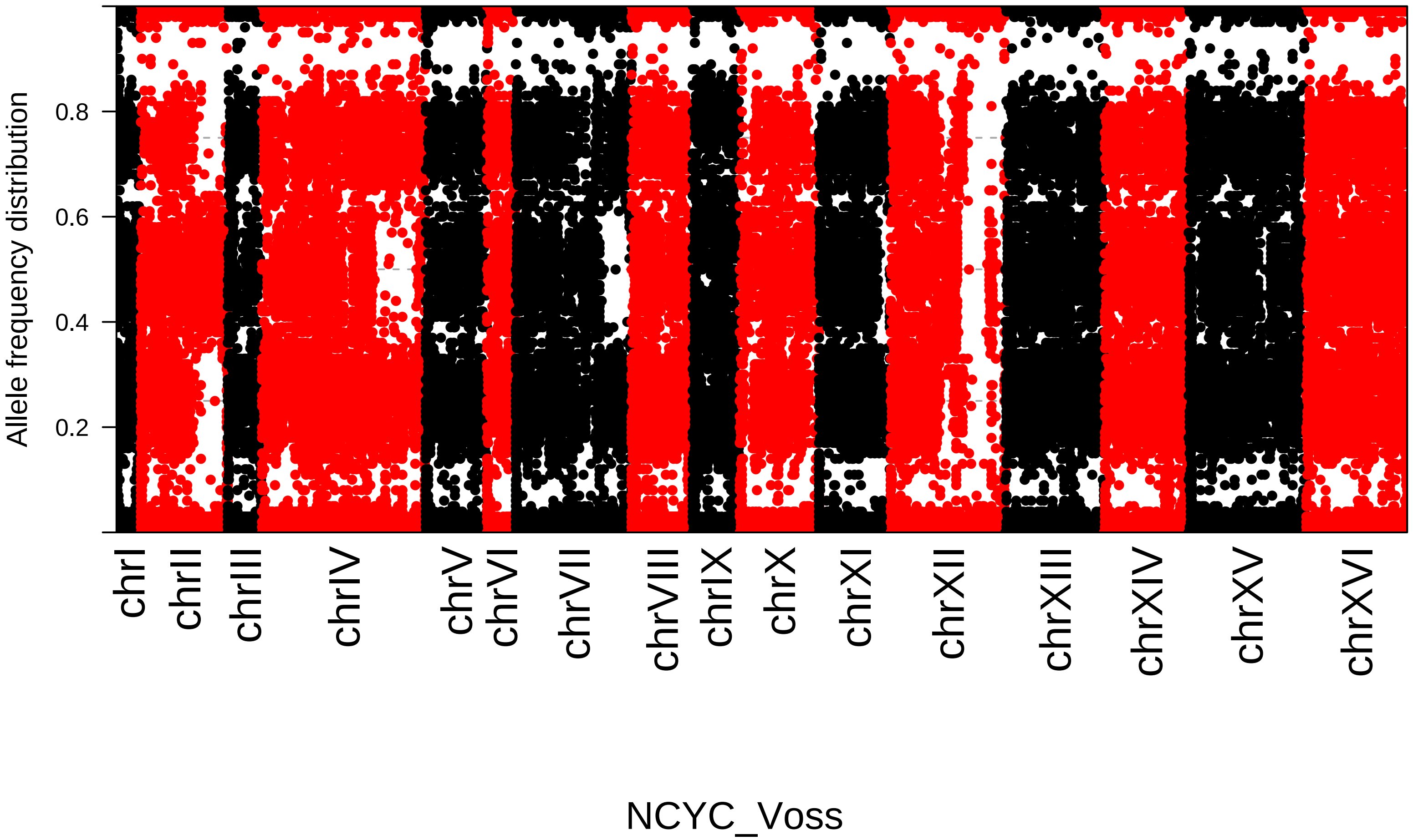

Allele frequency distribution
0.8
0.6
0.4
0.2
chrI
chrII
chrIII
chrIV
chrV
chrVI
chrVII
chrVIII
chrIX
chrX
chrXI
chrXII
chrXIII
chrXIV
chrXV
chrXVI
NCYC_Voss

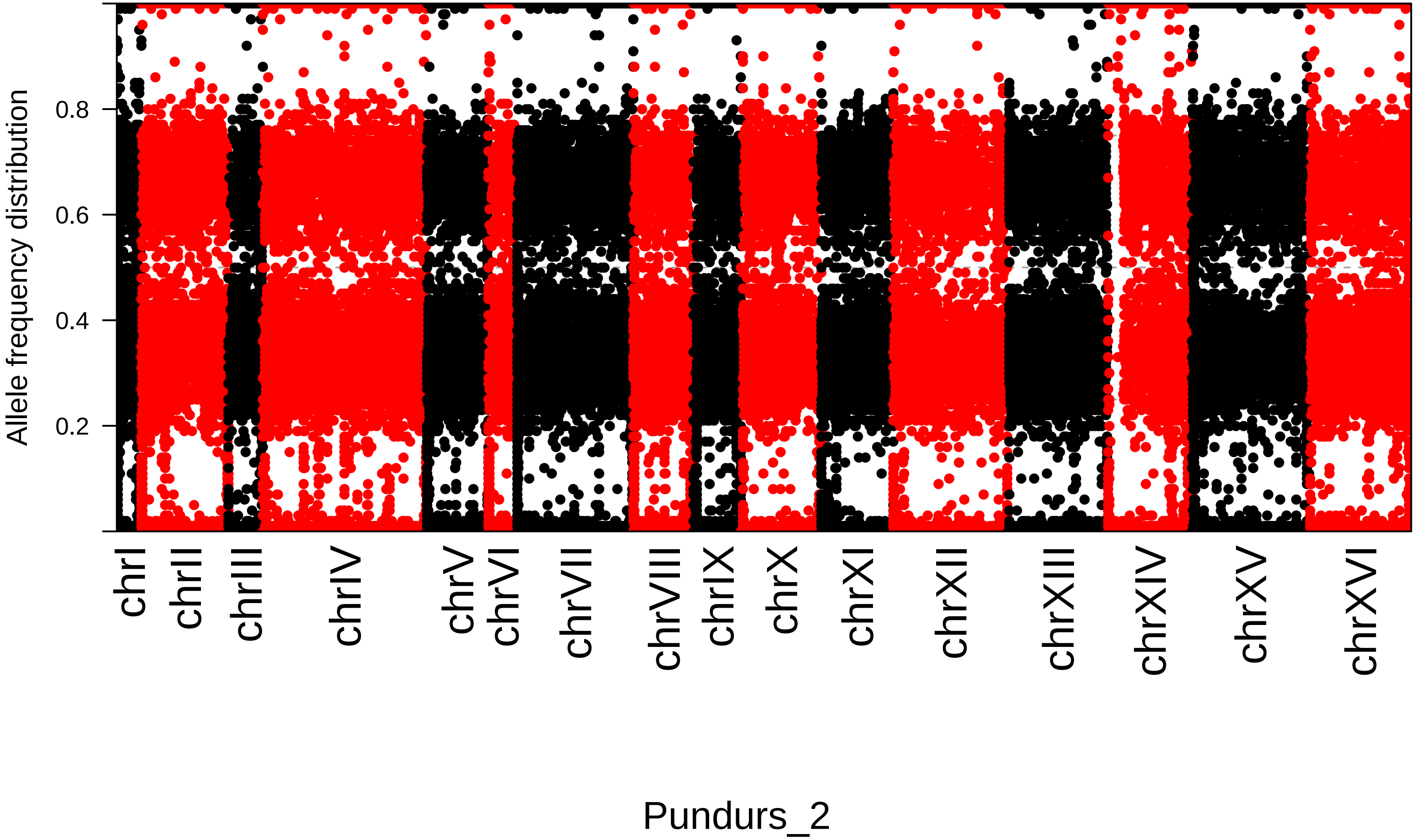

Allele frequency distribution
0.2
0.4
0.6
0.8
chrI
chrII
chrIII
chrIV
chrV
chrVI
chrVII
chrVIII
chrIX
chrX
chrXI
chrXII
chrXIII
chrXIV
chrXV
chrXVI
Pundurs_2

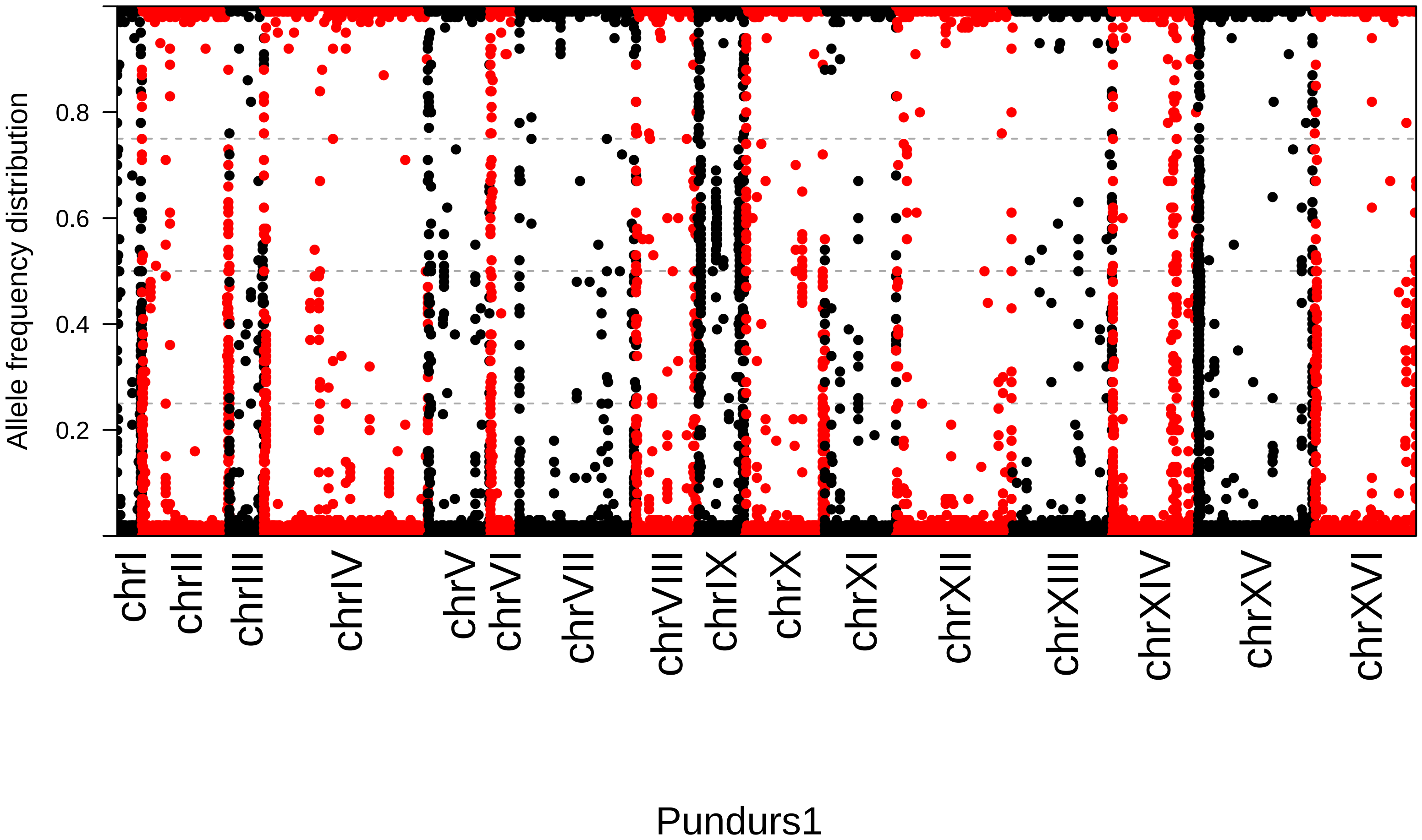

Allele frequency distribution
0.2
0.4
0.6
0.8
chrI
chrII
chrIII
chrIV
chrV
chrVI
chrVII
chrVIII
chrIX
chrX
chrXI
chrXII
chrXIII
chrXIV
chrXV
chrXVI
Pundurs1

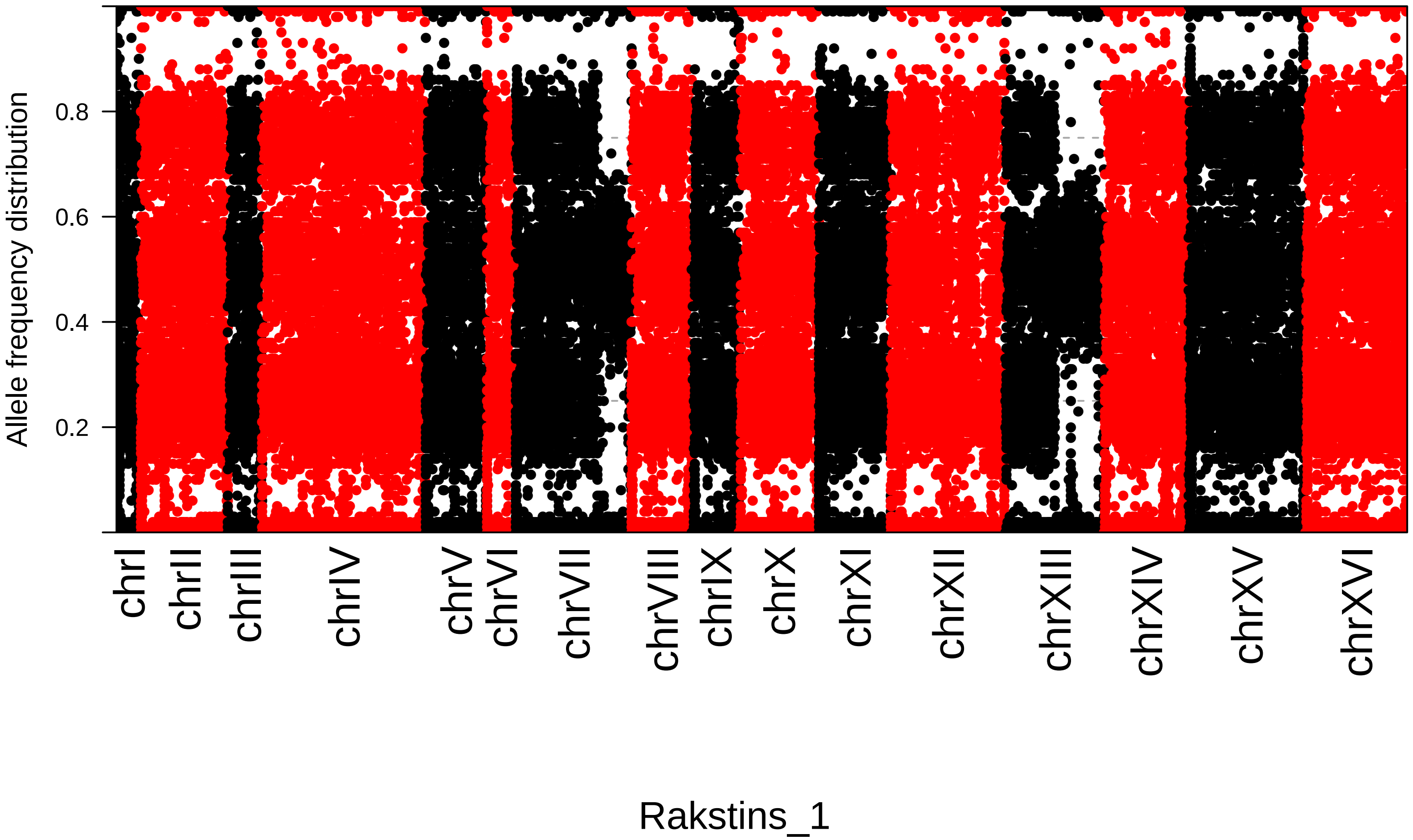
Allele frequency distribution
0.2
0.4
0.6
0.8
chrI
chrII
chrIII
chrIV
chrV
chrVI
chrVII
chrVIII
chrIX
chrX
chrXI
chrXII
chrXIII
chrXIV
chrXV
chrXVI
Rakstins_1

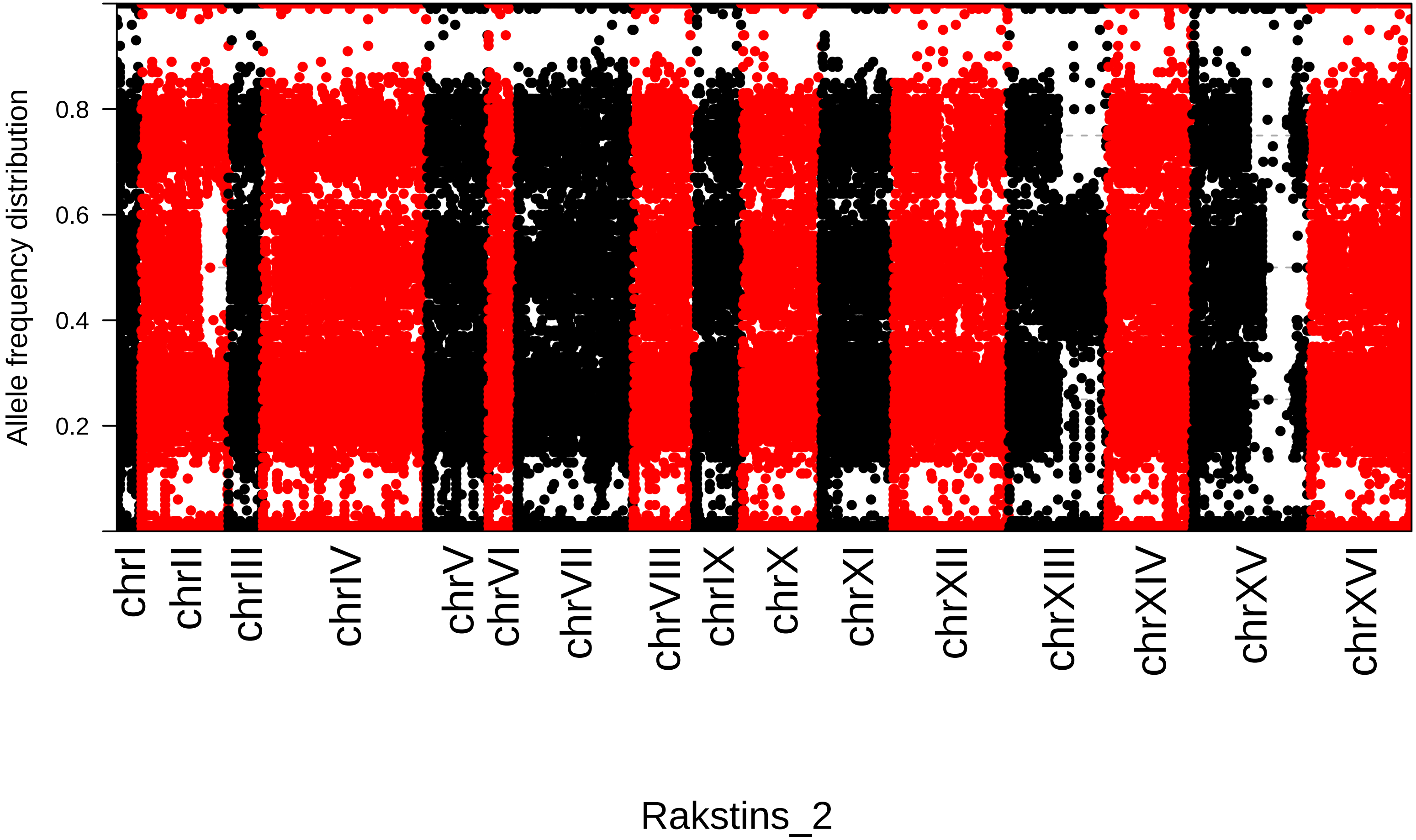
Allele frequency distribution
0.2
0.4
0.6
0.8
chrI
chrII
chrIII
chrIV
chrV
chrVI
chrVII
chrVIII
chrIX
chrX
chrXI
chrXII
chrXIII
chrXIV
chrXV
chrXVI
Rakstins_2

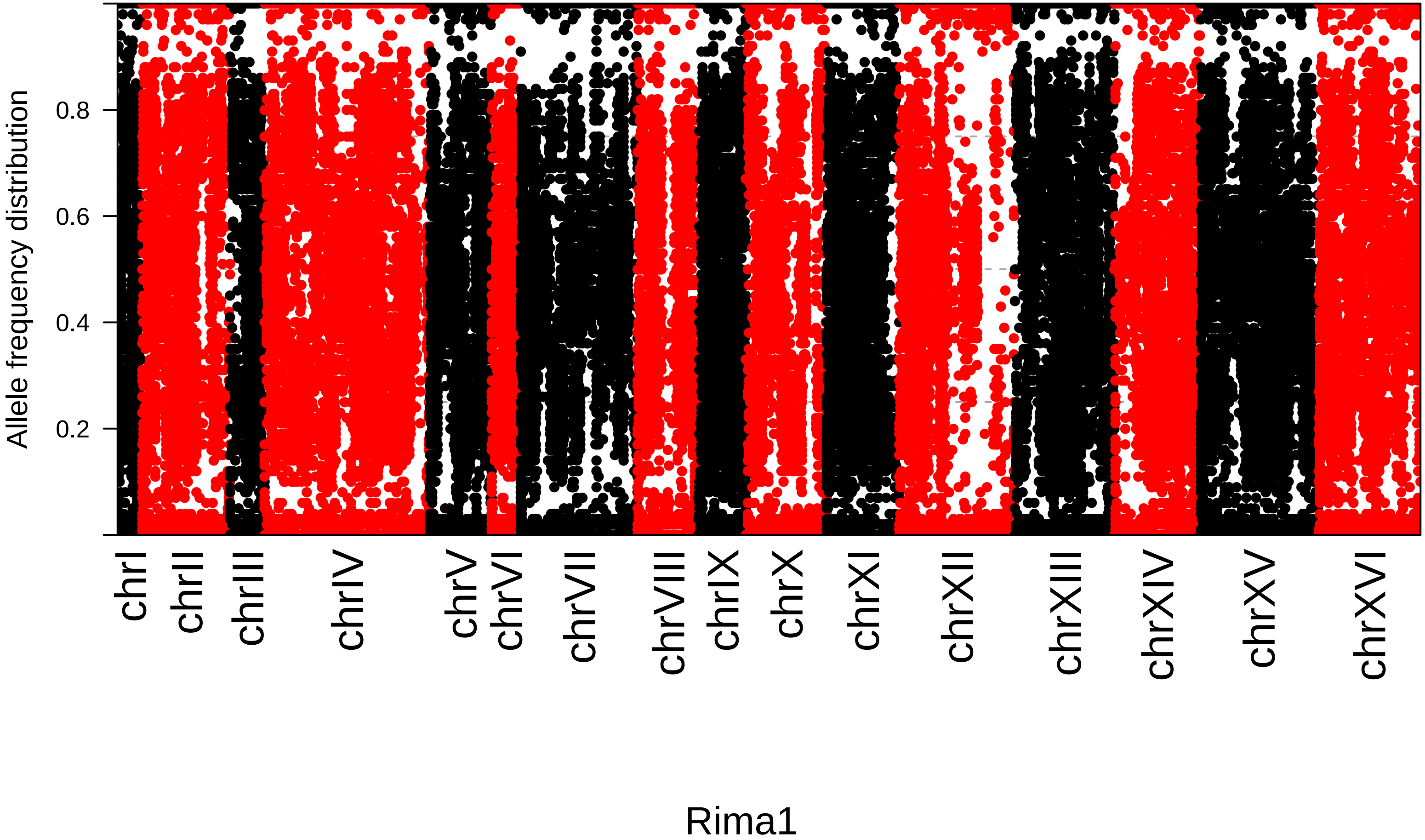

Allele frequency distribution
0.2
0.4
0.6
0.8
chrI
chrII
chrIII
chrIV
chrV
chrVI
chrVII
chrVIII
chrIX
chrX
chrXI
chrXII
chrXIII
chrXIV
chrXV
chrXVI
Rima1

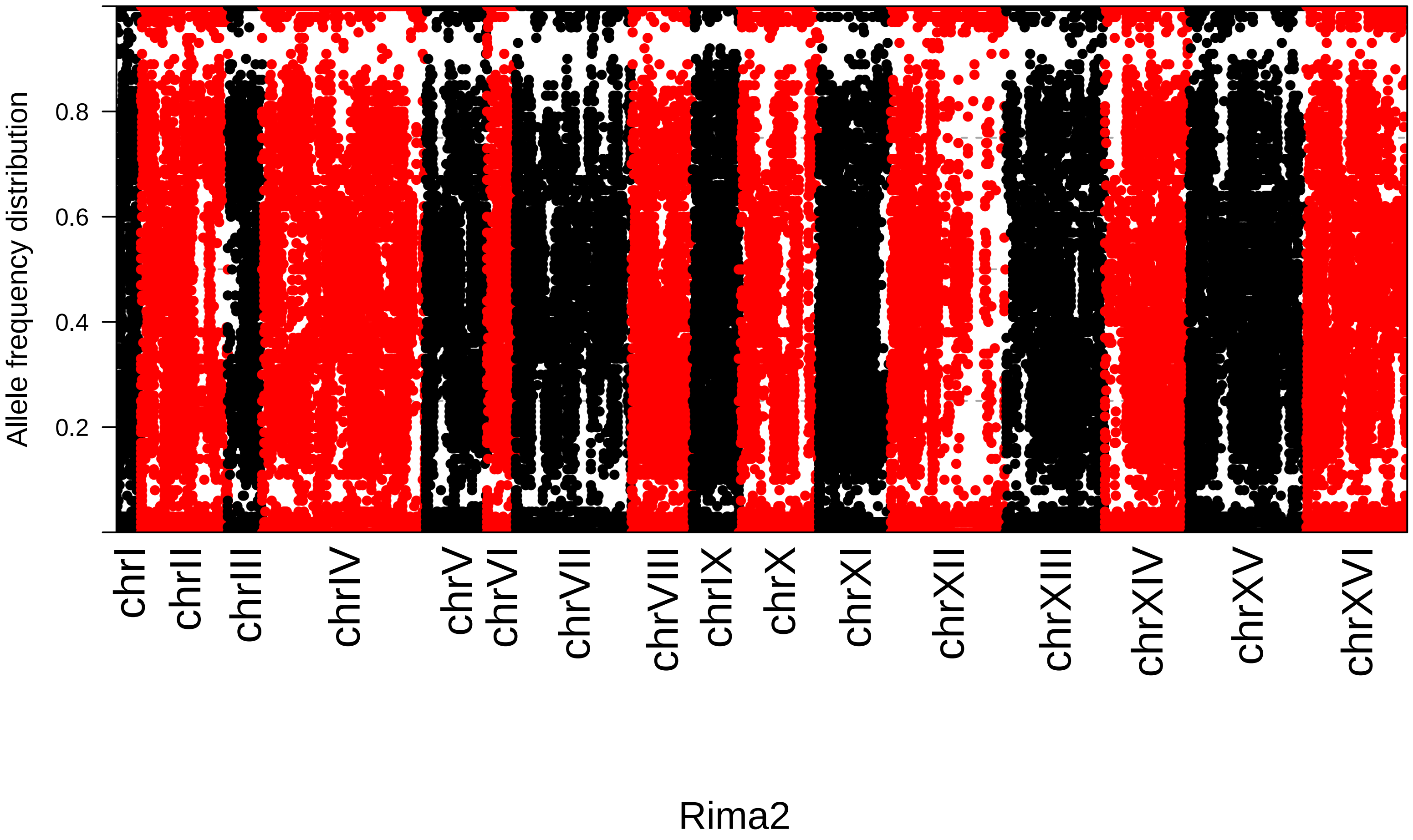
Allele frequency distribution
0.2
0.4
0.6
0.8
chrI
chrII
chrIII
chrIV
chrV
chrVI
chrVII
chrVIII
chrIX
chrX
chrXI
chrXII
chrXIII
chrXIV
chrXV
chrXVI
Rima2

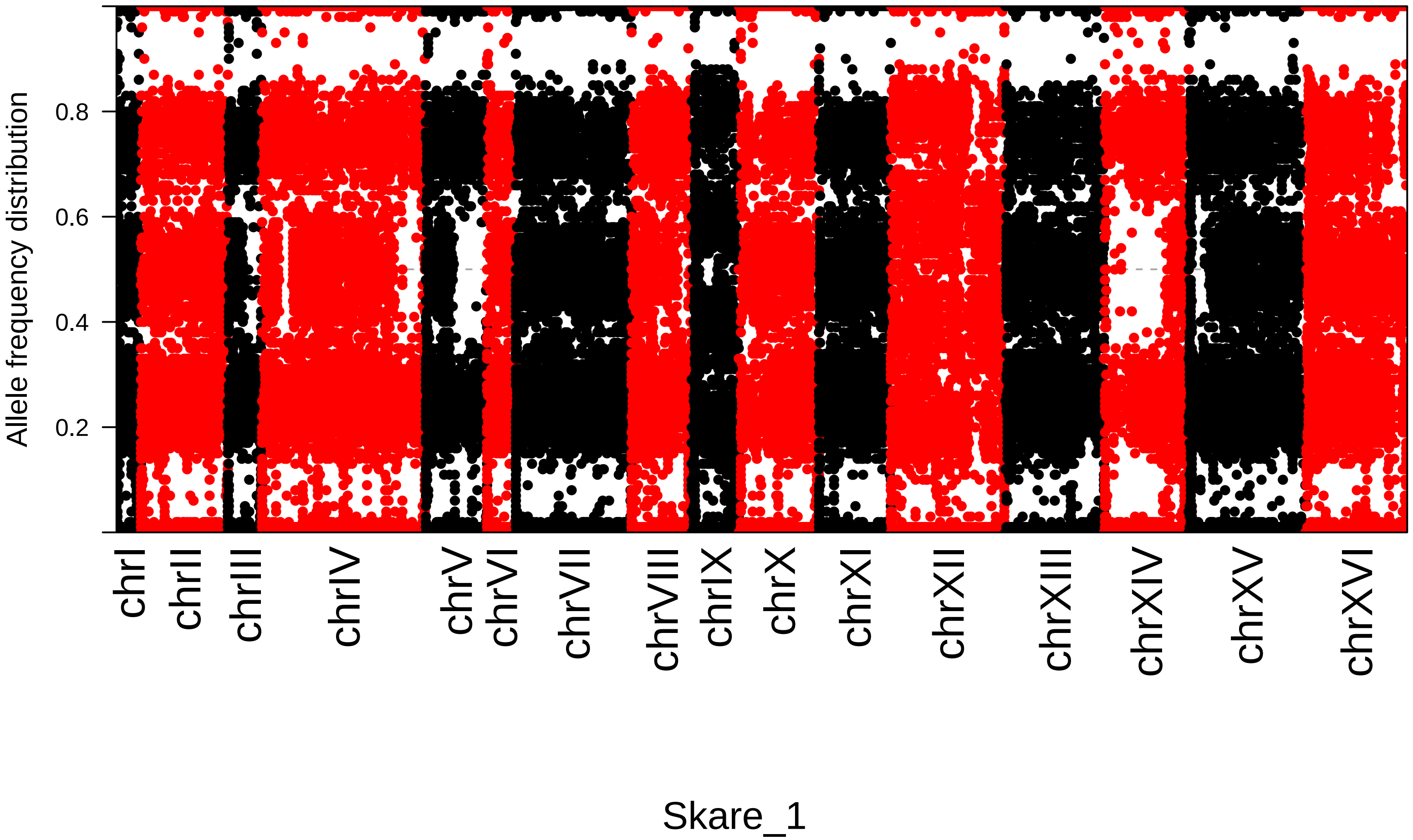

Allele frequency distribution
0.2
0.4
0.6
0.8
chrI
chrII
chrIII
chrIV
chrV
chrVI
chrVII
chrVIII
chrIX
chrX
chrXI
chrXII
chrXIII
chrXIV
chrXV
chrXVI
Skare_1

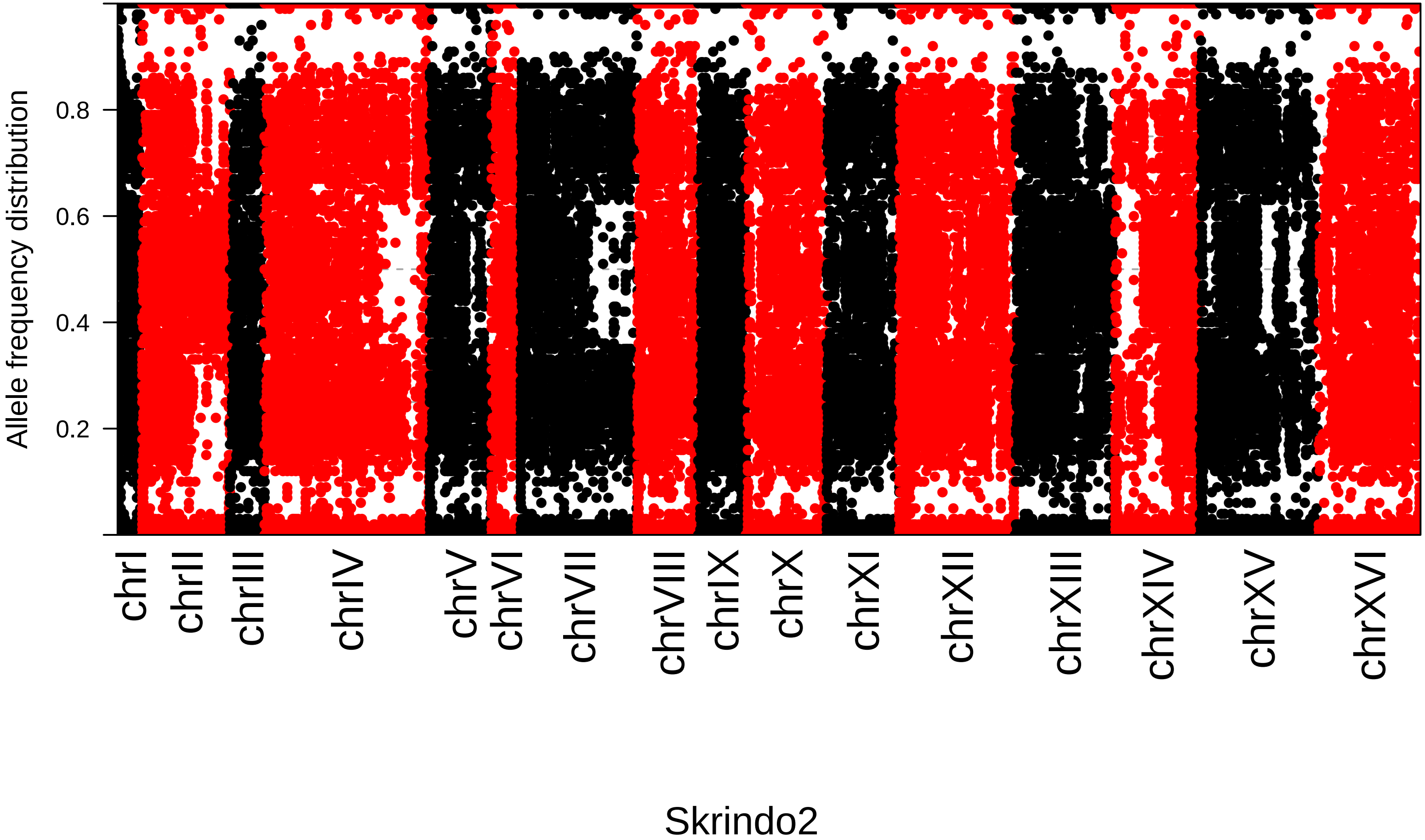

Allele frequency distribution
0.2
0.4
0.6
0.8
chrI
chrII
chrIII
chrIV
chrV
chrVI
chrVII
chrVIII
chrIX
chrX
chrXI
chrXII
chrXIII
chrXIV
chrXV
chrXVI
Skrindo2

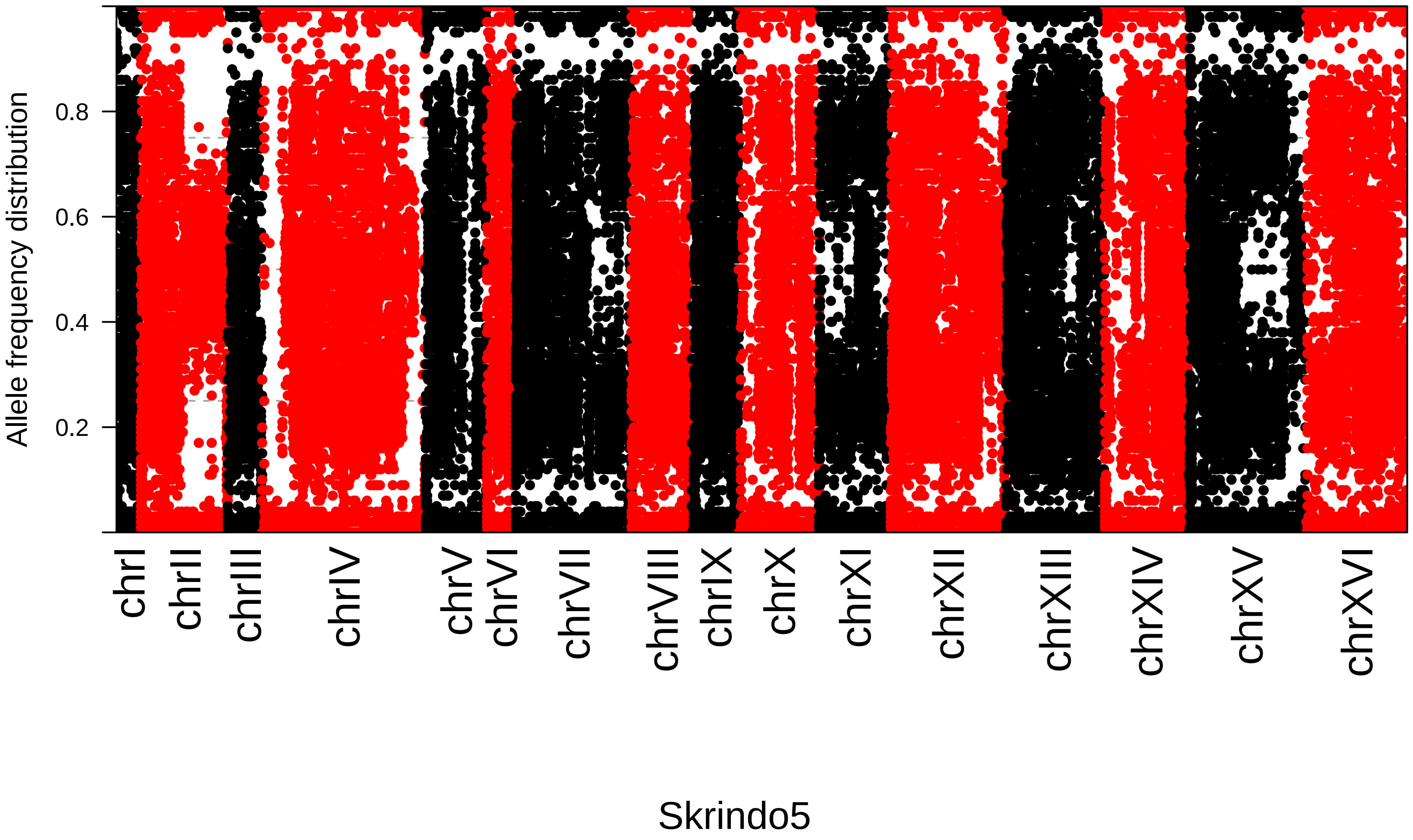

Allele frequency distribution
0.2
0.4
0.6
0.8
chrI
chrII
chrIII
chrIV
chrV
chrVI
chrVII
chrVIII
chrIX
chrX
chrXI
chrXII
chrXIII
chrXIV
chrXV
chrXVI
Skrindo5

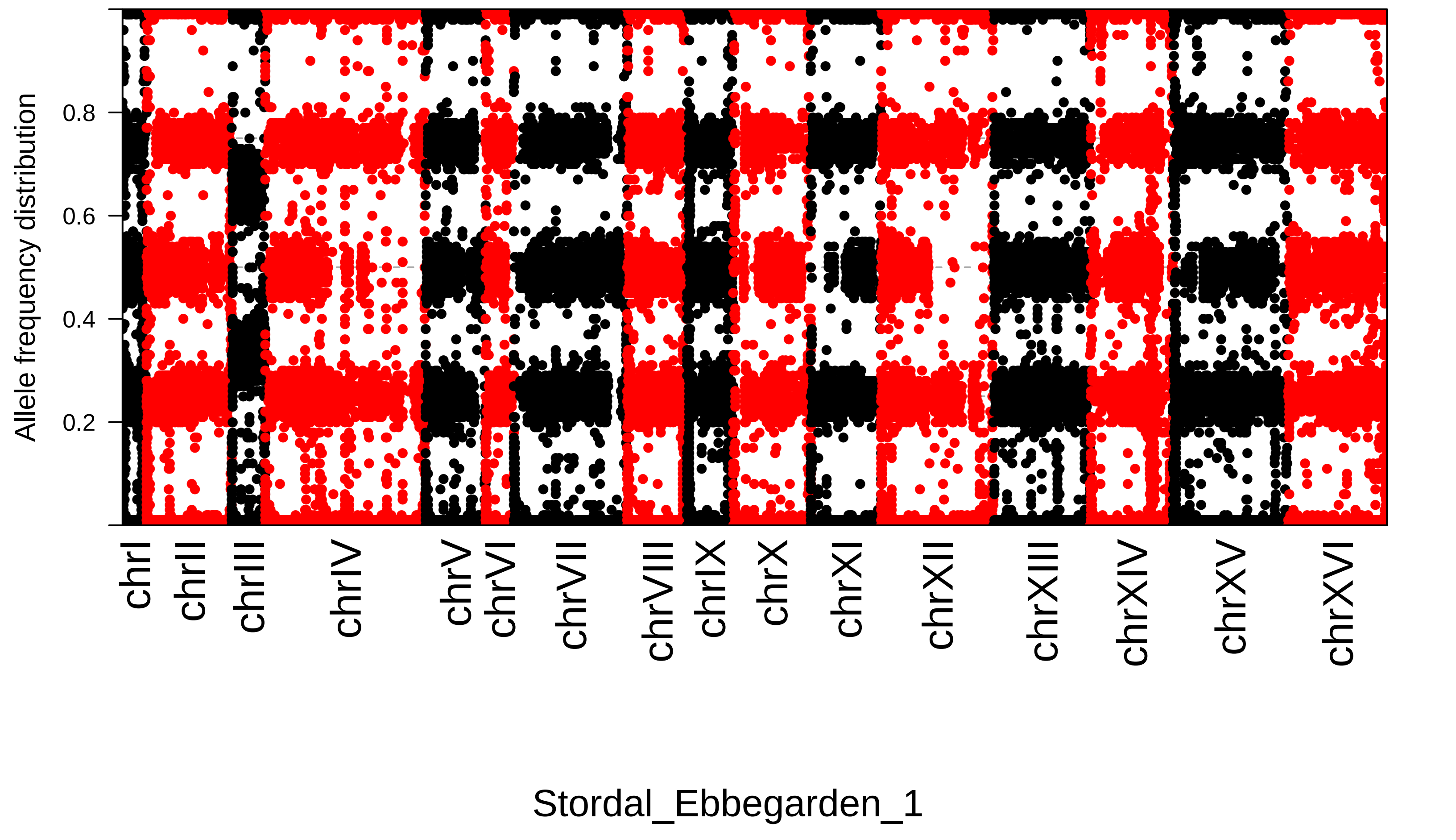

Allele frequency distribution
0.8
0.6
0.4
0.2
chrI
chrII
chrIII
chrIV
chrV
chrVI
chrVII
chrVIII
chrIX
chrX
chrXI
chrXII
chrXIII
chrXIV
chrXV
chrXVI
Stordal_Ebbegarden_1

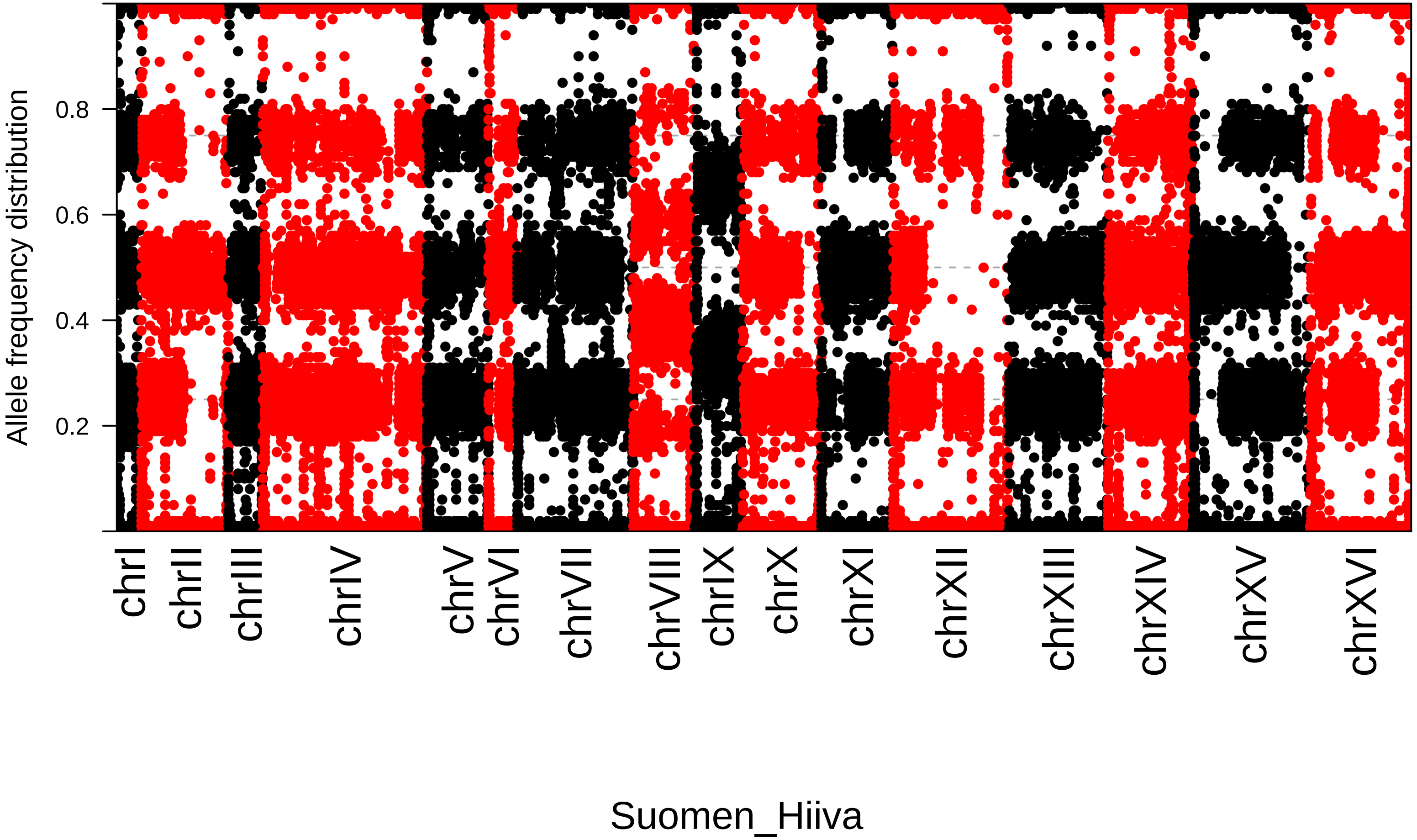

Allele frequency distribution
0.2
0.4
0.6
0.8
chrI
chrII
chrIII
chrIV
chrV
chrVI
chrVII
chrVIII
chrIX
chrX
chrXI
chrXII
chrXIII
chrXIV
chrXV
chrXVI
Suomen_Hiiva

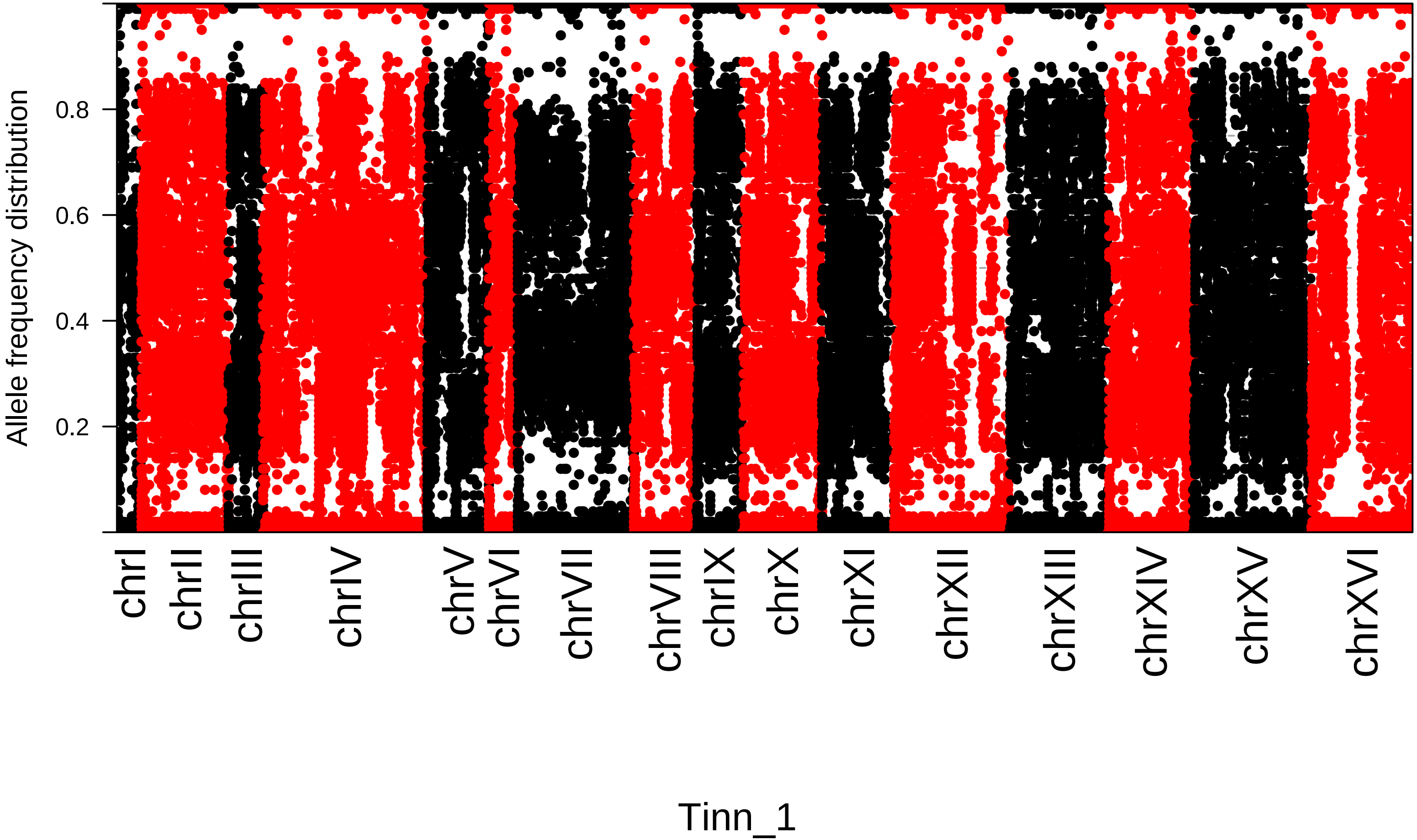

Allele frequency distribution
0.2
0.4
0.6
0.8
chrI
chrII
chrIII
chrIV
chrV
chrVI
chrVII
chrVIII
chrIX
chrX
chrXI
chrXII
chrXIII
chrXIV
chrXV
chrXVI
Tinn_1

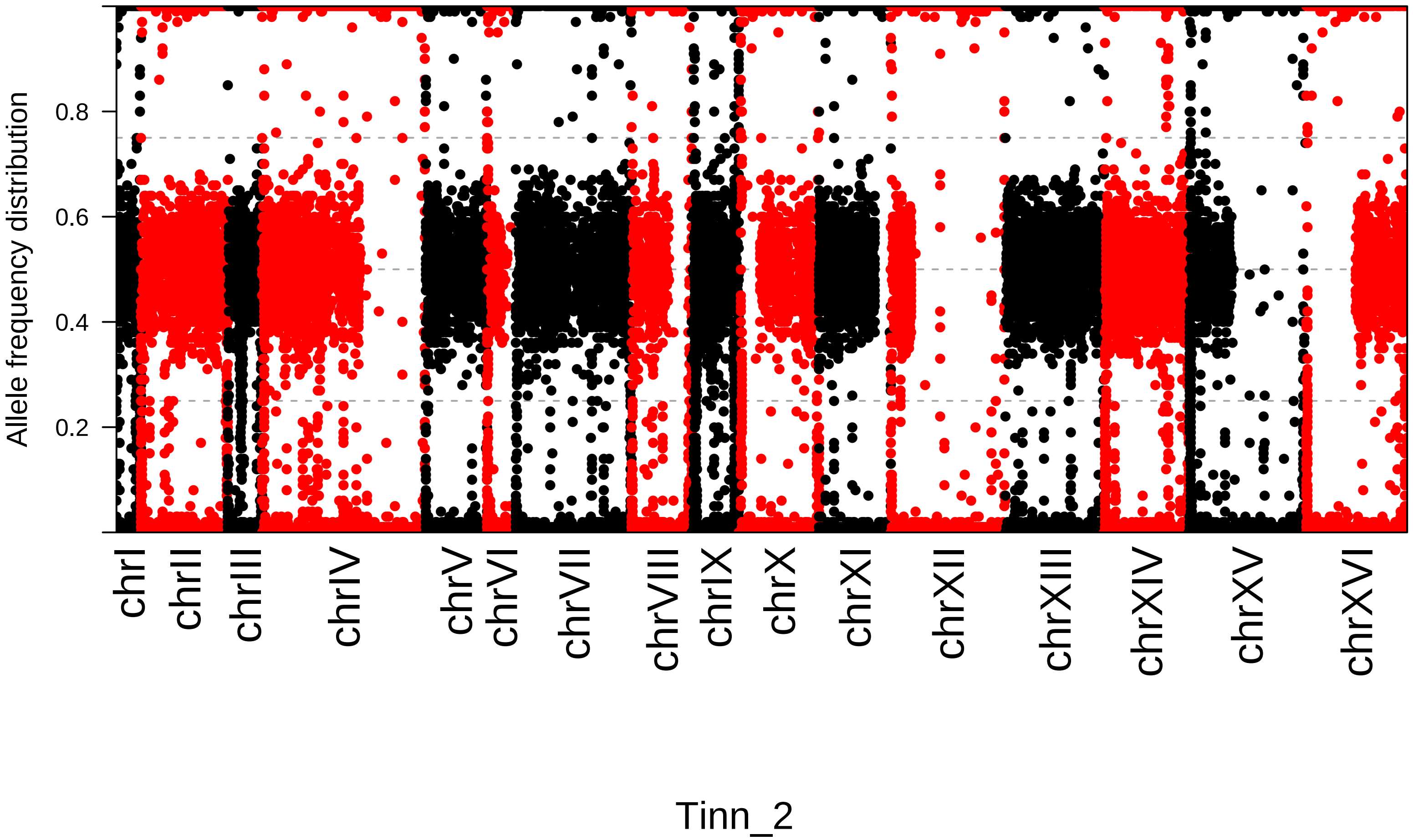

Allele frequency distribution
0.2
0.4
0.6
0.8
chrI
chrII
chrIII
chrIV
chrV
chrVI
chrVII
chrVIII
chrIX
chrX
chrXI
chrXII
chrXIII
chrXIV
chrXV
chrXVI
Tinn_2

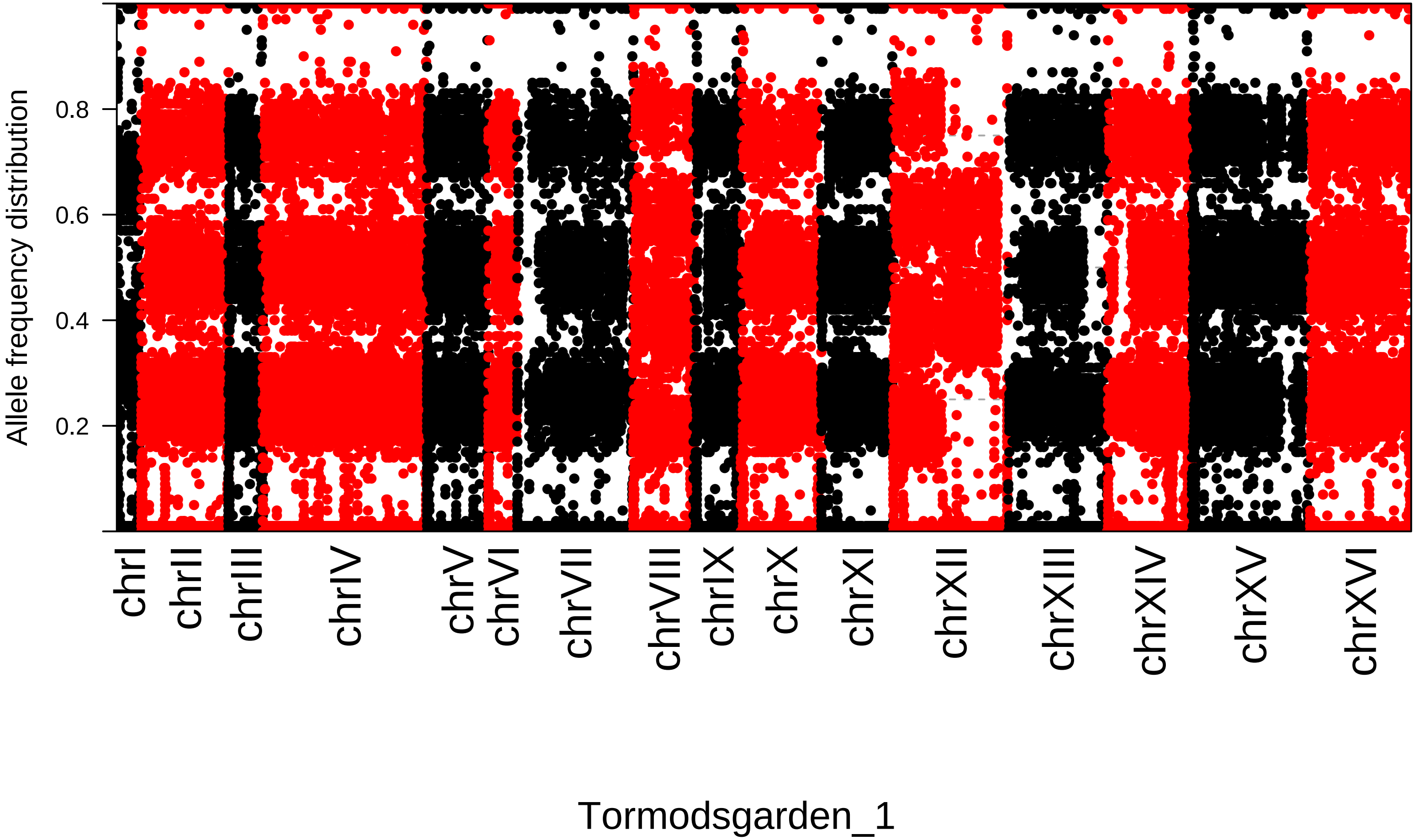
Allele frequency distribution
0.8
0.6
0.4
0.2
chrI
chrII
chrIII
chrIV
chrV
chrVI
chrVII
chrVIII
chrIX
chrX
chrXI
chrXII
chrXIII
chrXIV
chrXV
chrXVI
Tormodsgarden_1

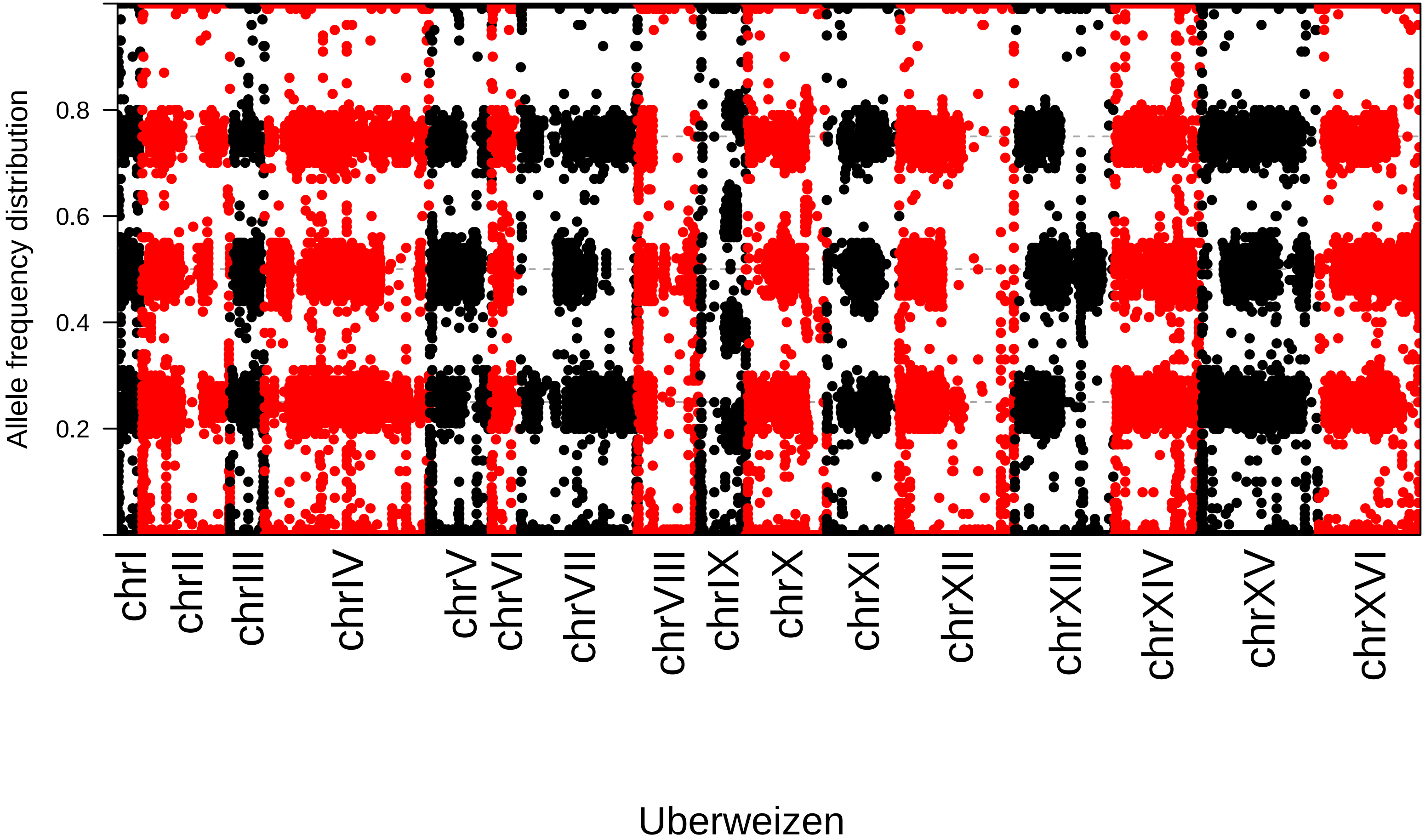

Allele frequency distribution
0.2
0.4
0.6
0.8
chrI
chrII
chrIII
chrIV
chrV
chrVI
chrVII
chrVIII
chrIX
chrX
chrXI
chrXII
chrXIII
chrXIV
chrXV
chrXVI
Uberweizen

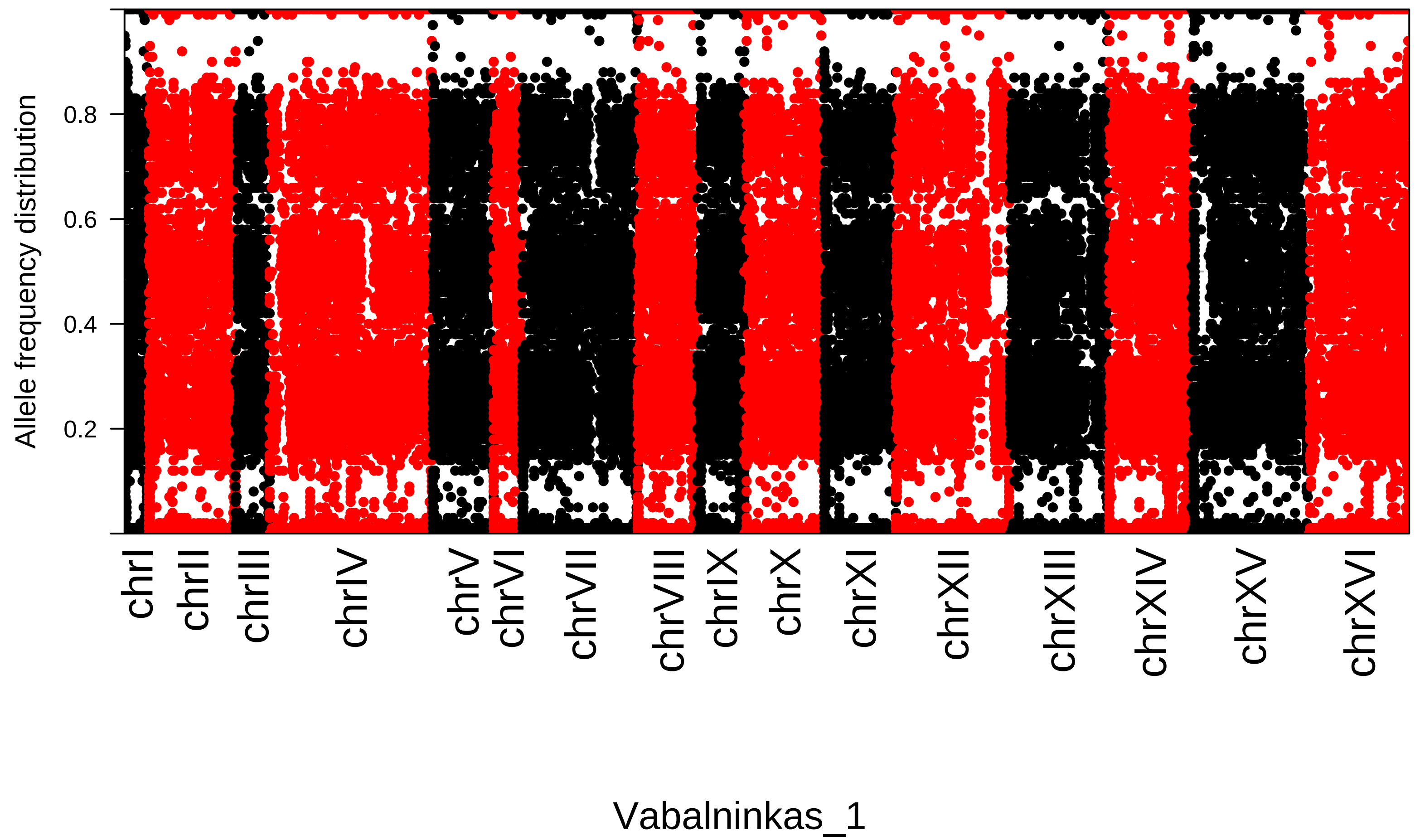

Allele frequency distribution
0.2
0.4
0.6
0.8
chrI
chrII
chrIII
chrIV
chrV
chrVI
chrVII
chrVIII
chrIX
chrX
chrXI
chrXII
chrXIII
chrXIV
chrXV
chrXVI
Vabalninkas_1

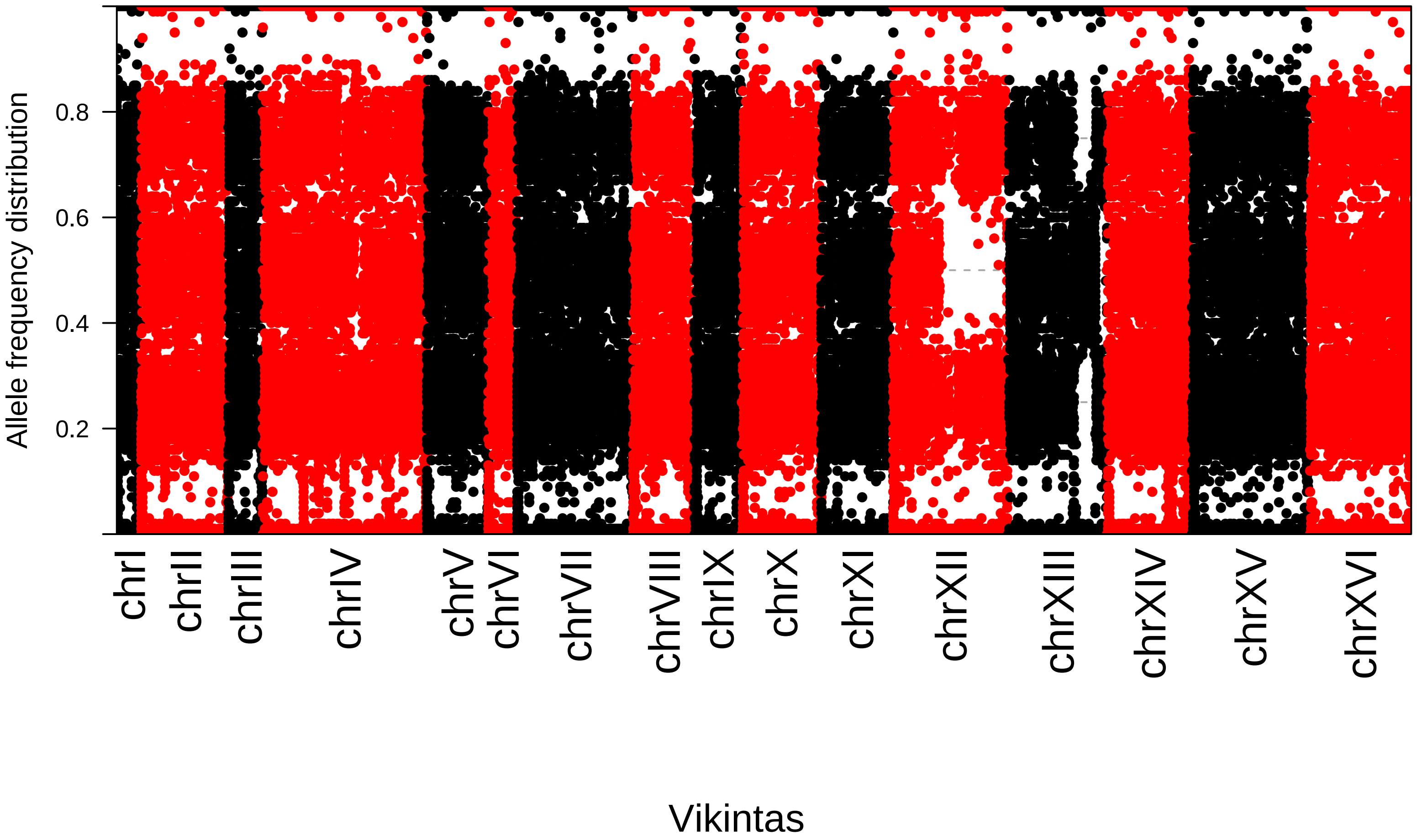
Allele frequency distribution
0.2
0.4
0.6
0.8
chrI
chrII
chrIII
chrIV
chrV
chrVI
chrVII
chrVIII
chrIX
chrX
chrXI
chrXII
chrXIII
chrXIV
chrXV
chrXVI
Vikintas

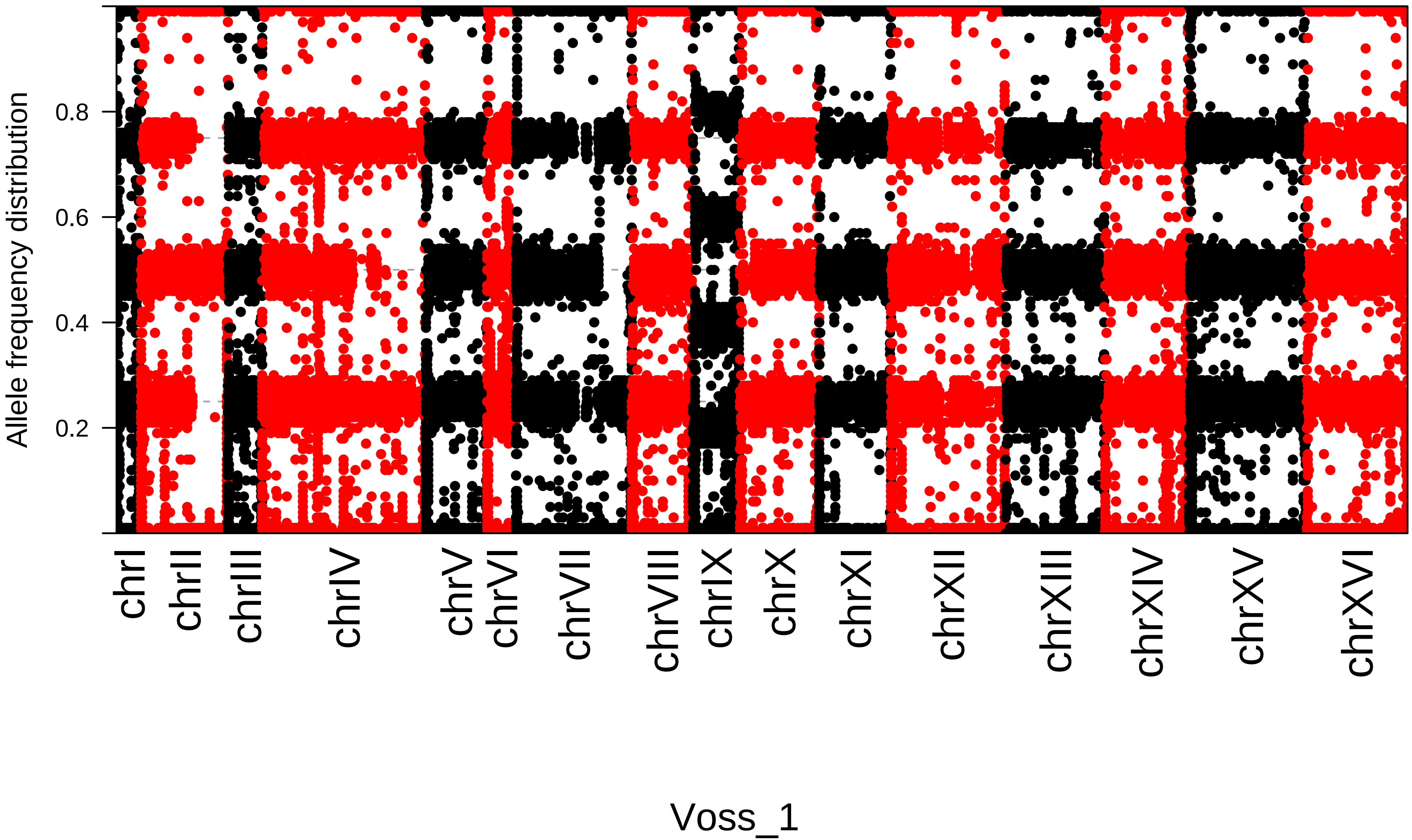
Allele frequency distribution
0.8
0.6
0.4
0.2
chrI
chrII
chrIII
chrIV
chrV
chrVI
chrVII
chrVIII
chrIX
chrX
chrXI
chrXII
chrXIII
chrXIV
chrXV
chrXVI
Voss_1

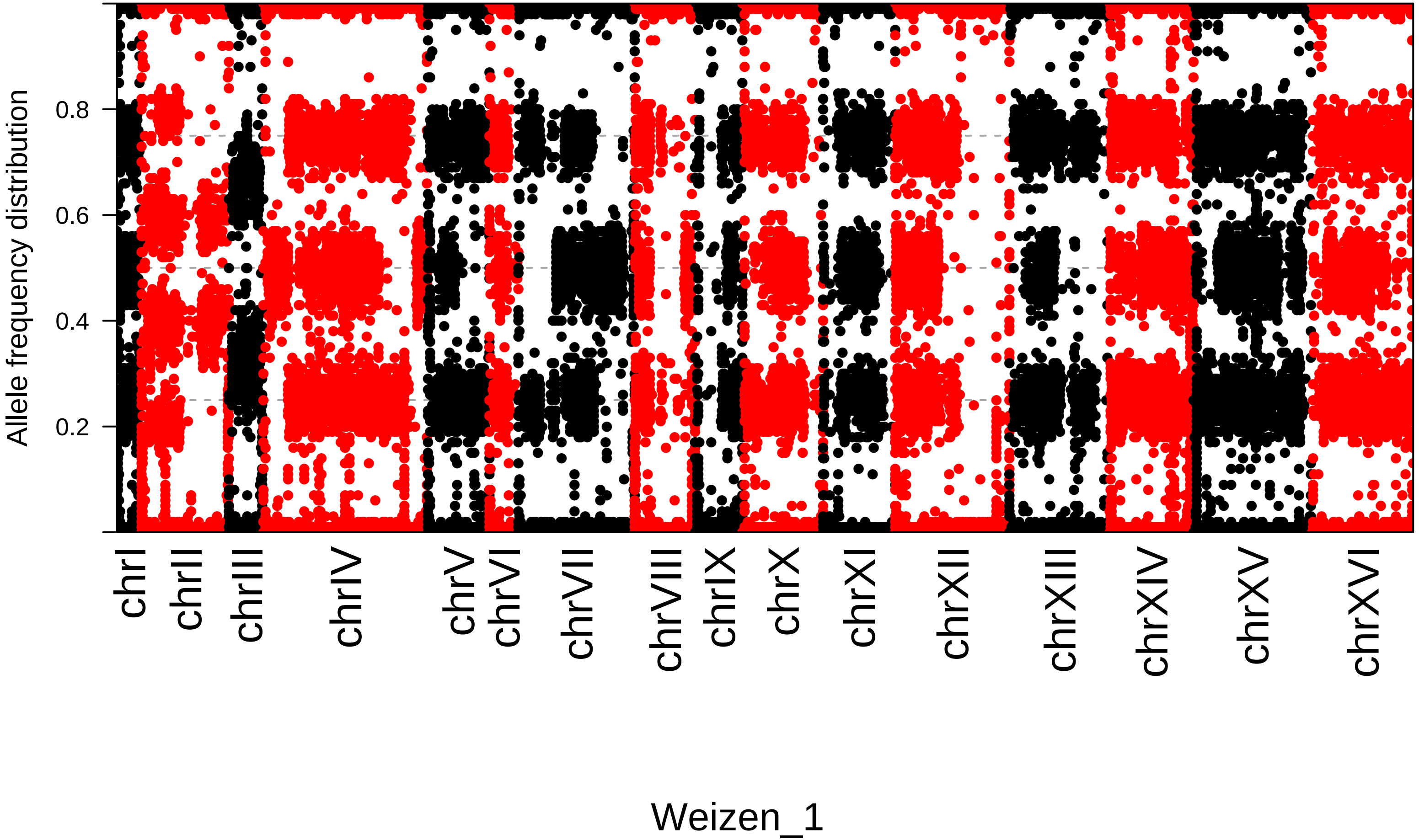
Allele frequency distribution
0.2
0.4
0.6
0.8
chrI
chrII
chrIII
chrIV
chrV
chrVI
chrVII
chrVIII
chrIX
chrX
chrXI
chrXII
chrXIII
chrXIV
chrXV
chrXVI
Weizen_1

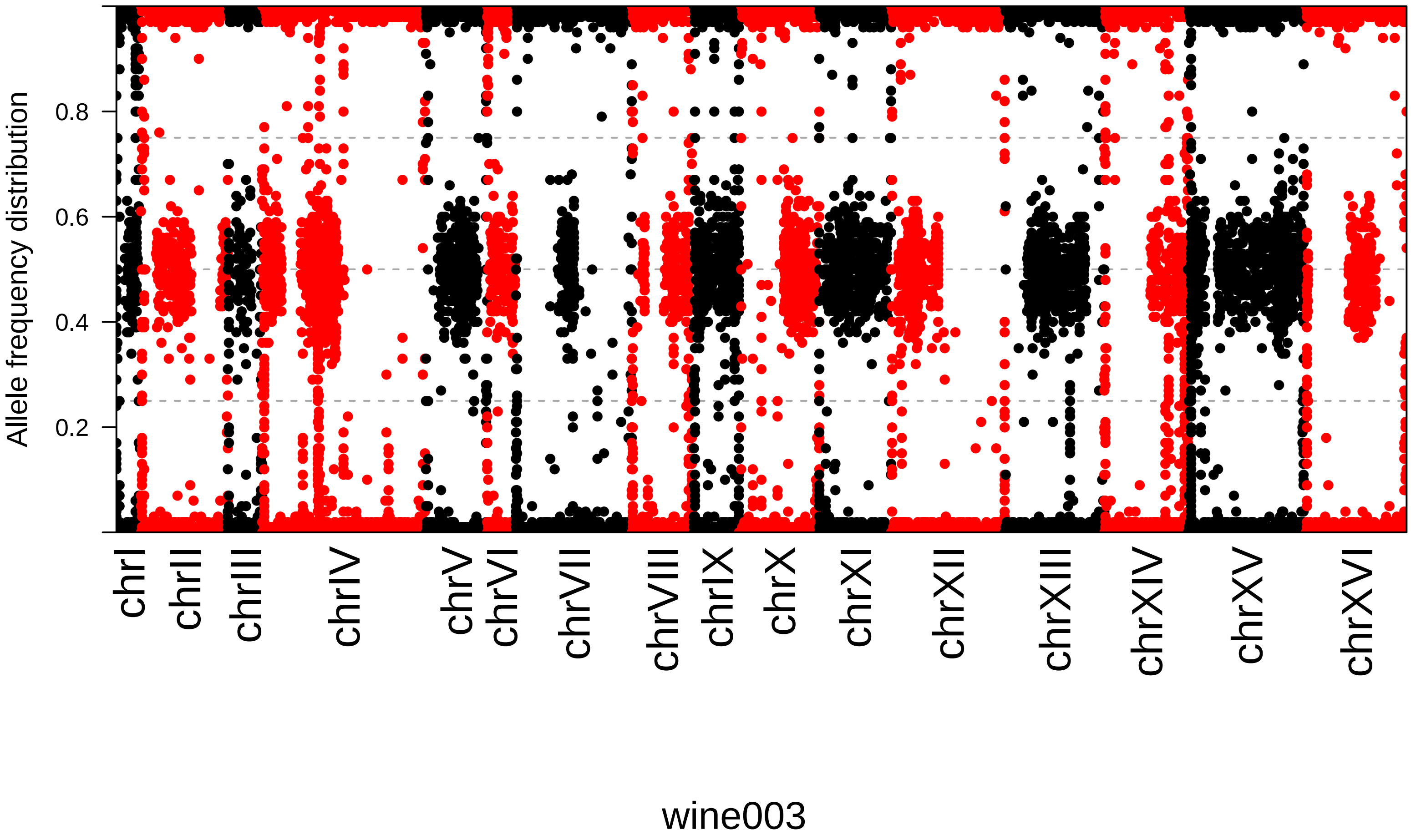

Allele frequency distribution
0.2
0.4
0.6
0.8
chrI
chrII
chrIII
chrIV
chrV
chrVI
chrVII
chrVIII
chrIX
chrX
chrXI
chrXII
chrXIII
chrXIV
chrXV
chrXVI
wine003

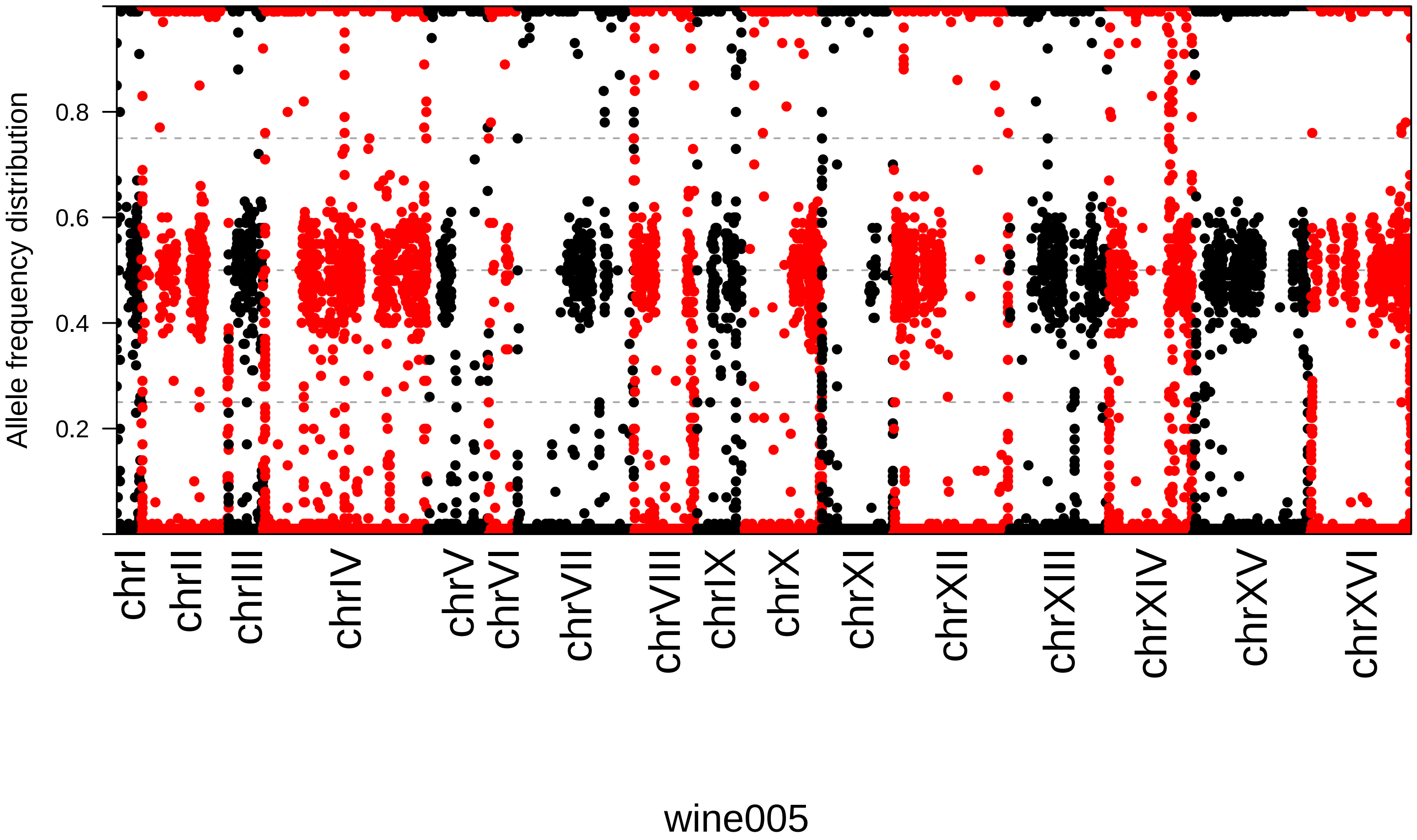
Allele frequency distribution
0.2
0.4
0.6
0.8
chrI
chrII
chrIII
chrIV
chrV
chrVI
chrVII
chrVIII
chrIX
chrX
chrXI
chrXII
chrXIII
chrXIV
chrXV
chrXVI
wine005
