## Supplementary figures and images for "European Farmhouse Brewing Yeasts Form a Distinct Genetic Group"

### Supplementary Figure S4

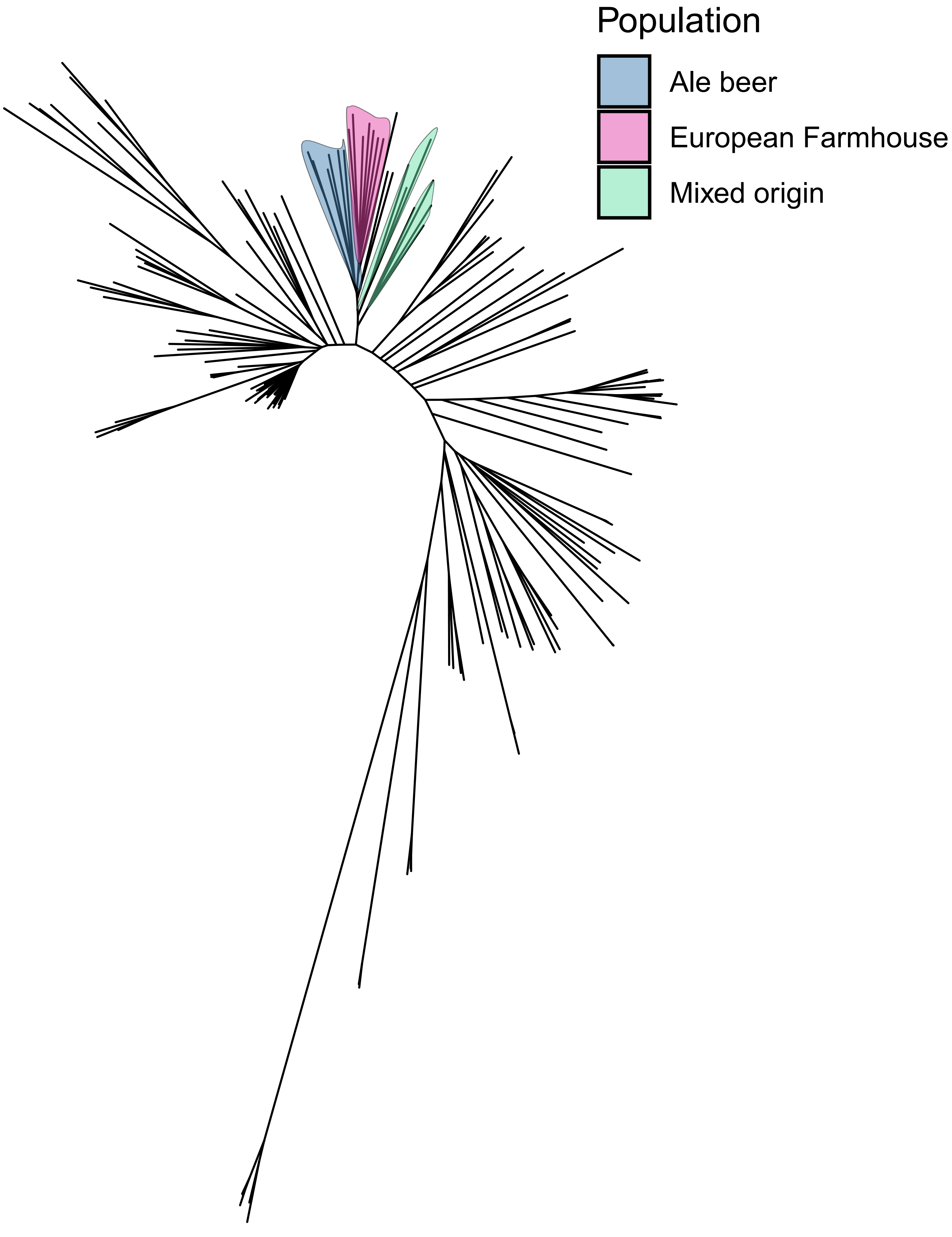

### Supplementary Figure S5

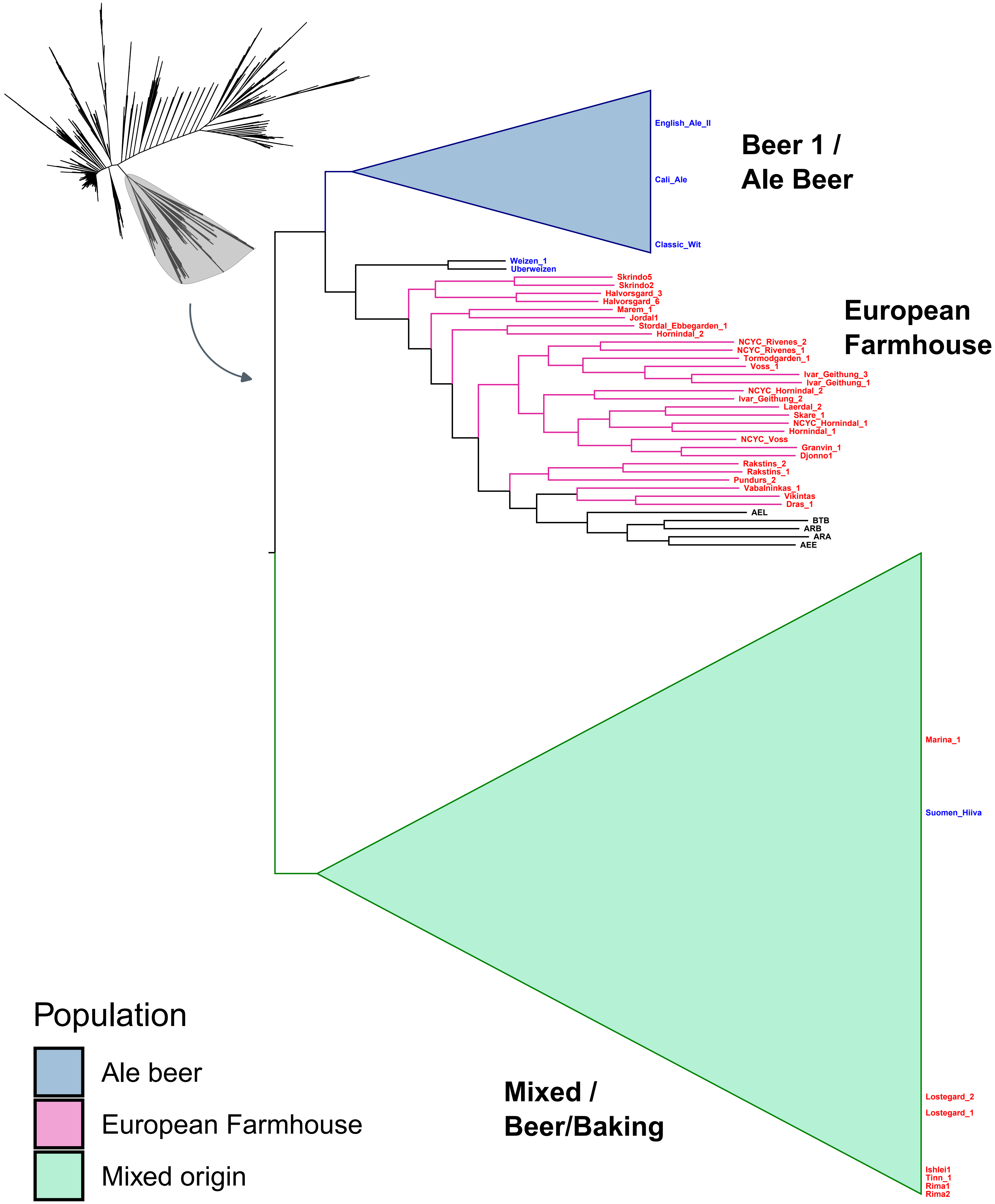

### Supplementary Figure S6

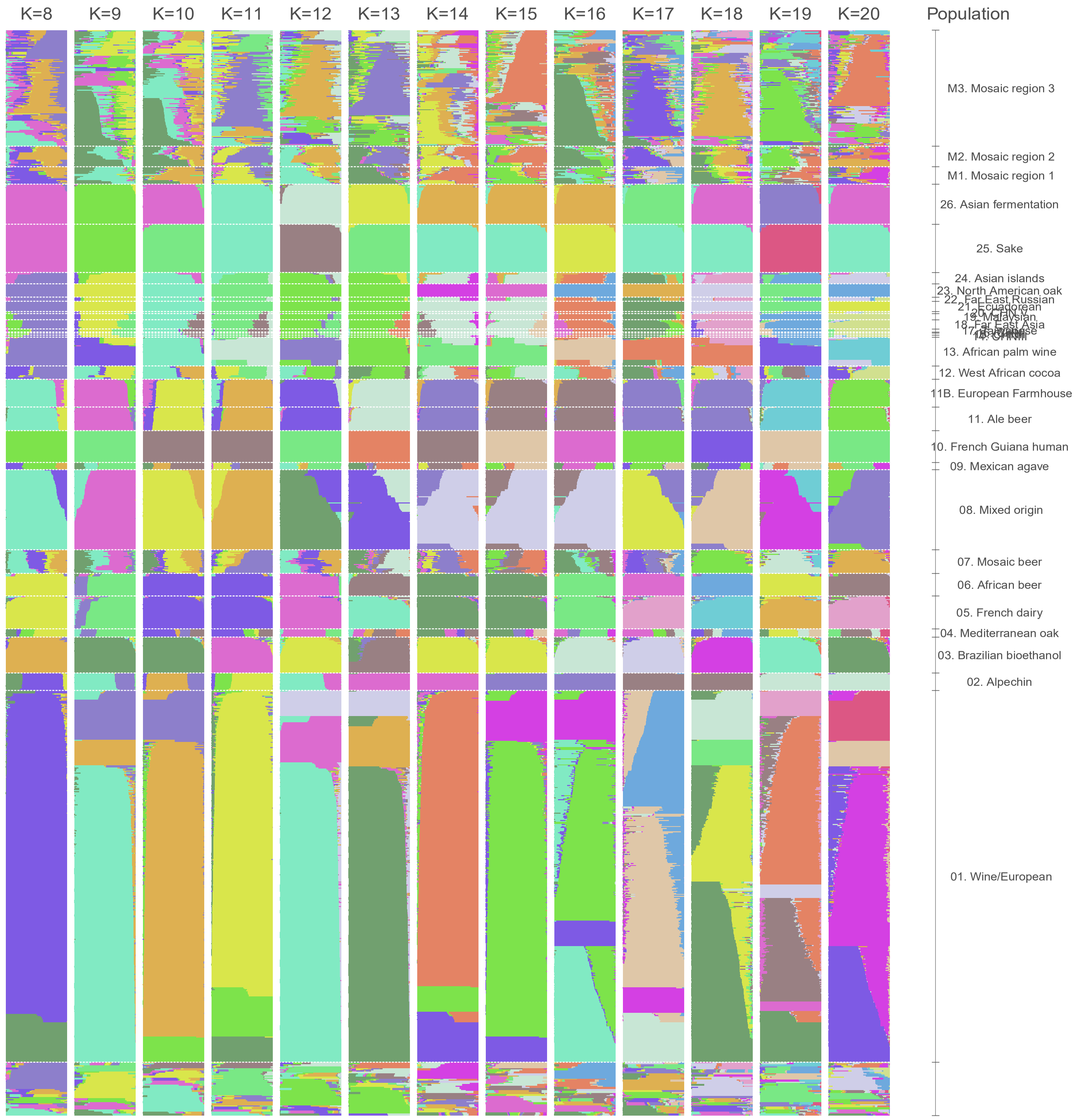

### Supplementary Figure S7

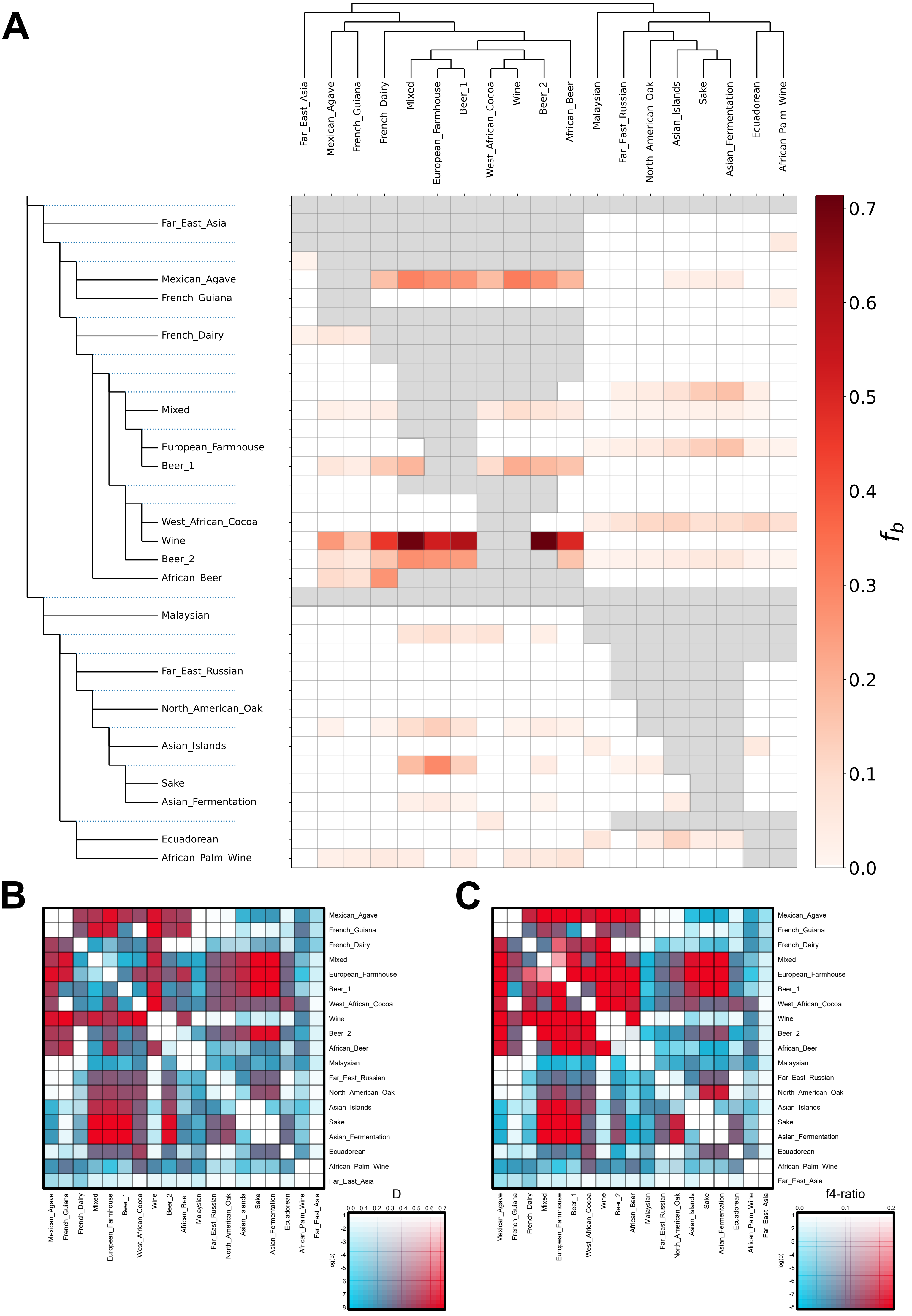

### Supplementary Figure S8

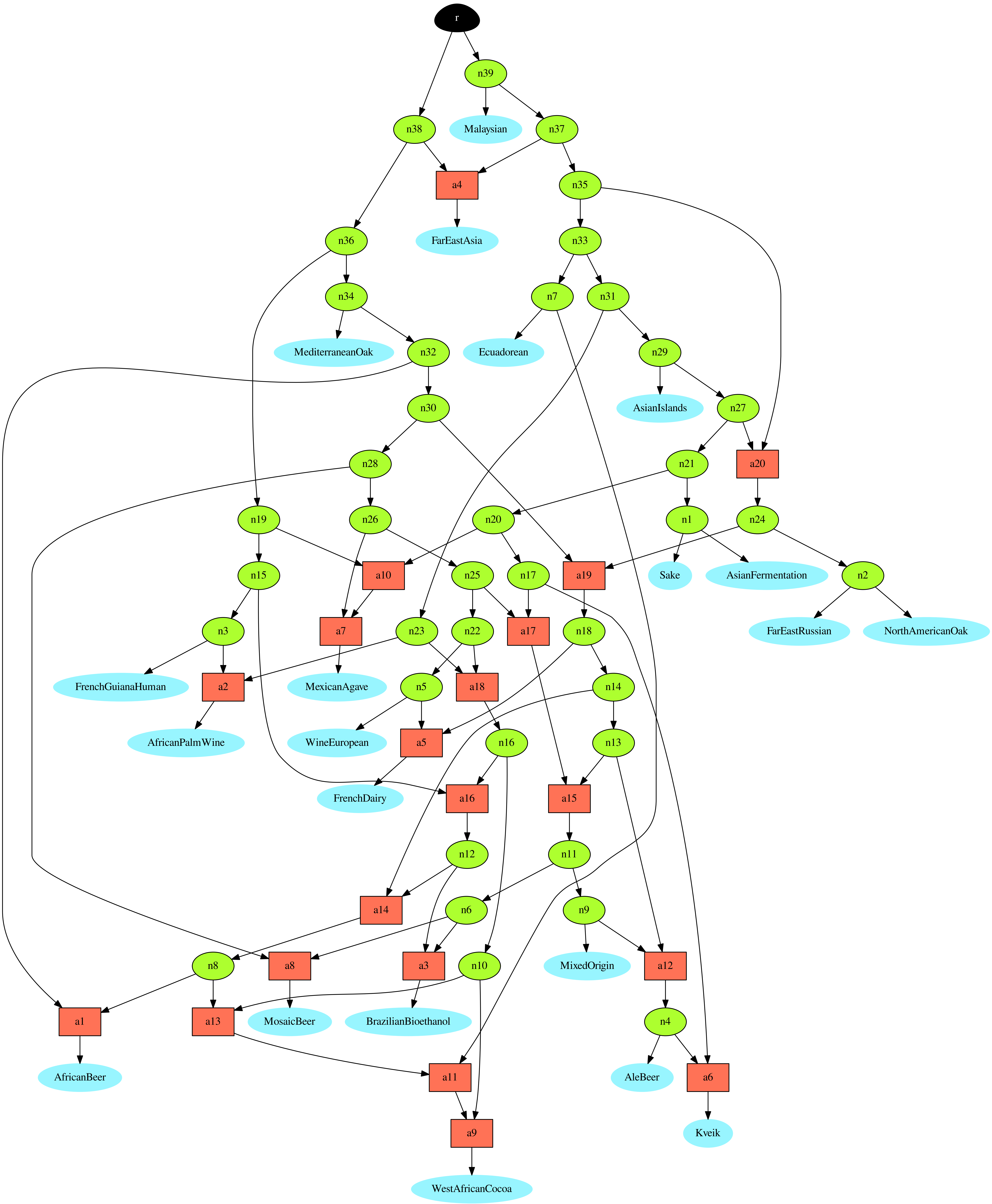

### Supplementary Figure S9

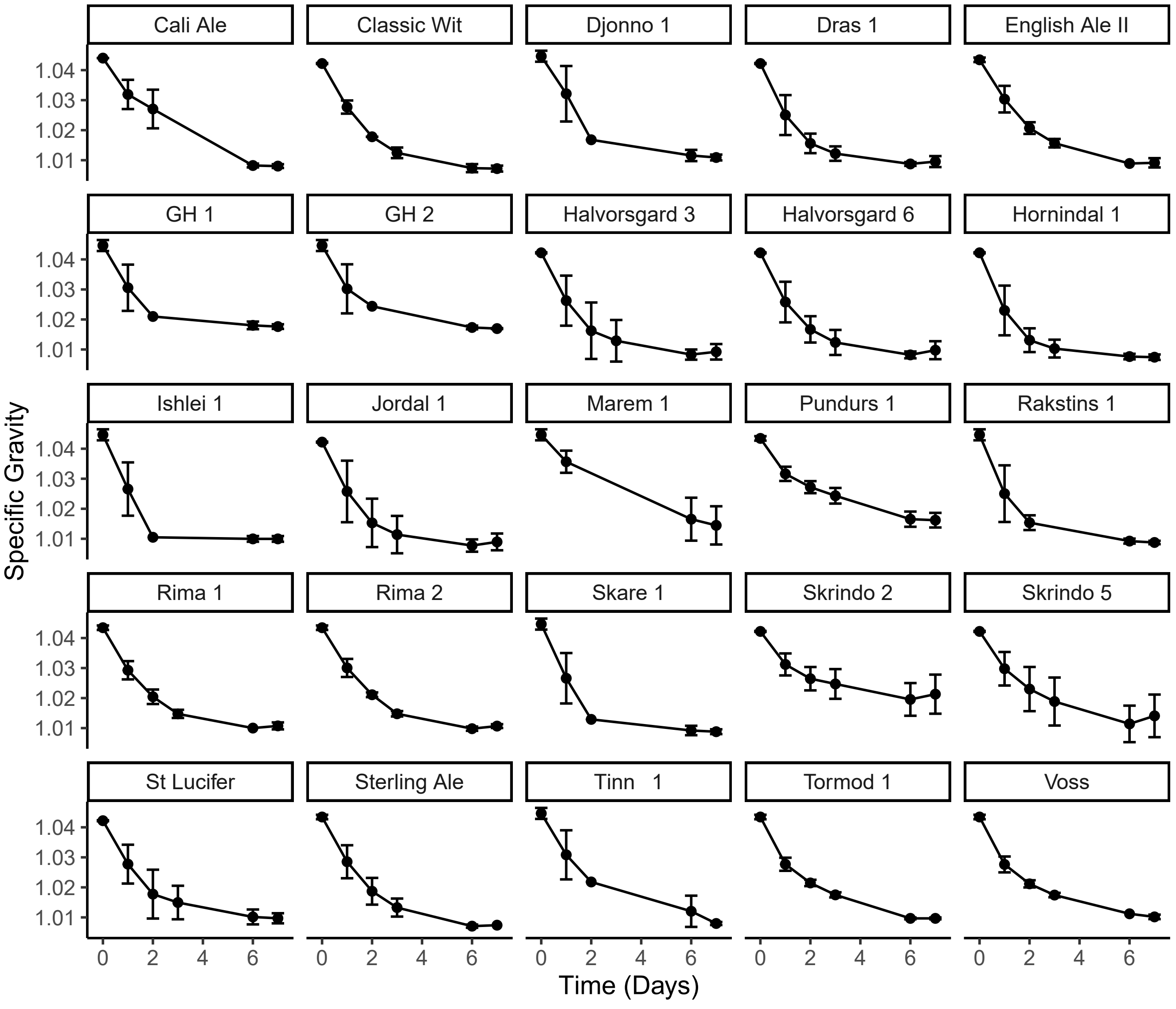

### Supplementary Figure S10

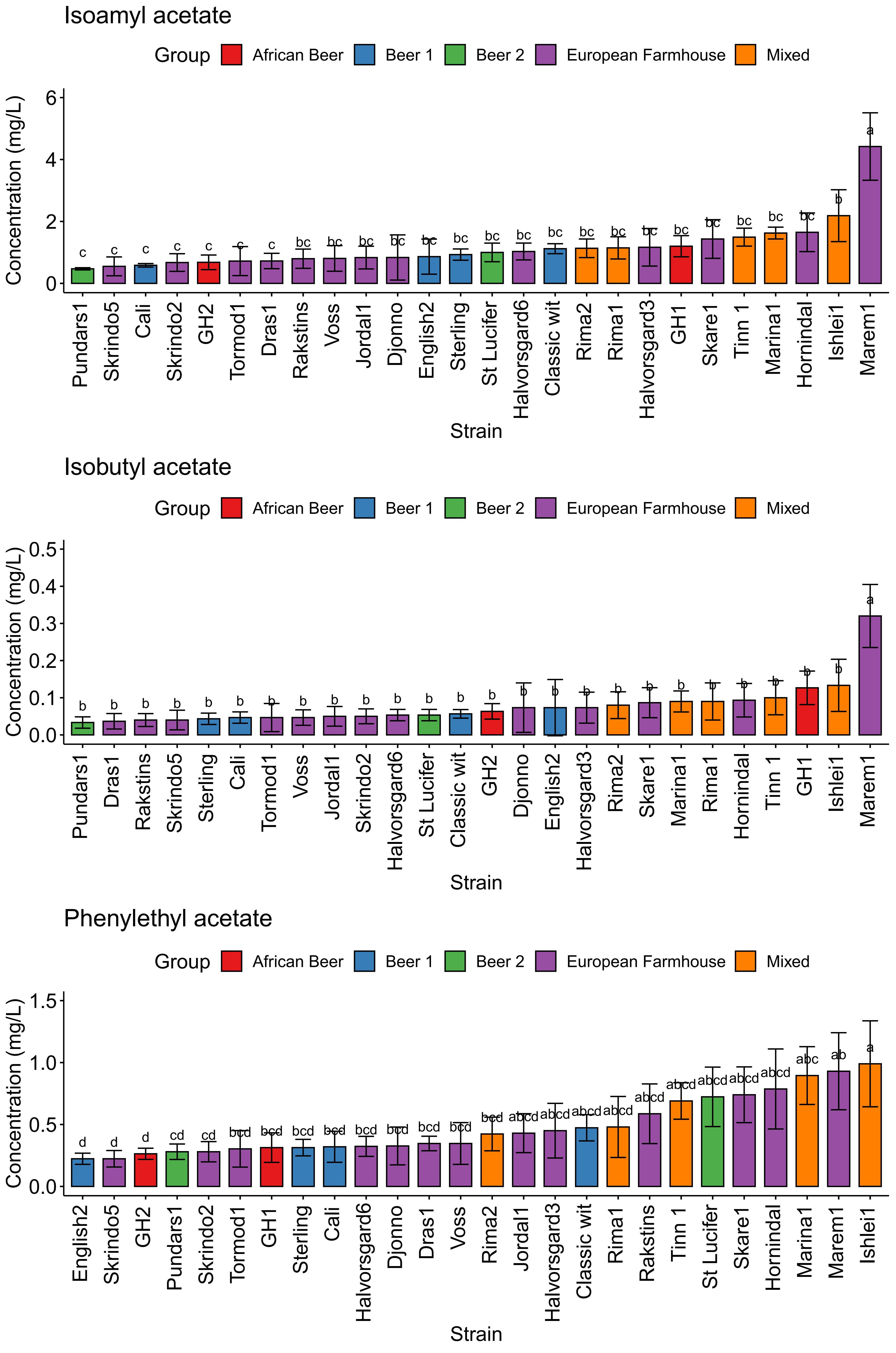

### Supplementary Figure S11

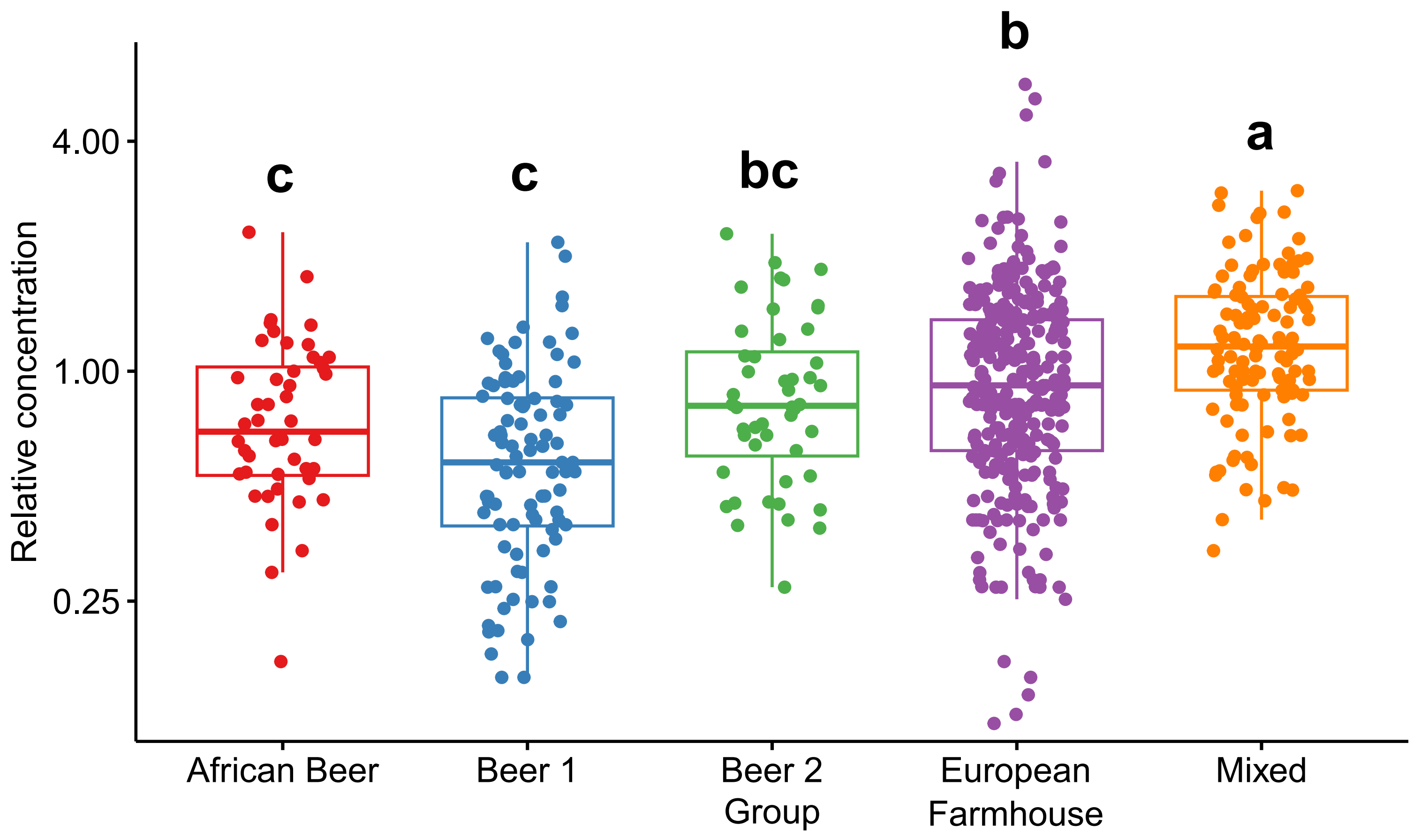

### Supplementary Figure S12

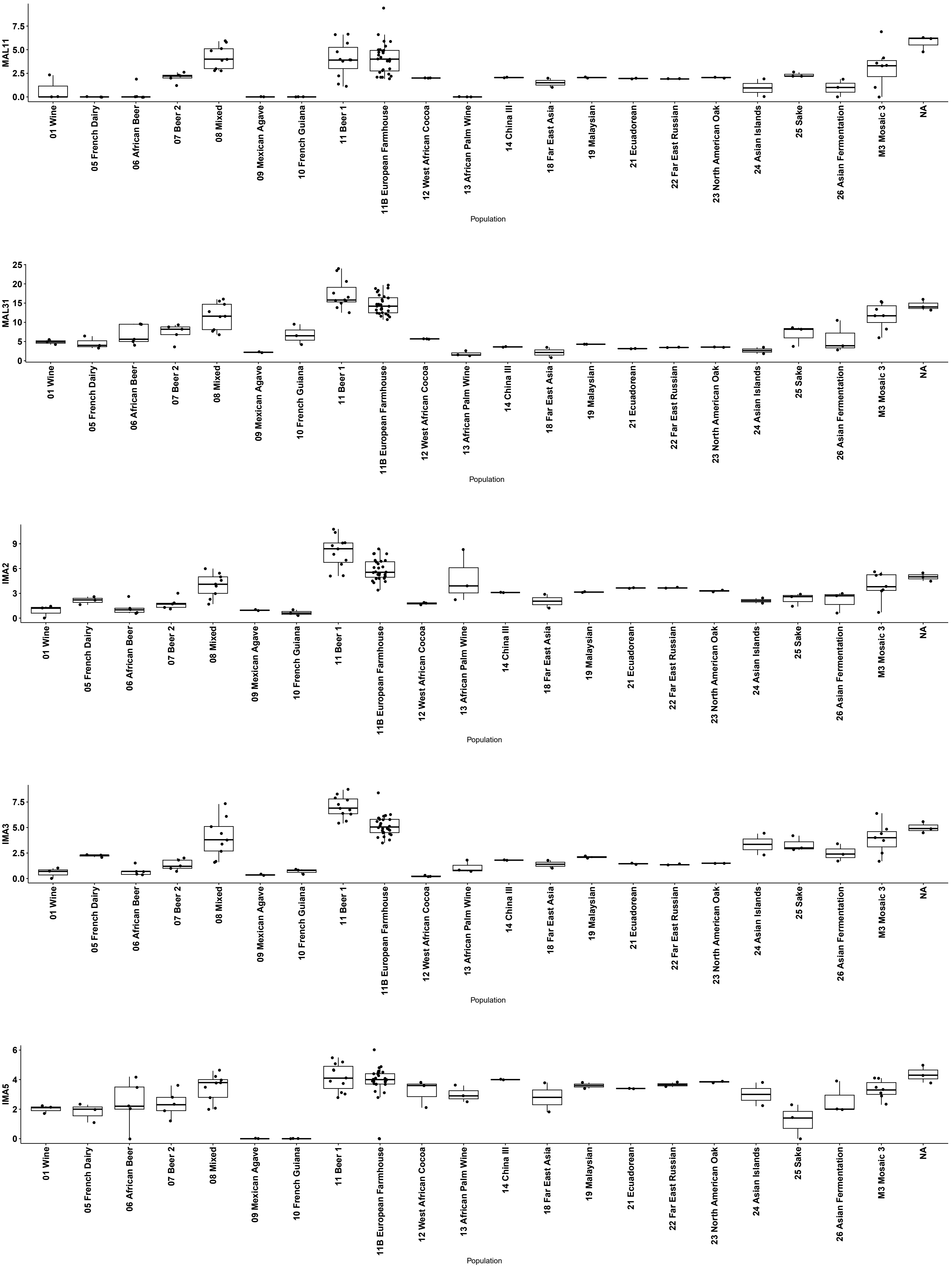

### Supplementary Figure S13

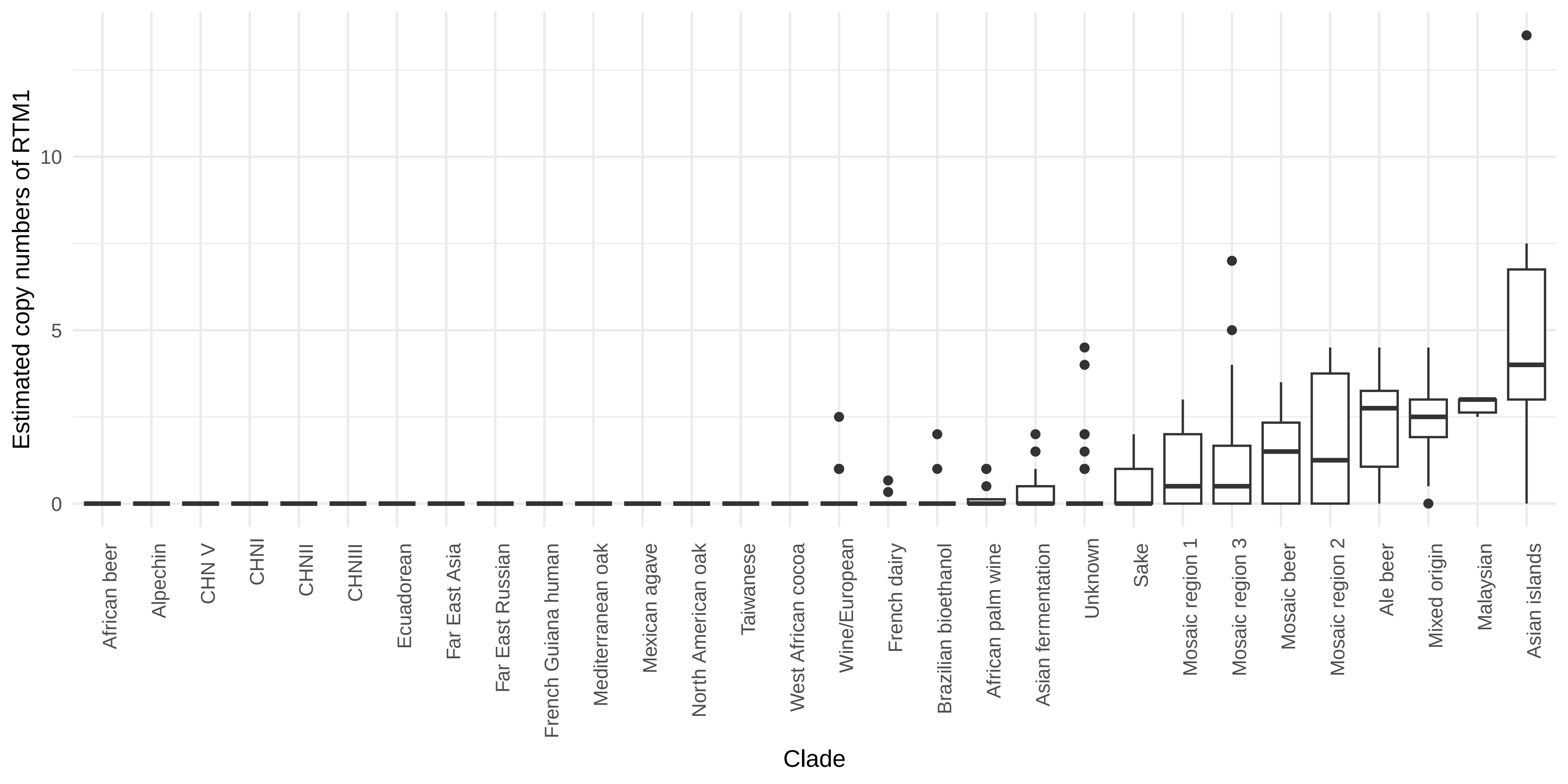

### Supplementary Figure S14

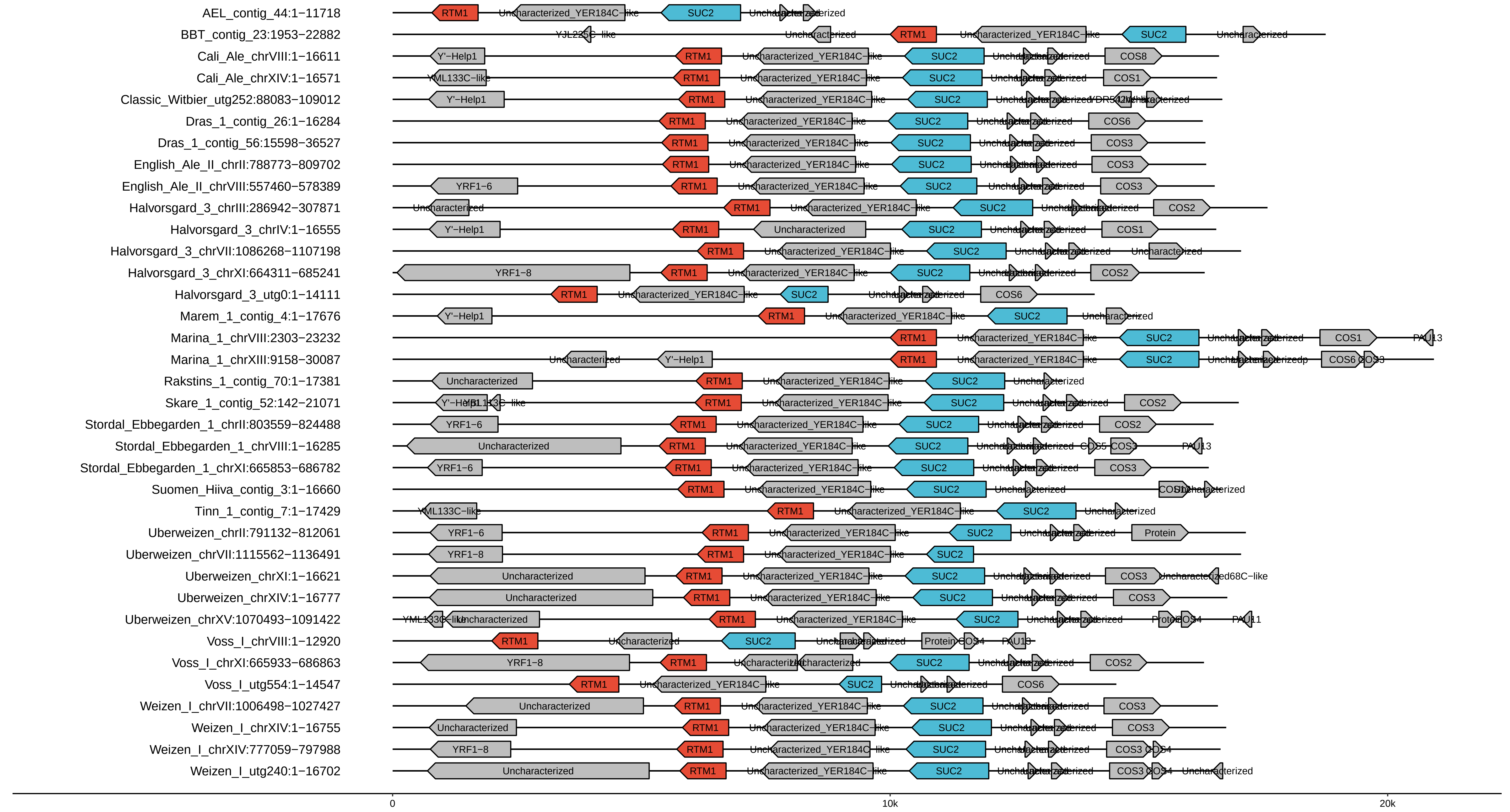
